# Separability of antibacterial and membranolytic activity in the human host defense peptide LL-37

**DOI:** 10.1101/2025.05.22.655608

**Authors:** John S. Albin, Dimuthu A. Vithanage, Wayne Vuong, Corey Johnson, Lael M. Yonker, Bradley L. Pentelute

## Abstract

The human *CAMP* gene product LL-37 is thought to exert direct antimicrobial activity against gram-negative bacteria via membrane disruption. In the course of structure-activity relationship studies of LL-37 aimed at developing peptidomimetic antibiotics, however, we incidentally noted mutations in LL-37 that globally inhibit membrane disruption in both mammalian and gram-negative bacterial cells. Despite their diminished capacity for membranolysis, these variants retained full antibacterial activity against gram-negative bacteria. While testing LL-37 and derivatives thereof against clinical isolates of *Pseudomonas aeruginosa* from patients with cystic fibrosis, we further noted unusually high rates of elevated minimum inhibitory concentrations for LL-37. Further evaluation of these clinical isolates revealed that they are fully permeabilized by LL-37 without being killed. Thus, we have identified variants of LL-37 that kill gram-negative bacteria without permeabilizing, and gram-negative bacteria that are permeabilized by LL-37 without being killed. This may suggest the existence of one or more mechanisms other than membrane disruption by which LL-37 can kill gram-negative bacteria, which may open up new avenues for antibiotic development based on the naturally evolved, non-membrane targets of human host defense peptides.

## Introduction

Most host defense peptides (HDPs) are thought to exert their direct antimicrobial activities against gram-negative bacteria through the disruption of bacterial membranes (1–3). A classic example this type of HDP is the human cathelicidin, LL-37, which is the most extensively studied of the human HDPs. Initially expressed as a 16 kDa proprotein, this *CAMP* gene product undergoes proteolytic processing following neutrophil degranulation to release an antimicrobial 37-residue helical peptide starting with two Leu residues (4, 5) LL-37 and many other HDPs are thought to be kill bacterial through membranolysis (6, 7), which is protective against certain types of bacterial infections *in vivo* (8, 9). It has been previously noted, however, that the minimum inhibitory concentrations (MIC) of a given membranolytic HDP may not necessarily correlate with the concentration at which it achieves membranolysis(10). Moreover, it is important to note that the ability of HDPs to permeabilize a membrane does not necessarily mean that permeabilization is the mechanism by which an HDP kills bacteria. Rather, it is possible that permeabilization serves to deliver a given HDP to the proper cell compartment for downstream biological function, as can occur in certain cell-penetrating peptides(11).

Although much of the investigation of antimicrobial mechanisms of action among HDPs over the past several decades has focused on membrane interactions of the kind thought to be critical for LL-37 function, multiple lines of evidence have also demonstrated that HDPs may function through targets other than membranes (*e.g.*, (12, 13)). Among the best examples of this in nature are bacterial microcins such as microcin J25, which is thought to inhibit the bacterial RNA polymerase(14–16), and insect proline-rich peptides such pyrrhocoricin, which is thought to act on bacterial DnaK and the bacterial ribosome(17–19). Within humans, the α defensin HD- 5 appears to exert its activity against gram-negative bacteria through an as-yet undefined non-membranolytic mechanism(20), while HD-6 forms nanonet structures that entrap intestinal microbes(21). Moreover, synthetic derivatives of the porcine cathelicidin protegrin that have advanced to Phase III clinical trials act through membrane protein targets such as BamA (22).

The original goal of our study was to identify derivatives of LL-37 with diminished activity against mammalian cell membranes and to then use these detoxified derivatives as the basis for lead antimicrobials based on the LL-37 scaffold. In the course of identifying these mutants, however, we noted that detoxified derivatives of LL-37 lose activity against both mammalian and bacterial cell membranes while retaining full antimicrobial activity against gram-negative bacteria, contradicting the common conception of LL-37 as acting primarily through membrane disruption. On testing these mutant derivatives for comparison with wild-type in clinical strains of *Pseudomonas aeruginosa* from patients with cystic fibrosis, we further noted unexpectedly high rates of elevated MIC to both wild-type and derivative LL-37, contradicting the common conception of the high barriers to resistance posed by classic membranolytic antimicrobials such as LL-37 – at least in the context of chronic infection. Surprisingly, these variants that avoided killing by LL-37 remained fully susceptible to permeabilization by LL-37, demonstrating that permeabilization alone is unlikely to be fatal in these cases. Noting both LL-37 variants that kill gram-negative bacteria without permeabilizing and gram-negative bacteria that are killed by LL-37 without being permeabilized, our data suggest that LL-37 may possess one or more mechanisms for killing gram-negative bacteria other than membranolysis.

## Results

### LL-37 hemolytic and antimicrobial functions are separable

Prior investigations into LL-37 structure-activity relationships have focused primarily on the study of LL-37 fragments(23). Though commonly deployed and often useful, the study of such fragments risks the loss of structural context found in the intact peptide. We therefore focused our efforts on the mutagenesis of structurally distinct surfaces within full-length LL-37. That is, in much the same way that one may mutagenize a particular surface in a larger protein to disrupt a particular function, we hypothesized that the same principle of structurally contiguous regions imparting function might apply to small but structured proteins like LL-37 and other HDPs.

To do this, we started by visually inspecting a prior NMR structure of LL-37 and identifying structurally contiguous residues that might cooperate in a given function(24). We then synthesized a series of LL-37 mutants in which every residue in each individual region was changed to Ala and used these mutants as the basis for initial studies; a schematic summary of this strategy is shown in **Figure 1A**. Importantly, evaluation by circular dichroism (CD) spectroscopy showed that all of these mutants retain characteristic minima at 208 and 222 nm suggestive of retained helical structure, and estimates of secondary structure content as calculated by BeStSel(25) indicated 100% helical content throughout. It is thus likely that these mutants retain helical structure comparable to wild-type LL-37, allowing for the isolation of sidechain effects through grouped Ala mutagenesis. **Supplementary Figure 1** shows the CD spectra for all mutants as well as wild-type LL-37.

**Figure 1.**
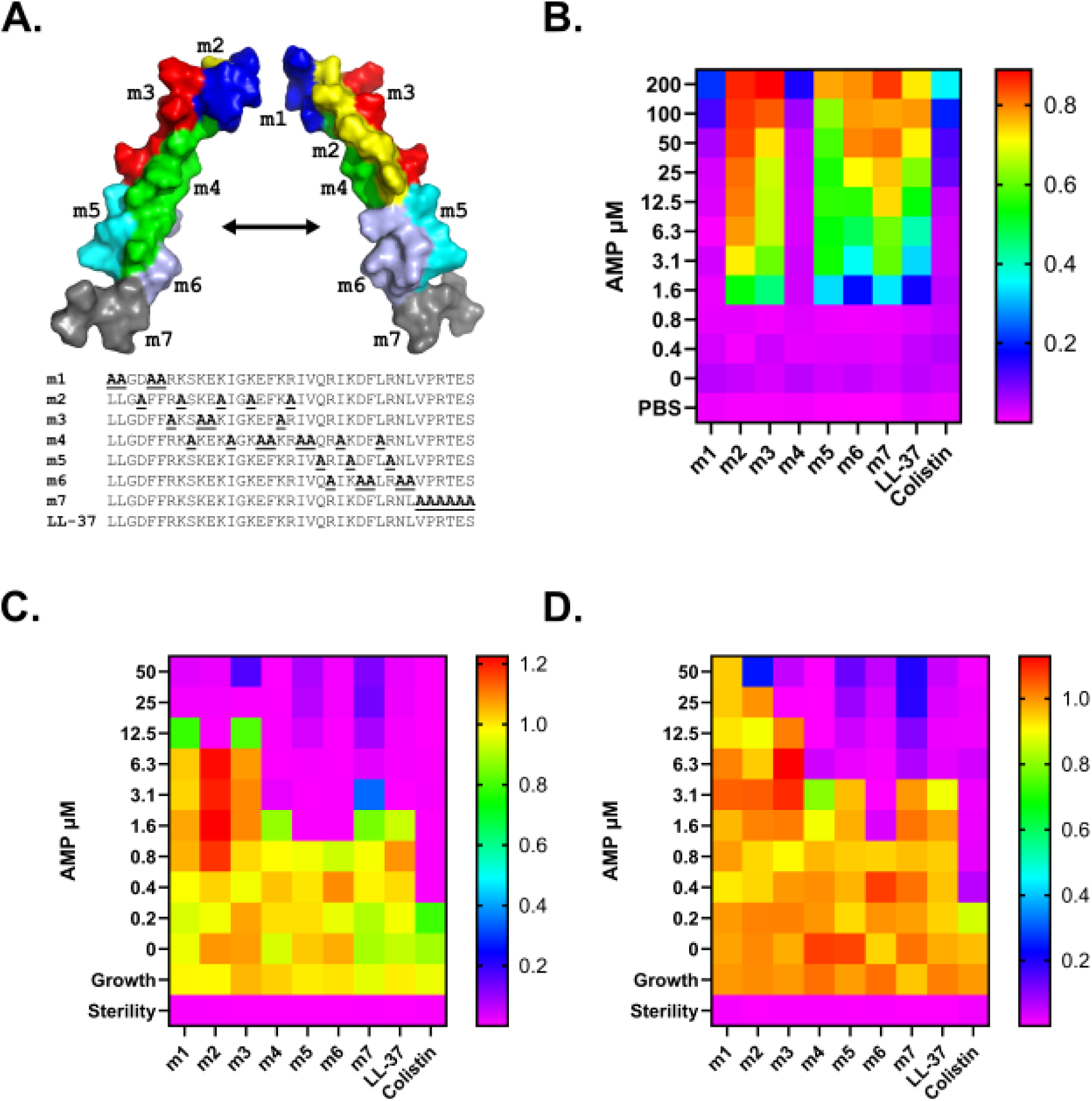
Mutations in region m4 separate LL-37 hemolytic and antimicrobial functions. **A.** Overview of mutagenesis strategy. Each mutant utilized has all residues within the putative structural surface changed to Ala, as shown in the associated sequences compared with wild-type LL-37. Regions are shown on a prior NRM structure 2K6O. **B.** Hemolysis assay quantifying the activity of each peptide relative to a 100% hemolysis control after 18 hours of incubation at 37 °C in the presence of different concentrations of each peptide. Warm colors indicate greater hemolysis as evidenced by absorbance at 414 nm for free hemoglobin in well supernatants; cool colors indicate low or no hemolysis. **C-D.** Antimicrobial susceptibility testing of *E. coli* (**C**) and *P. aeruginosa* (**D**), which shows the OD600 after 18 hours of incubation at 37 °C in the presence of different concentrations of each peptide. Warm colors indicate growth; cool colors indicate no or low growth. Data shown are the mean of three independent experiments.

Using hemolytic activity against sheep blood as a surrogate for activity against mammalian cell membranes, we first assessed the hemolytic activity of this set of mutants. As shown in **Figure 1B**, m1 and m4 within this group demonstrated substantial reductions in hemolytic activity compared with wild-type, making them comparable in this measure of toxicity to the control antibiotic colistin. Further evaluation of the antimicrobial activity of this complement of mutants via broth dilution antimicrobial susceptibility testing, however, showed that only m4 retained activity, while m1 as well as m2 and m3 lost antimicrobial activity against both *Escherichia coli* and *P. aeruginosa* (**Figure 1C-D**). **Supplementary Figure 2** shows the hemolysis and antimicrobial susceptibility endpoint assays from **Figure 1** in histogram format with error bars. **Supplementary Figures 3-11** (hemolysis) and **12-20** (antimicrobial susceptibility) show the 18-hour kinetic curves associated with each endpoint measurement. Mutations in region m4 thus ablate LL-37 hemolytic activity without affecting antimicrobial activity, demonstrating separability of these two functions.

### Separation of hemolytic and antibacterial functions in LL-37 is conferred primarily by a combination of two amino acids

LL-37 mutant m4 consists of eight residue changes in native LL-37 to Ala. To better define those responsible for separation of function between hemolytic and antibacterial functions in LL-37, we made a series of overlapping mutants as shown in **Figure 2A** and subjected these to testing for hemolytic and antibacterial activities in a fashion similar to **Figure 1**. As shown in **Figure 2B**, disruption of hemolytic activity was initially localized to the four residues of the m10 region – I20, V21, I24, and L28. Mutation of any one of those four residues to Ala as in m11-m14, however, did not ablate hemolytic activity. We noted, however, that only the m10 mutant among the overlapping mutants m8-m10 had diminished hemolytic activity, implicating the only mutations unique to m10 among these mutants – I24A and L28A. We also noted among size exclusion chromatography (SEC) experiments that the elution times and peak shapes associated with I20A (m14) and V21A (m13) were intermediate between those of LL-37 and m4, while those associated with I24A (m12) and L28A (m11) eluted with a retention time similar to m4 (see **Figure 3** below). We therefore prepared mutant m15, which combines I24A and L28A and demonstrates diminished hemolytic activity comparable to the m4 mutant (**Figure 2B**). Antimicrobial activity remained largely the same across mutants tested in this series of experiments as shown in **Figure 2C-D**. We did note, however, that most mutants had MICs roughly one dilution lower than LL-37, which may be expected in light of the findings presented in **Figure 3** below. As shown in **Supplementary Figure 21**, and similar to the data shown in **Supplementary Figure 2**, this series of mutants maintains helical content comparable to wild-type LL-37. **Supplementary Figure 22** provides a histogram depiction of the data in **Figure 2**. 18-hour kinetic curves associated with each endpoint measurement are provided in **Supplementary Figures 23-33** (hemolysis) and **34-44** (antimicrobial susceptibility testing).

**Figure 2.**
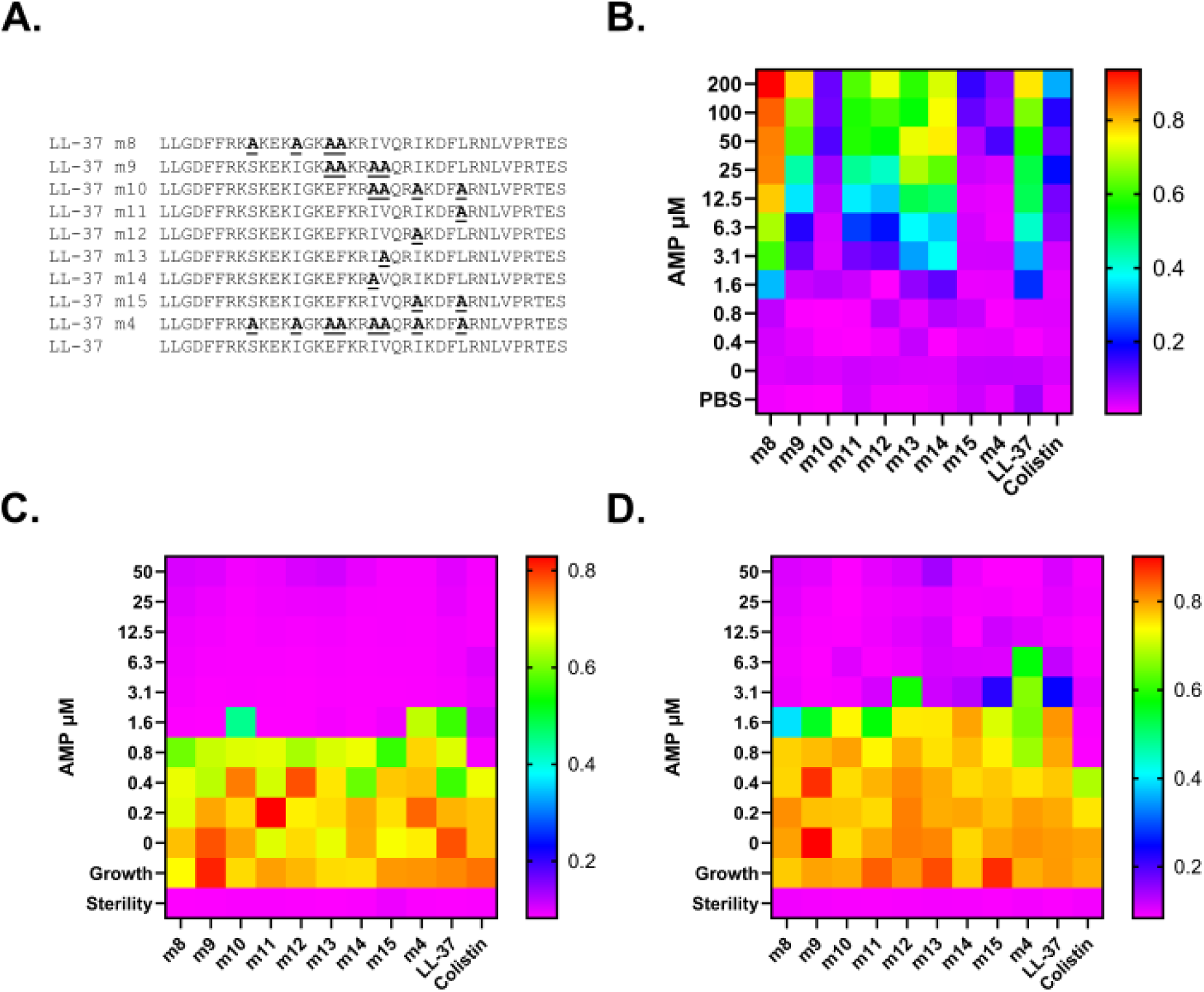
Separation of hemolytic and antimicrobial activities maps primarily to residues I24 and L28. **A.** Overlapping mutants were prepared as shown to localize the separation of function residues within LL-37 m4. **B.** Hemolysis assay quantifying the activity of each peptide relative to a freeze-thaw 100% hemolysis control. Warm colors indicate greater hemolysis as evidenced by absorbance at 414 nm for free hemoglobin in well supernatants; cool colors indicate low or no hemolysis. **C-D.** Antimicrobial susceptibility testing of *E. coli* (**C**) and *P. aeruginosa* (**D**), which shows the OD600 after 18 hours of incubation at 37 °C in the presence of different concentrations of each peptide. Warm colors indicate growth; cool colors indicate no or low growth.

**Figure 3.**
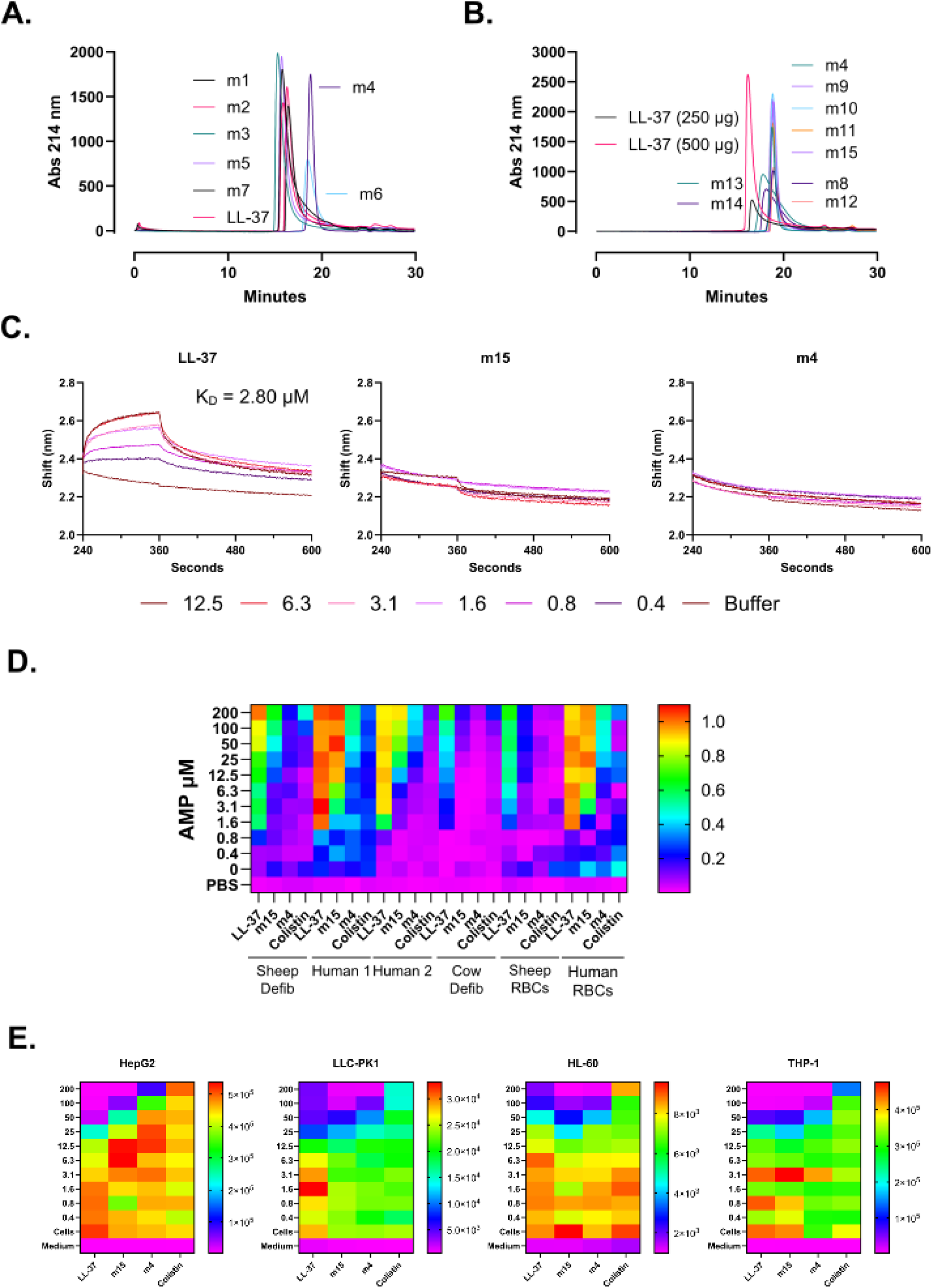
LL-37 mutants with decreased activity against mammalian cell membranes have a decreased propensity for oligomerization. **A.** Size exclusion chromatography (SEC) of LL-37 surface mutants as in Figure 1. **B.** SEC of overlapping mutants from Figure 2. **C.** Biolayer interferometry (BLI) showing the shifts associated with incubating the indicated analyte with a probe consisting of biotinylated LL-37. **D.** Extension of hemolysis assays to the sample types shown. Sheep and cow blood are defibrinated. Human blood is purchased whole blood from each of two donors. Sheep and human RBCs indicate isolated RBCs. **E.** Luminescence toxicity assay with each of the indicated cell types.

### LL-37 mutants with attenuated hemolytic activity have a diminished propensity for oligomerization

On comparison of our mutations of interest with prior crystal structures of LL-37, we noted that many of these were located in or adjacent to a core region of LL-37 previously associated with oligomerization(26, 27). We therefore hypothesized that mutants with diminished hemolytic activity might also have a diminished propensity for self-interaction. To test this hypothesis, we first subjected each of the surface mutants in **Figure 1** to size exclusion chromatography (SEC). As shown in **Figure 3A**, while most of these mutants migrated in a pattern similar to wild-type LL-37, m4 and m6, which also has residues in the region previously implicated in dimerization, had longer retention times and less tailing. By comparison to size controls, this pattern was suggestive of m4 migration as a monomer and LL-37 migration as a dimer under the conditions used (data not shown). This was further reinforced by evaluation of SEC traces among the extended list of mutants from **Figure 2**, where mutants with attenuated hemolytic activity such as m10 and m15 demonstrated peaks superimposable with that of m4.

We did note, however, that other mutants that retain hemolytic activity can also generate peaks comparable to m4 (*e.g.*, m9, m11, m8, and m12 in **Figure 3B**). Thus, while all observed mutants that lack hemolytic activity have SEC retention times suggestive of migration as monomers, not all mutants with similar retention times lose hemolytic activity.

To better quantify the propensity for self-interaction among LL-37 mutants, we prepared an N-biotinylated version of LL-37 and used this as a probe in biolayer interferometry (BLI) experiments with LL-37, m15, or m4 as analytes. As shown in **Figure 3C**, LL-37 demonstrated a characteristic on-off curve with an average KD across three independent experiments of 2.80 µM. By contrast, incubation of the LL-37 probe with m15 or m4 analytes yielded minimal shifts and no data sufficient for binding constant quantification, suggestive of a lack of self-interaction. Thus, decreased hemolytic activity in LL-37 is associated with a decreased propensity for self-interaction. **Supplementary Figure 45** shows the raw traces from three independent BLI experiments.

### Oligomerization-deficient LL-37 mutants demonstrate a range of activity against mammalian cell membranes

Within the standard model system that we use, the decreased activity of oligomerization-deficient LL-37 mutants toward sheep red blood cells (RBCs) was substantial. In **Figures 1** and **2**, for example, the ratio of the maximum peptide concentration with < 20% hemolysis for m4 relative to LL-37 is 125, slightly better than colistin at 62.5. To better define the activity spectrum of these mutants, we next carried out a series of hemolysis assays using a wider variety of sample types, including sheep and cow defibrinated blood, human whole blood, and isolated sheep and human RBCs. As shown in **Figure 3D**, detoxification effects varied by sample type, though the fold-improvement in hemolytic activity relative to wild-type LL-37 generally followed the trend of colistin > m4 > m15. This is quantified in **Supplementary Figure 46**, where the average fold-change in the highest peptide concentration yielding less < 20% hemolysis within each blood type relative to LL-37 was 51-fold for m15, 135-fold for m4, and 208-fold for colistin, albeit with substantial deviation due to differences among sample types. In particular, LL-37 derivatives showed less fold-improvement in human cell types relative to other species. It is unclear whether this is purely an effect of membrane interactions, or whether there are also naturally evolved LL-37-protein interaction in humans that may contribute. **Supplementary Figure 47** provides the data from **Figure 3D** in histogram format. Kinetic hemolysis curves for each condition from triplicate experiments are provided in **Supplementary Figures 48**-**71.**

To further assess whether the differential toxicity of different LL-37 mutants toward different mammalian cell membranes extends to different types of metabolically active cells, we carried out RealTime Glo (Promega) toxicity assays using HepG2, LLC-PK1, HL-60, and THP-1 cells. While some cell types such as HepG2 demonstrated a gradual decrease in toxicity by roughly one dilution each from LL-37 to m15 to m4 to colistin, the differences between oligomerization-deficient mutants and wild-type LL-37 were more muted in this context, though metabolically active cells were also less susceptible to LL-37 generally compared with RBCs, which may contribute to the lesser differences seen here (**Figure 3E**). Thus, residual toxicity of LL-37 derivatives in metabolically active cells may be mediated, in part, by interactions other than those occurring directly with membranes, which may be consistent with the reported interactions of LL-37 with protein receptors (*e.g.*, (28–32)). Histogram depictions of the data in **Figure 3E** are provided in **Supplementary Figure 72**. Kinetic curves leading into the endpoint measures in **Figure 3E** are shown in **Supplementary Figures 73-88**.

### Elevated LL-37 MICs are common among clinical isolates of *P. aeruginosa* from patients with cystic fibrosis

To better characterize the antimicrobial activity of LL-37 and oligomerization-deficient derivatives, we next tested LL-37 and m15 against a panel of clinical strains of *P. aeruginosa* from patients with cystic fibrosis using broth microdilution assays similar to those in **Figures 1-2**, in this case divided into four groups of eight strains each tested separately in duplicate for a total of 32 strains. As shown in **Figure 4A**, the modal MIC for LL-37 was 12.5 µM, and the MICs for LL-37 and m15 were within one dilution of each other in either direction in 30/32 (94%) strains, suggesting that LL-37 and m15 are generally comparable in their antimicrobial profiles against clinical strains of *P. aeruginosa* – albeit with a longer tail for m15 in the distributions shown. Although HDPs such as LL-37 are generally thought to pose high barriers to resistance due to their action on relatively immutable membrane targets(33–37), we found that 8/32 (25%) of strains in this collection had LL-37 MICs ≥ 50 µM, while 13/32 (41%) had m15 MICs ≥ 50 µM. Thus, elevated MICs are common among the clinical strains of *P. aeruginosa* tested in this study, approximating those observed for traditional antibiotics against *P. aeruginosa* among patients with cystic fibrosis, where chronic colonization and increasingly resistant infections are common(38). **Supplementary Figure 89** shows the endpoint assays contributing to the content of **Figure 4**, and **Supplementary Figures 90**-**121** show the kinetic curves associated with each endpoint divided by strain for each of the antimicrobials tested.

**Figure 4.**
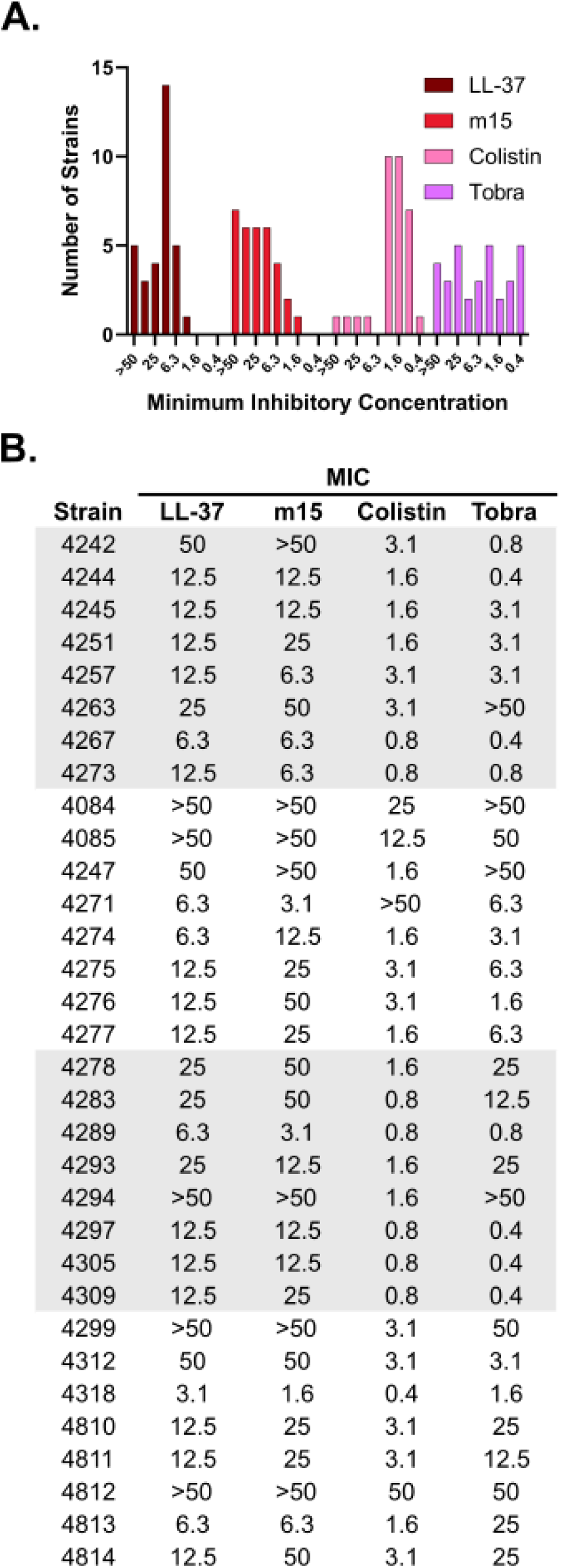
Elevated LL-37 and m15 MICs are common among clinical strains of. *P. aeruginosa* **from patients with cystic fibrosis. A.** Histogram distribution showing the frequency of each MIC among each of 32 strains for each of LL-37, m15, colistin, and tobramycin. **B.** Table showing relationships among MICs to each antimicrobial tested within each strain, grouped according to the order in which strains were tested. MICs at 48 hours of incubation are provided in µM.

### Oligomerization-deficient LL-37 mutants separate gram-negative cell killing from gram-negative cell permeabilization

The endpoint readout of the hemolysis assays described above depends on the release of free hemoglobin from RBCs in the presence of a given peptide. The attenuated release of free hemoglobin in the presence of oligomerization-deficient mutants thus implies that these mutants have a decreased capacity for the permeabilization of mammalian cell membranes.

Based on the generally accepted premise that LL-37 kills gram-negative bacteria via membrane disruption, we thought it unlikely that the same kind of diminished permeabilization activity would be observed with gram-negative bacterial cells. Rather, we hypothesized that the effect of diminished oligomerization must be to enhance cell type selectivity – decreasing activity against mammalian cell membranes while leaving activity against bacterial membranes intact.

To test this hypothesis, we subjected *E. coli* and *P. aeruginosa* strains to a dual membrane permeabilization assay in which N-Phenyl-1-Naphthylamine (NPN) serves as a marker of outer membrane disruption and propidium iodide (PI) serves as a marker of inner membrane disruption(39). As shown in **Figure 5A**, permeabilization of the inner membrane of *E. coli* by LL-37 is detectable down to 3.1 µM, which is also the MIC for LL-37 in this strain (**Figure 1C**). By contrast, permeabilization is markedly attenuated by approximately 16-fold for m4, with m15 staking out a middle position between these two (**Figure 5B-C**). Colistin included as a control appears to also effectively permeabilize *E. coli* under these conditions, albeit with delayed kinetics of uncertain significance (**Figure 5D**). This trend was also observed for outer membrane disruption under these conditions, though m4 permeabilization under these conditions was appreciable at concentrations as low as 12.5 µM as opposed to 50 µM for inner membrane permeabilization above (**Figure 5E-H**). Thus, despite the fact that LL-37 and m4 have the same MIC against *E. coli*, permeabilization of *E. coli* by m4 appears attenuated under the conditions used in our studies. Similar observations were made in *P. aeruginosa*, though permeabilization was globally more efficient and m4 was less attenuated by approximately four-fold in this context relative to LL-37 (*i.e.*, detectable permeabilization to 6.3 µM for m4 versus 1.6 µM for LL-37 in **Supplementary Figure 122**).

**Figure 5.**
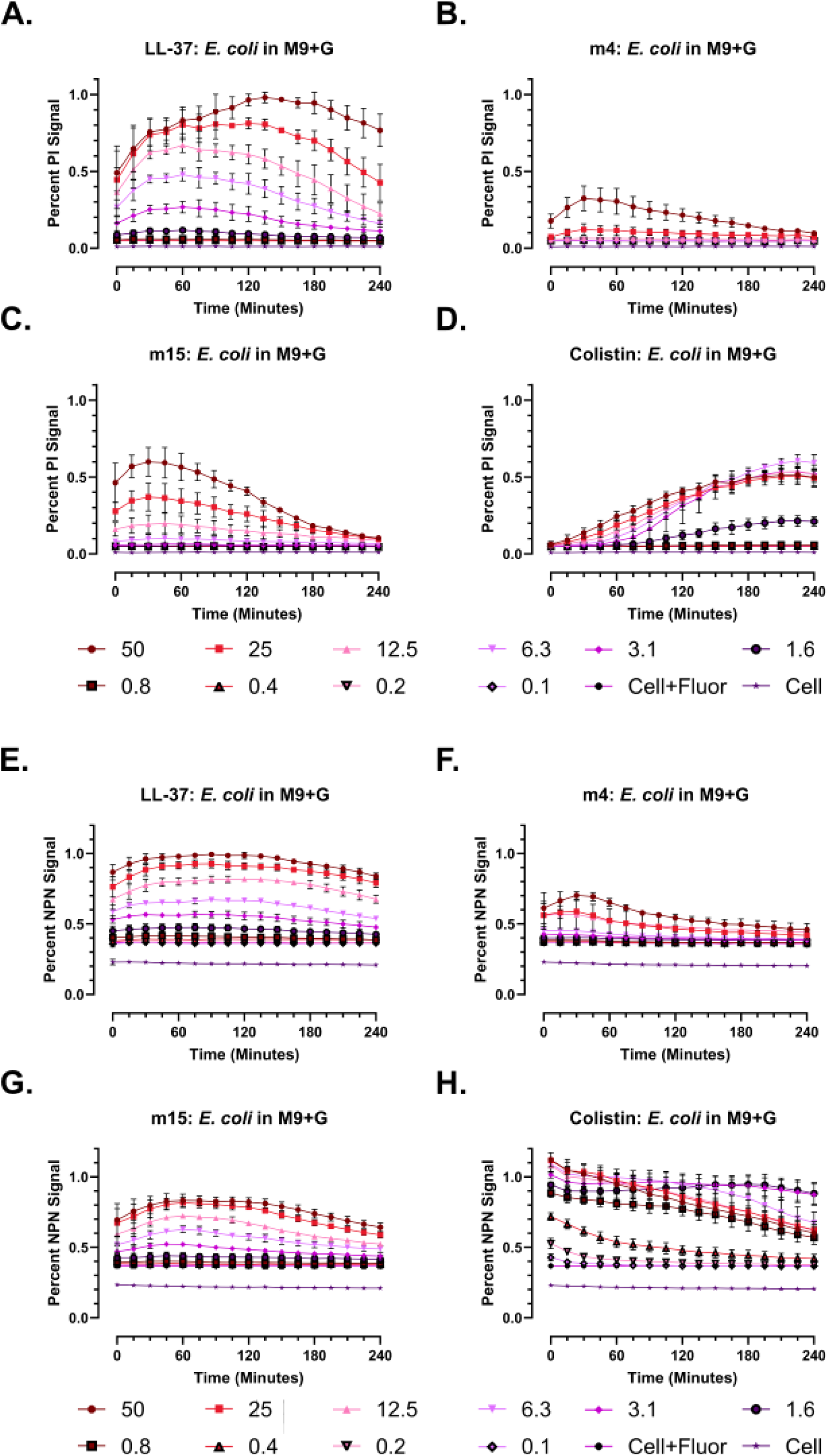
Oligomerization-deficient mutants demonstrate attenuated membrane permeabilization activity in gram-negative bacteria. A-D. Propidium iodide (PI) staining for inner membrane permeabilization over 4 hours in the presence of each peptide at the indicated concentrations given in µM. Quantification in these experiments proceeds by calculating a fold-change in PI fluorescent signal (Ex 535 / Em 617) over baseline within a condition (*e.g.*, *E. coli* ATCC 25922 in M9+glucose here), setting that fold-change to 100% in a given experiment (usually a high concentration LL-37 timepoint), and then expressing the remaining changes as a percentage of that high point. **E-H.** N-Phenyl-1-Naphthylamine (NPN) staining for outer membrane permeabilization over 4 hours in the presence of each peptide at the indicated concentration given in in µM. Quantification proceeds as for PI, except that the highest NPN signal (Ex 350 / Em 420) timepoint is set to 100%. Data throughout show the mean and standard deviation of three independent experiments in 50 µL total volume in black transparent-bottom 384-well plates incubated at 37 °C.

The experiments shown in **Figure 5** were completed in M9+glucose, which was chosen for its lack of fluorescent background. To determine the applicability of the above observations in different buffers, however, we also included PBS+glucose and unadjusted Mueller Hinton Broth (MHB) among our experimental conditions. Although the latter has a high fluorescent background, it is the medium in which the MIC assays used throughout were completed and is thus the most directly relevant to the functional phenotypes observed in **Figure 1**.

As shown in **Supplementary Figures 123**-**124**, permeabilization was globally somewhat more prominent with oligomerization-deficient mutants in PBS+glucose, indicating a degree of dependence on the specific buffer composition used in a given experimental setup. For example, the minimum detectable PI staining under these conditions in *E. coli* is around 0.8 µM for LL-37 versus 3.1 µM for m4, again with further narrowing of differences in *P. aeruginosa* to comparable levels of permeabilization between LL-37 and m4. In MHB, by contrast, m4 demonstrated appreciable permeabilization of *E. coli* only at 50 µM in MHB, consistent with the results observed in M9+glucose. There is thus likely at least a four-fold difference in permeabilization between m4 and LL-37 in MHB, though our ability to interpret anything beyond this level is limited by the background fluorescence in MHB (*e.g.*, 12.5 µM is the lowest level at which permeabilization is observed in MHB before signal falls into the background in **Supplementary Figure 125**). As in other conditions, phenotypes for LL-37 and oligomerization-deficient mutants were closer to each other in *P. aeruginosa* in MHB (**Supplementary Figure 126**), perhaps reflecting the relative permissiveness of the *P. aeruginosa* outer membrane for the passage of smaller hydrophilic molecules(40). Background fluorescence did not permit the quantification of outer membrane permeabilization in MHB (data not shown).

There are several degrees of permeabilization observed in these studies. At one extreme, the data in M9+glucose show markedly attenuated permeabilization activity in *E. coli* and to a lesser extent in *P. aeruginosa* for m4 relative to LL-37 – despite the fact that the MICs for LL-37 and m4 are the same in **Figures 1-2**. Attenuation is also seen in *E. coli* in MHB, but not with *P. aeruginosa*. Levels of permeabilization for LL-37 and m4 are closer to each other in PBS+glucose. Of note, all cells in these studies were initially grown in unadjusted MHB before being washed and diluted in the staining buffer of interest. Thus, at least at the start of each experiment, cells should be physiologically similar to those grown in MHB for susceptibility testing. Attempts to conduct susceptibility testing directly in M9+glucose have generally resulted in no growth of control cultures at a standard inoculum (data not shown). Within these limitations, the data obtained for *E. coli* in MHB in **Supplementary Figure 125** most closely resemble the **Figure 5** data in M9+glucose, and suggest a four-fold or greater decrease in permeabilization efficiency for m4 versus LL-37 in MHB. Because all of our antimicrobial susceptibility testing is conducted in MHB, it is likely that the diminished susceptibility to permeabilization that we see in **Supplementary Figure 125** (and in **Figure 5**) is reflective of what is happening during susceptibility testing in **Figures 1-2**, where m4 kills *E. coli* with the same efficiency as LL-37 despite its apparently attenuated membranolytic activity. Thus, although the level of permeabilization required to kill gram-negative bacteria remains undefined, there is a clear discrepancy between the permeabilization and killing activities of m4 and LL-37

### *P. aeruginosa* strains resistant to killing by LL-37 are efficiently permeabilized by LL-37

Although membrane permeabilization is generally thought to be the mechanism by which LL-37 kills gram-negative bacteria, the data in **Figure 5** may imply a mechanism of LL-37 killing of gram-negative bacteria other than permeabilization, particularly at the lower peptide concentrations used in *in vitro* susceptibility testing. To further investigate the potential for such a mechanism, we next asked the inverse question – if there are LL-37 variants that can kill gram-negative bacteria without permeabilizing, are there also gram-negative bacteria that can be permeabilized without being killed?

To address this, we chose a panel of seven strains from the collection of *P. aeruginosa* characterized in **Figure 4** that demonstrated the ability to grow in the presence of even high concentrations of LL-37 on the order of ≥ 50 µM. We then subjected these to permeabilization assays as in **Figure 5** for comparison with a control strain with an LL-37 MIC of 6.3 µM. As shown in **Figure 6**, these strains of *P. aeruginosa* were permeabilized by LL-37 with an efficiency similar to that observed in the control strain – despite the inability of LL-37 to efficiently kill these same strains in **Figure 4**. Similar results were observed for outer membrane permeabilization in **Supplementary Figure 127**. **Supplementary Figures 128**-**135** show the permeabilization patterns in each of these eight strains for LL-37, m4, m15, and colistin; with some exceptions, patterns are generally as demonstrated in **Figure 4** in M9+glucose, featuring prominent distinctions between LL-37 and m4. In summary, the strains of *P. aeruginosa* analyzed here are efficiently permeabilized by LL-37 without being killed. Because membrane disruption clearly occurs without being lethal in these cases, it is possible that even strains that are killed by LL-37 may not be killed by membrane disruption itself.

**Figure 6.**
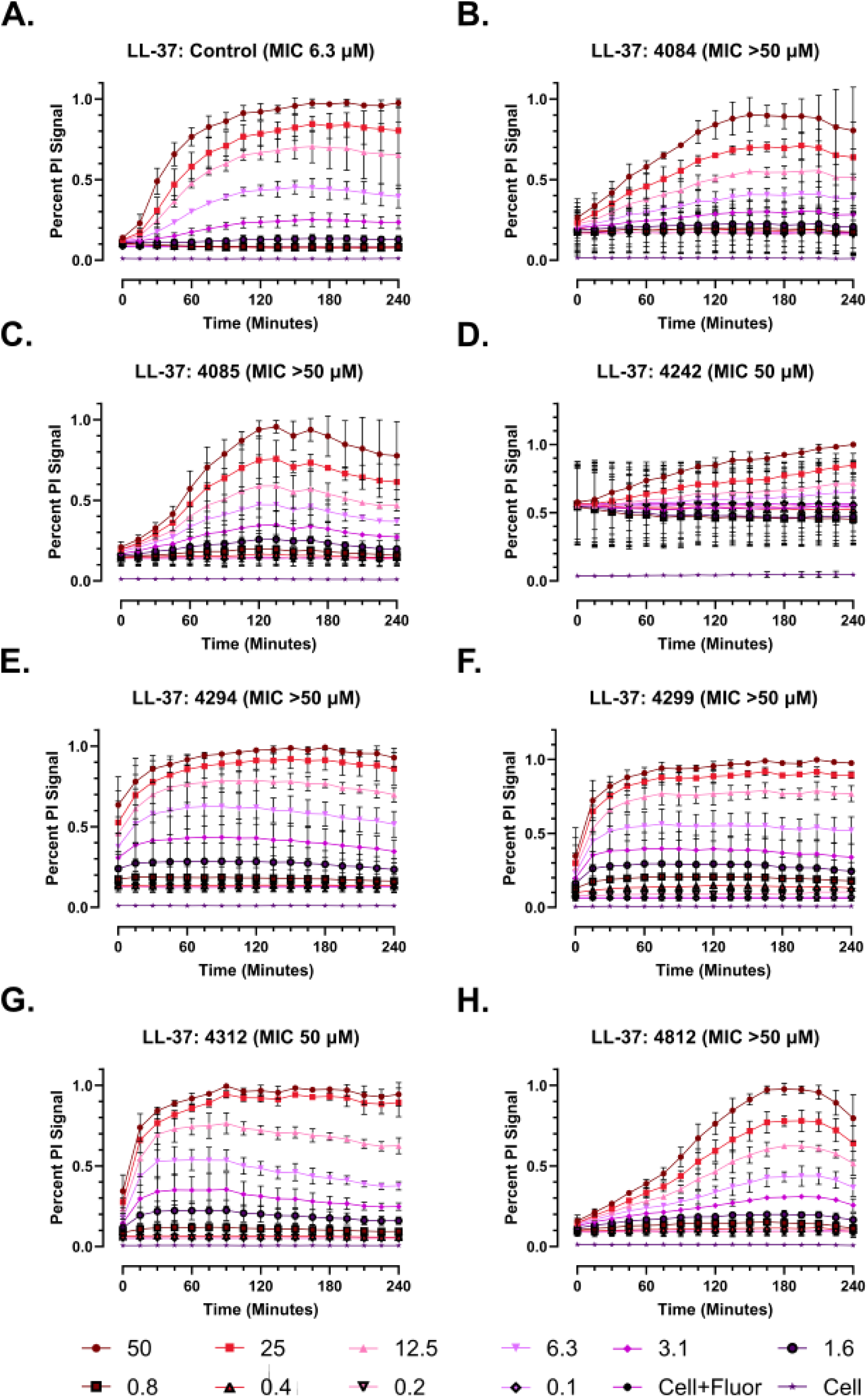
***P. aeruginosa* isolates with elevated LL-37 MICs are efficiently permeabilized by LL-37.** Propidium iodide (PI) staining for inner membrane permeabilization over 4 hours in the presence of LL-37. The strain and its MIC from Figure 4 for LL-37 are given in parentheses. **A.** Control *P. aeruginosa* strain, ATCC 27853. **B-I.** Clinical strains of *P. aeruginosa* from patients with cystic fibrosis. Peptide concentrations are given in µM.

## Discussion

The studies described here were initiated to identify determinants of mammalian cell toxicity and potency in the human HDP LL-37 with the goal of using this information for the development of lead antimicrobials based on the LL-37 template. On this front, we have identified oligomerization-deficient derivatives of LL-37 with improved *in vitro* therapeutic indices compared with wild-type LL-37, which may form the basis of further iterations on the LL-37 template for the development of novel antibiotics. Indeed, on the question of pure membranolytic activity against mammalian cell membranes, these mutants are approaching the levels of toxicity seen with colistin, which is included throughout as a comparator. This does not make them candidates for imminent clinical translation, but it does make them useful starting points for further development efforts starting from a less toxic baseline, as we have been working toward in other studies(41).

When starting this work, we had no particular interest in questions of LL-37 mechanisms of action against gram-negative bacteria, which we considered settled science. Indeed, there are few if any HDPs that better exemplify the characteristics of a stereotypical helical, amphipathic, membranolytic HDP than the extensively-studied human cathelicidin. In the course of our experiments, however, we encountered a series of three surprising findings that raised broader questions about our understanding of how LL-37 kills gram-negative bacteria.

In **Figure 4**, we present data to suggest that elevated LL-37 MICs are common in strains of *P. aeruginosa* from patients with cystic fibrosis, where colonization and chronic infection are common. This contradicts the general conception of LL-37 and other membranolytic peptides as posing high barriers to resistance by virtue of targeting relatively immutable membranes.

Rather, these data would suggest that it may be possible for *P. aeruginosa* to develop resistance to LL-37 over time, whether via direct adaptation to LL-37 itself or via cross-resistance in the setting of repeated antibiotic courses, though this has not been noted previously in smaller collection evaluations of which we are aware (42). We are currently working on better defining the frequency of elevated LL-37 MICs in clinical strains of *P. aeruginosa* from patients with cystic fibrosis, and it stands to reason that similar studies at other body sites, for example among gut *Enterobacteriaceae*, may also be informative.

In retrospect, this observation was perhaps predictable. When antibiotic resistance is encountered in nature, it typically occurs originally in the context of ecological conflict. When looking for resistance to human HDPs, then, it may stand to reason that it will be easier to find in the context in which ecological conflict involving an HDP might occur – namely, colonization and chronic infection. Regardless of the selective pressures to which *P. aeruginosa* was adapting in these strains, it is clear that they have not evolved to evade membrane permeabilization by LL- 37, which is as robust in these strains as it is in a sensitive control strain (**Figure 6**). If the selective pressures encountered *in vivo* caused these strains to develop elevated LL-37 MICs without developing resistance to permeabilization, then we would take this to mean that there is something else that LL-37 can do directly to a gram-negative cell that would be detrimental to pathogen survival.

Finally, in **Figure 6**, we show that *P. aeruginosa* isolates that are not killed even by high concentrations of LL-37 are nevertheless efficiently permeabilized by those same concentrations of LL-37. This may suggest that membrane disruption, even when present and fully active, is not necessarily fatal, which may be broadly applicable even among sensitive bacterial strains. Moreover, if membrane disruption by LL-37 is not fatal to these strains, this would seemingly imply that resistance is occurring through distinct, non-membrane target sites. These putative targets, in turn, may implicate one or more mechanisms by which LL-37 can kill gram-negative bacteria independent of direct membranolytic effects.

Based on the data presented here – variants of LL-37 that kill gram-negative bacteria without permeabilizing and variants of *P. aeruginosa* that are permeabilized by LL-37 without being killed – we hypothesize that there may be mechanism(s) by which LL-37 can kill gram-negative bacteria other than membrane disruption. That is, permeabilization certainly occurs and is likely relevant, at a minimum, for raising the resistance barrier and controlling high bacterial inocula, but it may not be the only way that LL-37 has evolved to kill gram-negative bacteria. It is possible, for example, that local concentrations of LL-37 at sites of infection with many neutrophils emptying the contents of their granules may be sufficiently high that membranolysis is a dominant mechanism of action. When concentrations are low, however, it be that LL-37 has at least one additional mechanism of action by which to kill gram-negative bacteria, and perhaps others to be discovered in its sequence space.

We would propose that even intact membranolytic activity in a given peptide does not automatically imply that this is its mechanism of bacterial cell killing. It is possible that one driver for the evolution of permeabilization activity among HDPs, for example, is self-promoted transport across membranes to a defined target rather than membranolytic killing. It is also difficult to distinguish membrane permeabilization as a downstream effect of cell death from membrane permeabilization as a consequence of direct permeabilization, though kinetics may assist in this. It may be worthwhile to reevaluate other HDPs to better define how commonly permeabilization and action at non-membrane targets occur in the same peptide.

A number of non-membrane targets have been described previously for HDPs, albeit less commonly with human HDPs – in part because studies specifically looking for this are relatively uncommon. Further evaluating this possibility will be important not only for our ability to better develop peptidomimetics based on the LL-37 scaffold, but also to better understand the antimicrobial targets selected for in the course of human evolution, which may serve as the basis for target-based development of both peptidomimetic and traditional small molecule antibiotics. In particular, the problem of achieving clinically-relevant selective membranolysis has been outstanding now for many decades. It is unclear whether it is solvable. Even if it is not, however, it may still be possible to mine LL-37 and other HDPs for non-membrane targets to serve as the basis for non-membranolytic therapeutics.

## Methods

### Peptide synthesis and purification –

Methods for peptide synthesis and purification are generally as previously described (*e.g.*, (43–45)). Peptides used in this study were made on a third-generation automated flow synthesizer following preloading of 0.45 mmol/g 4-(4-Hydroxymethyl-3-methoxyphenoxy)butyric acid (HMPB) resin (ChemMatrix) with the relevant C-terminal amino acid using overnight reaction with N,N’-Diisopropylcarbodiimide (DIC) and 4-dimethylamino pyridine (DMAP).

Following flow synthesis, resin was cleaved by treatment with Reagent K (82.5% trifluoroacetic acid (TFA), 5% water, 5% phenol, 2.5% 1,2-ethanedithiol) for 2 hours at room temperature followed by peptide precipitation with ether chilled on dry ice. Precipitated peptide was then resuspended in 50% water / 50% acetonitrile with 0.1% TFA, flash frozen, and lyophilized. Crude peptide was purified on reverse phase columns using a Biotage Selekt flash chromatography system.

### Circular dichroism spectroscopy –

Samples were prepared for circular dichroism spectroscopy by dissolving peptides at 0.5 mg/mL in water and then diluting with 20% v/v trifluoroethanol. Data were collected on an Aviv 420 CD Spectrometer at the indicated temperatures in 0.1 cm pathlength cuvettes. Calculation of secondary structural content was completed using BeStSel(25).

### Size exclusion chromatography –

Size exclusion chromatography was performed on an Agilent 1260 Infinity II Bio-Inert liquid chromatography system using a Superdex 75 100/300 column and PBS buffer supplemented with 3.5% isopropanol with injection of approximately 25-50 µg per peptide and a flow rate of 0.8 mL / minute over 30 minutes and monitoring at 214 nm.

### Biolayer interferometry –

Biolayer interferometry was performed on a GatorPlus bio-layer interferometry system (GatorBio) using streptavidin probes. Following an initial baseline measure for 30 seconds, probes were loaded with 800 nM N-biotinylated LL-37 in PBS for 180 seconds. Following another 30 second baseline read, association was measured for 120 seconds followed by dissociation for 240 seconds. Analyte concentration series of LL-37, m4, or m15 were 12.5 µM, 6.3 µM, 3.1 µM, 1.6 µM, or 0.8 µM with no analyte (buffer) and no probe (analyte only at 12.5 µM) controls. Calculation of interaction kinetics was completed in GatorOne software associated with the instrument.

### Antimicrobial susceptibility testing –

Testing for antimicrobial activity of peptides of interest was completed as per Clinical and Laboratory Standards Institute methods with previously described modifications for the use of cationic peptides, specifically the use of polypropylene plates (Greiner) and unadjusted Mueller Hinton Broth (Difco) (46). These methods were scaled down to either 35 or 50 µL per well in 384-well plates. In general, peptide was added initially as a 10x concentrate in water followed by the addition of a concentrated broth and cell mix to a final 1x concentration of all reagents. Following plating, growth was monitored by OD600 on a Tecan Spark plate reader over the course of 18 hours at 37 °C, during which time the plate was incubated in a Spark Large Humidity Cassette to minimize evaporation.

### Strains –

The primary bacterial strains used in these studies were *E. coli* ATCC 25922 and *P. aeruginosa* ATCC 27853, which are quality control strains for antimicrobial susceptibility testing. Strains of *P. aeruginosa* isolated from patients with cystic fibrosis were chosen at random from a collection of previously published clinical strains(47) maintained at the Massachusetts General Hospital Cystic Fibrosis Center.

### Hemolysis Assays –

Hemolysis assays were carried out by treating blood samples with a peptide of interest over a range of concentrations. In general, blood was spun for 10 minutes at 800 rcf and supernatant decanted to clearance, after which cells were brought up in PBS at 1% v/v for eventual final dilution to 0.5% with a 2x concentration of peptide dissolved in PBS. Final well volumes were 50 µL each in 384-well polypropylene plates (Greiner). Reactions were then monitored over 18 hours by OD600 as a kinetic marker of hemolysis. In addition to this kinetic marker, endpoint measures of hemolysis were made by adding 25 µL of PBS to each well at the end of incubation, spinning plates for 10 minutes at 800 rcf, and then removing 25 µL of supernatant for evaluation of free hemoglobin levels by absorbance at 414 nm. Types of blood used in these studies included defibrinated sheep and cow blood (Hardy Diagnostics), purchased human whole blood from two separate donors (ZenBio), sheep RBCs (Innovative Research), and human RBCs (Rockland Immunochemicals).

Quantification of hemolysis was completed by comparison to a thrice freeze-thawed control of the same blood used in plating set to 100%. Thus, percentages are not a literal percentage of cells, but rather a bulk percentage of signal. The choice of the percentage cutoff for determining samples with substantial hemolysis (20% for most blood types with the exception of two samples set to 40%) was based on the percentage at there was a clear cutoff between signal and noise in the colistin concentration gradient, the premise being that colistin can be used *in vivo* and thus provides a rough surrogate for approximating where a peptide is on the toxicity spectrum from LL-37 to colistin.

### Luminescence toxicity assays –

Luminescence toxicity assays were completed using RealTime Glo reagents per the manufacturer’s instructions. Following initial pilot studies, HepG2 cells (maintained in DMEM with 10% FBS) were plated at a density of 5,000 cells per well, while LLC-PK1 (maintained in Medium 199 with 5% FBS), HL-60 (IMDM with 20% FBS), and THP-1 (maintained cells were plated at 20,000 cells/well in 25 µL per well of white clear-bottom plates. These were then allowed to settle for approximately 12 hours, after which 25 µL of RealTime Glo reagents in the appropriate medium and 5 µL of a 10x concentration of peptide in water. Wells were then monitored over 18 hours in a Tecan Spark Plate reader and incubated in a Small Humidity Cassette to minimize evaporation over time. HepG2 cells were maintained in EMEM medium (ATCC) with 10% FBS. LLC-PK1 cells were maintained in Medium 199 (Sigma) with 5% FBS. THP-1 cells were maintained in RPMI medium (ATCC) with 10% FBS and supplemented with fresh β mercaptoethanol to 0.05 mM final concentration at the time of usage (Gibco). HL-60 cells were maintained in IMDM medium (ATCC) with 20% FBS. Adherent cells were split using Trypsin-EDTA (ATCC).

### Dual membrane permeabilization assays –

Dual membrane permeabilization assays were based on prior descriptions(*e.g.*, (48)), and scaled down to take place in 50 µL total volume in 384-well black plates with transparent bottoms (Greiner). In brief, cells were grown in unadjusted MHB to a density of OD600 ≥ 0.5. Cells were then spun down at 10,000 rcf for 1 minute, washed with the buffer of interest, spun again, and then resuspend at OD600 0.5 in the buffer of interest. 50 µL of cells in buffer were then plated directly onto 5 µL of 10x peptide over a range of concentrations as shown, with monitoring every 15 minutes over a course of 4 hours – beyond which time we had observed little change in pilot experiments (data not shown). For outer membrane permeabilization, N-Phenyl-1-Naphthylamine (NPN, TCI) was dissolved in acetone and then in buffer to a final concentration of 10 µM. For inner membrane permeabilization, propidium iodide (Cayman Chemical) was dissolved in water and used at a final concentration of 5 µM. Plates were maintained at 37 °C over the course of the experiment in a Large Humidity Cassett to minimize evaporation. Signal was monitored over time in a Tecan Spark plate reader at Ex 350 / Em 420 for NPN and Ex 535 / Em 617 for PI.

## Acknowledgements

This work has been funded in by the National Institute of Allergy and Infectious Diseases (T32 AI007061 and K08 AI166345 to JSA; U19 AI142780 to BLP) and by the Cystic Fibrosis Foundation (ALBIN19F0, ALBIN21Q0, ALBIN22A0-KB to JSA).

## Author Contributions

JSA designed and conducted the experiments and wrote the paper. DAV, WV, and CJ performed the experiments. BLP guided the experiments and wrote the paper.

