## Supplementary Figures 1-135 for "Separability of antibacterial and membranolytic activity in the human host defense peptide LL-37"

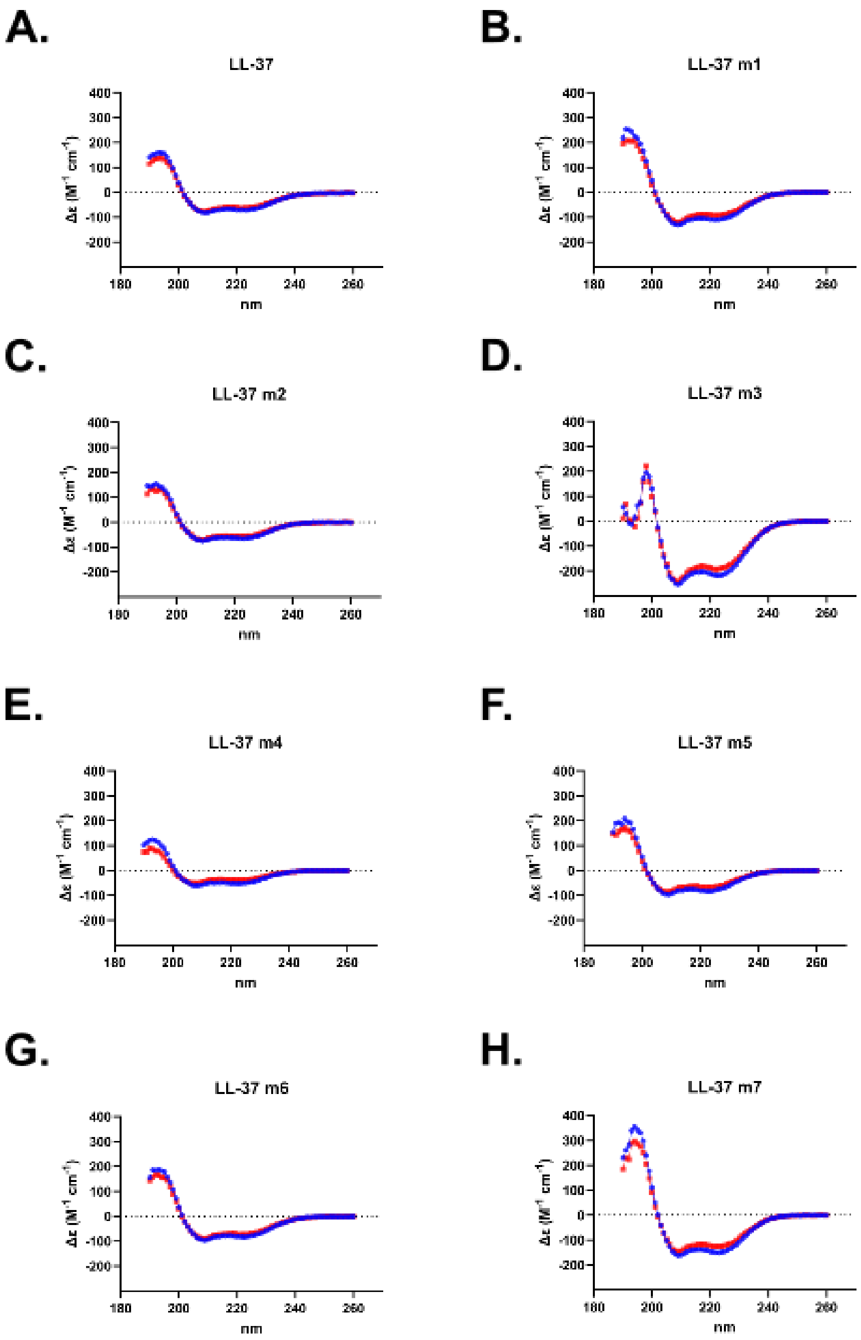


**I.**

| | % Helix | |
|---|---|---|
| | 20 °C | 37 °C |
| LL-37 | 100 | 100 |
| LL-37 m1 | 100 | 100 |
| LL-37 m2 | 100 | 100 |
| LL-37 m3 | 100 | 100 |
| LL-37 m4 | 100 | 100 |
| LL-37 m5 | 100 | 100 |
| LL-37 m6 | 100 | 100 |
| LL-37 m7 | 100 | 100 |

**Supplementary Figure 1 – Circular dichroism (CD) spectroscopy of Figure 1 mutants.** **A-H.** CD spectra are displayed for each mutant dissolved in water plus 20% trifluoroethanol, with spectra covering 190-260 nm at both 20 °C (Blue) and 37 °C (Red). **I.** Quantification of helical content as determined by BeStSel for each mutant.

A.

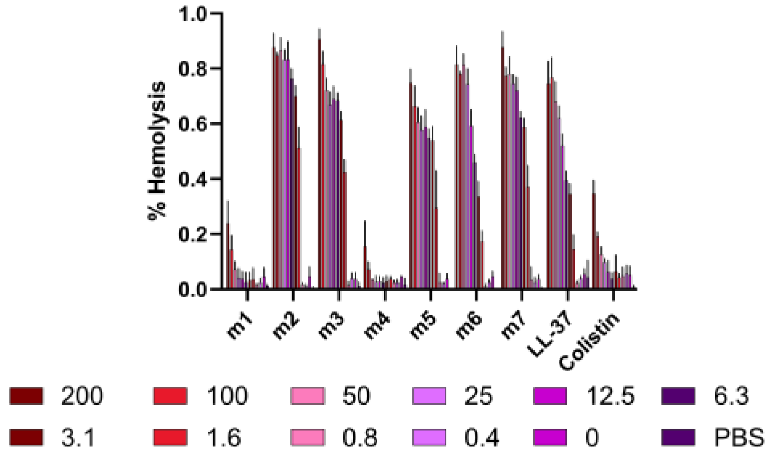
% Hemolysis
1.0
0.8
0.6
0.4
0.2
0.0
m1
m2
m3
m4
m5
m6
m7
LL-37
Colistin
200
100
50
25
12.5
6.3
3.1
1.6
0.8
0.4
0
PBS

B.

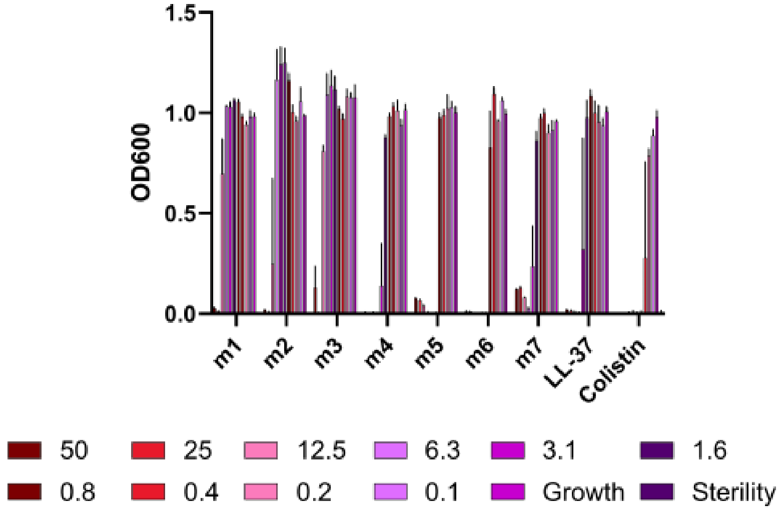
OD600
1.5
1.0
0.5
0.0
m1
m2
m3
m4
m5
m6
m7
LL-37
Colistin
50
25
12.5
6.3
3.1
1.6
0.8
0.4
0.2
0.1
Growth
Sterility

C.

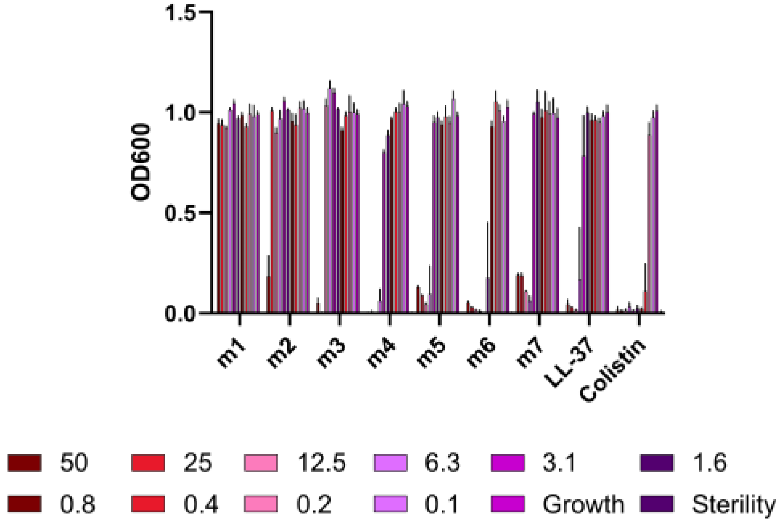
OD600
1.5
1.0
0.5
0.0
m1
m2
m3
m4
m5
m6
m7
LL-37
Colistin
50
25
12.5
6.3
3.1
1.6
0.8
0.4
0.2
0.1
Growth
Sterility

**Supplementary Figure 2 – Histogram depiction of Figure 1 data. A.** Endpoint hemolytic activity of each peptide against defibrinated sheep blood relative to a 100% hemolysis thrice freeze-thawed control after 18 hours at 37 °C. Raw data were obtained by absorbance at 414 nm in well supernatants for free hemoglobin. **B-C.** Endpoint activity of each peptide against *E. coli* (**B**) or *Pseudomonas* (**C**) after 18 hours of incubation at 37 °C in the presence of the indicated concentrations of each peptide. Data are the raw OD600 after 18 hours at 37 °C for three independent experiments. Peptide concentrations are given in μM.

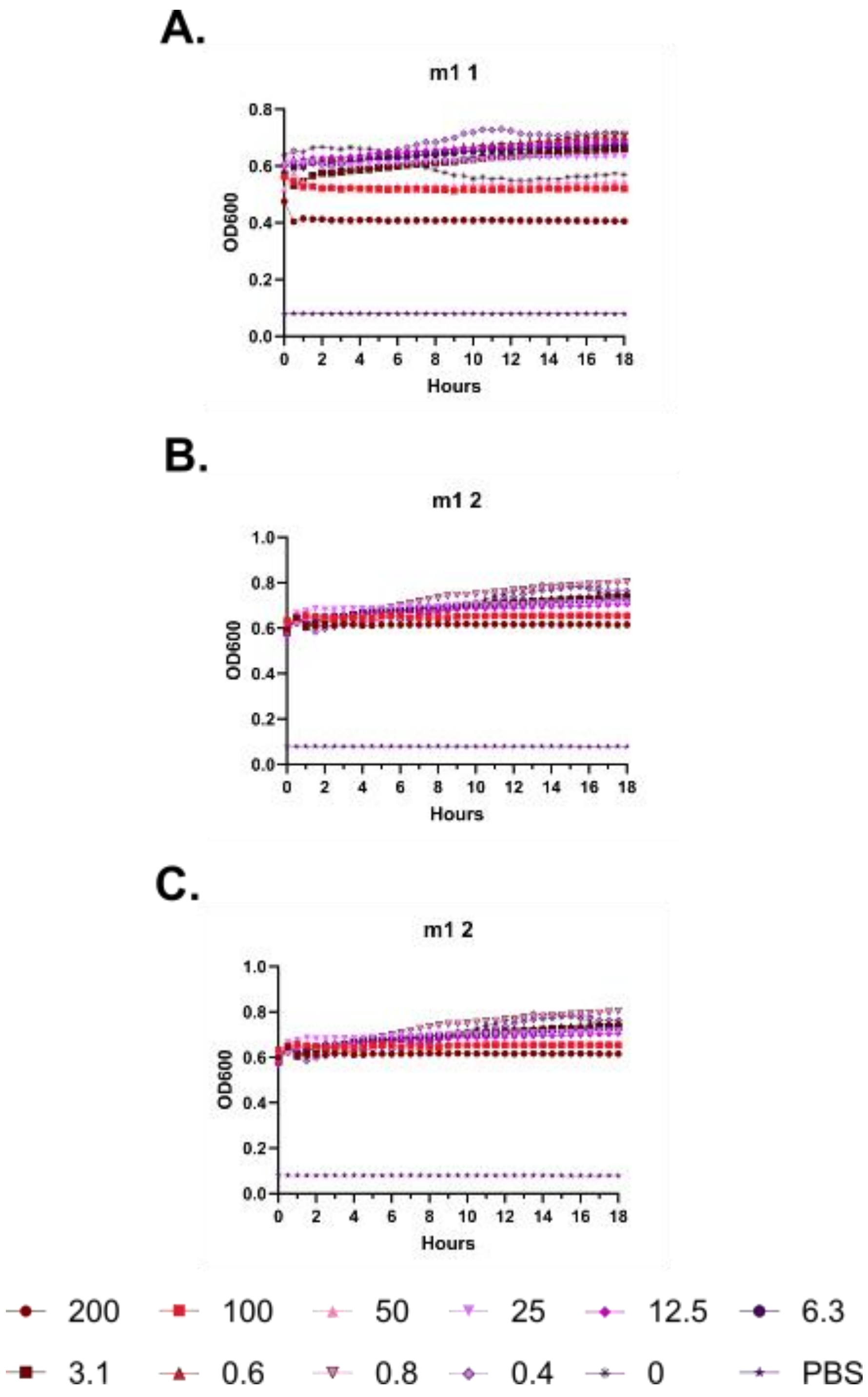
A.
m1 1
OD600
0.0
0.2
0.4
0.6
0.8
Hours
0 2 4 6 8 10 12 14 16 18
B.
m1 2
OD600
0.0
0.2
0.4
0.6
0.8
1.0
Hours
0 2 4 6 8 10 12 14 16 18
C.
m1 2
OD600
0.0
0.2
0.4
0.6
0.8
1.0
Hours
0 2 4 6 8 10 12 14 16 18
200
100
50
25
12.5
6.3
3.1
0.6
0.8
0.4
0
PBS

**Supplementary Figure 3 – Kinetic hemolysis assays for (m1). A.** Hemolysis was monitored by OD600 over 18 hours at 37 °C in the presence of the indicated peptide concentrations. Decreasing OD600 indicates increasing hemolysis. Peptide concentrations are given in μM. Each curve indicates an independent experiment.

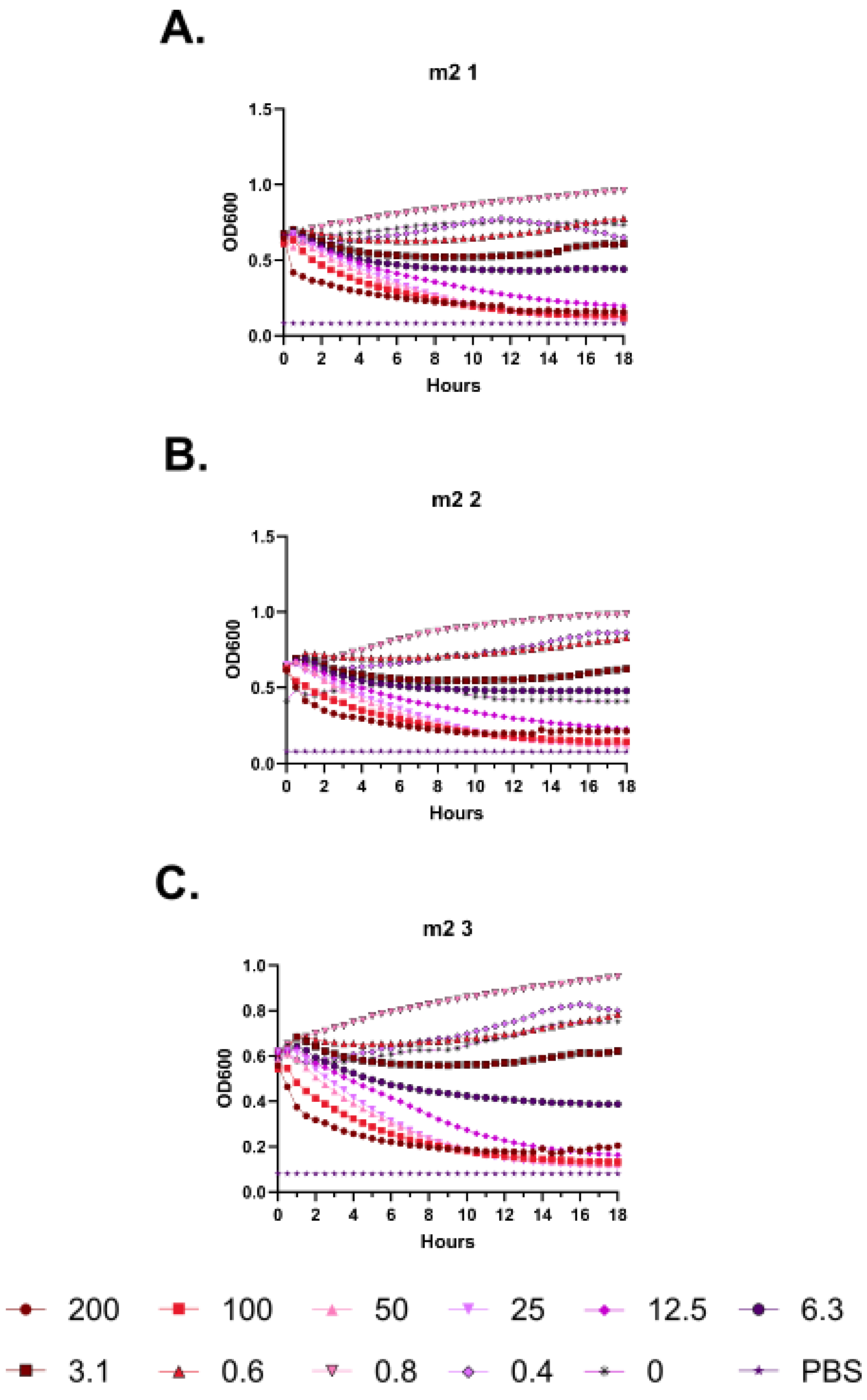
A.
m2 1
1.5
1.0
0.5
0.0
OD600
0 2 4 6 8 10 12 14 16 18
Hours
B.
m2 2
1.5
1.0
0.5
0.0
OD600
0 2 4 6 8 10 12 14 16 18
Hours
C.
m2 3
1.0
0.8
0.6
0.4
0.2
0.0
OD600
0 2 4 6 8 10 12 14 16 18
Hours
200
100
50
25
12.5
6.3
3.1
0.6
0.8
0.4
0
PBS

**Supplementary Figure 4 – Kinetic hemolysis assays for (m2). A-C.** Hemolysis was monitored by OD600 over 18 hours at 37 °C in the presence of the indicated peptide concentrations. Decreasing OD600 indicates increasing hemolysis. Peptide concentrations are given in μM. Each curve indicates an independent experiment.

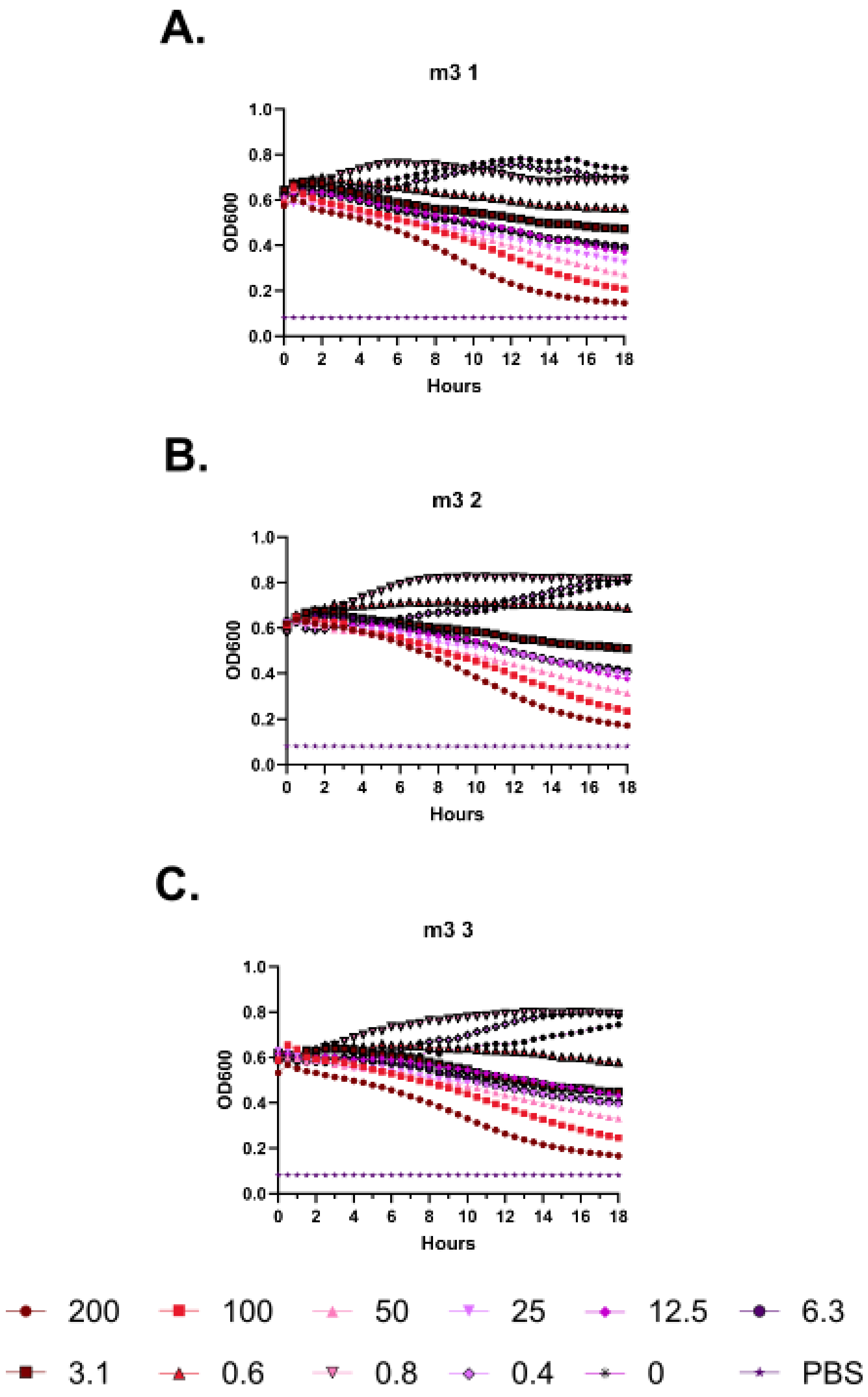
A.
m3 1
OD600
Hours
B.
m3 2
OD600
Hours
C.
m3 3
OD600
Hours
1.0
0.8
0.6
0.4
0.2
0.0
0 2 4 6 8 10 12 14 16 18
200
100
50
25
12.5
6.3
3.1
0.6
0.8
0.4
0
PBS

**Supplementary Figure 5 – Kinetic hemolysis assays for (m3). A-C.** Hemolysis was monitored by OD600 over 18 hours at 37 °C in the presence of the indicated peptide concentrations. Decreasing OD600 indicates increasing hemolysis. Peptide concentrations are given in μM. Each curve indicates an independent experiment.

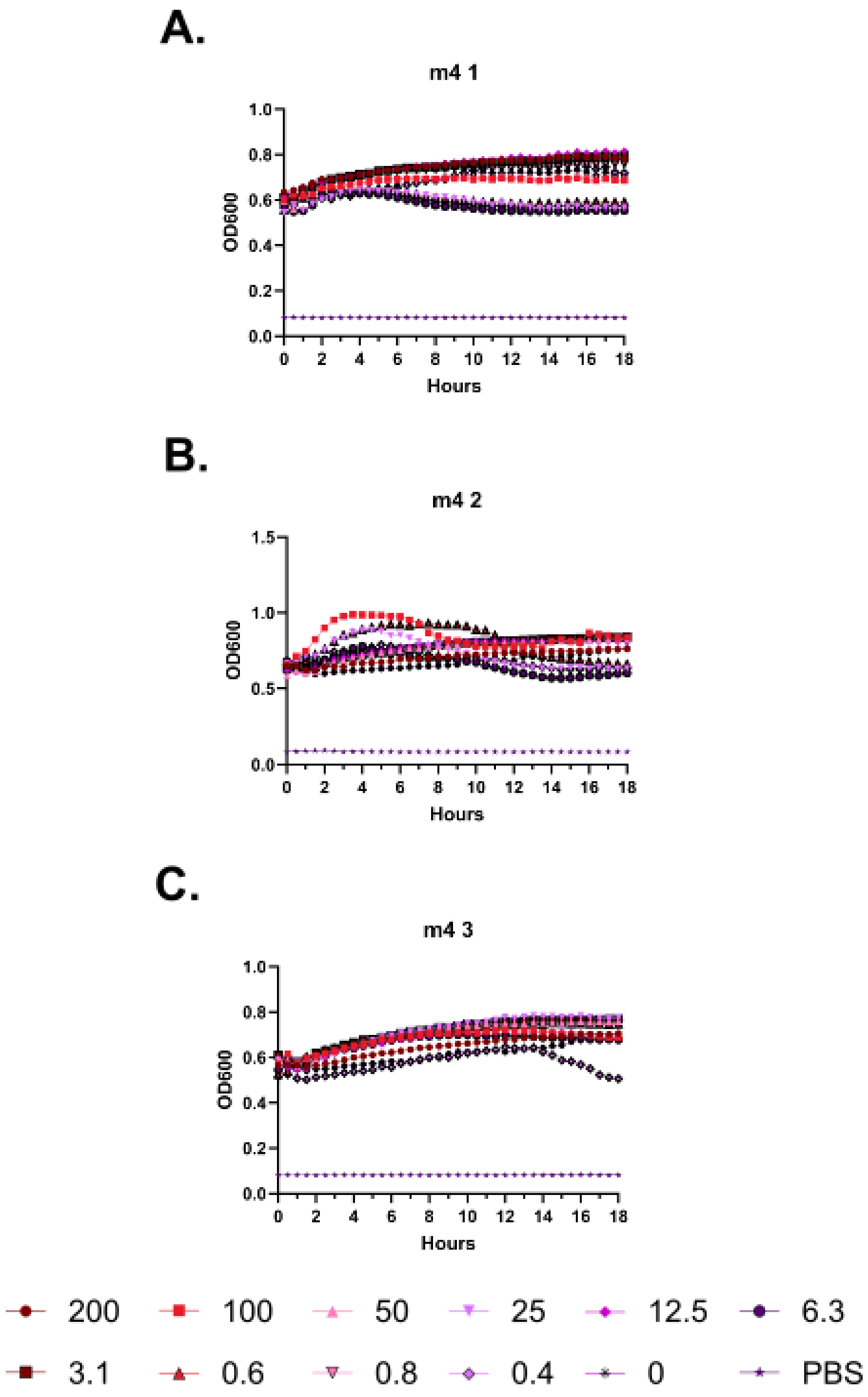
A.
m4 1
OD600
Hours
B.
m4 2
OD600
Hours
C.
m4 3
OD600
Hours
200
100
50
25
12.5
6.3
3.1
0.6
0.8
0.4
0
PBS

**Supplementary Figure 6 – Kinetic hemolysis assays for (m4). A-C.** Hemolysis was monitored by OD600 over 18 hours at 37 °C in the presence of the indicated peptide concentrations. Decreasing OD600 indicates increasing hemolysis. Peptide concentrations are given in µM. Each curve indicates an independent experiment.

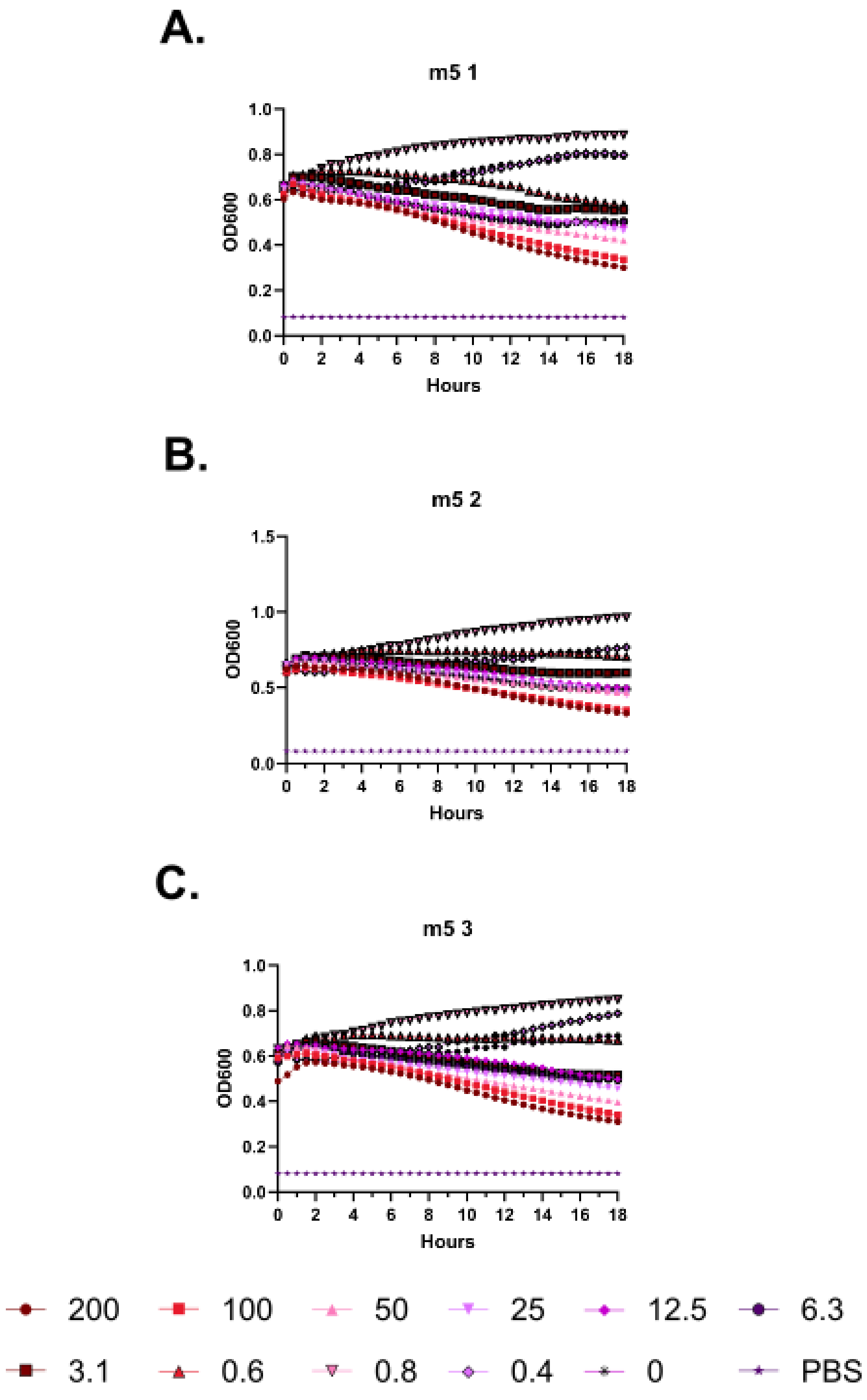
A.
m5 1
OD600
Hours
B.
m5 2
OD600
Hours
C.
m5 3
OD600
Hours
200
100
50
25
12.5
6.3
3.1
0.6
0.8
0.4
0
PBS

**Supplementary Figure 7 – Kinetic hemolysis assays for (m5). A-C.** Hemolysis was monitored by OD600 over 18 hours at 37 °C in the presence of the indicated peptide concentrations. Decreasing OD600 indicates increasing hemolysis. Peptide concentrations are given in µM. Each curve indicates an independent experiment.

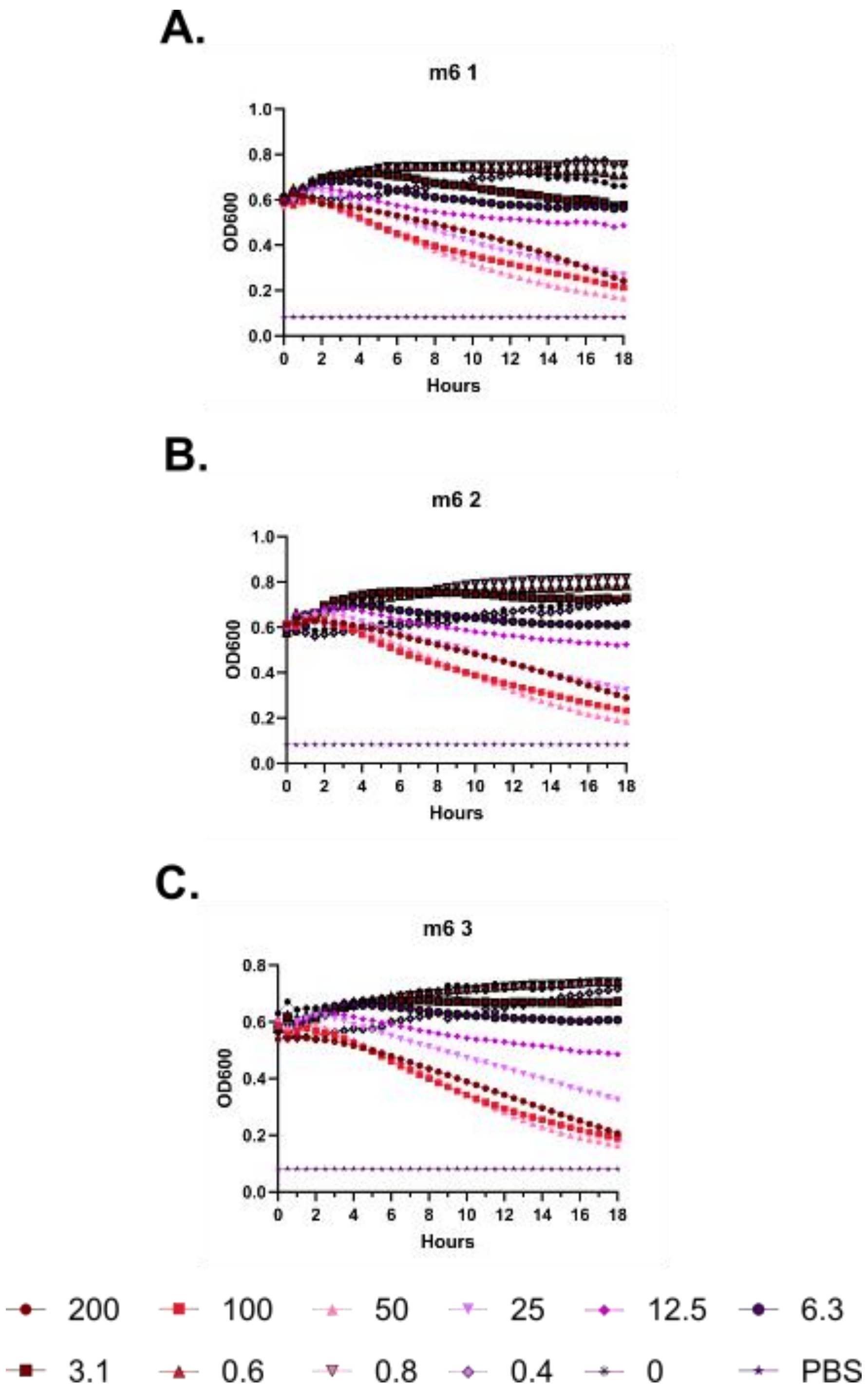
A.
m6 1
OD600
Hours
B.
m6 2
OD600
Hours
C.
m6 3
OD600
Hours
200
100
50
25
12.5
6.3
3.1
0.6
0.8
0.4
0
PBS

**Supplementary Figure 8 – Kinetic hemolysis assays for (m6). A-C.** Hemolysis was monitored by OD600 over 18 hours at 37 °C in the presence of the indicated peptide concentrations. Decreasing OD600 indicates increasing hemolysis. Peptide concentrations are given in μM. Each curve indicates an independent experiment.

**A.**

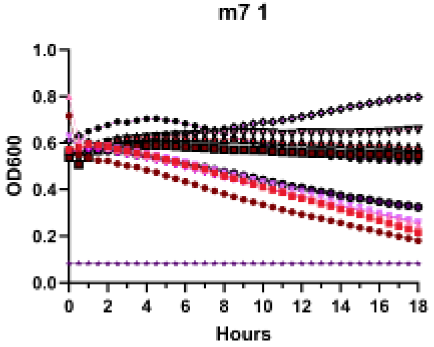


**B.**

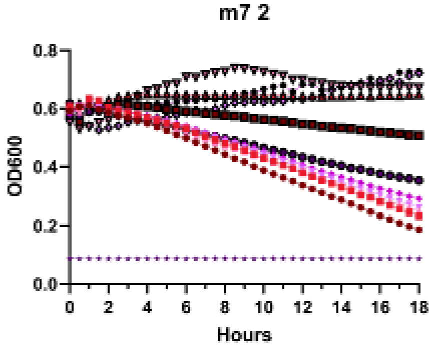


**C.**

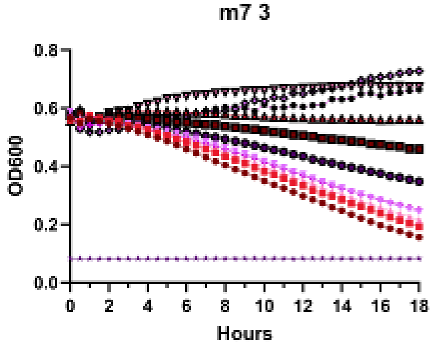


| | | | | | |
|---|---|---|---|---|---|
| 200 | 100 | 50 | 25 | 12.5 | 6.3 |
| 3.1 | 0.6 | 0.8 | 0.4 | 0 | PBS |

**Supplementary Figure 9 – Kinetic hemolysis assays for (m7). A-C.** Hemolysis was monitored by OD600 over 18 hours at 37 °C in the presence of the indicated peptide concentrations. Decreasing OD600 indicates increasing hemolysis. Peptide concentrations are given in μM. Each curve indicates an independent experiment.

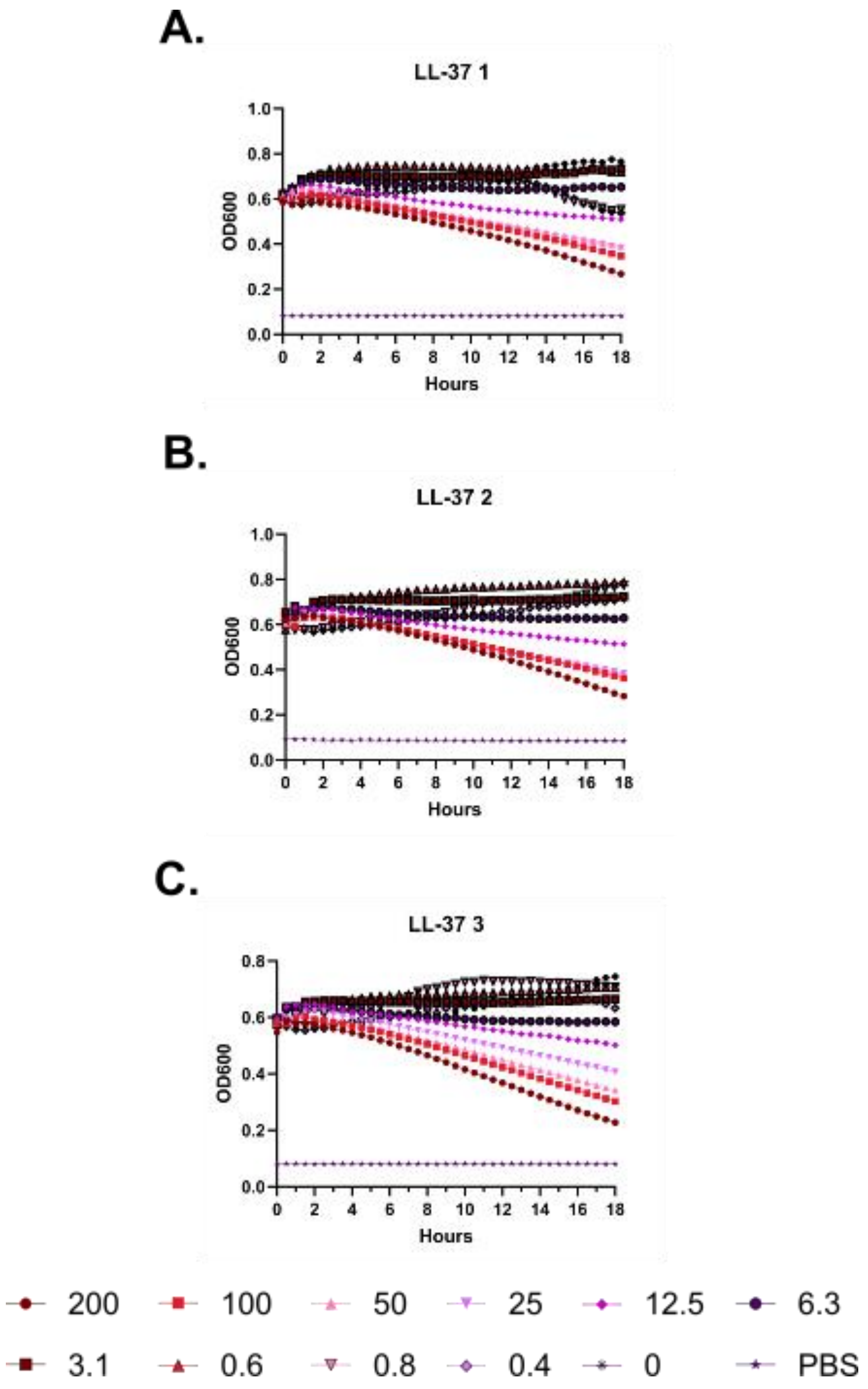
A.
LL-37 1
OD600
Hours
B.
LL-37 2
OD600
Hours
C.
LL-37 3
OD600
Hours
200
100
50
25
12.5
6.3
3.1
0.6
0.8
0.4
0
PBS

**Supplementary Figure 10 – Kinetic hemolysis assays for (LL-37). A-C.** Hemolysis was monitored by OD600 over 18 hours at 37 °C in the presence of the indicated peptide concentrations. Decreasing OD600 indicates increasing hemolysis. Peptide concentrations are given in μM. Each curve indicates an independent experiment.

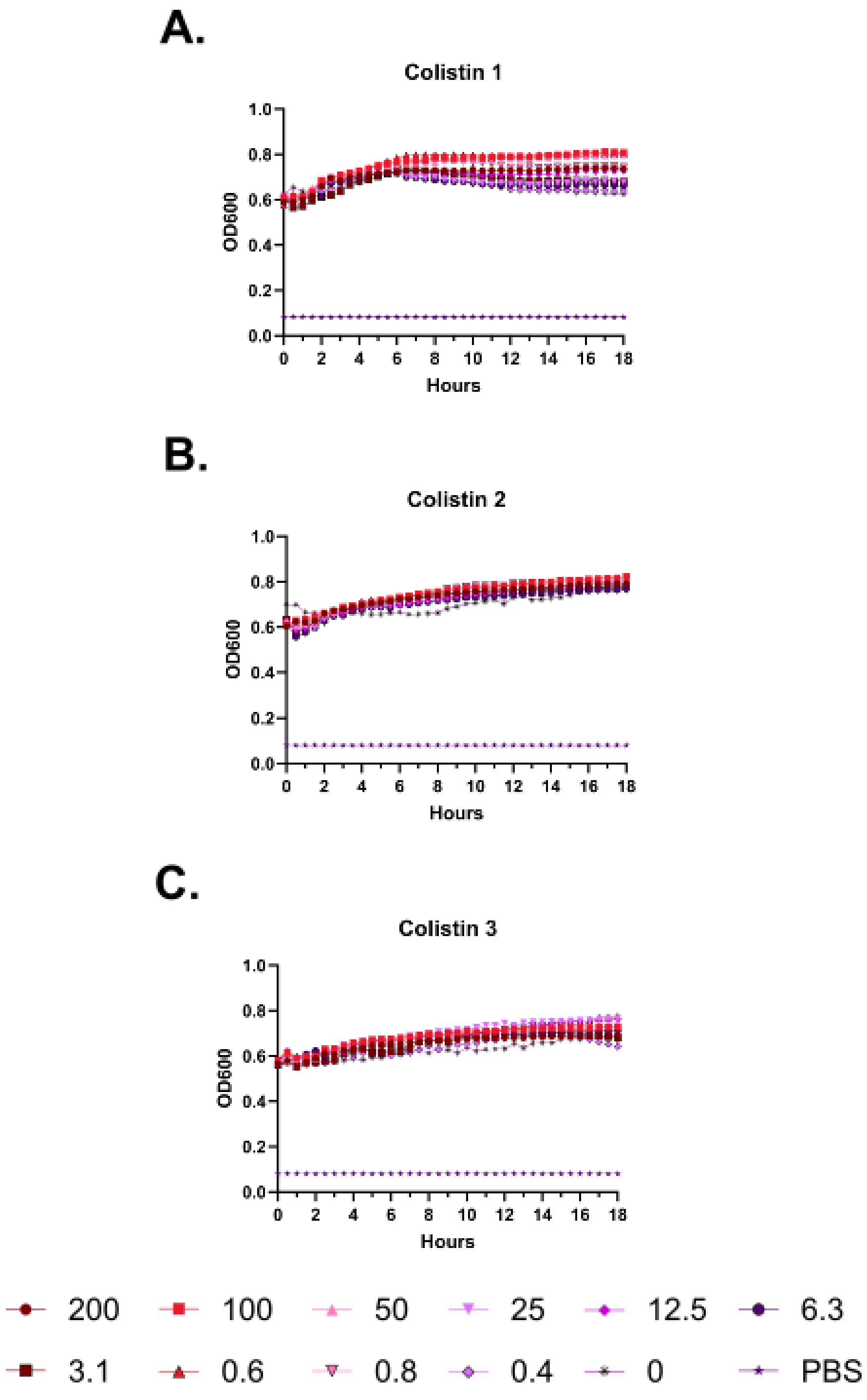
A.
Colistin 1
OD600
Hours
1.0
0.8
0.6
0.4
0.2
0.0
0
2
4
6
8
10
12
14
16
18
B.
Colistin 2
OD600
Hours
C.
Colistin 3
OD600
Hours
200
100
50
25
12.5
6.3
3.1
0.6
0.8
0.4
0
PBS

**Supplementary Figure 11 – Kinetic hemolysis assays for (Colistin). A-C.** Hemolysis was monitored by OD600 over 18 hours at 37 °C in the presence of the indicated peptide concentrations. Decreasing OD600 indicates increasing hemolysis. Peptide concentrations are given in µM. Each curve indicates an independent experiment.

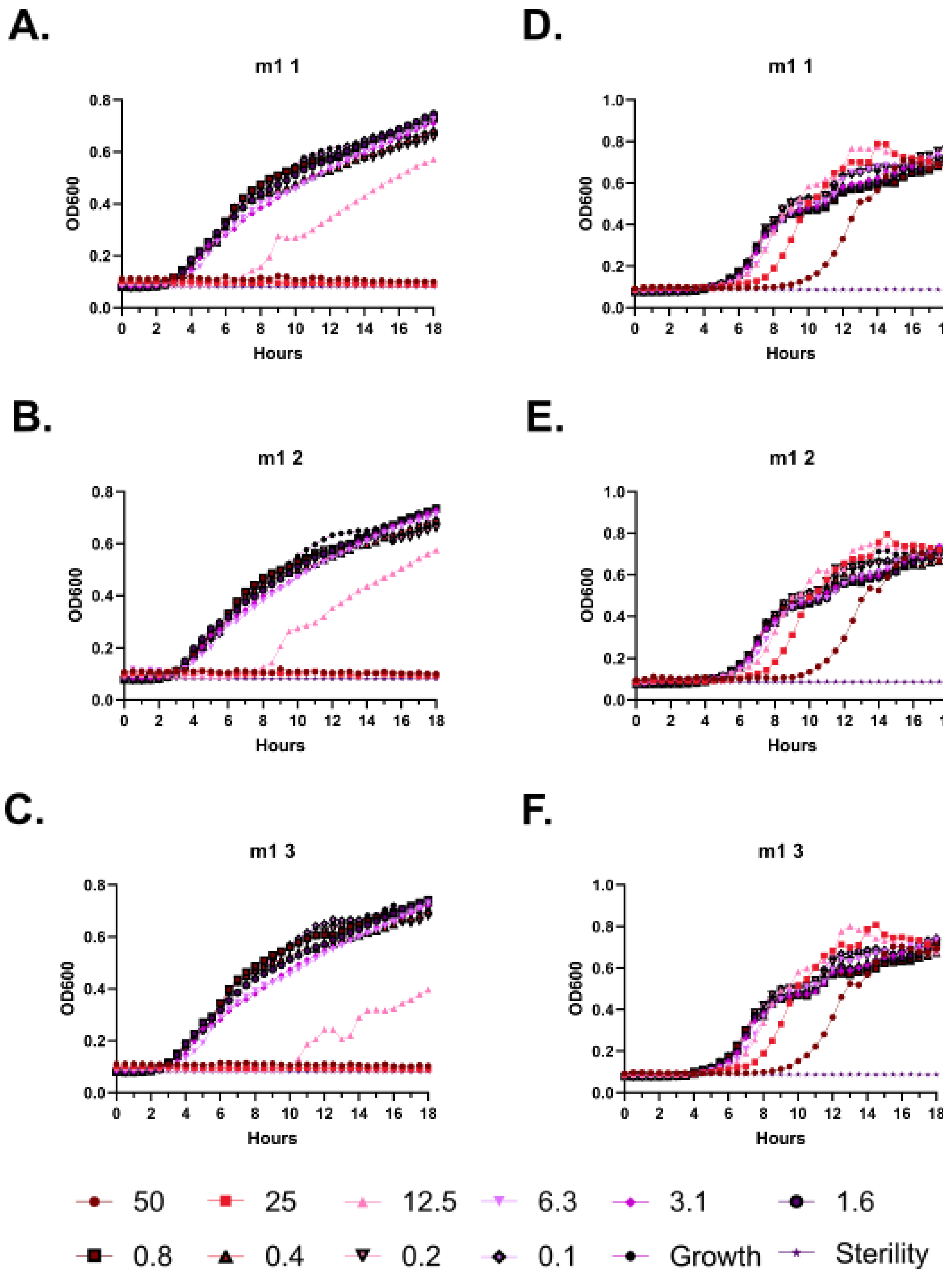
A.
m1 1
B.
m1 2
C.
m1 3
D.
m1 1
E.
m1 2
F.
m1 3
OD600
Hours
50
25
12.5
6.3
3.1
1.6
0.8
0.4
0.2
0.1
Growth
Sterility

**Supplementary Figure 12 – Kinetic antimicrobial susceptibility testing for (m1). A-C.** *E. coli*. **D-F.** *Pseudomonas.* Bacterial growth was monitored by OD600 over 18 hours at 37 °C in the presence of the indicated peptide concentrations. Increasing OD600 indicates increasing bacterial growth. Peptide concentrations are given in μM. Each curve indicates an independent experiment.

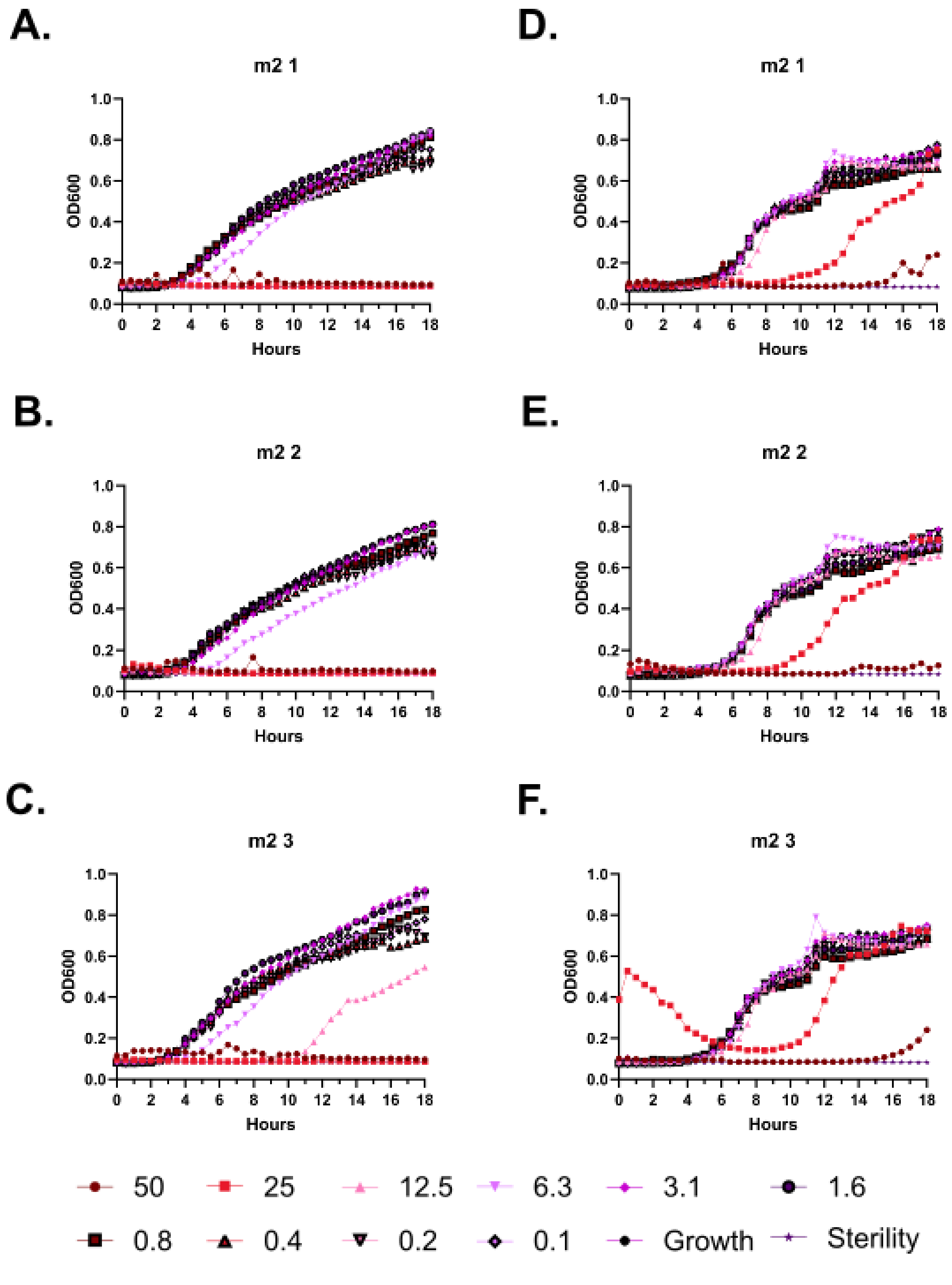

**Supplementary Figure 13 – Kinetic antimicrobial susceptibility testing for (m2). A-C.** *E. coli*. **D-F.** *Pseudomonas.* Bacterial growth was monitored by OD600 over 18 hours at 37 °C in the presence of the indicated peptide concentrations. Increasing OD600 indicates increasing bacterial growth. Peptide concentrations are given in µM. Each curve indicates an independent experiment.

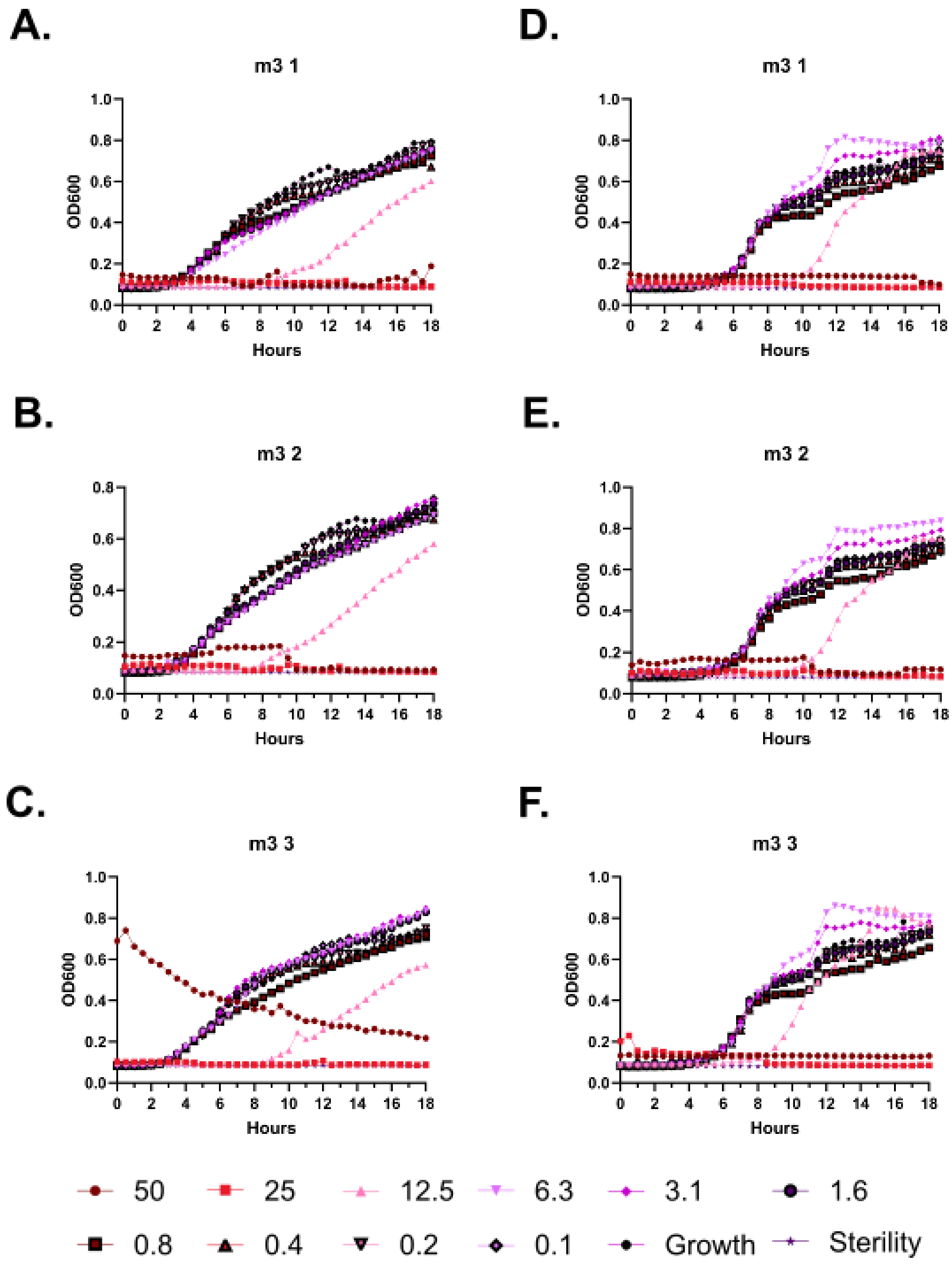
A.
m3 1
B.
m3 2
C.
m3 3
D.
m3 1
E.
m3 2
F.
m3 3
OD600
Hours
50
25
12.5
6.3
3.1
1.6
0.8
0.4
0.2
0.1
Growth
Sterility

**Supplementary Figure 14 – Kinetic antimicrobial susceptibility testing for (m3). A-C.** *E. coli*. **D-F.** *Pseudomonas.* Bacterial growth was monitored by OD600 over 18 hours at 37 °C in the presence of the indicated peptide concentrations. Increasing OD600 indicates increasing bacterial growth. Peptide concentrations are given in µM. Each curve indicates an independent experiment.

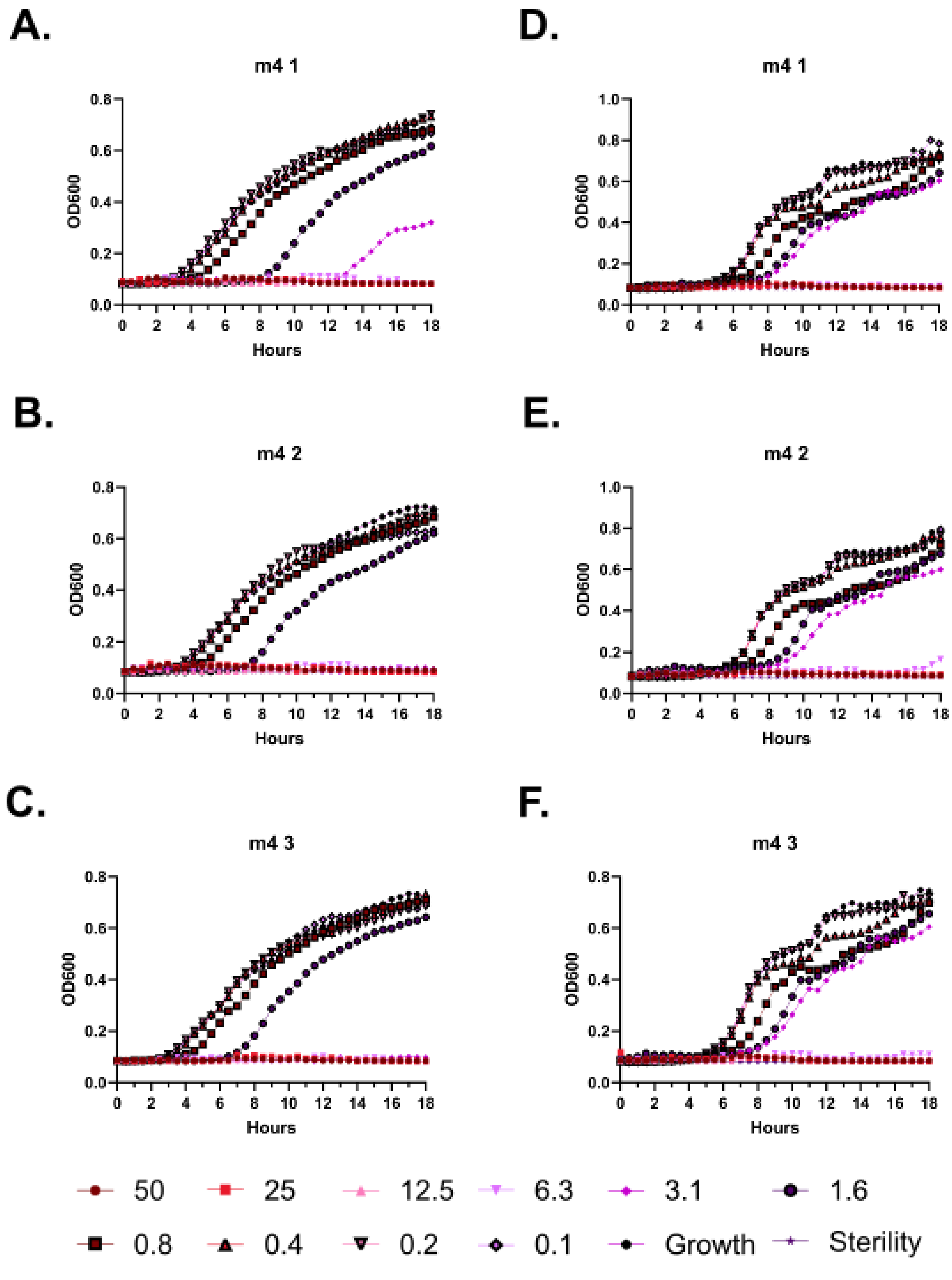
A.
m4 1
B.
m4 2
C.
m4 3
D.
m4 1
E.
m4 2
F.
m4 3
OD600
Hours
50
25
12.5
6.3
3.1
1.6
0.8
0.4
0.2
0.1
Growth
Sterility

**Supplementary Figure 15 – Kinetic antimicrobial susceptibility testing for (m4). A-C.** *E. coli*. **D-F.** *Pseudomonas.* Bacterial growth was monitored by OD600 over 18 hours at 37 °C in the presence of the indicated peptide concentrations. Increasing OD600 indicates increasing bacterial growth. Peptide concentrations are given in µM. Each curve indicates an independent experiment.

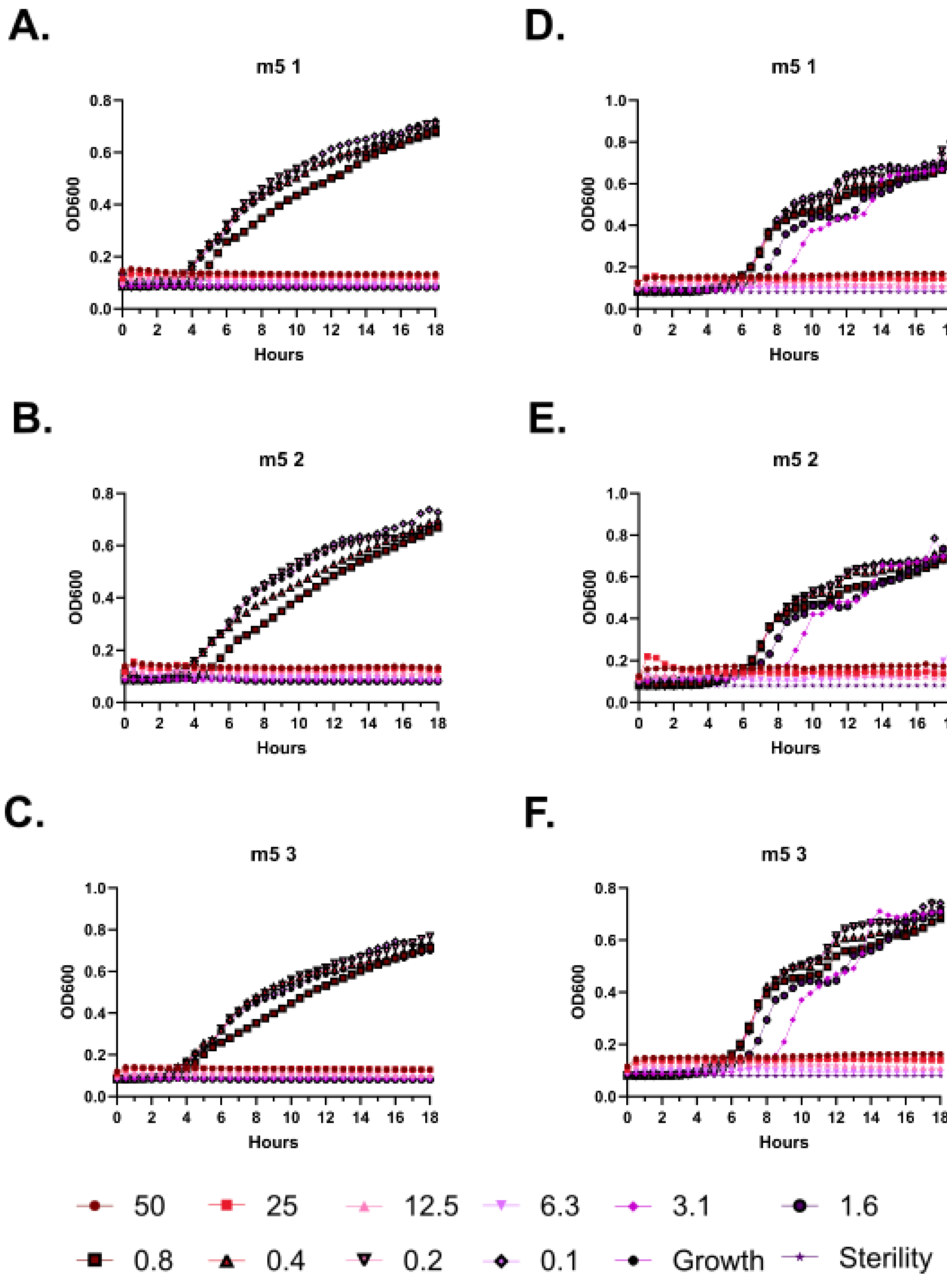
A.
m5 1
OD600
Hours
D.
m5 1
OD600
Hours
B.
m5 2
OD600
Hours
E.
m5 2
OD600
Hours
C.
m5 3
OD600
Hours
F.
m5 3
OD600
Hours
50
25
12.5
6.3
3.1
1.6
0.8
0.4
0.2
0.1
Growth
Sterility

**Supplementary Figure 16 – Kinetic antimicrobial susceptibility testing for (m5). A-C.** *E. coli*. **D-F.** *Pseudomonas.* Bacterial growth was monitored by OD600 over 18 hours at 37 °C in the presence of the indicated peptide concentrations. Increasing OD600 indicates increasing bacterial growth. Peptide concentrations are given in µM. Each curve indicates an independent experiment.

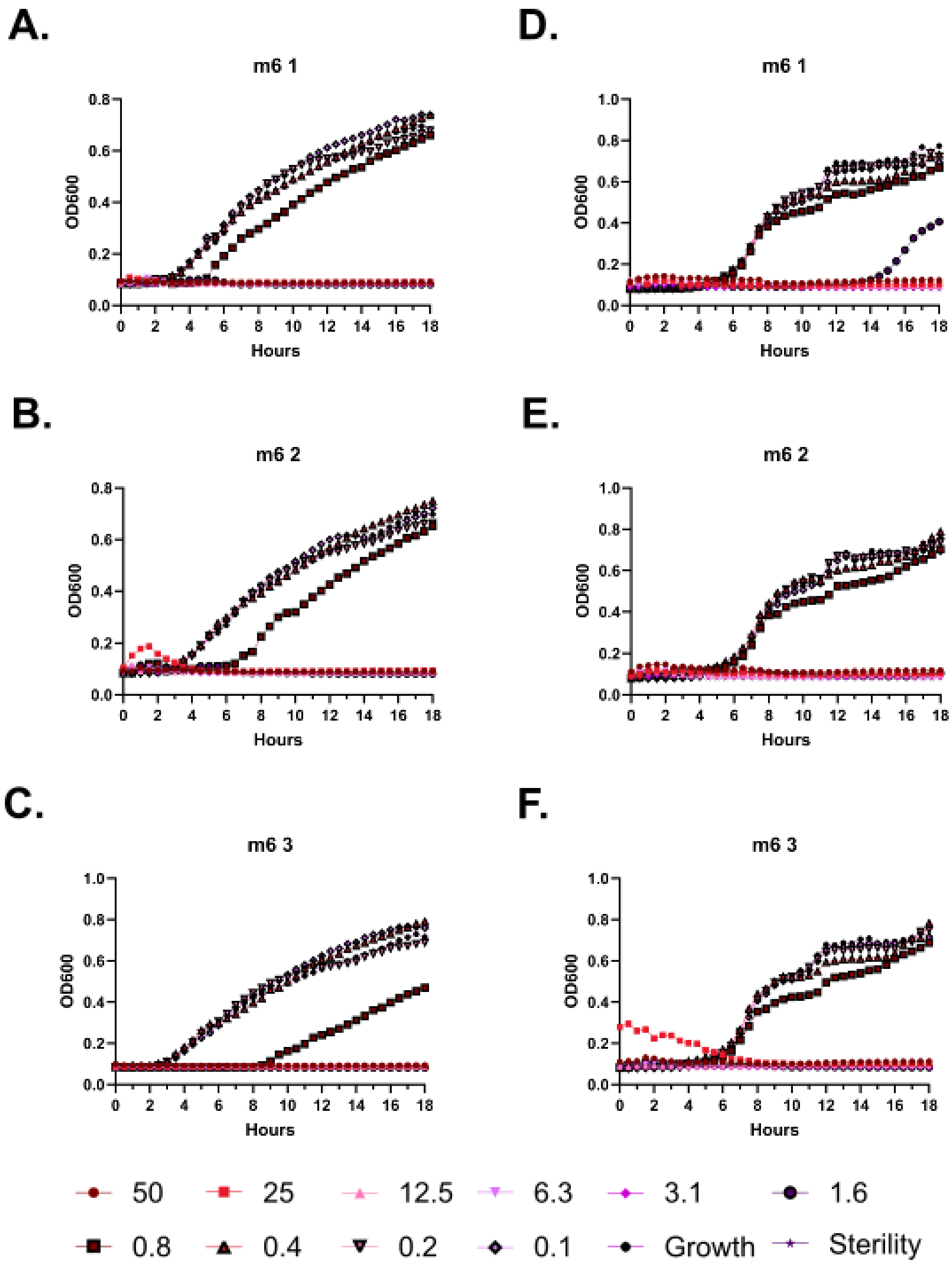
A.
m6 1
B.
m6 2
C.
m6 3
D.
m6 1
E.
m6 2
F.
m6 3
OD600
Hours
50
25
12.5
6.3
3.1
1.6
0.8
0.4
0.2
0.1
Growth
Sterility

**Supplementary Figure 17 – Kinetic antimicrobial susceptibility testing for (m6). A-C.** *E. coli*. **D-F.** *Pseudomonas.* Bacterial growth was monitored by OD600 over 18 hours at 37 °C in the presence of the indicated peptide concentrations. Increasing OD600 indicates increasing bacterial growth. Peptide concentrations are given in μM. Each curve indicates an independent experiment.

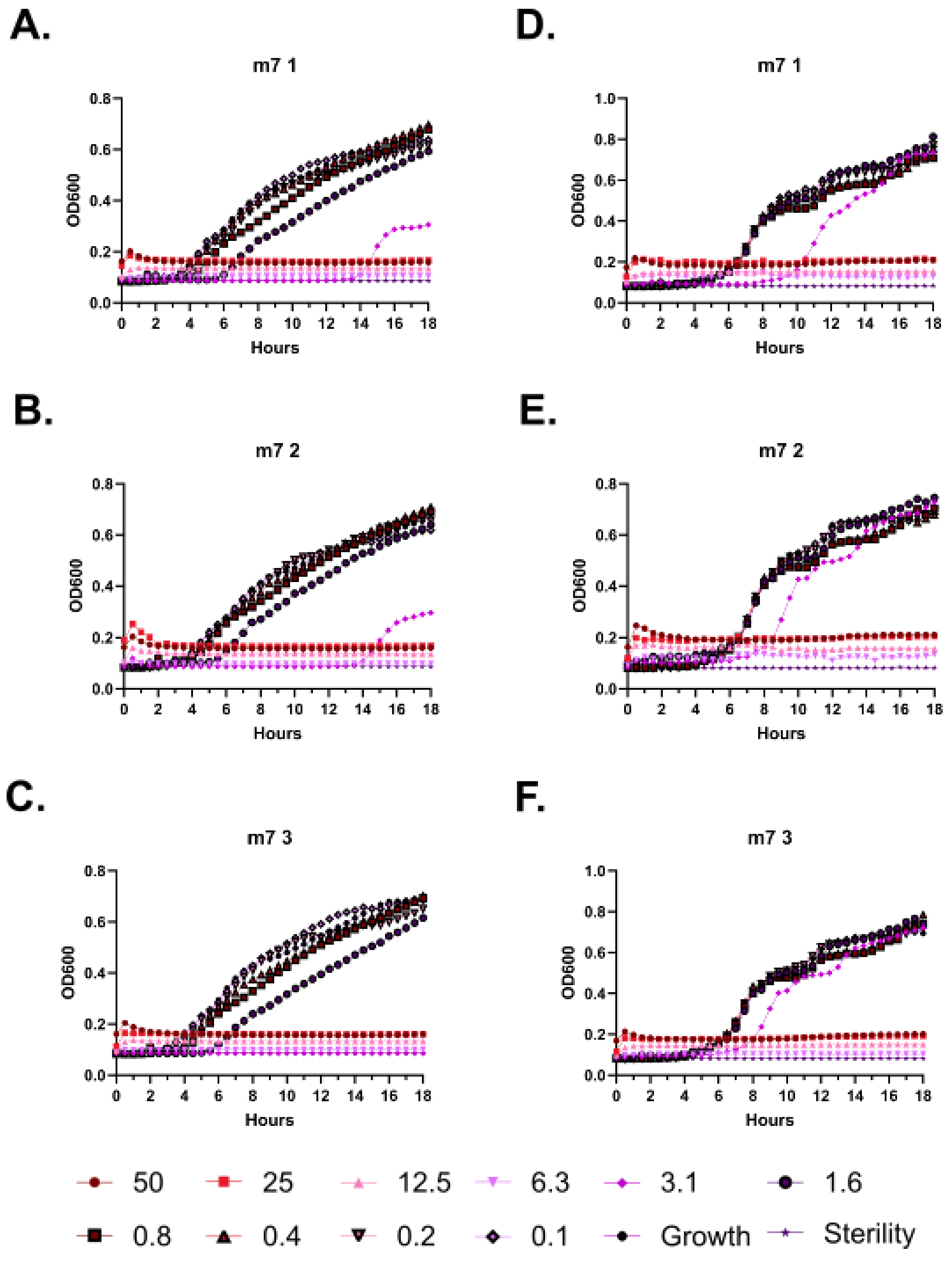
A.
m7 1
B.
m7 2
C.
m7 3
D.
m7 1
E.
m7 2
F.
m7 3
OD600
Hours
50
25
12.5
6.3
3.1
1.6
0.8
0.4
0.2
0.1
Growth
Sterility

**Supplementary Figure 18 – Kinetic antimicrobial susceptibility testing for (m7). A-C.** *E. coli*. **D-F.** *Pseudomonas.* Bacterial growth was monitored by OD600 over 18 hours at 37 °C in the presence of the indicated peptide concentrations. Increasing OD600 indicates increasing bacterial growth. Peptide concentrations are given in µM. Each curve indicates an independent experiment.

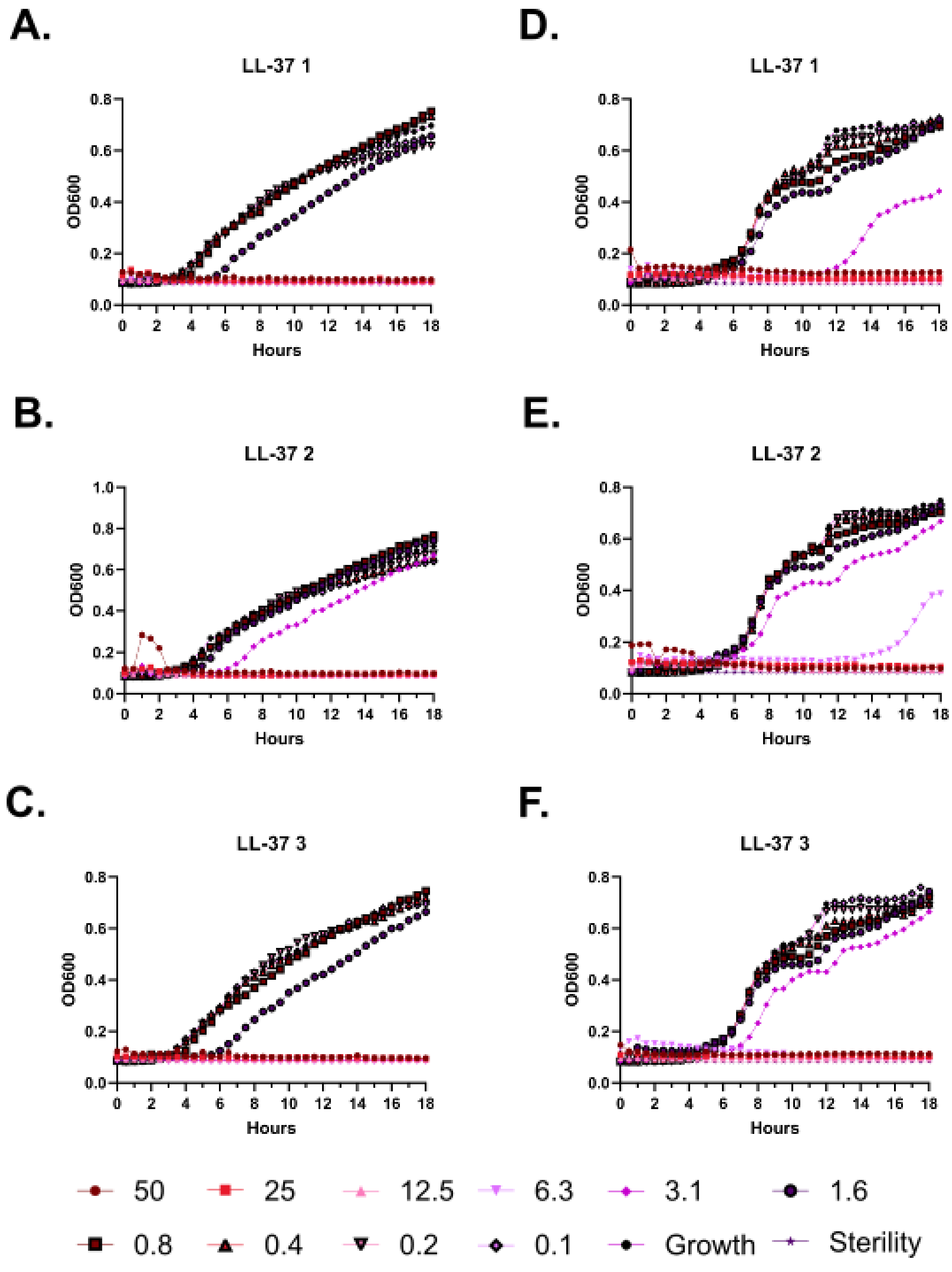

A.
LL-37 1
B.
LL-37 2
C.
LL-37 3
D.
LL-37 1
E.
LL-37 2
F.
LL-37 3
OD600
Hours
50
25
12.5
6.3
3.1
1.6
0.8
0.4
0.2
0.1
Growth
Sterility

**Supplementary Figure 19 – Kinetic antimicrobial susceptibility testing for (LL-37). A-C.** *E. coli*. **D-F.** *Pseudomonas.* Bacterial growth was monitored by OD600 over 18 hours at 37 °C in the presence of the indicated peptide concentrations. Increasing OD600 indicates increasing bacterial growth. Peptide concentrations are given in µM. Each curve indicates an independent experiment.

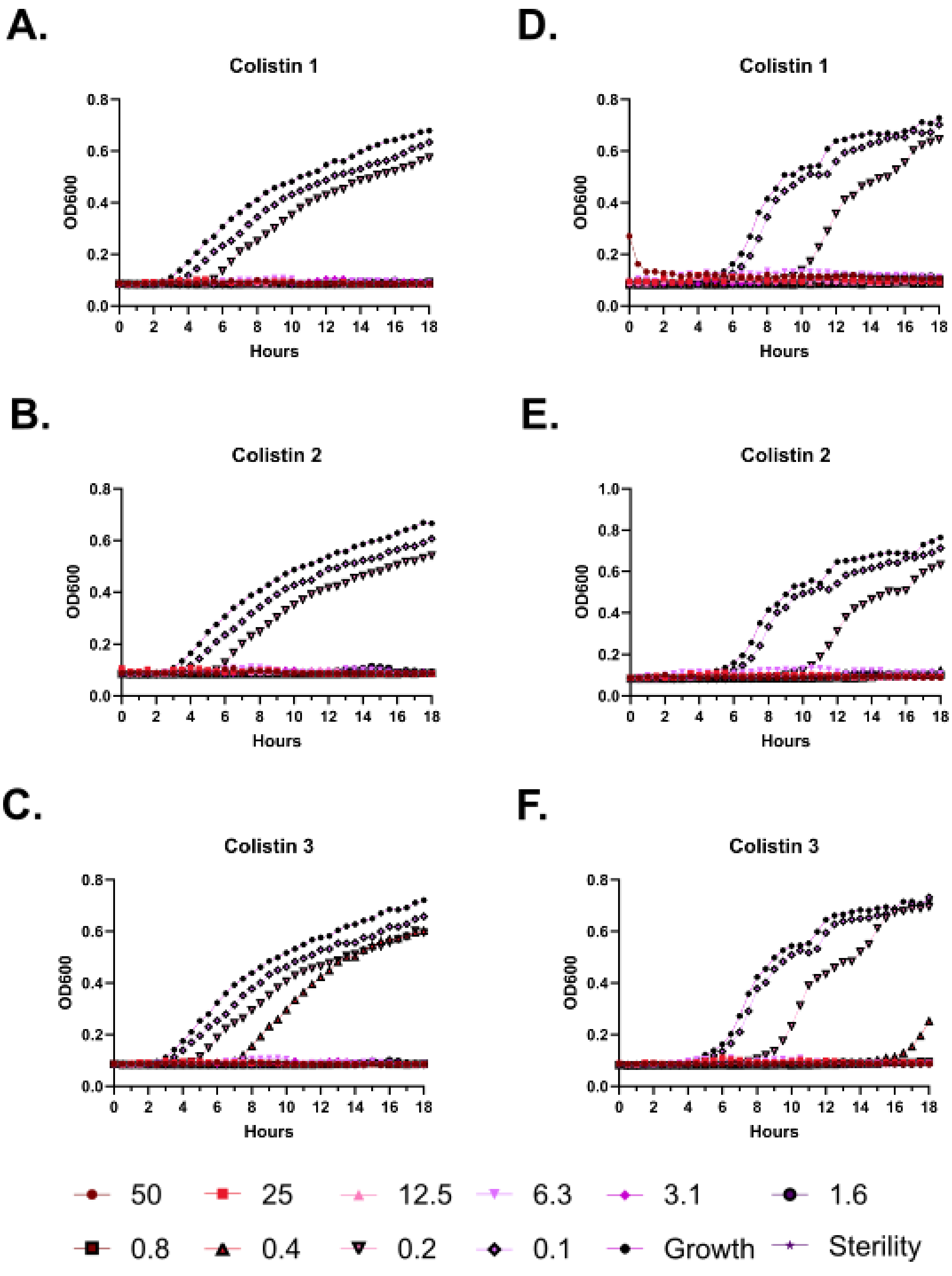

A.
Colistin 1
OD600
Hours
B.
Colistin 2
C.
Colistin 3
D.
Colistin 1
E.
Colistin 2
F.
Colistin 3
50
25
12.5
6.3
3.1
1.6
0.8
0.4
0.2
0.1
Growth
Sterility

**Supplementary Figure 20 – Kinetic antimicrobial susceptibility testing for (Colistin). A-C.** *E. coli*. **D-F.** *Pseudomonas.* Bacterial growth was monitored by OD600 over 18 hours at 37 °C in the presence of the indicated peptide concentrations. Increasing OD600 indicates increasing bacterial growth. Peptide concentrations are given in µM. Each curve indicates an independent experiment.

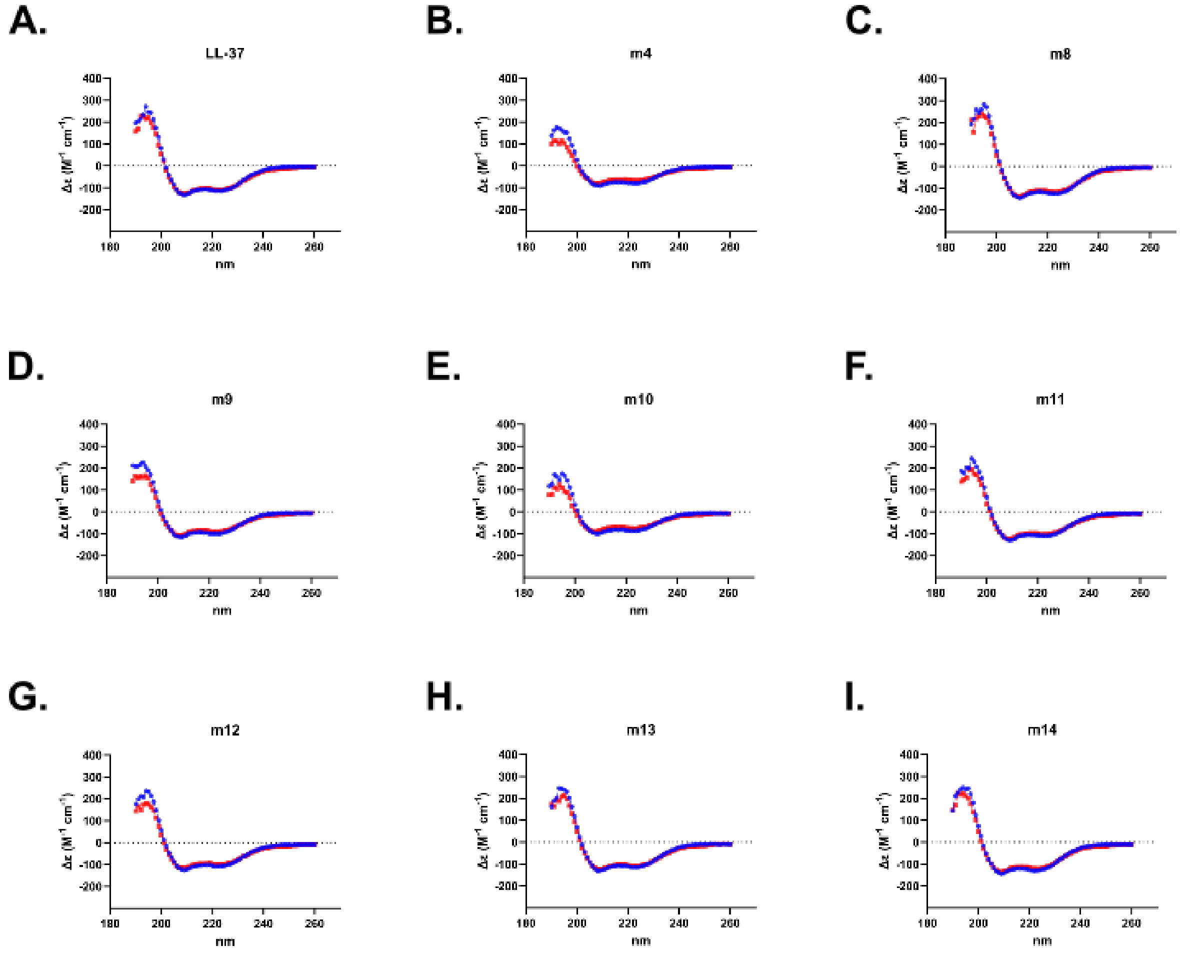


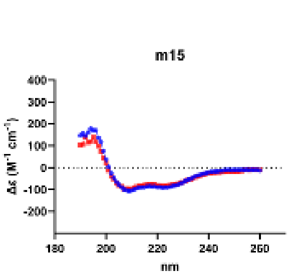


K.

| | % Helix | |
|---|---|---|
| | 20 °C | 37 °C |
| LL-37 | 100 | 100 |
| LL-37 m4 | 100 | 100 |
| LL-37 m8 | 100 | 100 |
| LL-37 m9 | 100 | 100 |
| LL-37 m10 | 100 | 100 |
| LL-37 m11 | 100 | 100 |
| LL-37 m12 | 100 | 100 |
| LL-37 m13 | 100 | 100 |
| LL-37 m14 | 100 | 100 |
| LL-37 m15 | 100 | 100 |

**Supplementary Figure 21 – Circular dichroism (CD) spectroscopy of Figure 2 mutants.** **A-J.** CD spectra are displayed for each mutant dissolved in water plus 20% trifluoroethanol. Data were collected from 190-260 nm at both 20 °C (Blue) and 37 °C (Red). **K.** Quantification of helical content as determined by BeStSel for each mutant.

A.

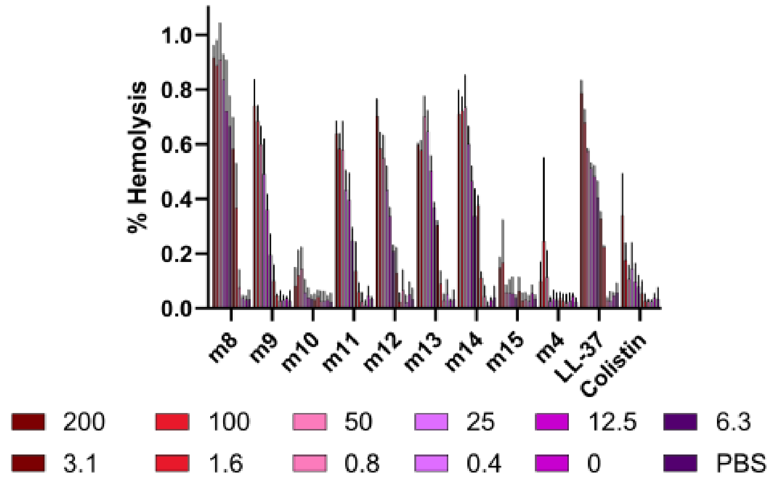
% Hemolysis
1.0
0.8
0.6
0.4
0.2
0.0
m8
m9
m10
m11
m12
m13
m14
m15
m4
LL-37
Colistin
200
100
50
25
12.5
6.3
3.1
1.6
0.8
0.4
0
PBS

B.

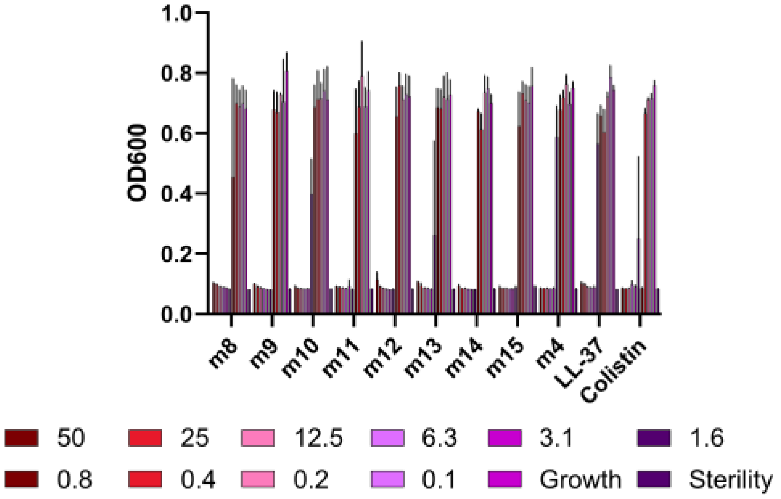
OD600
1.0
0.8
0.6
0.4
0.2
0.0
m8
m9
m10
m11
m12
m13
m14
m15
m4
LL-37
Colistin
50
25
12.5
6.3
3.1
1.6
0.8
0.4
0.2
0.1
Growth
Sterility

C.

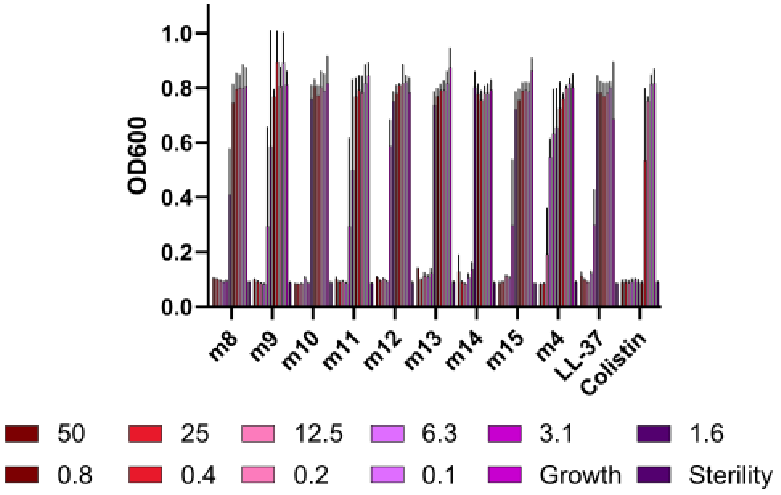
OD600
1.0
0.8
0.6
0.4
0.2
0.0
m8
m9
m10
m11
m12
m13
m14
m15
m4
LL-37
Colistin
50
25
12.5
6.3
3.1
1.6
0.8
0.4
0.2
0.1
Growth
Sterility

**Supplementary Figure 22 – Histogram depiction of Figure 2 data. A.** Endpoint hemolytic activity of each peptide relative to a 100% hemolysis freeze-thaw control after 18 hours at 37 °C. Raw data were obtained by absorbance at 414 nm in well supernatants for free hemoglobin signal. **B-C.** Endpoint activity of each peptide against *E. coli* (**B**) or *Pseudomonas* (**C**). Data are the raw OD600 after 18 hours at 37 °C. Peptide concentrations are given in μM. Data throughout show the mean and standard deviation of three independent experiments.

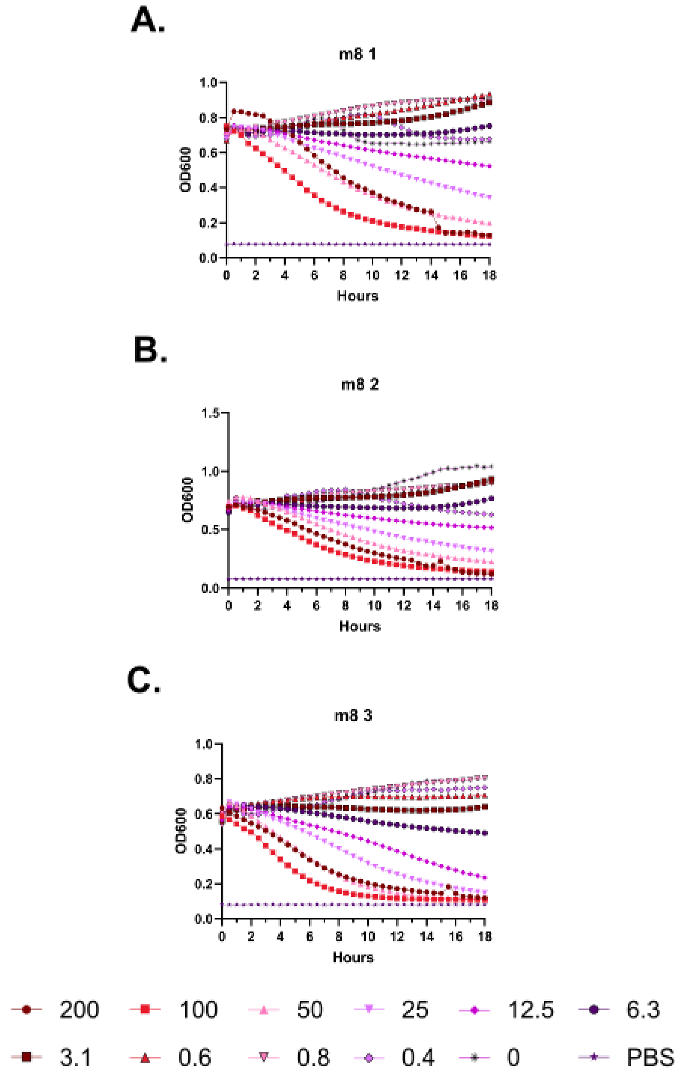
A.
m8 1
OD600
Hours
B.
m8 2
OD600
Hours
C.
m8 3
OD600
Hours
200
100
50
25
12.5
6.3
3.1
0.6
0.8
0.4
0
PBS

**Supplementary Figure 23 – Kinetic hemolysis assays for (m8). A-C.** Hemolysis was monitored by OD600 over 18 hours at 37 °C in the presence of the indicated peptide concentrations. Decreasing OD600 indicates increasing hemolysis. Peptide concentrations are given in μM. Each curve indicates an independent experiment.

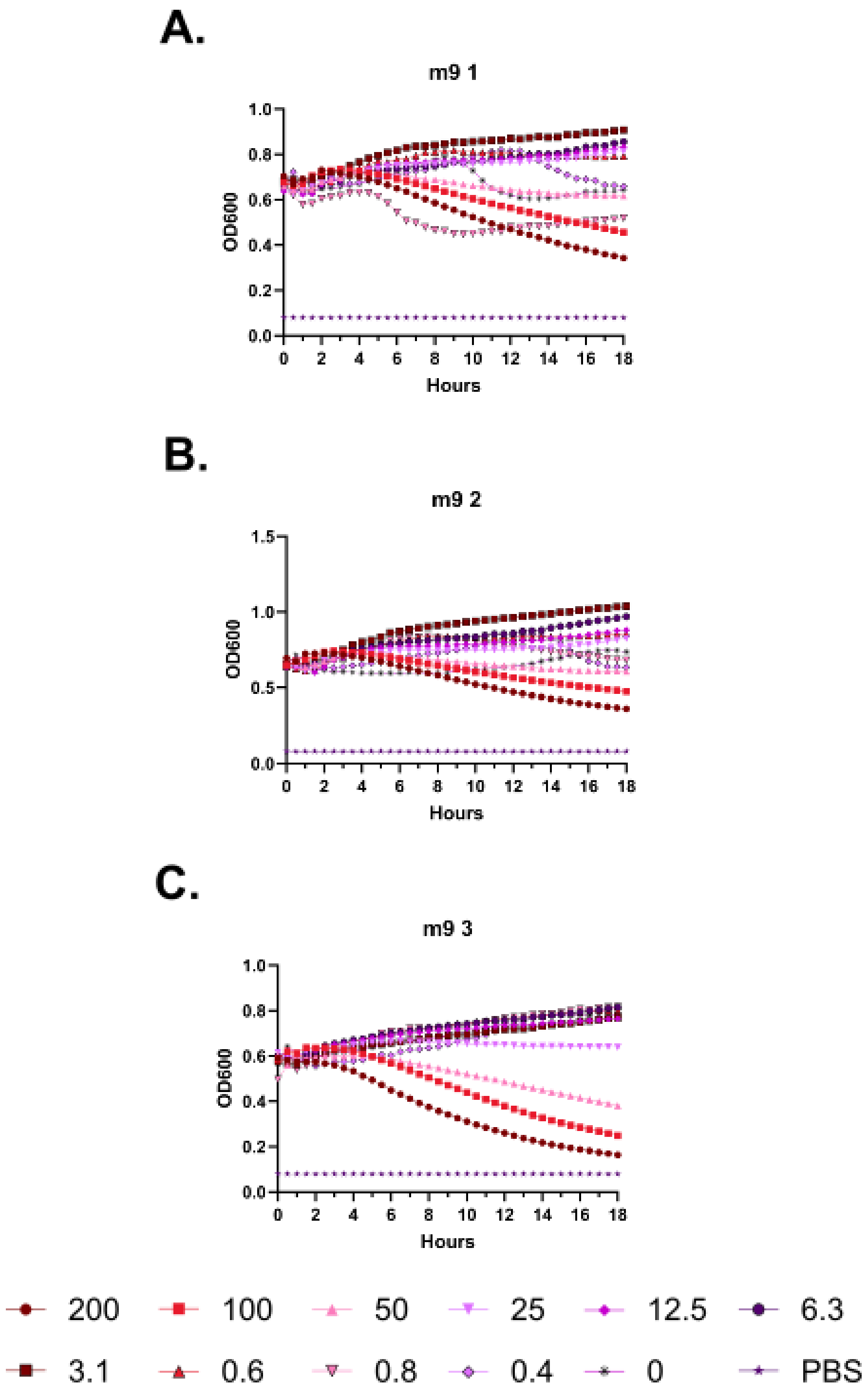
A.
m9 1
OD600
1.0
0.8
0.6
0.4
0.2
0.0
0 2 4 6 8 10 12 14 16 18
Hours
B.
m9 2
OD600
1.5
1.0
0.5
0.0
0 2 4 6 8 10 12 14 16 18
Hours
C.
m9 3
OD600
1.0
0.8
0.6
0.4
0.2
0.0
0 2 4 6 8 10 12 14 16 18
Hours
200
100
50
25
12.5
6.3
3.1
0.6
0.8
0.4
0
PBS

**Supplementary Figure 24 – Kinetic hemolysis assays for (m9). A-C.** Hemolysis was monitored by OD600 over 18 hours at 37 °C in the presence of the indicated peptide concentrations. Decreasing OD600 indicates increasing hemolysis. Peptide concentrations are given in μM. Each curve indicates an independent experiment.

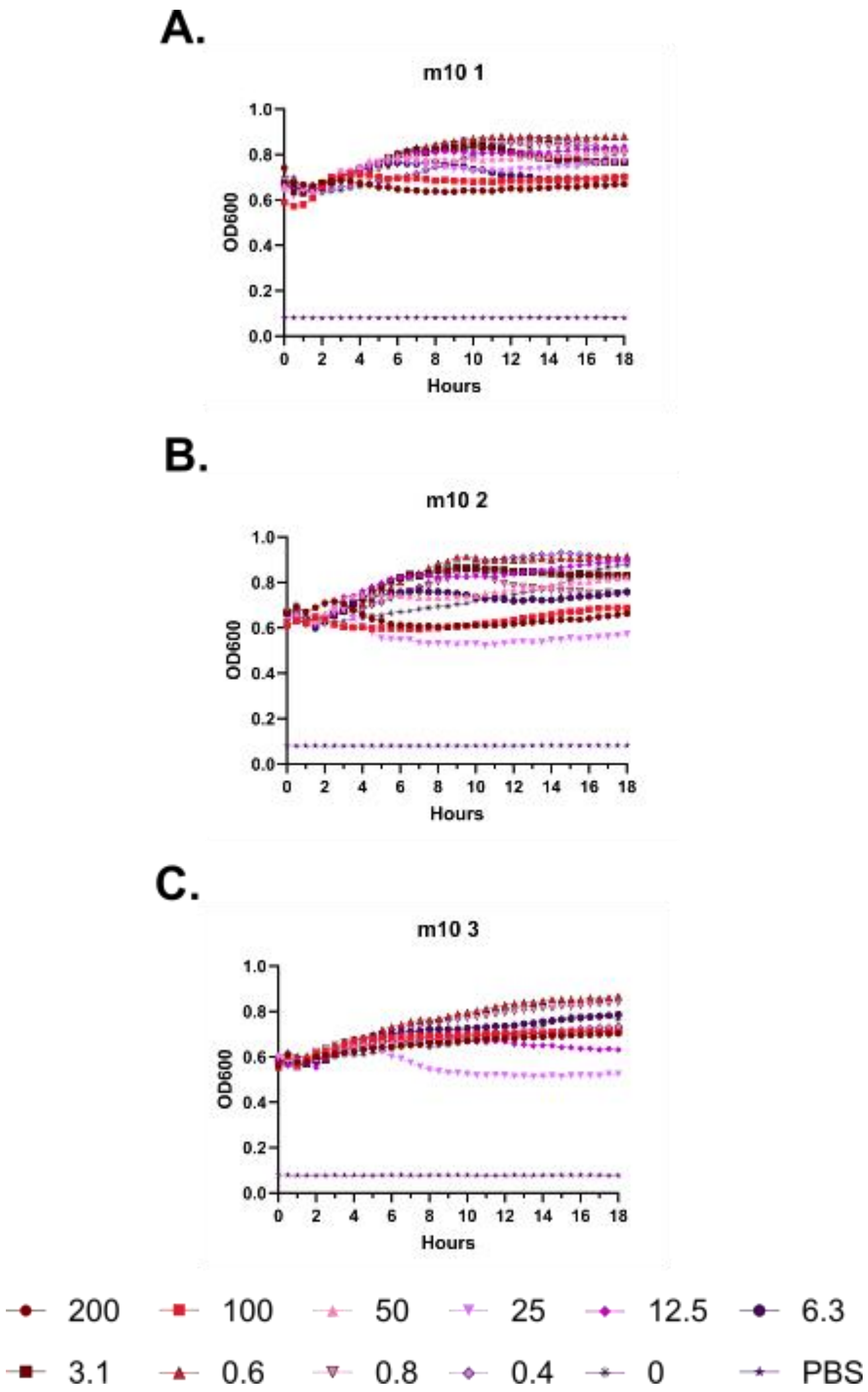
A.
m10 1
OD600
Hours
B.
m10 2
OD600
Hours
C.
m10 3
OD600
Hours
1.0
0.8
0.6
0.4
0.2
0.0
0 2 4 6 8 10 12 14 16 18
200
100
50
25
12.5
6.3
3.1
0.6
0.8
0.4
0
PBS

**Supplementary Figure 25 – Kinetic hemolysis assays for (m10). A-C.** Hemolysis was monitored by OD600 over 18 hours at 37 °C in the presence of the indicated peptide concentrations. Decreasing OD600 indicates increasing hemolysis. Peptide concentrations are given in μM. Each curve indicates an independent experiment.

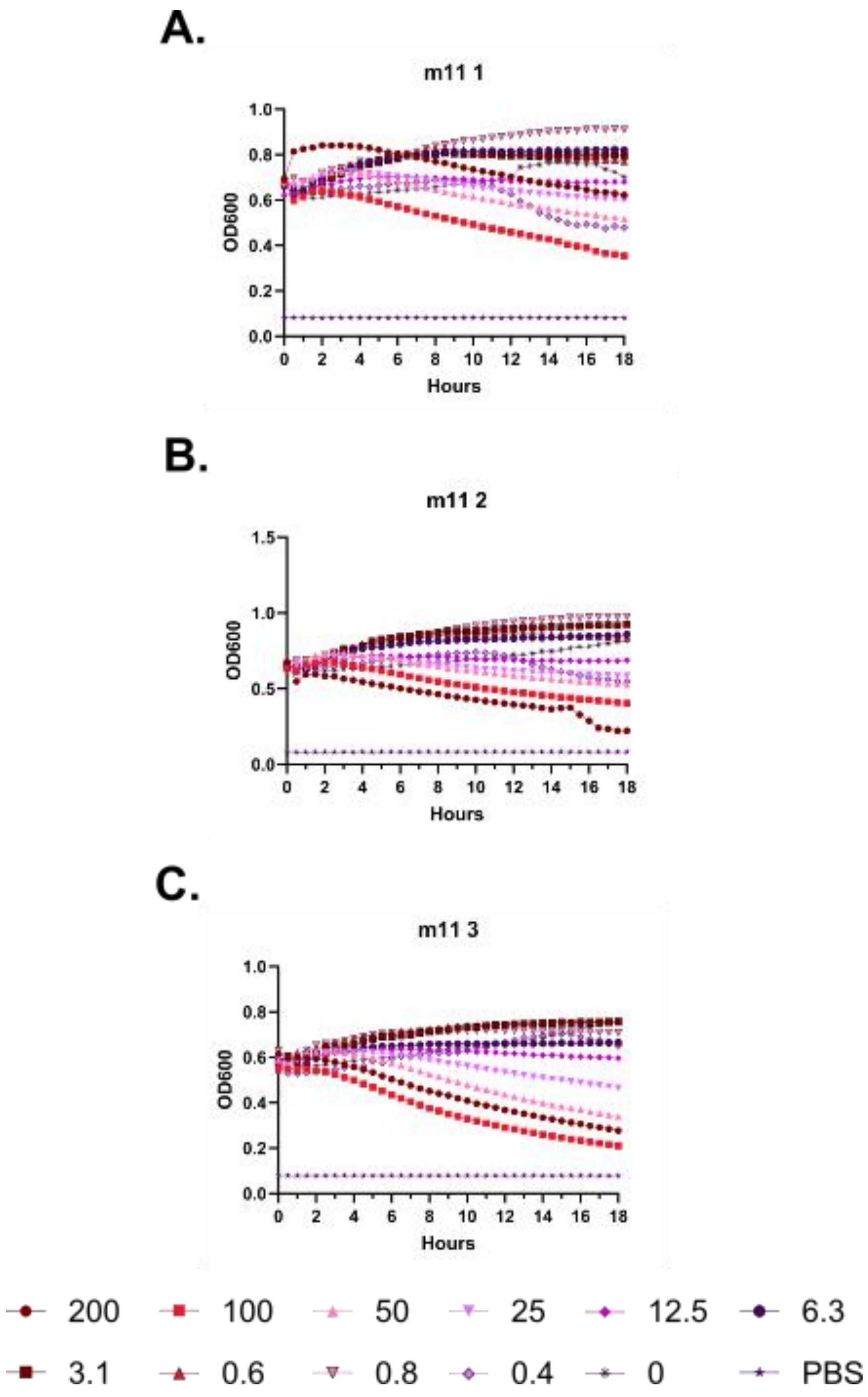
A.
m11 1
OD600
Hours
B.
m11 2
OD600
Hours
C.
m11 3
OD600
Hours
200
100
50
25
12.5
6.3
3.1
0.6
0.8
0.4
0
PBS

**Supplementary Figure 26 – Kinetic hemolysis assays for (m11). A-C.** Hemolysis was monitored by OD600 over 18 hours at 37 °C in the presence of the indicated peptide concentrations. Decreasing OD600 indicates increasing hemolysis. Peptide concentrations are given in µM. Each curve indicates an independent experiment.

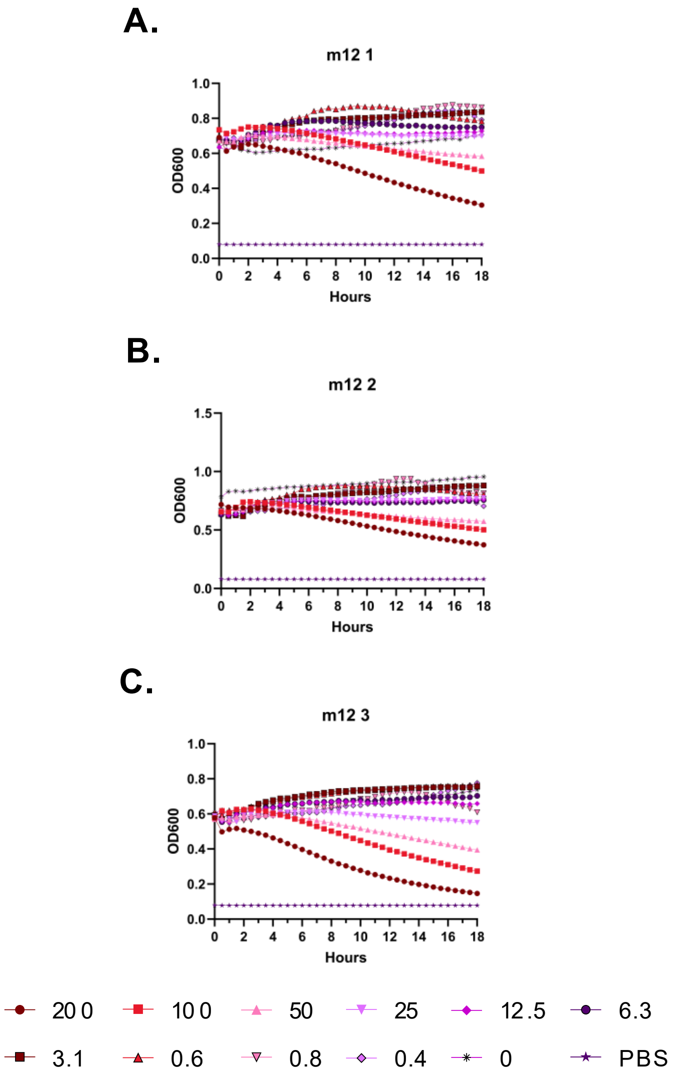
A.
m12 1
OD600
Hours
B.
m12 2
OD600
Hours
C.
m12 3
OD600
Hours
20 0
10 0
50
25
12.5
6.3
3.1
0.6
0.8
0.4
0
PBS

**Supplementary Figure 27 – Kinetic hemolysis assays for (m12). A-C.** Hemolysis was monitored by OD600 over 18 hours at 37 °C in the presence of the indicated peptide concentrations. Decreasing OD600 indicates increasing hemolysis. Peptide concentrations are given in μM. Each curve indicates an independent experiment.

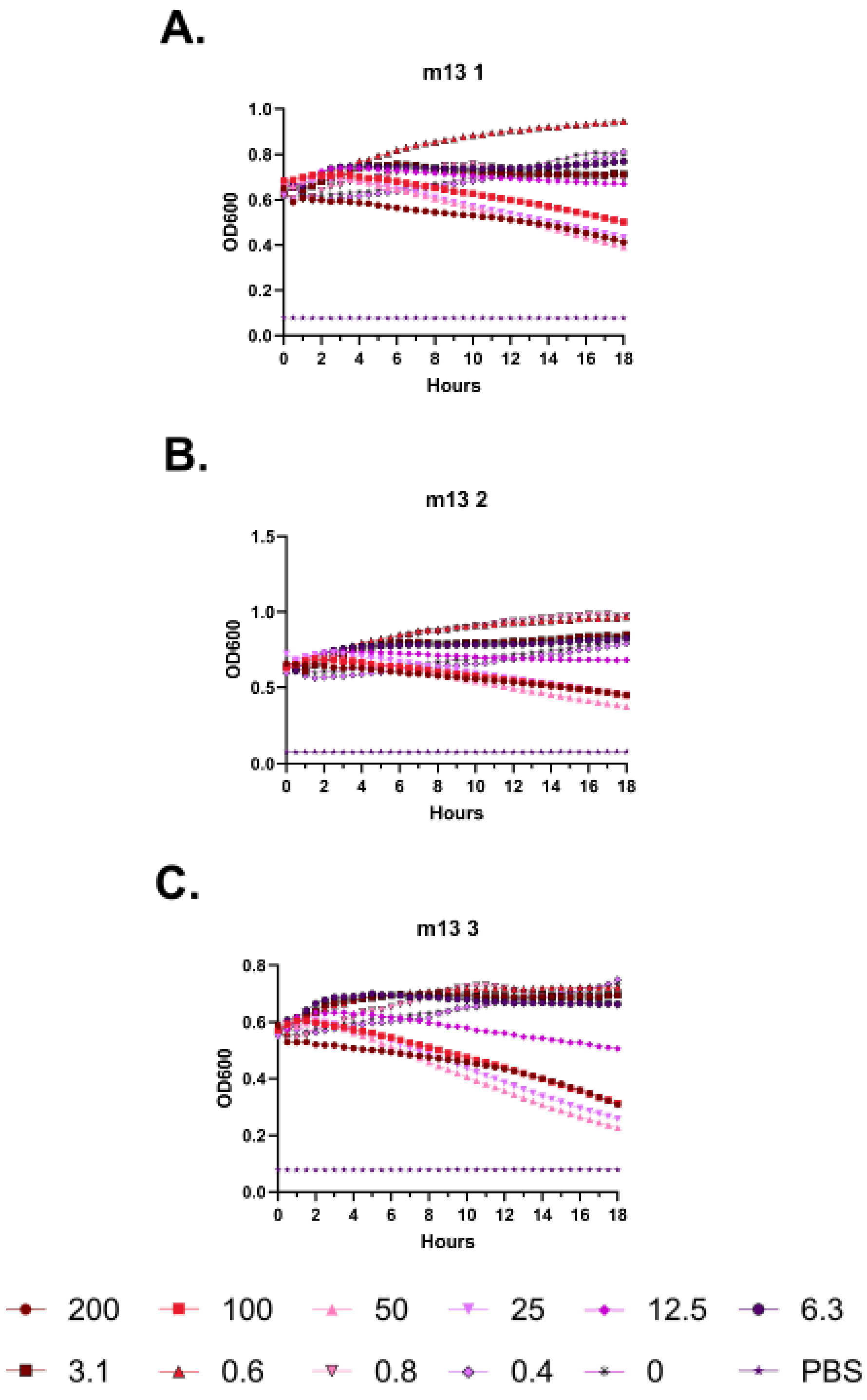
A.
m13 1
OD600
Hours
B.
m13 2
OD600
Hours
C.
m13 3
OD600
Hours
200
100
50
25
12.5
6.3
3.1
0.6
0.8
0.4
0
PBS

**Supplementary Figure 28 – Kinetic hemolysis assays for (m13). A-C.** Hemolysis was monitored by OD600 over 18 hours at 37 °C in the presence of the indicated peptide concentrations. Decreasing OD600 indicates increasing hemolysis. Peptide concentrations are given in μM. Each curve indicates an independent experiment.

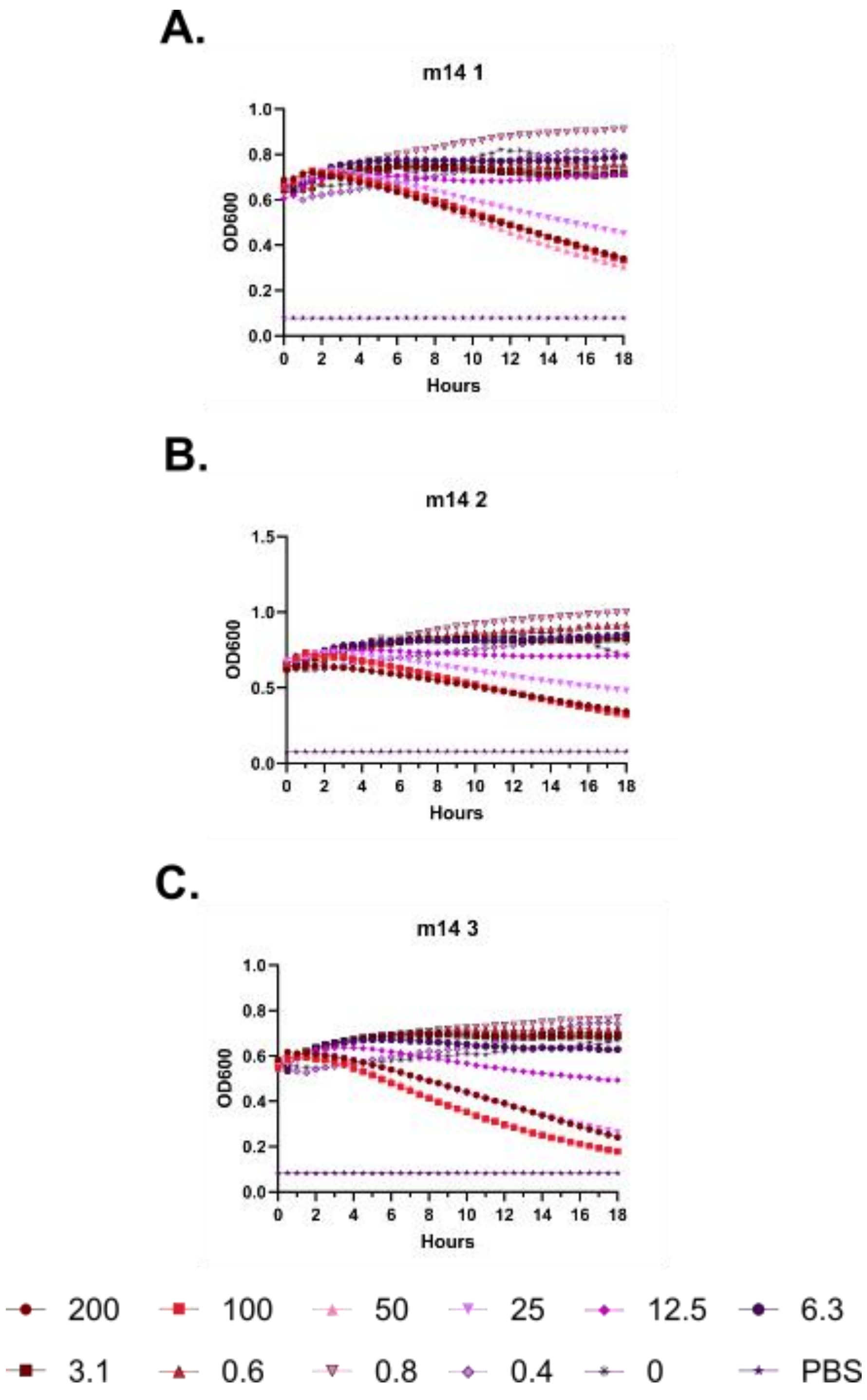
A.
m14 1
OD600
Hours
B.
m14 2
OD600
Hours
C.
m14 3
OD600
Hours
200
100
50
25
12.5
6.3
3.1
0.6
0.8
0.4
0
PBS

**Supplementary Figure 29 – Kinetic hemolysis assays for (m14). A-C.** Hemolysis was monitored by OD600 over 18 hours at 37 °C in the presence of the indicated peptide concentrations. Decreasing OD600 indicates increasing hemolysis. Peptide concentrations are given in µM. Each curve indicates an independent experiment.

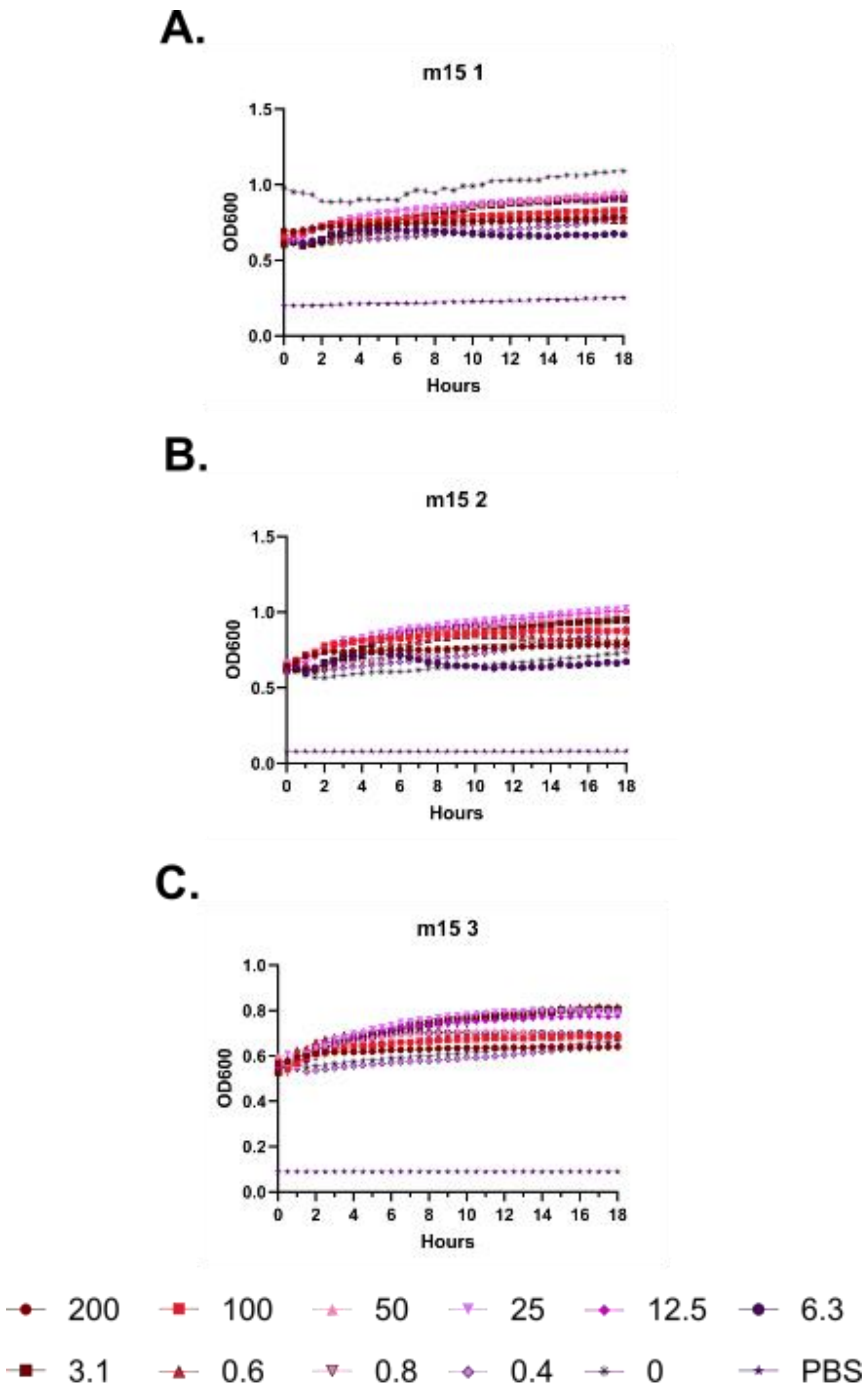
A.
m15 1
OD600
Hours
1.5
1.0
0.5
0.0
0 2 4 6 8 10 12 14 16 18
B.
m15 2
OD600
Hours
1.5
1.0
0.5
0.0
0 2 4 6 8 10 12 14 16 18
C.
m15 3
OD600
Hours
1.0
0.8
0.6
0.4
0.2
0.0
0 2 4 6 8 10 12 14 16 18
200
100
50
25
12.5
6.3
3.1
0.6
0.8
0.4
0
PBS

**Supplementary Figure 30 – Kinetic hemolysis assays for (m15). A-C.** Hemolysis was monitored by OD600 over 18 hours at 37 °C in the presence of the indicated peptide concentrations. Decreasing OD600 indicates increasing hemolysis. Peptide concentrations are given in µM. Each curve indicates an independent experiment.

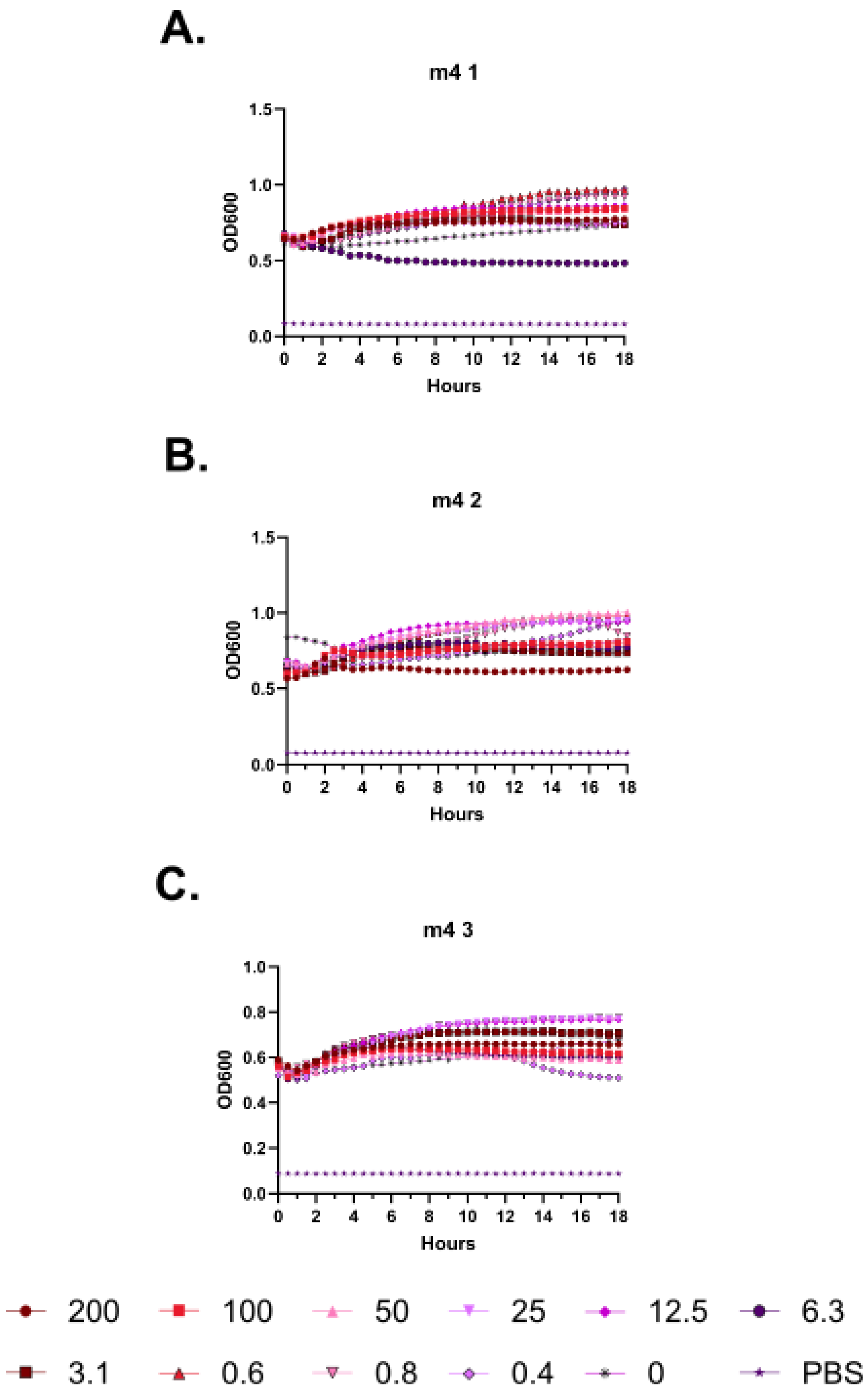
A.
m4 1
OD600
Hours
B.
m4 2
OD600
Hours
C.
m4 3
OD600
Hours
200
100
50
25
12.5
6.3
3.1
0.6
0.8
0.4
0
PBS

**Supplementary Figure 31 – Kinetic hemolysis assays for (m4). A-C.** Hemolysis was monitored by OD600 over 18 hours at 37 °C in the presence of the indicated peptide concentrations. Decreasing OD600 indicates increasing hemolysis. Peptide concentrations are given in μM. Each curve indicates an independent experiment.

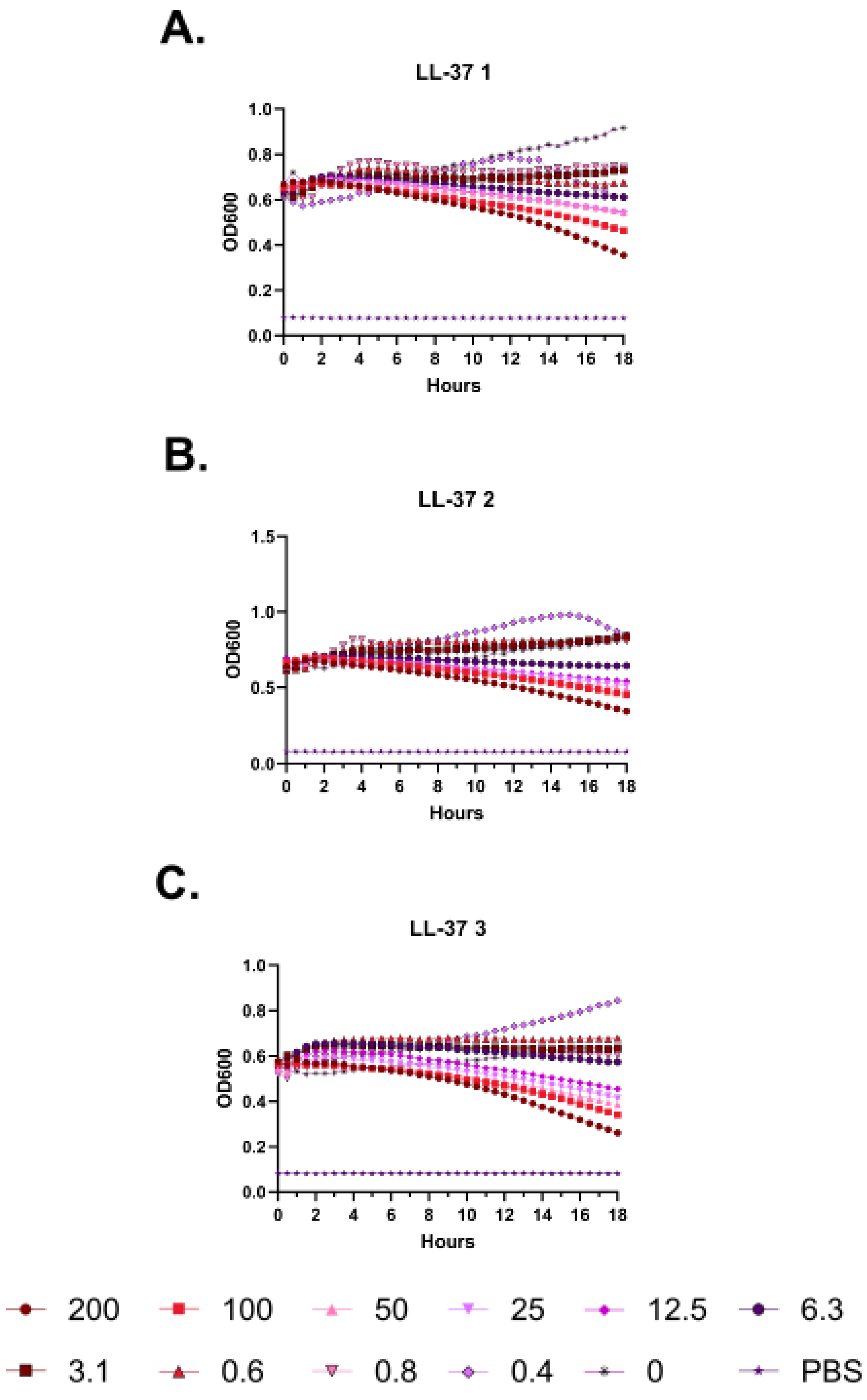
A.
LL-37 1
OD600
Hours
B.
LL-37 2
OD600
Hours
C.
LL-37 3
OD600
Hours
200
100
50
25
12.5
6.3
3.1
0.6
0.8
0.4
0
PBS

**Supplementary Figure 32 – Kinetic hemolysis assays for (LL-37). A-C.** Hemolysis was monitored by OD600 over 18 hours at 37 °C in the presence of the indicated peptide concentrations. Decreasing OD600 indicates increasing hemolysis. Peptide concentrations are given in μM. Each curve indicates an independent experiment.

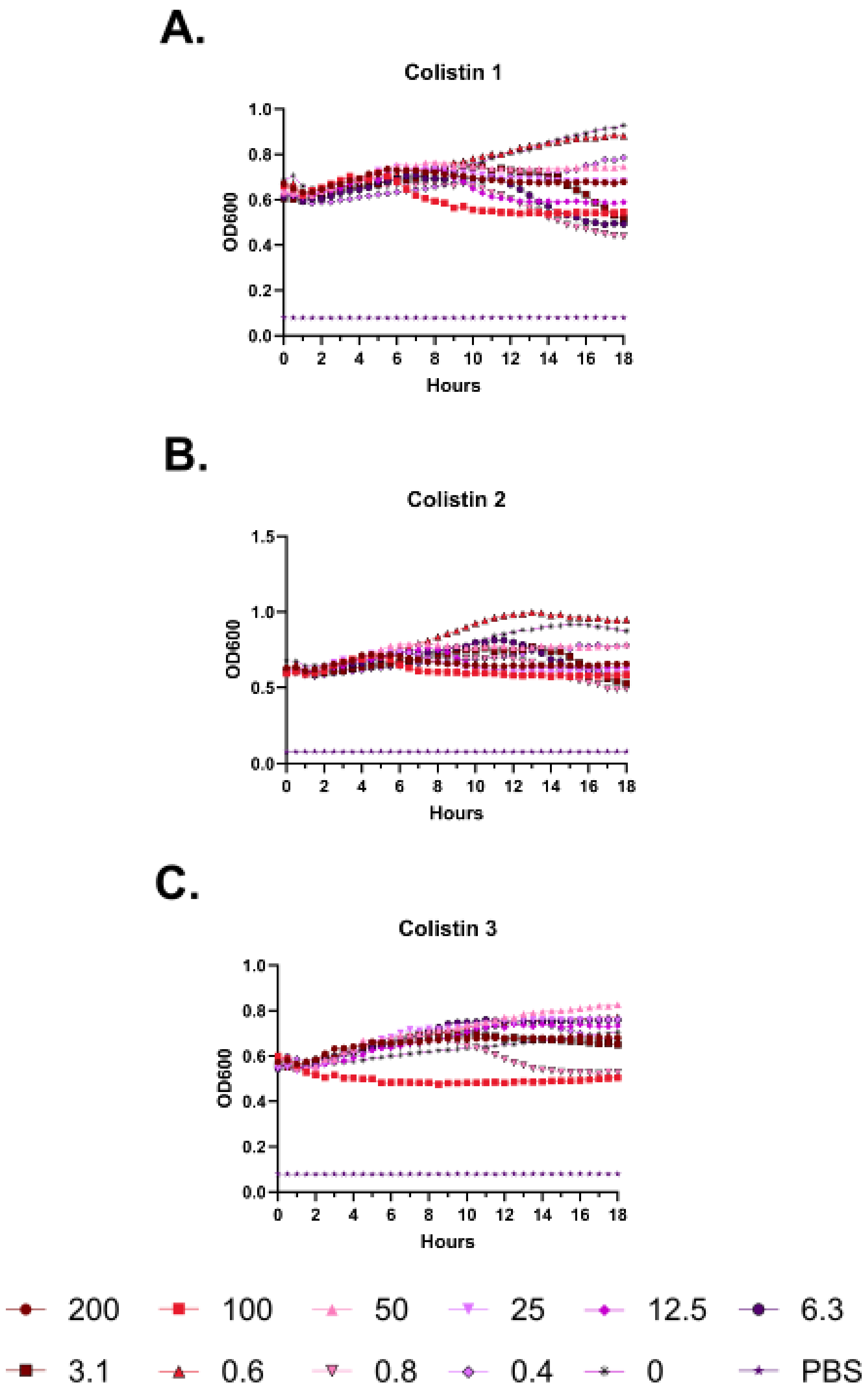
A.
Colistin 1
OD600
Hours
B.
Colistin 2
OD600
Hours
C.
Colistin 3
OD600
Hours
200
100
50
25
12.5
6.3
3.1
0.6
0.8
0.4
0
PBS

**Supplementary Figure 33 – Kinetic hemolysis assays for (Colistin). A-C.** Hemolysis was monitored by OD600 over 18 hours at 37 °C in the presence of the indicated peptide concentrations. Decreasing OD600 indicates increasing hemolysis. Peptide concentrations are given in µM. Each curve indicates an independent experiment.

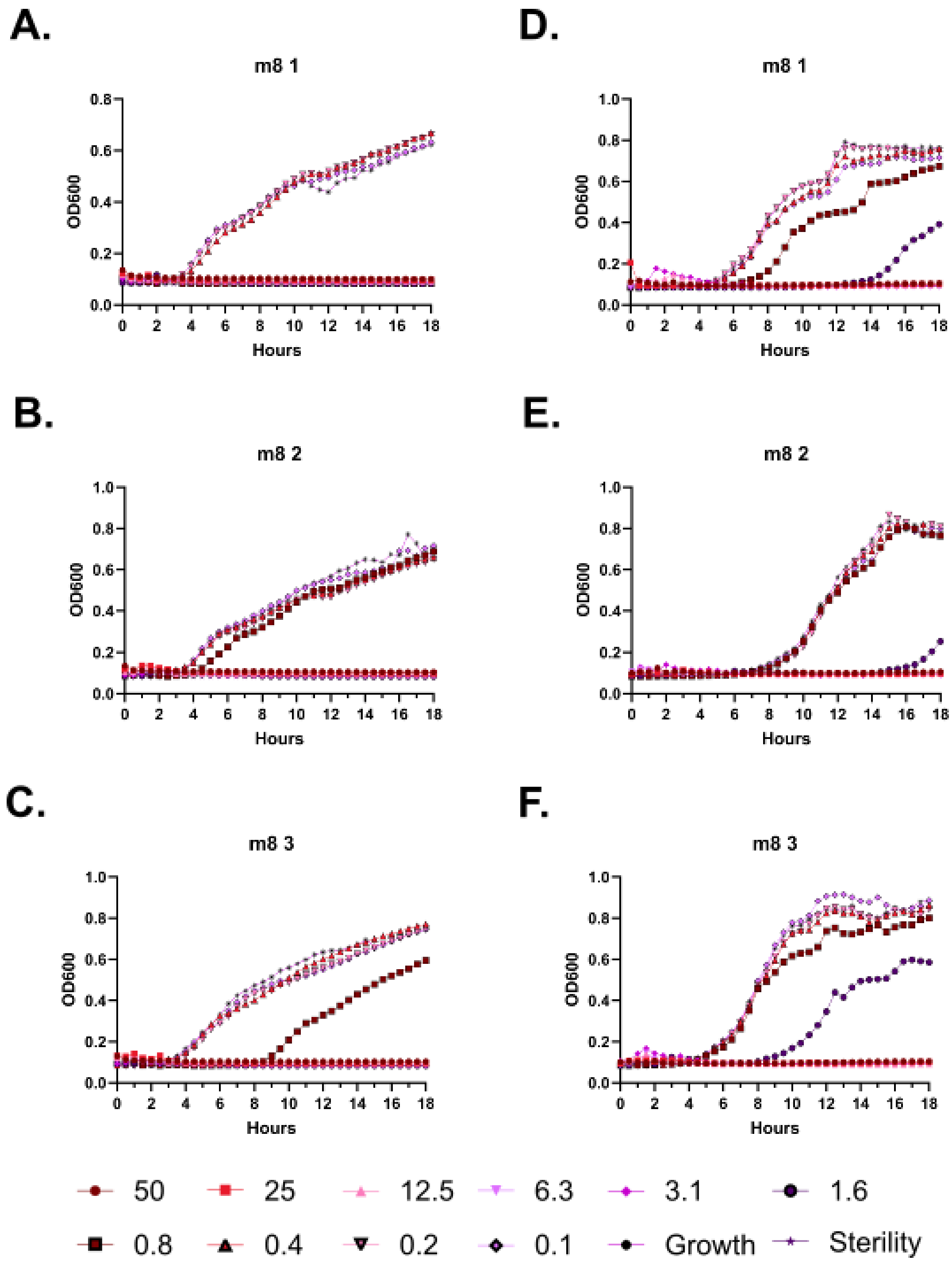
A.
m8 1
OD600
Hours
D.
m8 1
OD600
Hours
B.
m8 2
OD600
Hours
E.
m8 2
OD600
Hours
C.
m8 3
OD600
Hours
F.
m8 3
OD600
Hours
50
25
12.5
6.3
3.1
1.6
0.8
0.4
0.2
0.1
Growth
Sterility

**Supplementary Figure 34 – Kinetic antimicrobial susceptibility testing for (m8). A-C.** *E. coli*. **D-F.** *Pseudomonas.* Bacterial growth was monitored by OD600 over 18 hours at 37 °C in the presence of the indicated peptide concentrations. Increasing OD600 indicates increasing bacterial growth. Peptide concentrations are given in μM. Each curve indicates an independent experiment.

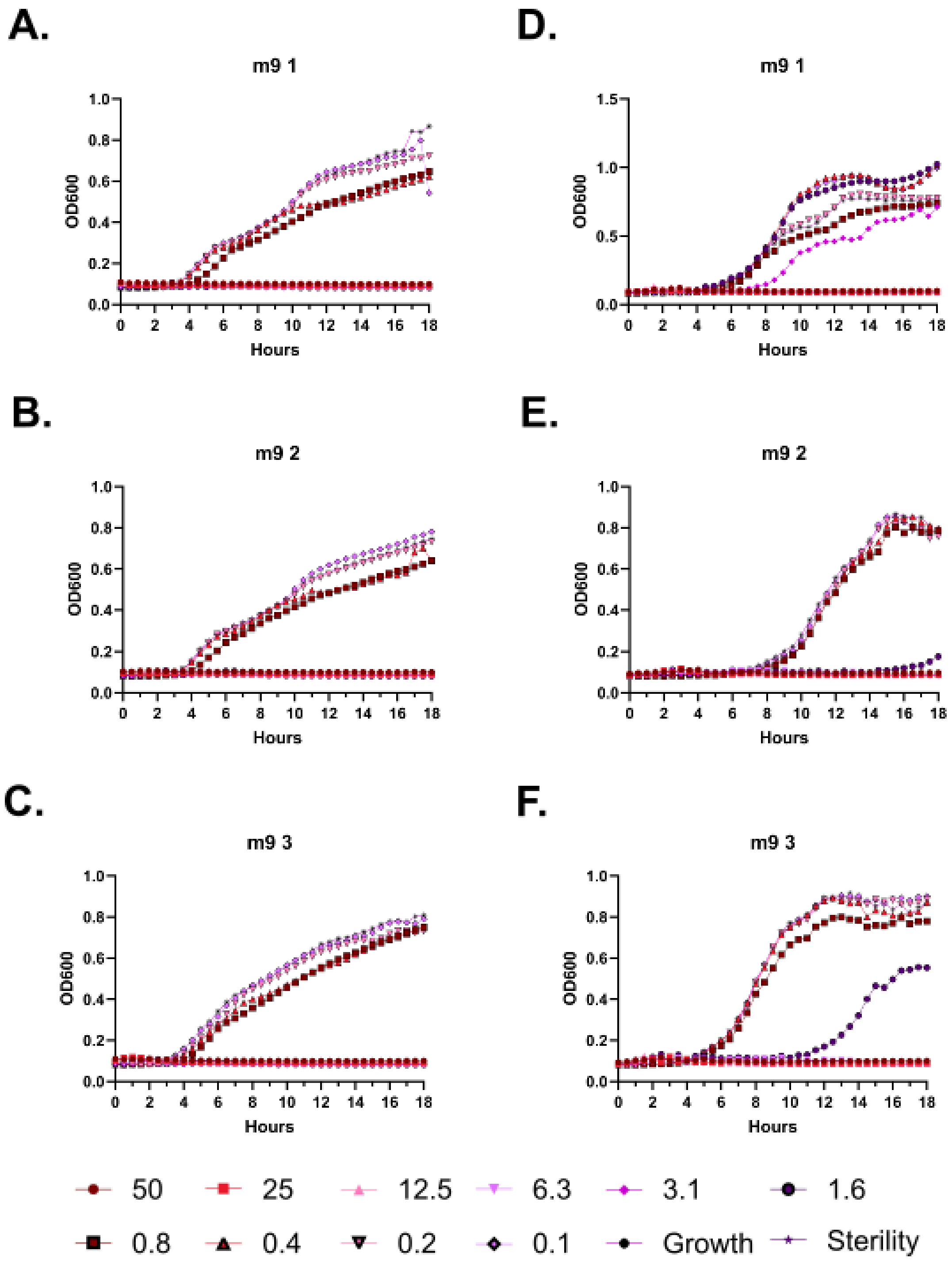
A.
m9 1
OD600
Hours
D.
m9 1
OD600
Hours
B.
m9 2
OD600
Hours
E.
m9 2
OD600
Hours
C.
m9 3
OD600
Hours
F.
m9 3
OD600
Hours
50
25
12.5
6.3
3.1
1.6
0.8
0.4
0.2
0.1
Growth
Sterility

**Supplementary Figure 35 – Kinetic antimicrobial susceptibility testing for (m9). A-C.** *E. coli*. **D-F.** *Pseudomonas.* Bacterial growth was monitored by OD600 over 18 hours at 37 °C in the presence of the indicated peptide concentrations. Increasing OD600 indicates increasing bacterial growth. Peptide concentrations are given in µM. Each curve indicates an independent experiment.

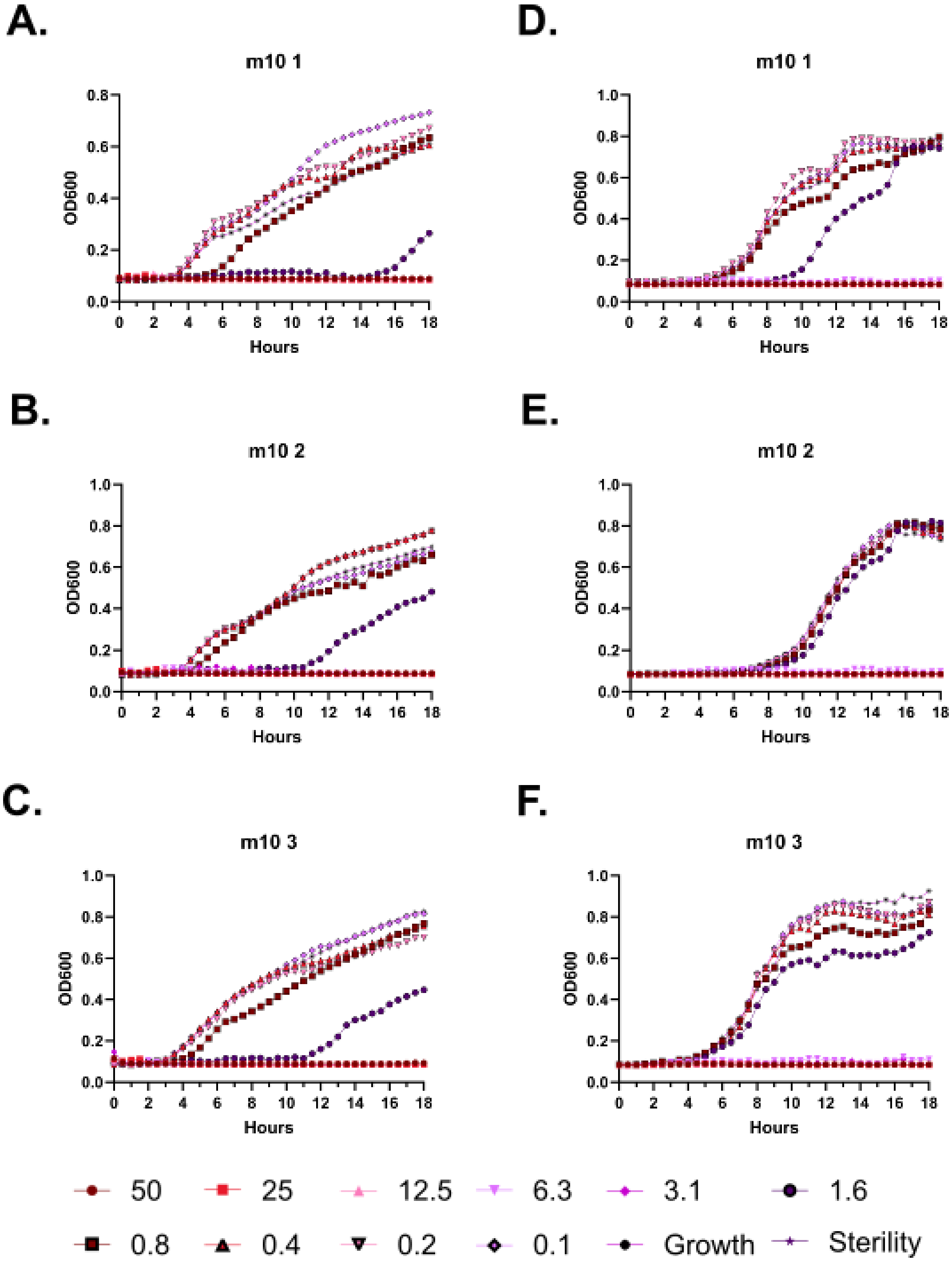
A.
m10 1
OD600
Hours
D.
m10 1
OD600
Hours
B.
m10 2
OD600
Hours
E.
m10 2
OD600
Hours
C.
m10 3
OD600
Hours
F.
m10 3
OD600
Hours
50
25
12.5
6.3
3.1
1.6
0.8
0.4
0.2
0.1
Growth
Sterility

**Supplementary Figure 36 – Kinetic antimicrobial susceptibility testing for (m10). A-C.** *E. coli*. **D-F.** *Pseudomonas.* Bacterial growth was monitored by OD600 over 18 hours at 37 °C in the presence of the indicated peptide concentrations. Increasing OD600 indicates increasing bacterial growth. Peptide concentrations are given in µM. Each curve indicates an independent experiment.

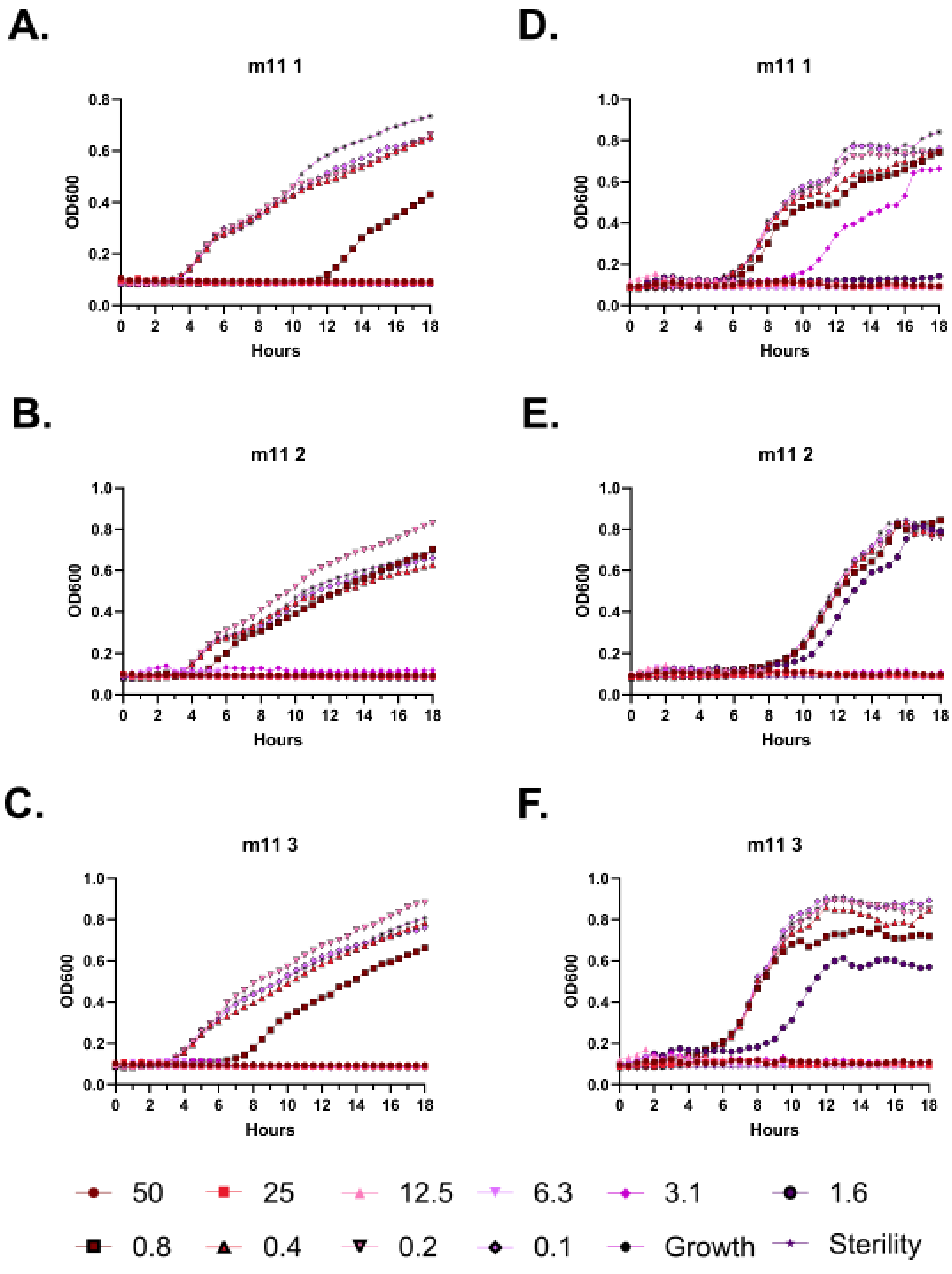
A.
m11 1
OD600
Hours
B.
m11 2
OD600
Hours
C.
m11 3
OD600
Hours
D.
m11 1
OD600
Hours
E.
m11 2
OD600
Hours
F.
m11 3
OD600
Hours
50
25
12.5
6.3
3.1
1.6
0.8
0.4
0.2
0.1
Growth
Sterility

**Supplementary Figure 37 – Kinetic antimicrobial susceptibility testing for (m11). A-C.** *E. coli*. **D-F.** *Pseudomonas.* Bacterial growth was monitored by OD600 over 18 hours at 37 °C in the presence of the indicated peptide concentrations. Increasing OD600 indicates increasing bacterial growth. Peptide concentrations are given in μM. Each curve indicates an independent experiment.

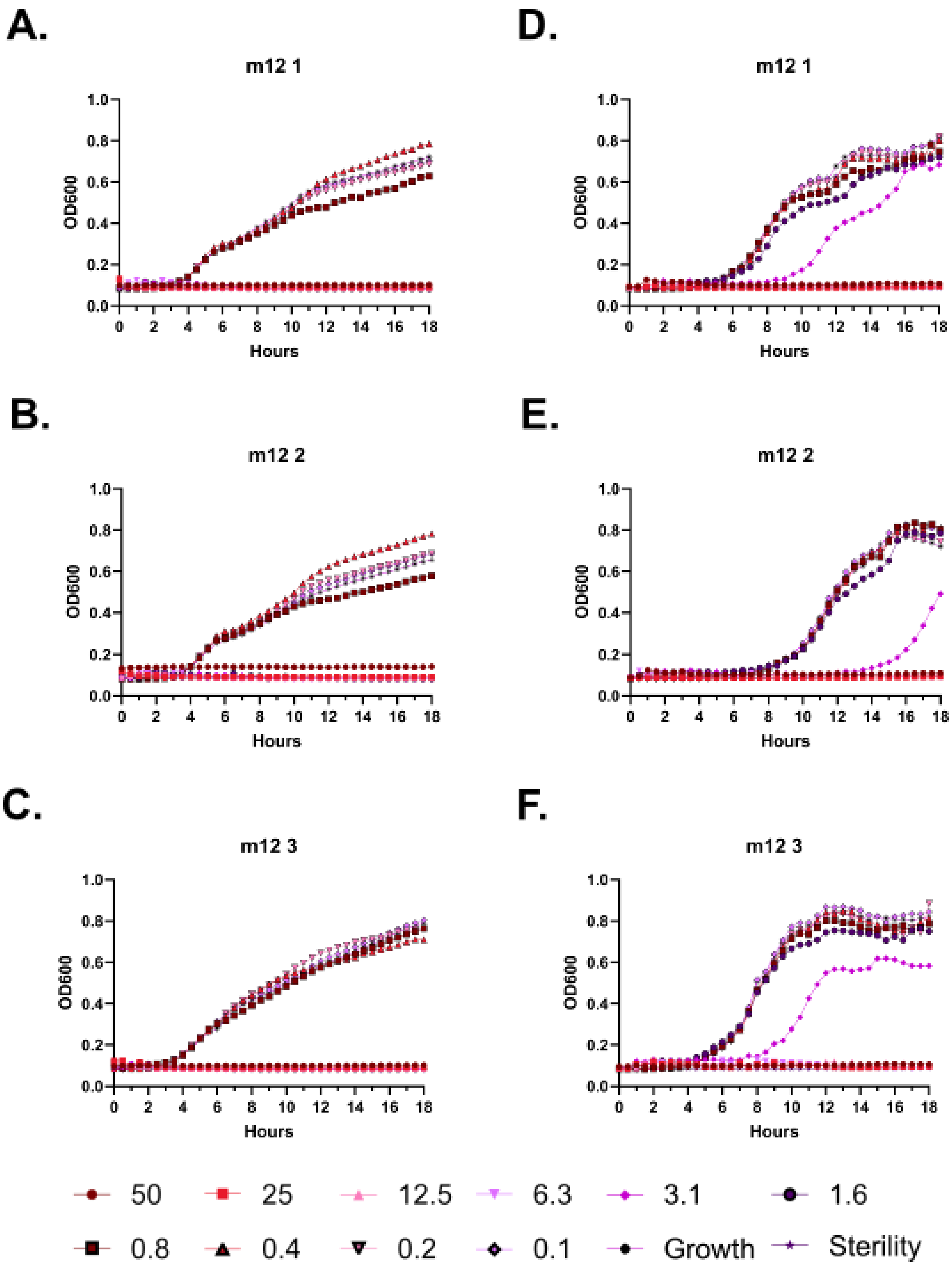
A.
m12 1
B.
m12 2
C.
m12 3
D.
m12 1
E.
m12 2
F.
m12 3
OD600
Hours
0.0
0.2
0.4
0.6
0.8
1.0
0
2
4
6
8
10
12
14
16
18
50
25
12.5
6.3
3.1
1.6
0.8
0.4
0.2
0.1
Growth
Sterility

**Supplementary Figure 38 – Kinetic antimicrobial susceptibility testing for (m12). A-C.** *E. coli*. **D-F.** *Pseudomonas.* Bacterial growth was monitored by OD600 over 18 hours at 37 °C in the presence of the indicated peptide concentrations. Increasing OD600 indicates increasing bacterial growth. Peptide concentrations are given in µM. Each curve indicates an independent experiment.

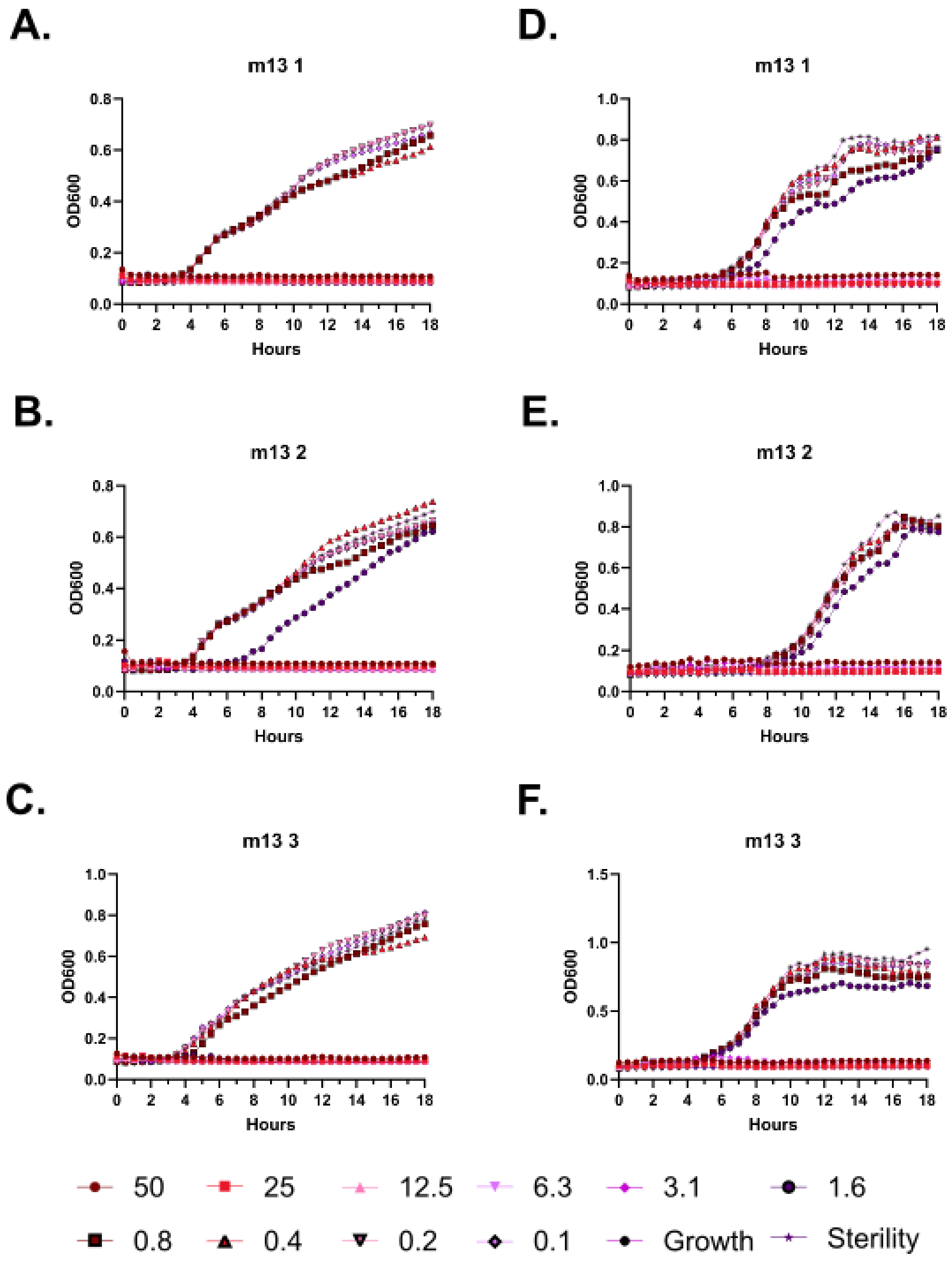
A.
m13 1
OD600
Hours
D.
m13 1
B.
m13 2
E.
m13 2
C.
m13 3
F.
m13 3
50
25
12.5
6.3
3.1
1.6
0.8
0.4
0.2
0.1
Growth
Sterility

**Supplementary Figure 39 – Kinetic antimicrobial susceptibility testing for (m13). A-C.** *E. coli*. **D-F.** *Pseudomonas.* Bacterial growth was monitored by OD600 over 18 hours at 37 °C in the presence of the indicated peptide concentrations. Increasing OD600 indicates increasing bacterial growth. Peptide concentrations are given in µM. Each curve indicates an independent experiment.

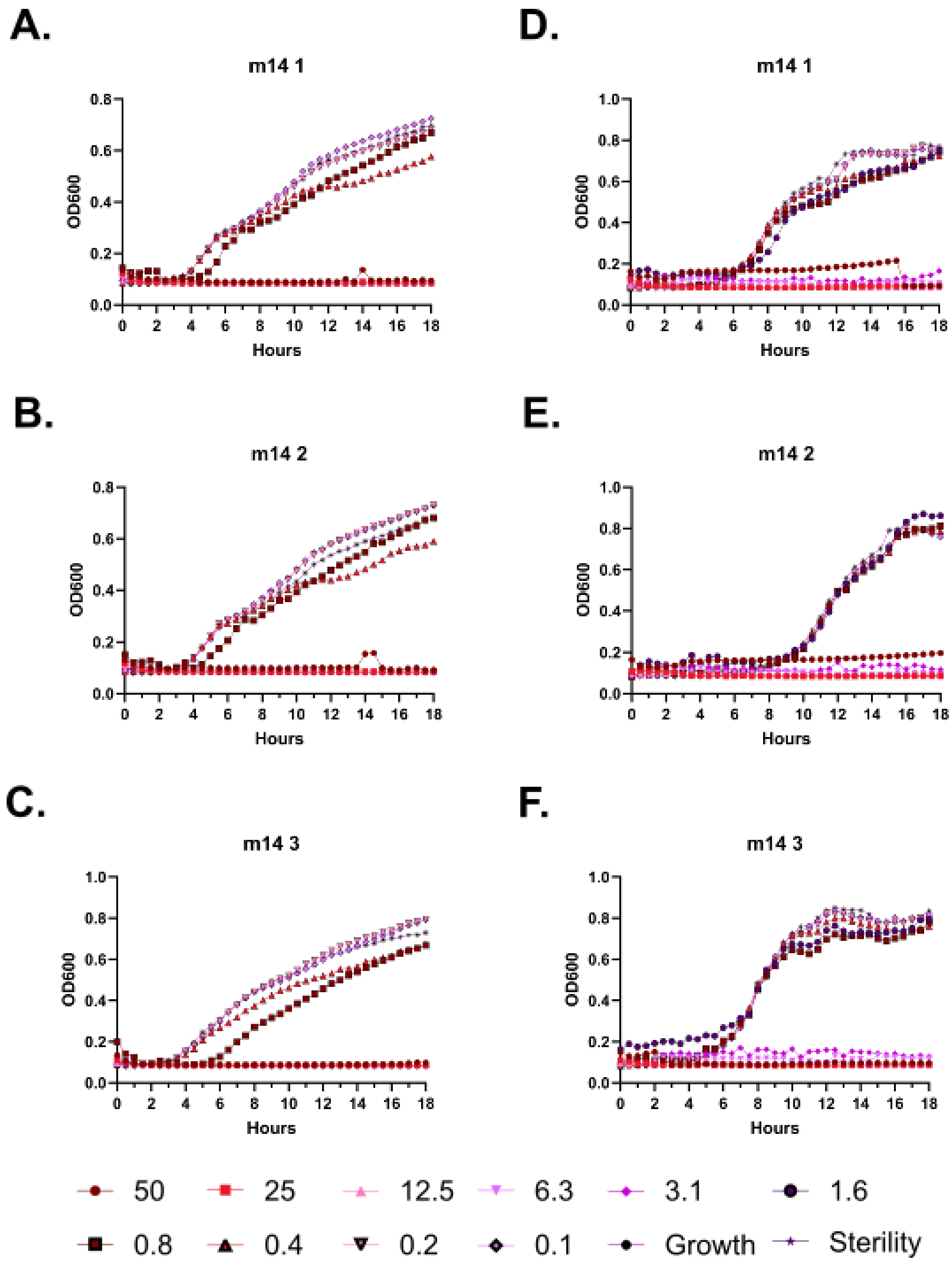

A.
m14 1
OD600
Hours
B.
m14 2
OD600
Hours
C.
m14 3
OD600
Hours
D.
m14 1
OD600
Hours
E.
m14 2
OD600
Hours
F.
m14 3
OD600
Hours
50
25
12.5
6.3
3.1
1.6
0.8
0.4
0.2
0.1
Growth
Sterility

**Supplementary Figure 40 – Kinetic antimicrobial susceptibility testing for (m14). A-C.** *E. coli*. **D-F.** *Pseudomonas.* Bacterial growth was monitored by OD600 over 18 hours at 37 °C in the presence of the indicated peptide concentrations. Increasing OD600 indicates increasing bacterial growth. Peptide concentrations are given in μM. Each curve indicates an independent experiment.

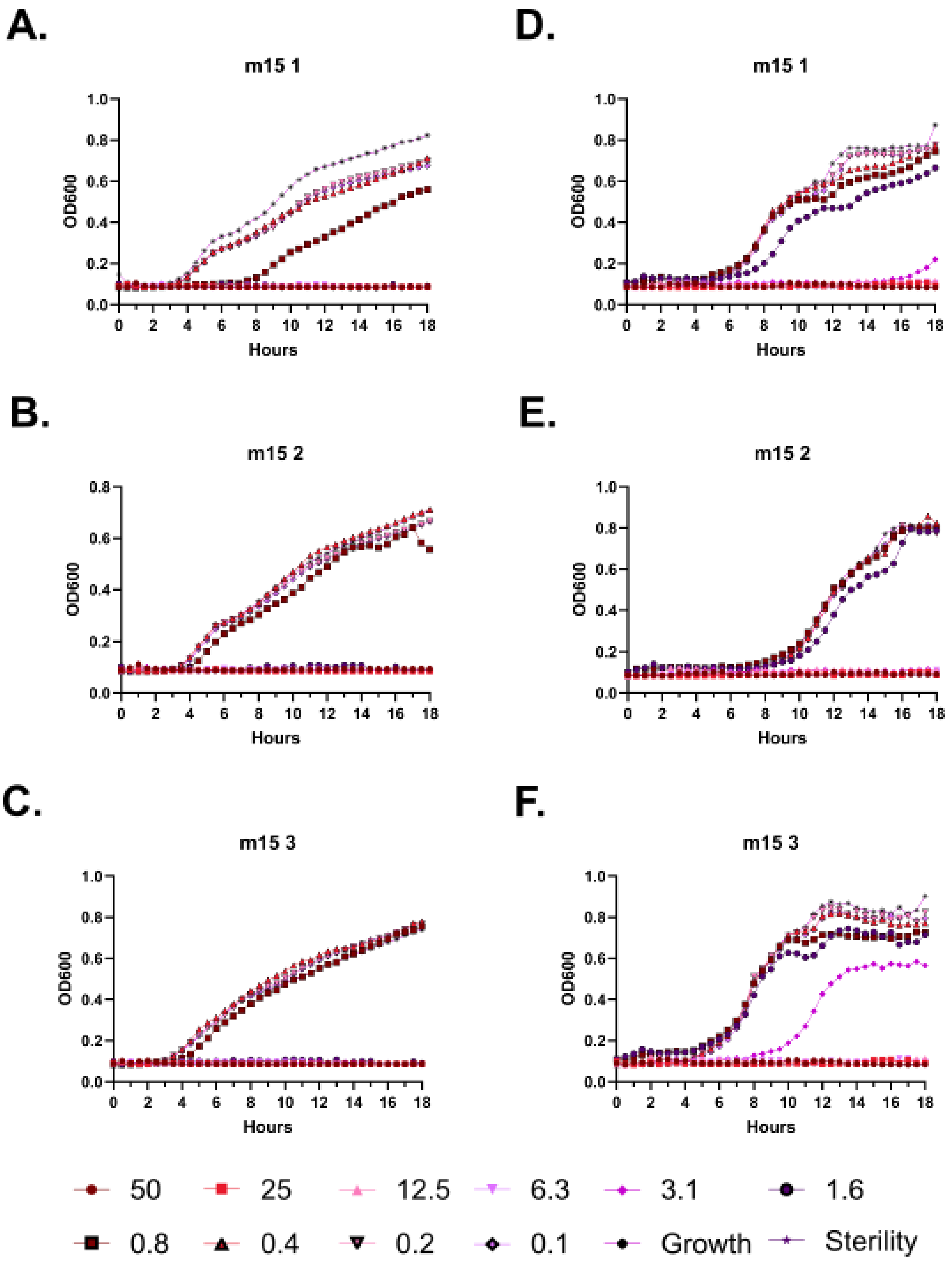
A.
m15 1
B.
m15 2
C.
m15 3
D.
m15 1
E.
m15 2
F.
m15 3
OD600
Hours
50
25
12.5
6.3
3.1
1.6
0.8
0.4
0.2
0.1
Growth
Sterility

**Supplementary Figure 41 – Kinetic antimicrobial susceptibility testing for (m15). A-C.** *E. coli*. **D-F.** *Pseudomonas.* Bacterial growth was monitored by OD600 over 18 hours at 37 °C in the presence of the indicated peptide concentrations. Increasing OD600 indicates increasing bacterial growth. Peptide concentrations are given in μM. Each curve indicates an independent experiment.

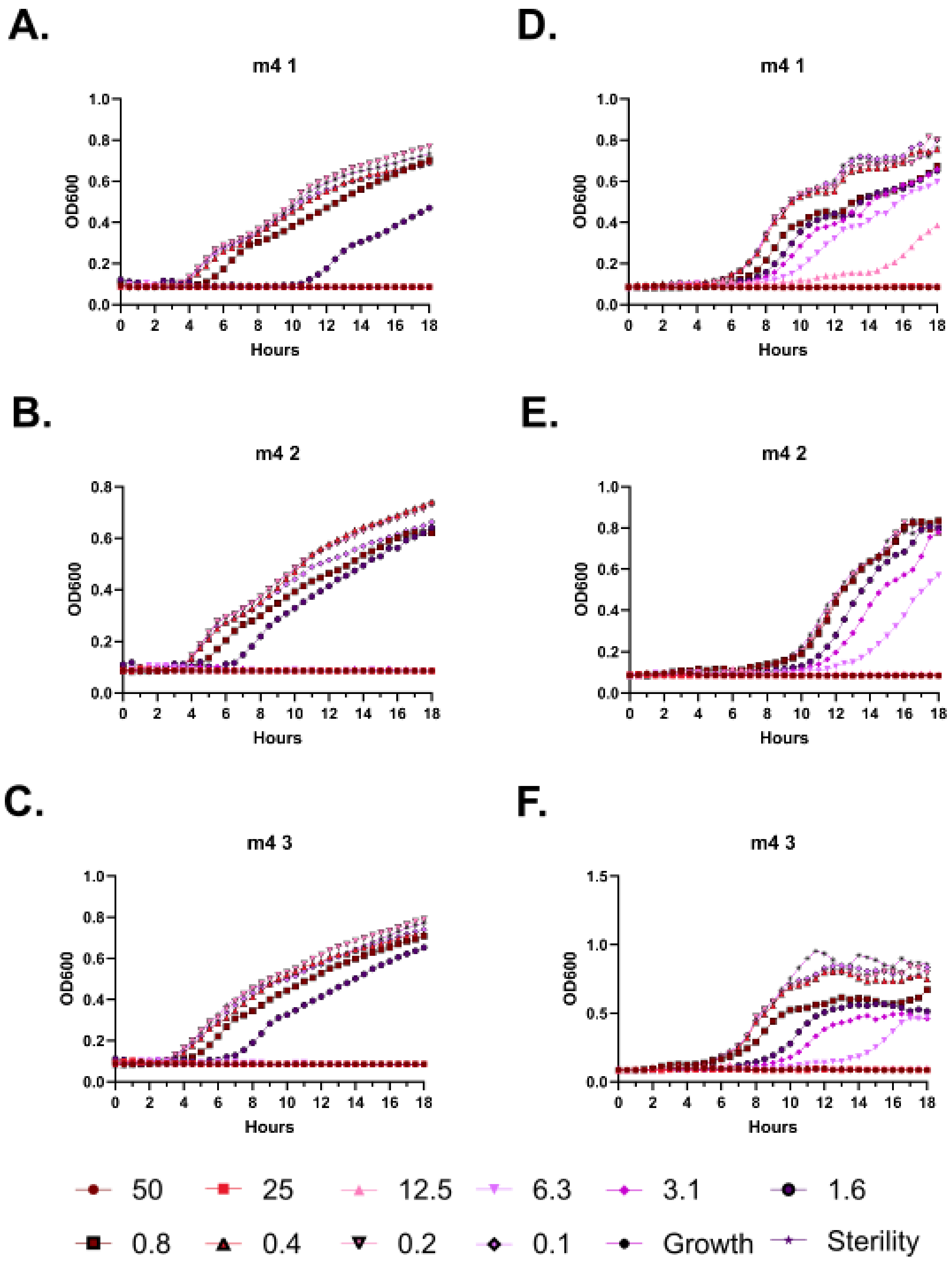

A.
m4 1
OD600
Hours
D.
m4 1
OD600
Hours
B.
m4 2
OD600
Hours
E.
m4 2
OD600
Hours
C.
m4 3
OD600
Hours
F.
m4 3
OD600
Hours
50
25
12.5
6.3
3.1
1.6
0.8
0.4
0.2
0.1
Growth
Sterility

**Supplementary Figure 42 – Kinetic antimicrobial susceptibility testing for (m4). A-C.** *E. coli*. **D-F.** *Pseudomonas.* Bacterial growth was monitored by OD600 over 18 hours at 37 °C in the presence of the indicated peptide concentrations. Increasing OD600 indicates increasing bacterial growth. Peptide concentrations are given in μM. Each curve indicates an independent experiment.

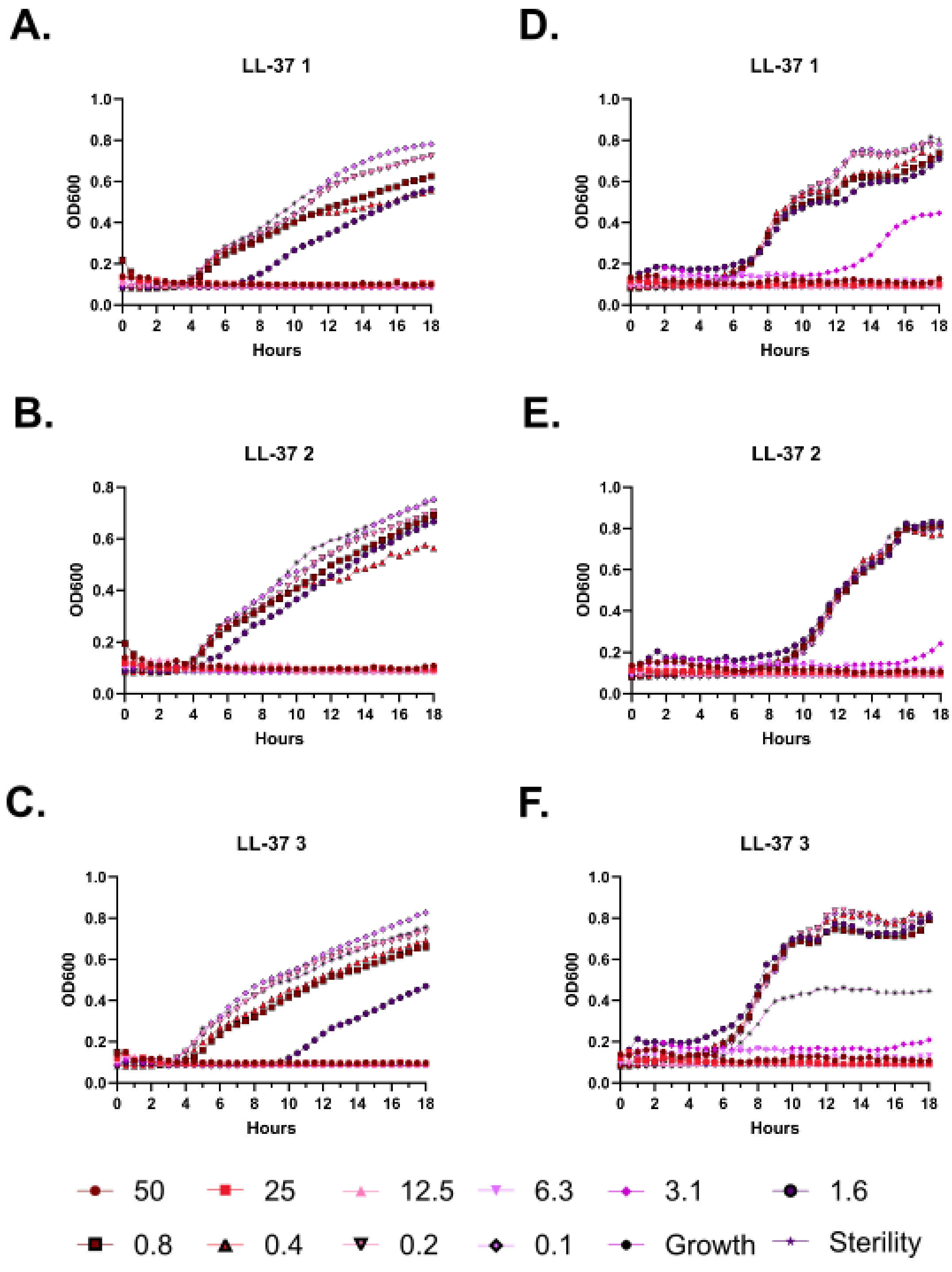
A.
LL-37 1
OD600
Hours
B.
LL-37 2
OD600
Hours
C.
LL-37 3
OD600
Hours
D.
LL-37 1
OD600
Hours
E.
LL-37 2
OD600
Hours
F.
LL-37 3
OD600
Hours
50
25
12.5
6.3
3.1
1.6
0.8
0.4
0.2
0.1
Growth
Sterility

**Supplementary Figure 43 – Kinetic antimicrobial susceptibility testing for (LL-37). A-C.** *E. coli*. **D-F.** *Pseudomonas.* Bacterial growth was monitored by OD600 over 18 hours at 37 °C in the presence of the indicated peptide concentrations. Increasing OD600 indicates increasing bacterial growth. Peptide concentrations are given in µM. Each curve indicates an independent experiment.

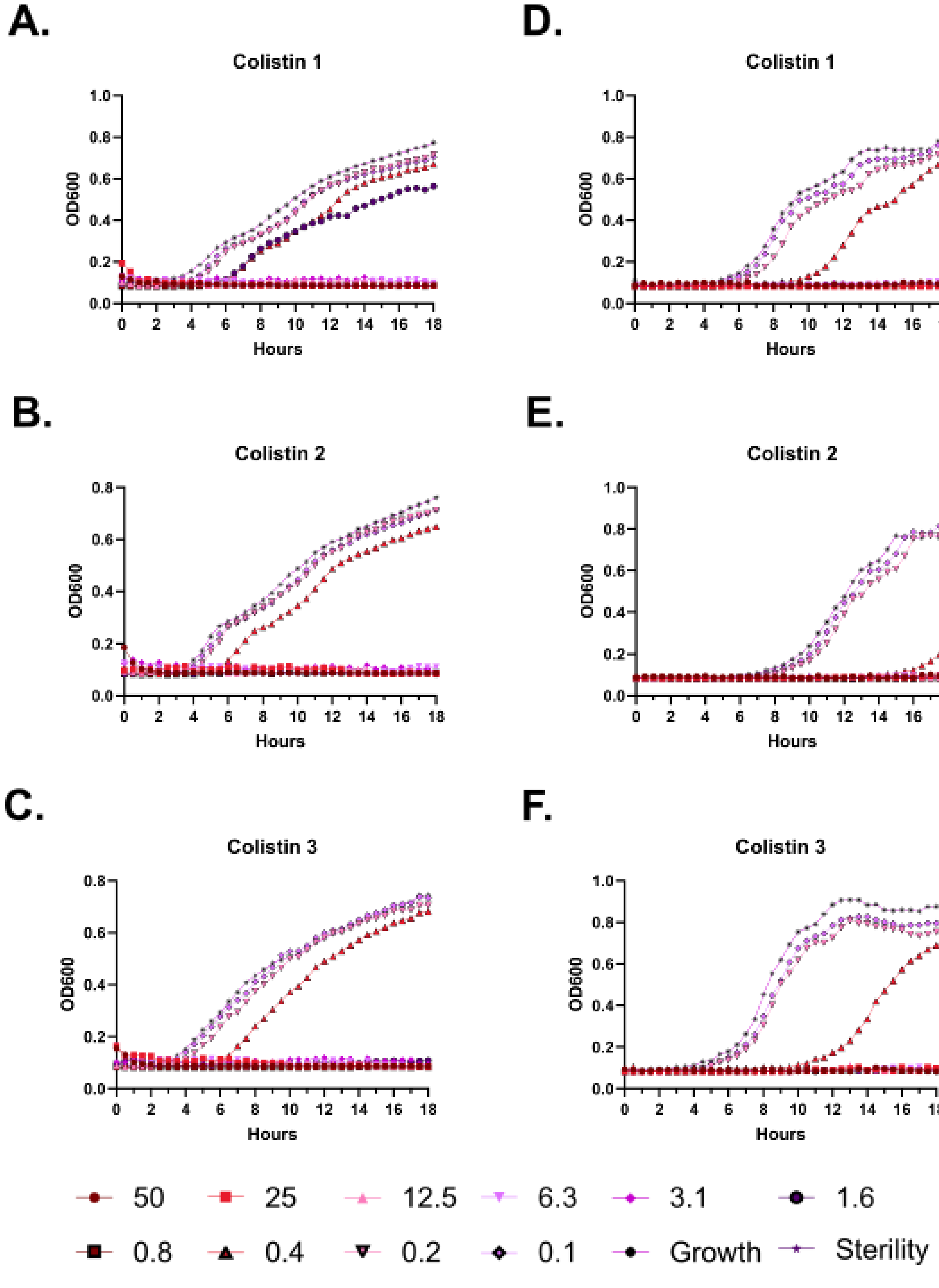
A.
Colistin 1
D.
Colistin 1
B.
Colistin 2
E.
Colistin 2
C.
Colistin 3
F.
Colistin 3
OD600
Hours
50
25
12.5
6.3
3.1
1.6
0.8
0.4
0.2
0.1
Growth
Sterility

**Supplementary Figure 44 – Kinetic antimicrobial susceptibility testing for (Colistin). A-C.** *E. coli*. **D-F.** *Pseudomonas.* Bacterial growth was monitored by OD600 over 18 hours at 37 °C in the presence of the indicated peptide concentrations. Increasing OD600 indicates increasing bacterial growth. Peptide concentrations are given in µM. Each curve indicates an independent experiment.

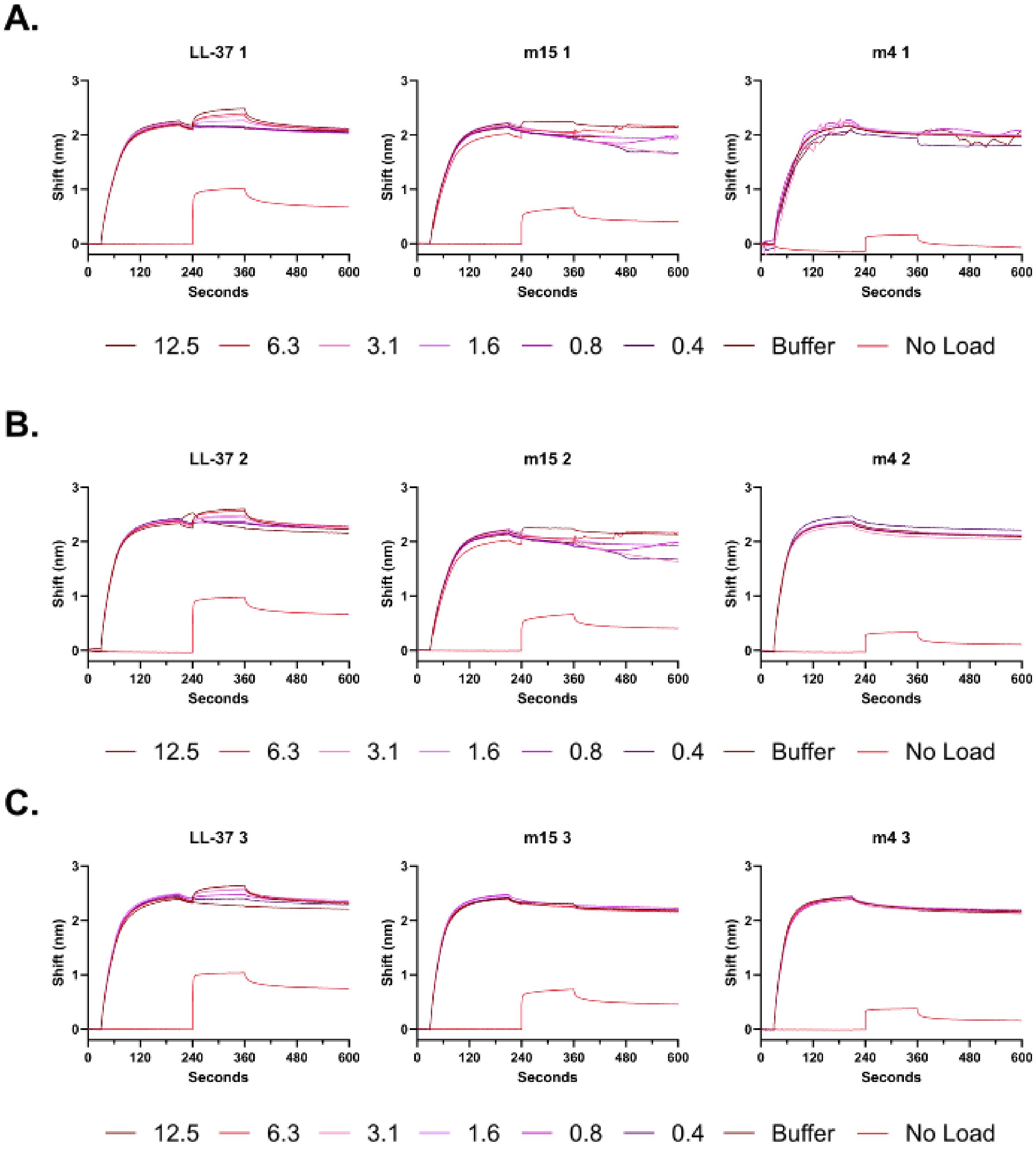
A.
LL-37 1
m15 1
m4 1
B.
LL-37 2
m15 2
m4 2
C.
LL-37 3
m15 3
m4 3
Shift (nm)
Seconds
0 120 240 360 480 600
12.5
6.3
3.1
1.6
0.8
0.4
Buffer
No Load

**Supplementary Figure 45 – Raw biolayer interferometry (BLI) traces for interactions between LL-37 probe and LL-37, m15, or m4 analytes. A.** BLI traces for Experiment 1. Buffer indicates a tip loaded with biotinylated LL-37 and then treated with buffer analyte (PBS). No load indicates a probe without any biotinylated LL-37 that is subsequently incubated with the corresponding peptide analyte at 12.5 µM. **B.** BLI traces for Experiment 2. **C.** BLI traces for Experiment 3. Peptide concentrations are given in µM.

| Sheep Defib | < 20% Fold Δ | |
|---|---|---|
| LL-37 | 0.4 | |
| m15 | 6.3 | 15.8 |
| m4 | 100 | 250 |
| Colistin | 50 | 125 |

| Cow Defib | < 20% Fold Δ | |
|---|---|---|
| LL-37 | 0.8 | |
| m15 | 200 | 250 |
| m4 | 200 | 250 |
| Colistin | 100 | 125 |

| Human 1 | < 40% Fold Δ | |
|---|---|---|
| LL-37 | 0.8 | |
| m15 | 1.6 | 2 |
| m4 | 25 | 31.3 |
| Colistin | 200 | 250 |

| Sheep RBCs | < 20% Fold Δ | |
|---|---|---|
| LL-37 | 0.8 | |
| m15 | 25 | 31.3 |
| m4 | 200 | 250 |
| Colistin | 200 | 250 |

| Human 2 | < 20% Fold Δ | |
|---|---|---|
| LL-37 | 0.8 | |
| m15 | 3.1 | 3.9 |
| m4 | 12.5 | 15.6 |
| Colistin | 200 | 250 |

| Human RBCs | < 40% Fold Δ | |
|---|---|---|
| LL-37 | 0.8 | |
| m15 | 1.6 | 2 |
| m4 | 12.5 | 15.6 |
| Colistin | 200 | 250 |

| | Avg. | SD |
|---|---|---|
| m15 | 51 | 98 |
| m4 | 135 | 126 |
| Colistin | 208 | 65 |

**Supplementary Figure 46 – Fold-improvement in hemolytic activity relative to LL-37 across species.** For each sample type, a number indicating the greatest peptide concentration in μM at which hemolysis levels are < 20% or < 40% is given. The choice of threshold is generally based on the overall propensity of a given sample for hemolysis in the presence of colistin, which despite itself being a relatively toxic compound *in vivo* serves as a reference point. A second column divides this concentration by the value obtained for LL-37 on that sample type to estimate a fold-improvement in hemolytic activity relative to LL-37.

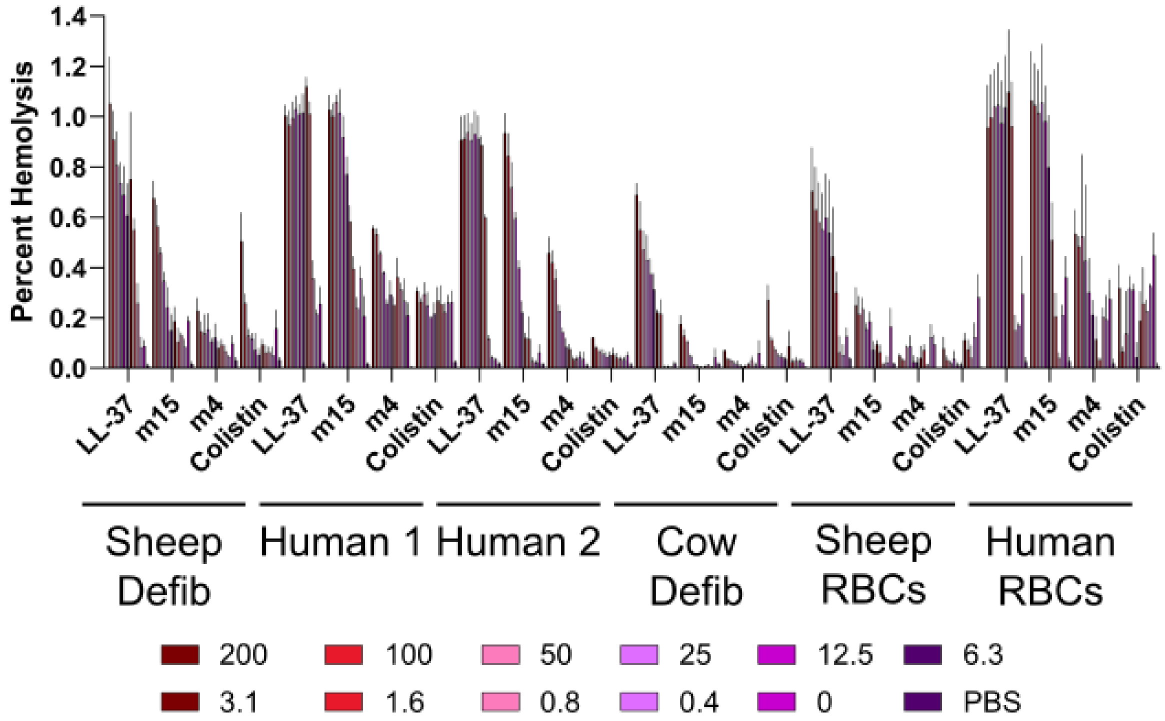

Percent Hemolysis
1.4
1.2
1.0
0.8
0.6
0.4
0.2
0.0
LL-37
m15
m4
Colistin
Sheep Defib
Human 1
Human 2
Cow Defib
Sheep RBCs
Human RBCs
200
100
50
25
12.5
6.3
3.1
1.6
0.8
0.4
0
PBS

**Supplementary Figure 47 – Histogram depiction of Figure 3D data. A.** Endpoint hemolytic activity of each peptide relative to a 100% hemolysis freeze-thaw control after 18 hours at 37 °C. Free hemoglobin levels in cell supernatants were assessed by measuring absorbance at 414 nm.

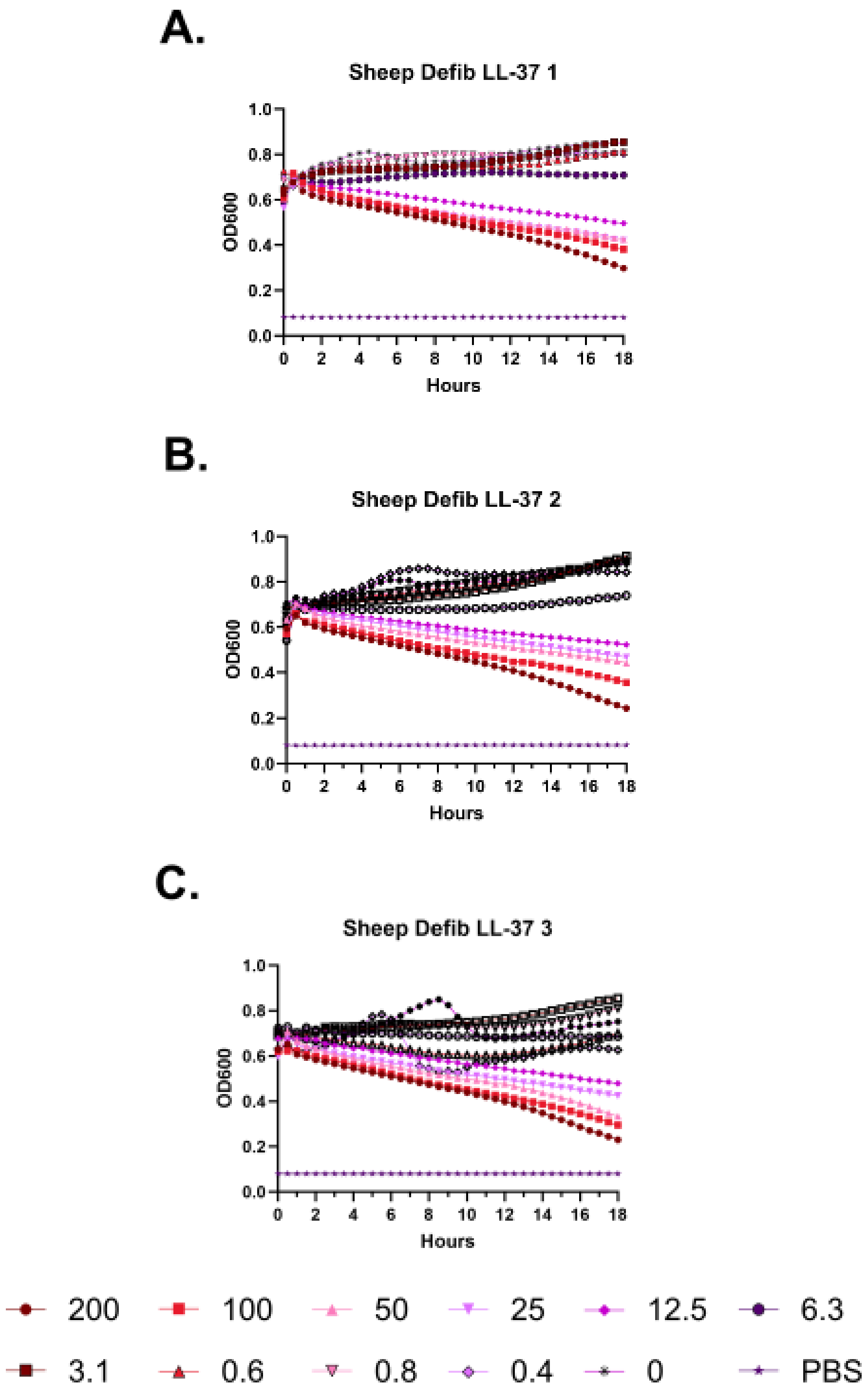
A.
Sheep Defib LL-37 1
OD600
Hours
B.
Sheep Defib LL-37 2
OD600
Hours
C.
Sheep Defib LL-37 3
OD600
Hours
200
100
50
25
12.5
6.3
3.1
0.6
0.8
0.4
0
PBS

**Supplementary Figure 48 – Kinetic hemolysis assays for LL-37 in defibrinated sheep blood. A-C.** Hemolysis was monitored by OD600 over 18 hours at 37 °C in the presence of the indicated peptide concentrations. Decreasing OD600 indicates increasing hemolysis. Peptide concentrations are given in μM. Each curve indicates an independent experiment.

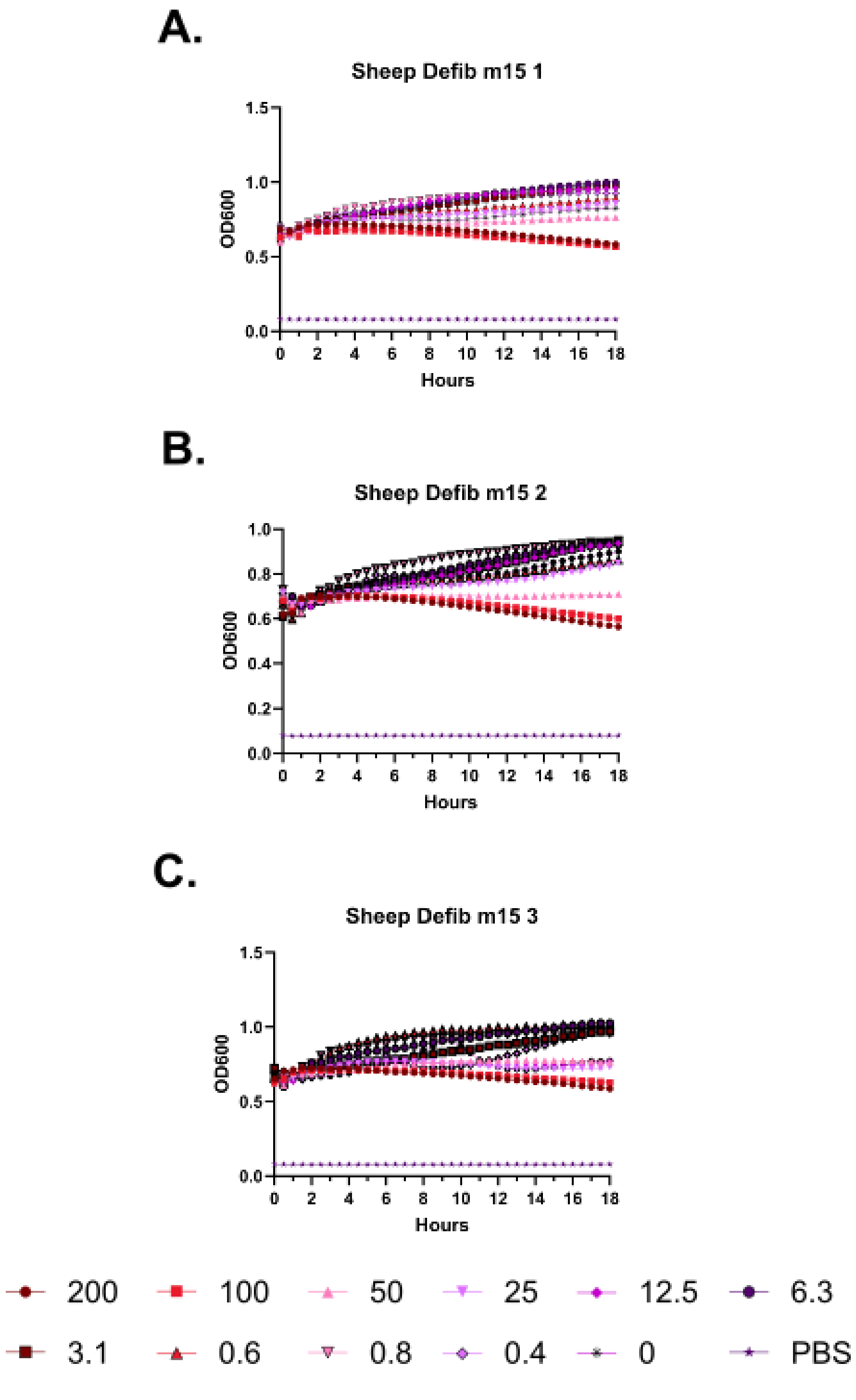
A.
Sheep Defib m15 1
OD600
Hours
B.
Sheep Defib m15 2
OD600
Hours
C.
Sheep Defib m15 3
OD600
Hours
200
100
50
25
12.5
6.3
3.1
0.6
0.8
0.4
0
PBS

**Supplementary Figure 49 – Kinetic hemolysis assays for m15 in defibrinated sheep blood. A-C.** Hemolysis was monitored by OD600 over 18 hours at 37 °C in the presence of the indicated peptide concentrations. Decreasing OD600 indicates increasing hemolysis. Peptide concentrations are given in μM. Each curve indicates an independent experiment.

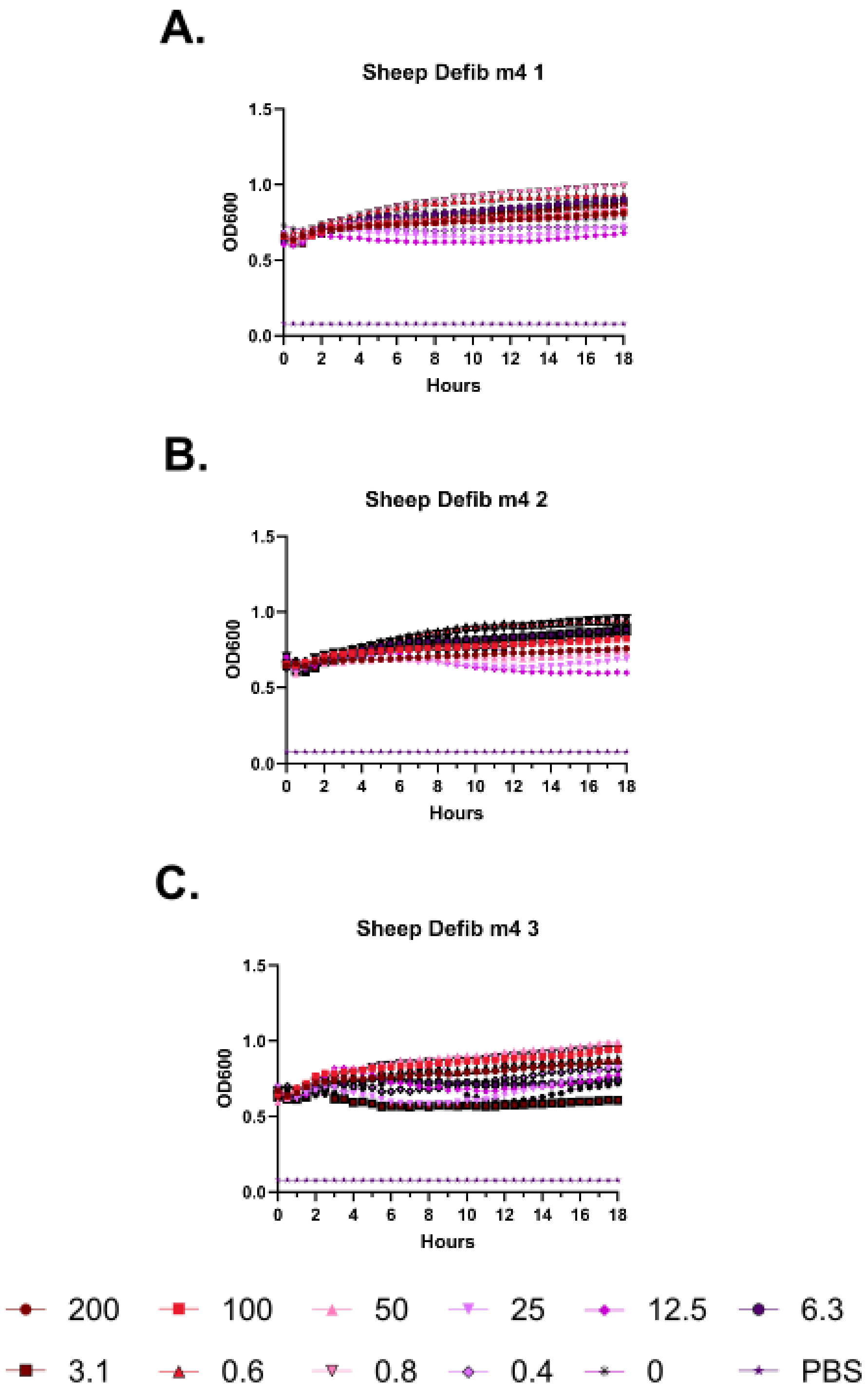
A.
Sheep Defib m4 1
OD600
Hours
B.
Sheep Defib m4 2
OD600
Hours
C.
Sheep Defib m4 3
OD600
Hours
200
100
50
25
12.5
6.3
3.1
0.6
0.8
0.4
0
PBS

**Supplementary Figure 50 – Kinetic hemolysis assays for m4 in defibrinated sheep blood. A-C.** Hemolysis was monitored by OD600 over 18 hours at 37 °C in the presence of the indicated peptide concentrations. Decreasing OD600 indicates increasing hemolysis. Peptide concentrations are given in μM. Each curve indicates an independent experiment.

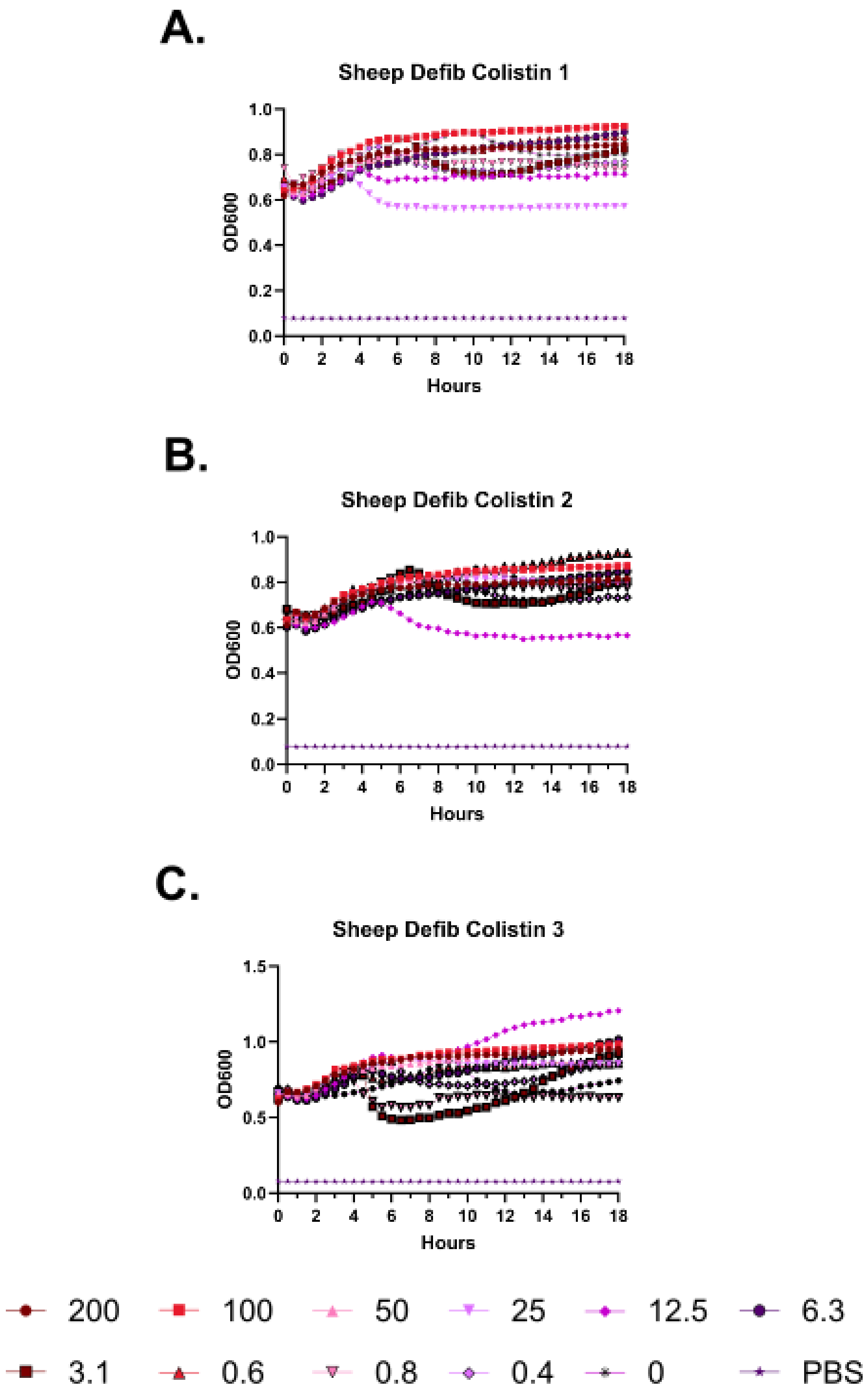
A.
Sheep Defib Colistin 1
OD600
Hours
B.
Sheep Defib Colistin 2
OD600
Hours
C.
Sheep Defib Colistin 3
OD600
Hours
200
100
50
25
12.5
6.3
3.1
0.6
0.8
0.4
0
PBS

**Supplementary Figure 51 – Kinetic hemolysis assays for colistin in defibrinated sheep blood. A-C.** Hemolysis was monitored by OD600 over 18 hours at 37 °C in the presence of the indicated peptide concentrations. Decreasing OD600 indicates increasing hemolysis. Peptide concentrations are given in μM. Each curve indicates an independent experiment.

A.

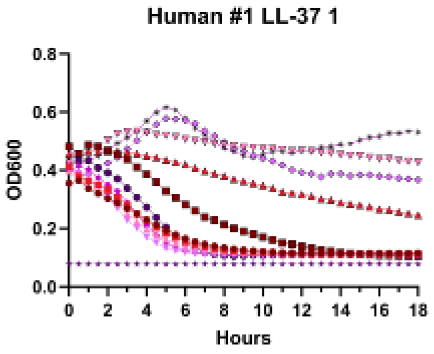
Human #1 LL-37 1
OD600
Hours
0.0
0.2
0.4
0.6
0.8
0
2
4
6
8
10
12
14
16
18

B.

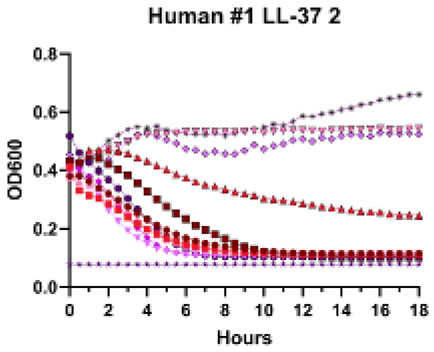
Human #1 LL-37 2
OD600
Hours
0.0
0.2
0.4
0.6
0.8
0
2
4
6
8
10
12
14
16
18

C.

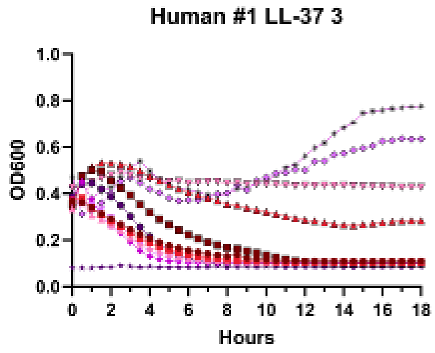
Human #1 LL-37 3
OD600
Hours
0.0
0.2
0.4
0.6
0.8
1.0
0
2
4
6
8
10
12
14
16
18

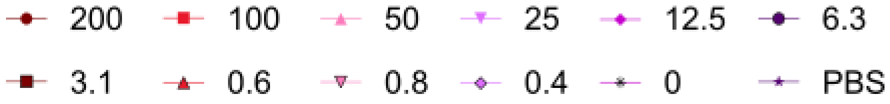
200
100
50
25
12.5
6.3
3.1
0.6
0.8
0.4
0
PBS

**Supplementary Figure 52 – Kinetic hemolysis assays for LL-37 in human whole blood donor #1. A-C.** Hemolysis was monitored by OD600 over 18 hours at 37 °C in the presence of the indicated peptide concentrations. Decreasing OD600 indicates increasing hemolysis. Peptide concentrations are given in μM. Each curve indicates an independent experiment.

**A.**

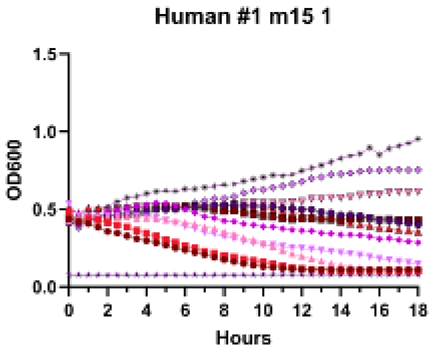

Human #1 m15 1
OD600
Hours


**B.**

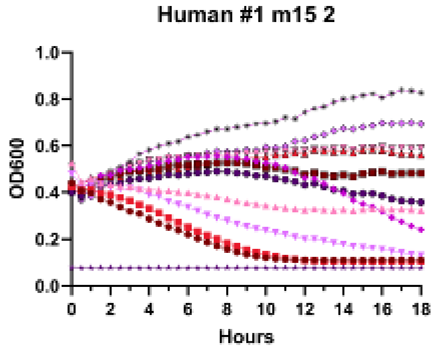

Human #1 m15 2
OD600
Hours


**C.**

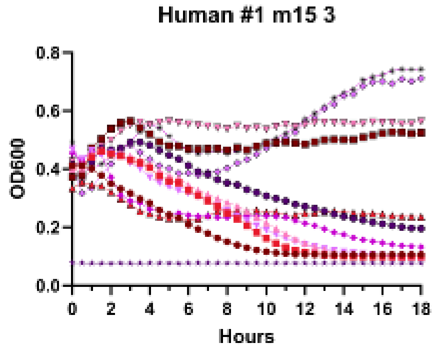

Human #1 m15 3
OD600
Hours


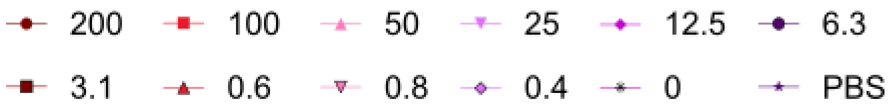

200
100
50
25
12.5
6.3
3.1
0.6
0.8
0.4
0
PBS

**Supplementary Figure 53 – Kinetic hemolysis assays for m15 in human whole blood donor #1. A-C.** Hemolysis was monitored by OD600 over 18 hours at 37 °C in the presence of the indicated peptide concentrations. Decreasing OD600 indicates increasing hemolysis. Peptide concentrations are given in μM. Each curve indicates an independent experiment.

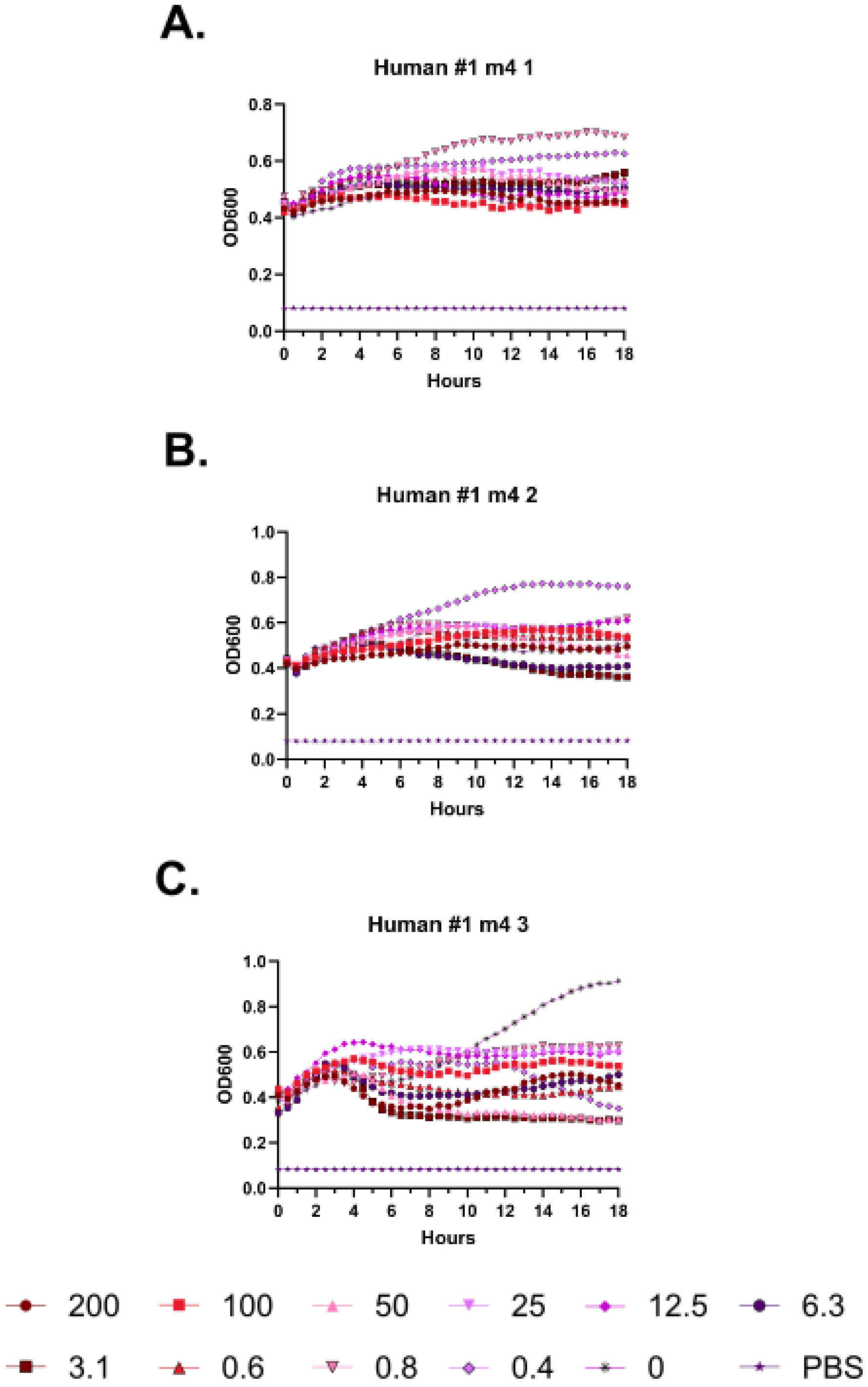
A.
Human #1 m4 1
OD600
Hours
B.
Human #1 m4 2
OD600
Hours
C.
Human #1 m4 3
OD600
Hours
200
100
50
25
12.5
6.3
3.1
0.6
0.8
0.4
0
PBS

**Supplementary Figure 54 – Kinetic hemolysis assays for m4 in human whole blood donor #1. A-C.** Hemolysis was monitored by OD600 over 18 hours at 37 °C in the presence of the indicated peptide concentrations. Decreasing OD600 indicates increasing hemolysis. Peptide concentrations are given in μM. Each curve indicates an independent experiment.

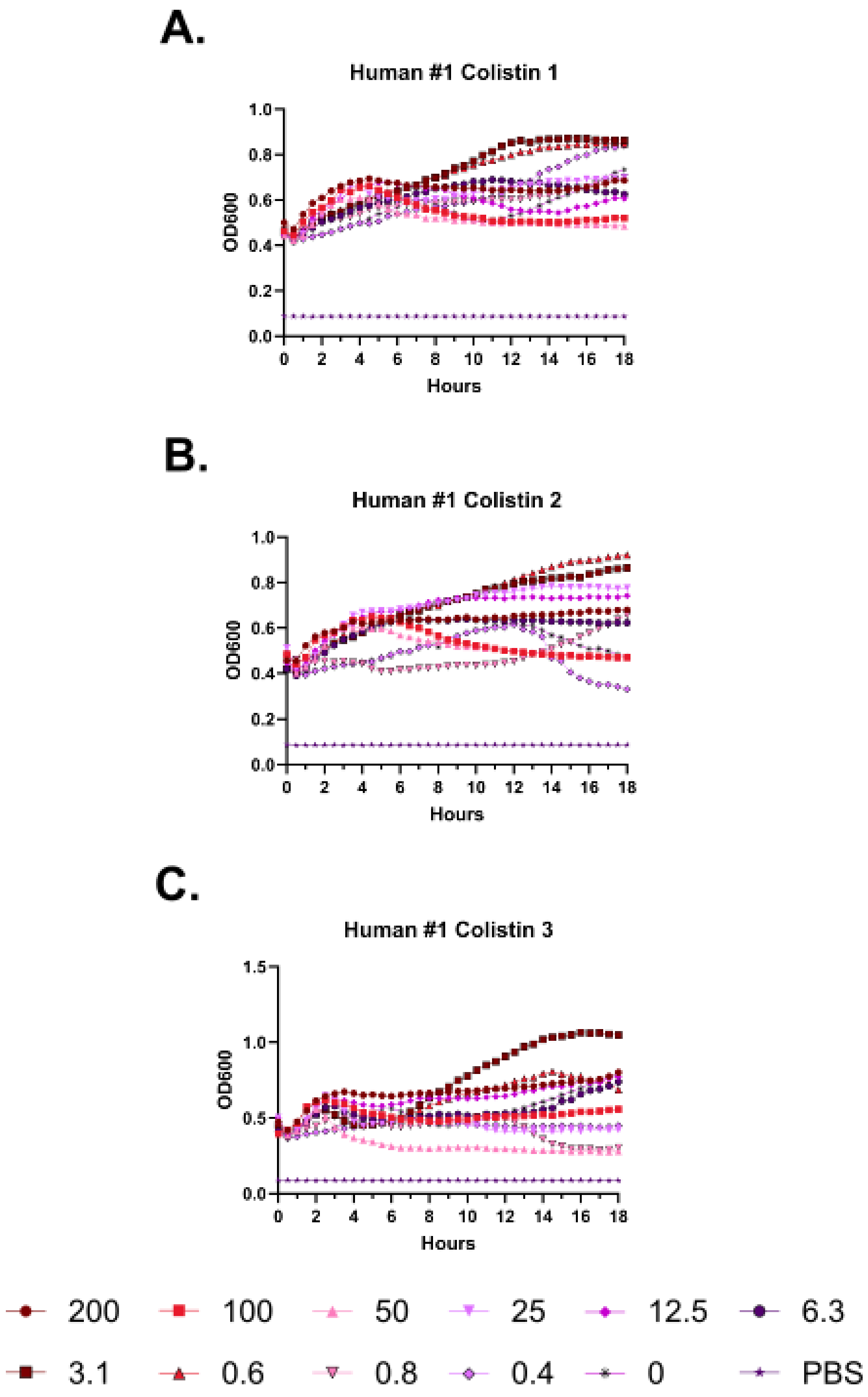
A.
Human #1 Colistin 1
OD600
Hours
B.
Human #1 Colistin 2
OD600
Hours
C.
Human #1 Colistin 3
OD600
Hours
200
100
50
25
12.5
6.3
3.1
0.6
0.8
0.4
0
PBS

**Supplementary Figure 55 – Kinetic hemolysis assays for colistin in human whole blood donor #1. A-C.** Hemolysis was monitored by OD600 over 18 hours at 37 °C in the presence of the indicated peptide concentrations. Decreasing OD600 indicates increasing hemolysis. Peptide concentrations are given in μM. Each curve indicates an independent experiment.

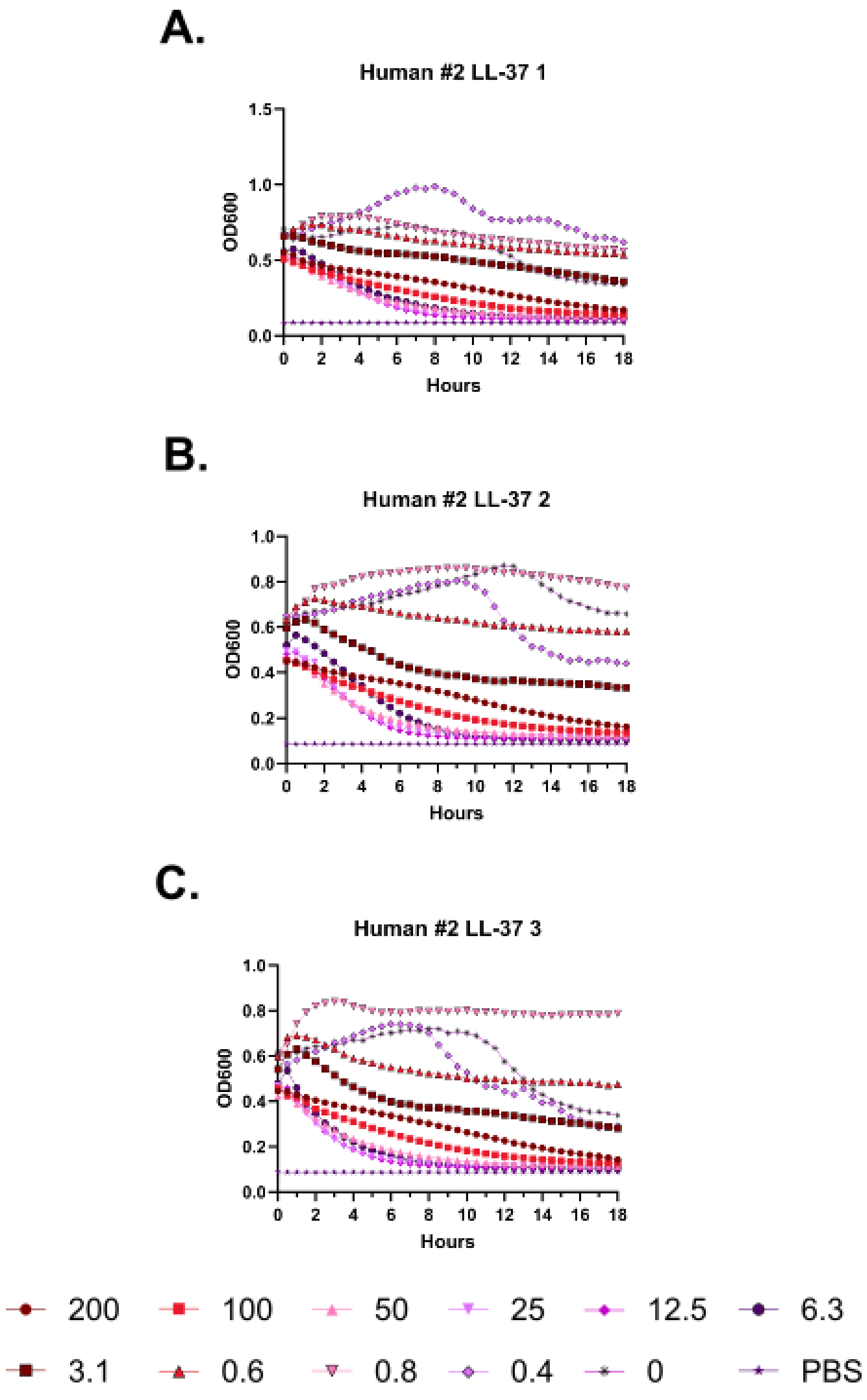
A.
Human #2 LL-37 1
OD600
Hours
B.
Human #2 LL-37 2
OD600
Hours
C.
Human #2 LL-37 3
OD600
Hours
200
100
50
25
12.5
6.3
3.1
0.6
0.8
0.4
0
PBS

**Supplementary Figure 56 – Kinetic hemolysis assays for LL-37 in human whole blood donor #2. A-C.** Hemolysis was monitored by OD600 over 18 hours at 37 °C in the presence of the indicated peptide concentrations. Decreasing OD600 indicates increasing hemolysis. Peptide concentrations are given in μM. Each curve indicates an independent experiment.

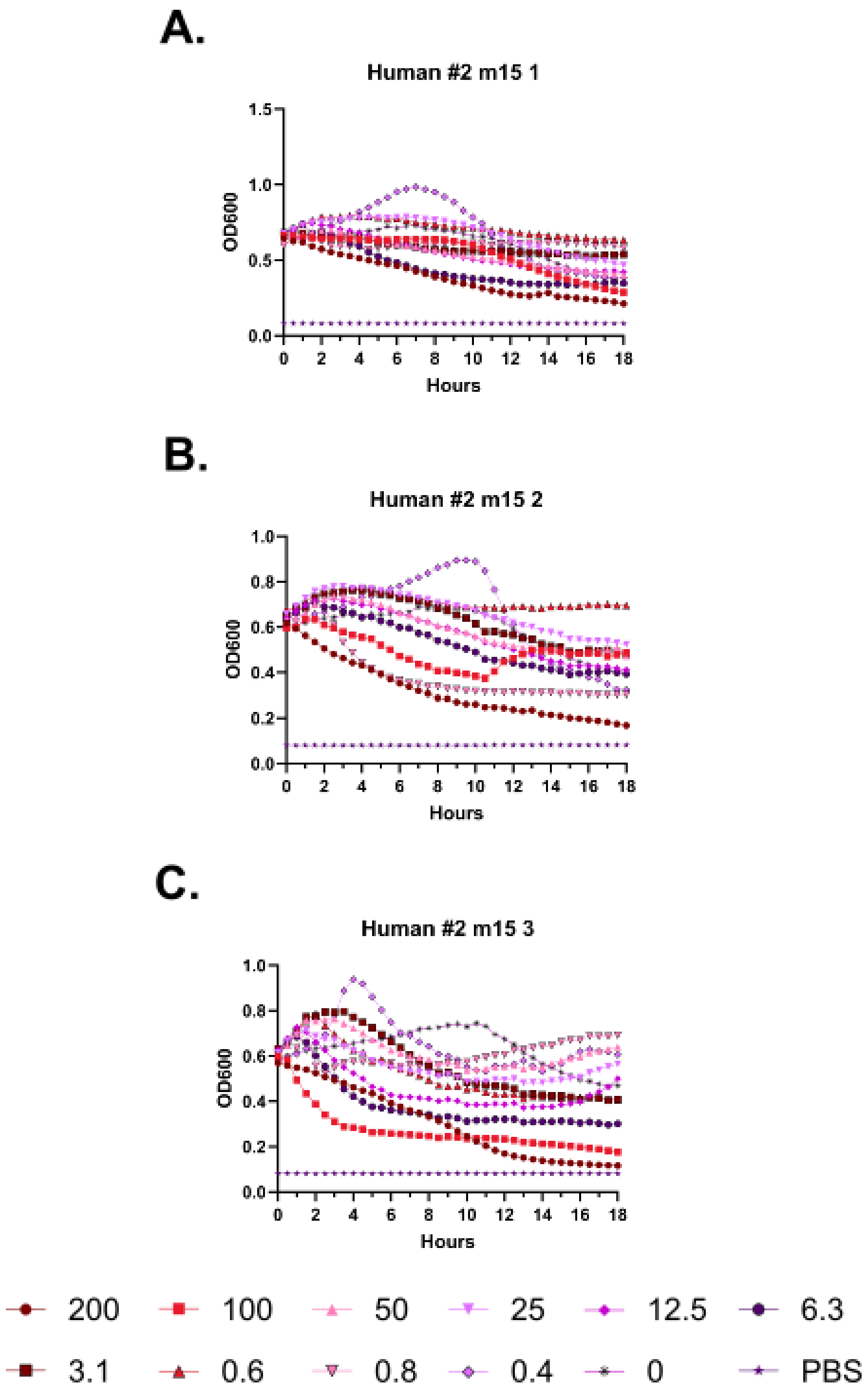
A.
Human #2 m15 1
OD600
Hours
B.
Human #2 m15 2
OD600
Hours
C.
Human #2 m15 3
OD600
Hours
200
100
50
25
12.5
6.3
3.1
0.6
0.8
0.4
0
PBS

**Supplementary Figure 57 – Kinetic hemolysis assays for m15 in human whole blood donor #2. A-C.** Hemolysis was monitored by OD600 over 18 hours at 37 °C in the presence of the indicated peptide concentrations. Decreasing OD600 indicates increasing hemolysis. Peptide concentrations are given in μM. Each curve indicates an independent experiment.

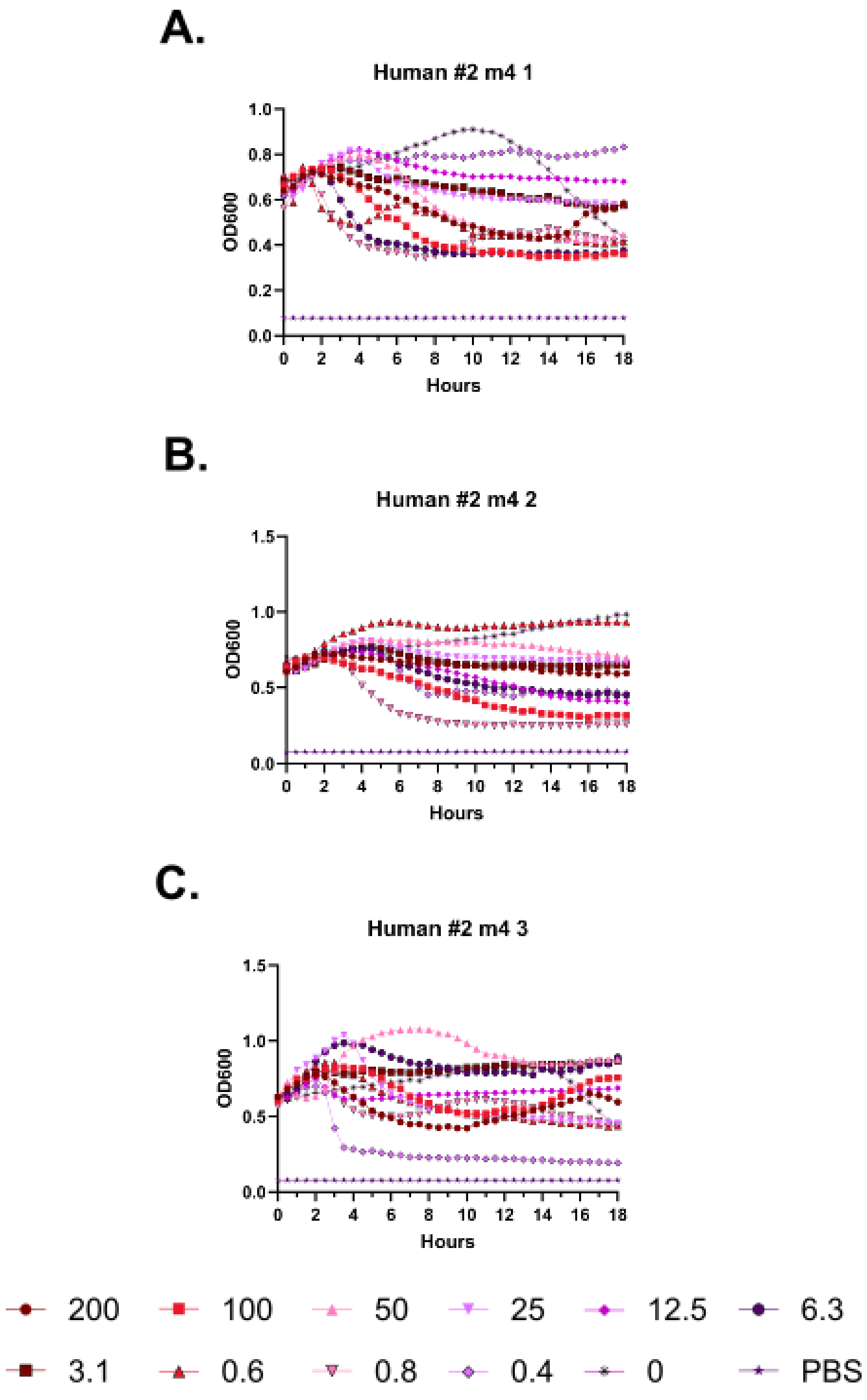
A.
Human #2 m4 1
OD600
Hours
B.
Human #2 m4 2
OD600
Hours
C.
Human #2 m4 3
OD600
Hours
200
100
50
25
12.5
6.3
3.1
0.6
0.8
0.4
0
PBS

**Supplementary Figure 58 – Kinetic hemolysis assays for m4 in human whole blood donor #2. A-C.** Hemolysis was monitored by OD600 over 18 hours at 37 °C in the presence of the indicated peptide concentrations. Decreasing OD600 indicates increasing hemolysis. Peptide concentrations are given in μM. Each curve indicates an independent experiment.

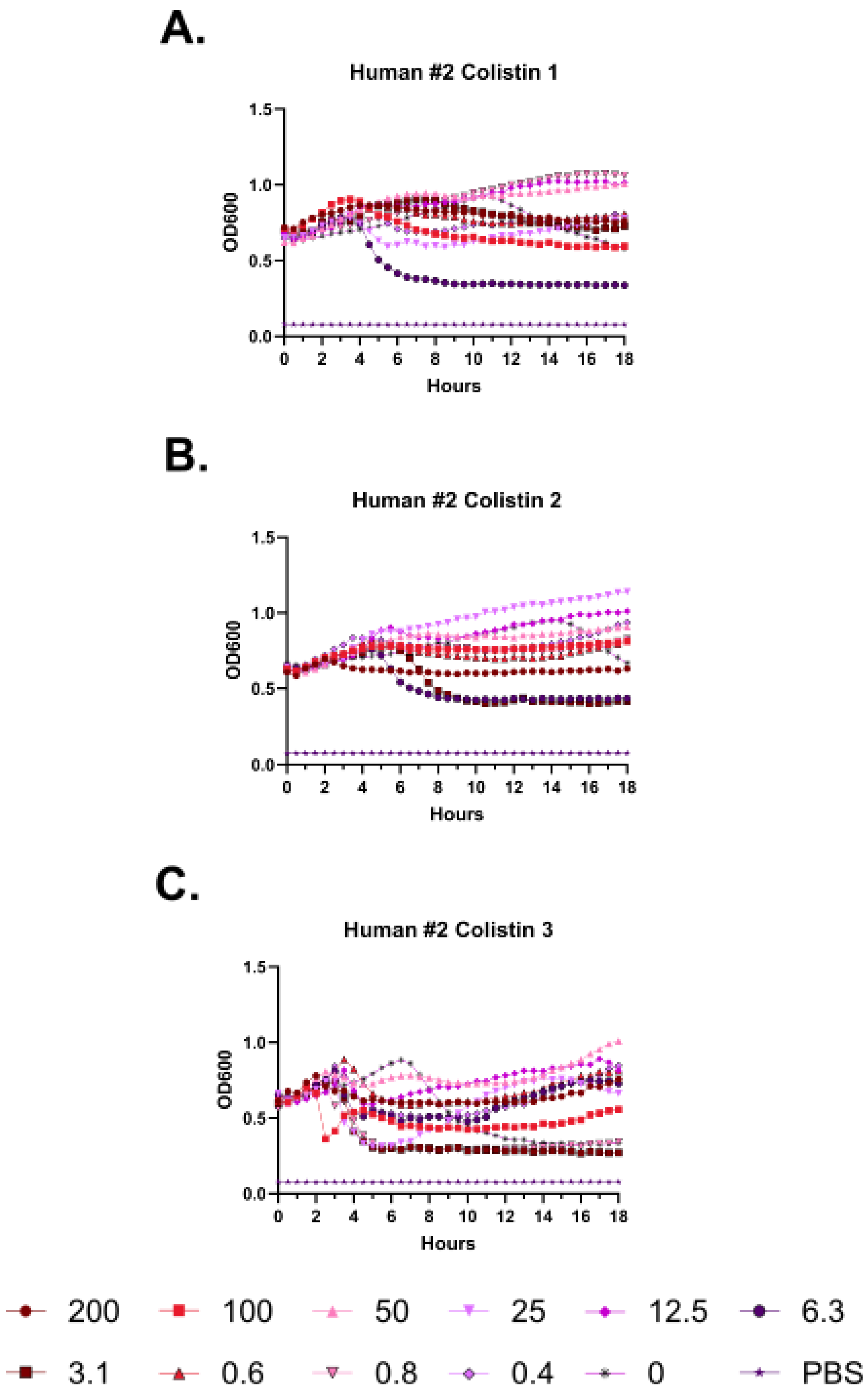
A.
Human #2 Colistin 1
OD600
Hours
B.
Human #2 Colistin 2
OD600
Hours
C.
Human #2 Colistin 3
OD600
Hours
200
100
50
25
12.5
6.3
3.1
0.6
0.8
0.4
0
PBS

**Supplementary Figure 59 – Kinetic hemolysis assays for colistin in human whole blood donor #2. A-C.** Hemolysis was monitored by OD600 over 18 hours at 37 °C in the presence of the indicated peptide concentrations. Decreasing OD600 indicates increasing hemolysis. Peptide concentrations are given in μM. Each curve indicates an independent experiment.

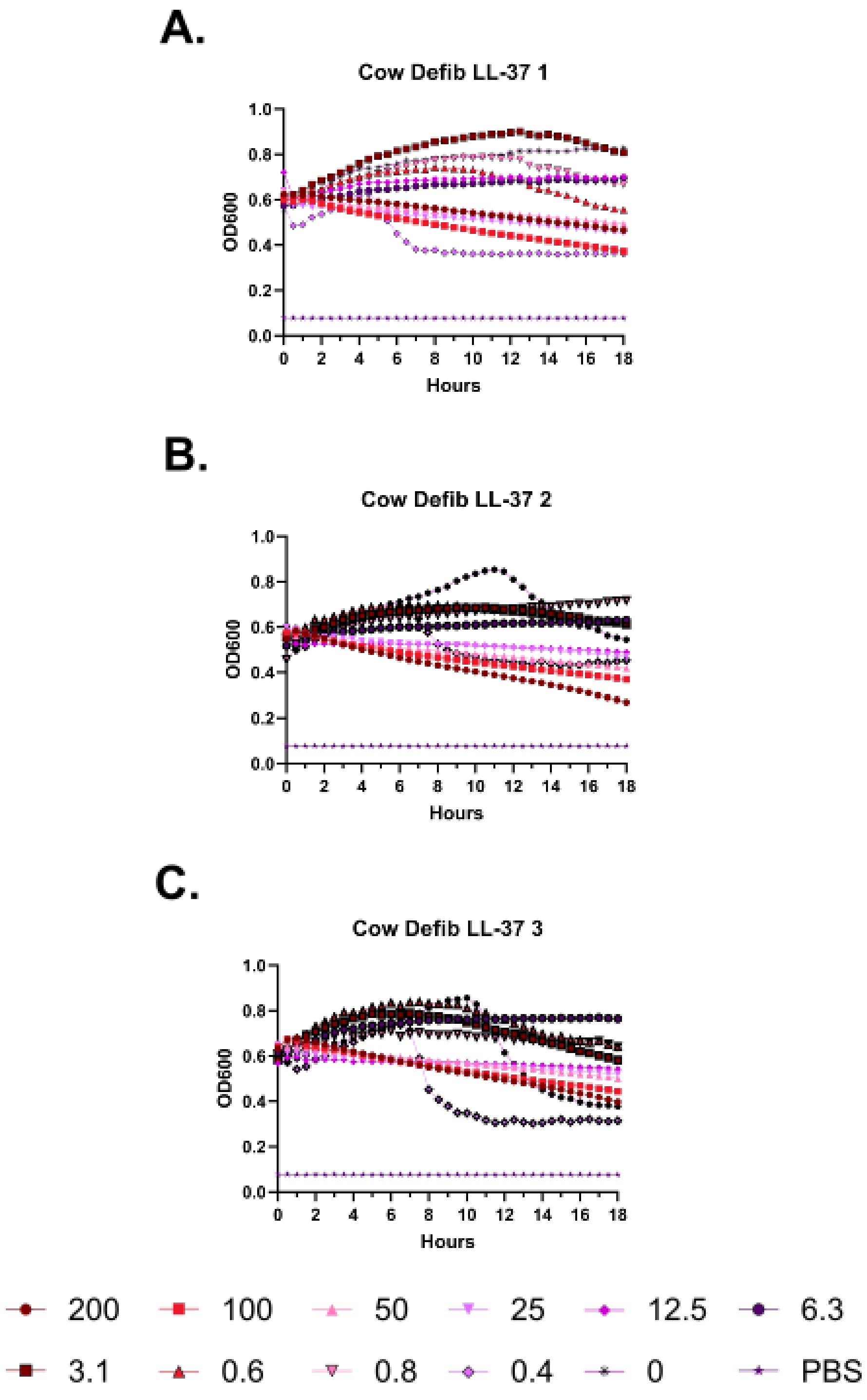
A.
Cow Defib LL-37 1
OD600
Hours
B.
Cow Defib LL-37 2
OD600
Hours
C.
Cow Defib LL-37 3
OD600
Hours
200
100
50
25
12.5
6.3
3.1
0.6
0.8
0.4
0
PBS

**Supplementary Figure 60 – Kinetic hemolysis assays for LL-37 in defibrinated cow blood. A-C.** Hemolysis was monitored by OD600 over 18 hours at 37 °C in the presence of the indicated peptide concentrations. Decreasing OD600 indicates increasing hemolysis. Peptide concentrations are given in μM. Each curve indicates an independent experiment.

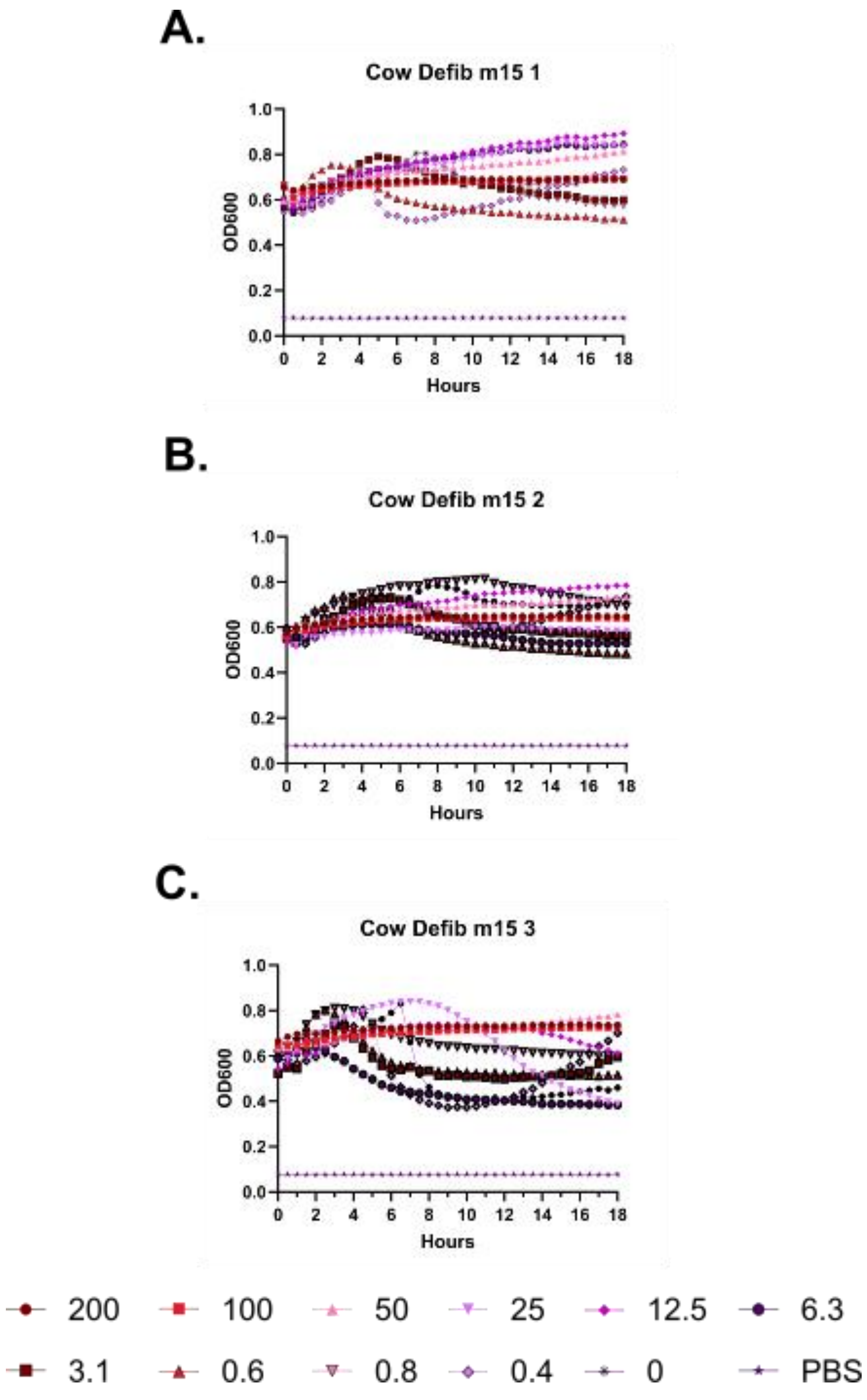
A.
Cow Defib m15 1
OD600
Hours
B.
Cow Defib m15 2
OD600
Hours
C.
Cow Defib m15 3
OD600
Hours
200
100
50
25
12.5
6.3
3.1
0.6
0.8
0.4
0
PBS

**Supplementary Figure 61 – Kinetic hemolysis assays for m15 in defibrinated cow blood. A-C.** Hemolysis was monitored by OD600 over 18 hours at 37 °C in the presence of the indicated peptide concentrations. Decreasing OD600 indicates increasing hemolysis. Peptide concentrations are given in μM. Each curve indicates an independent experiment.

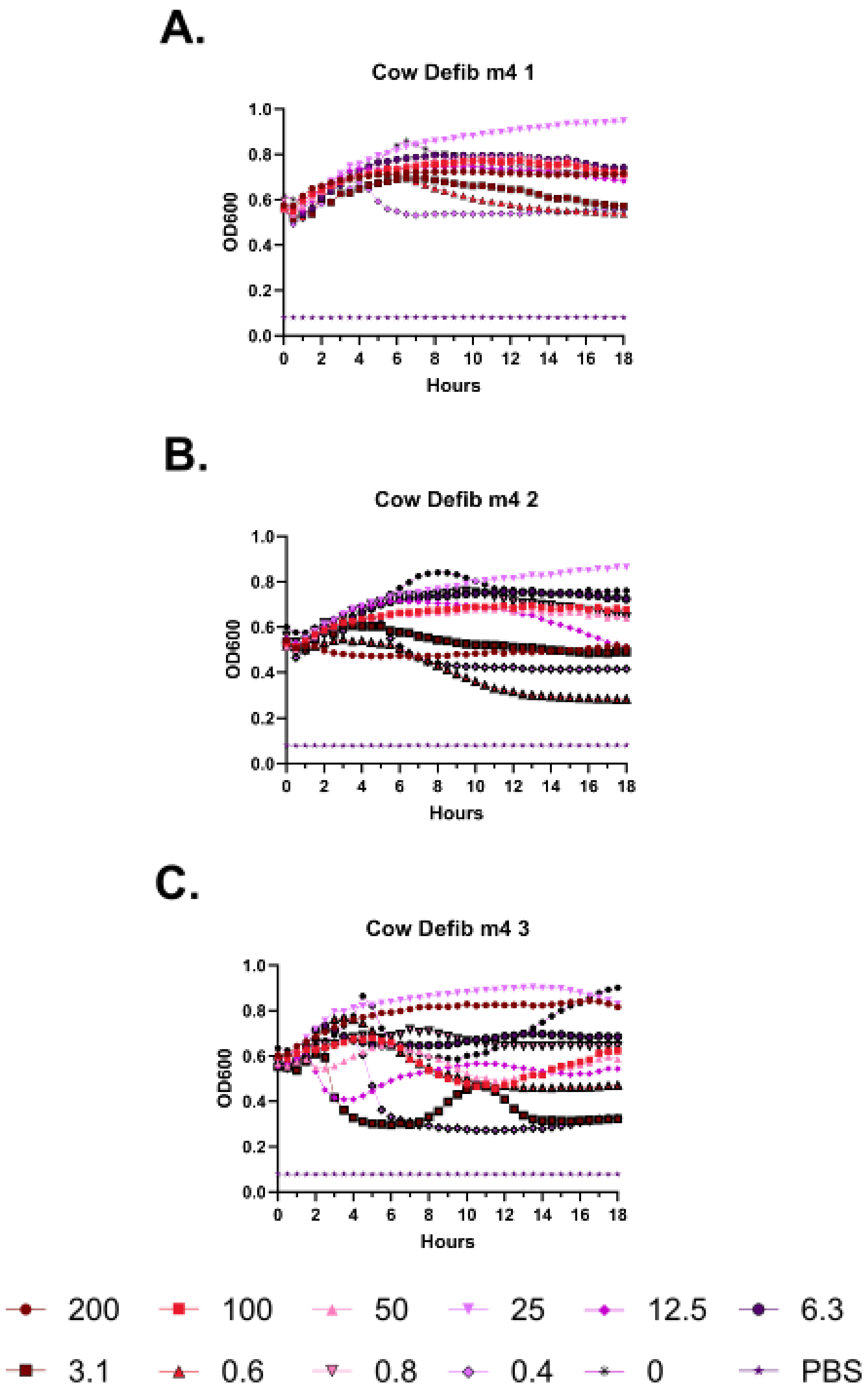
A.
Cow Defib m4 1
OD600
Hours
0.0
0.2
0.4
0.6
0.8
1.0
0
2
4
6
8
10
12
14
16
18
B.
Cow Defib m4 2
OD600
Hours
C.
Cow Defib m4 3
OD600
Hours
200
100
50
25
12.5
6.3
3.1
0.6
0.8
0.4
0
PBS

**Supplementary Figure 62 – Kinetic hemolysis assays for m4 in defibrinated cow blood. A-C.** Hemolysis was monitored by OD600 over 18 hours at 37 °C in the presence of the indicated peptide concentrations. Decreasing OD600 indicates increasing hemolysis. Peptide concentrations are given in μM. Each curve indicates an independent experiment.

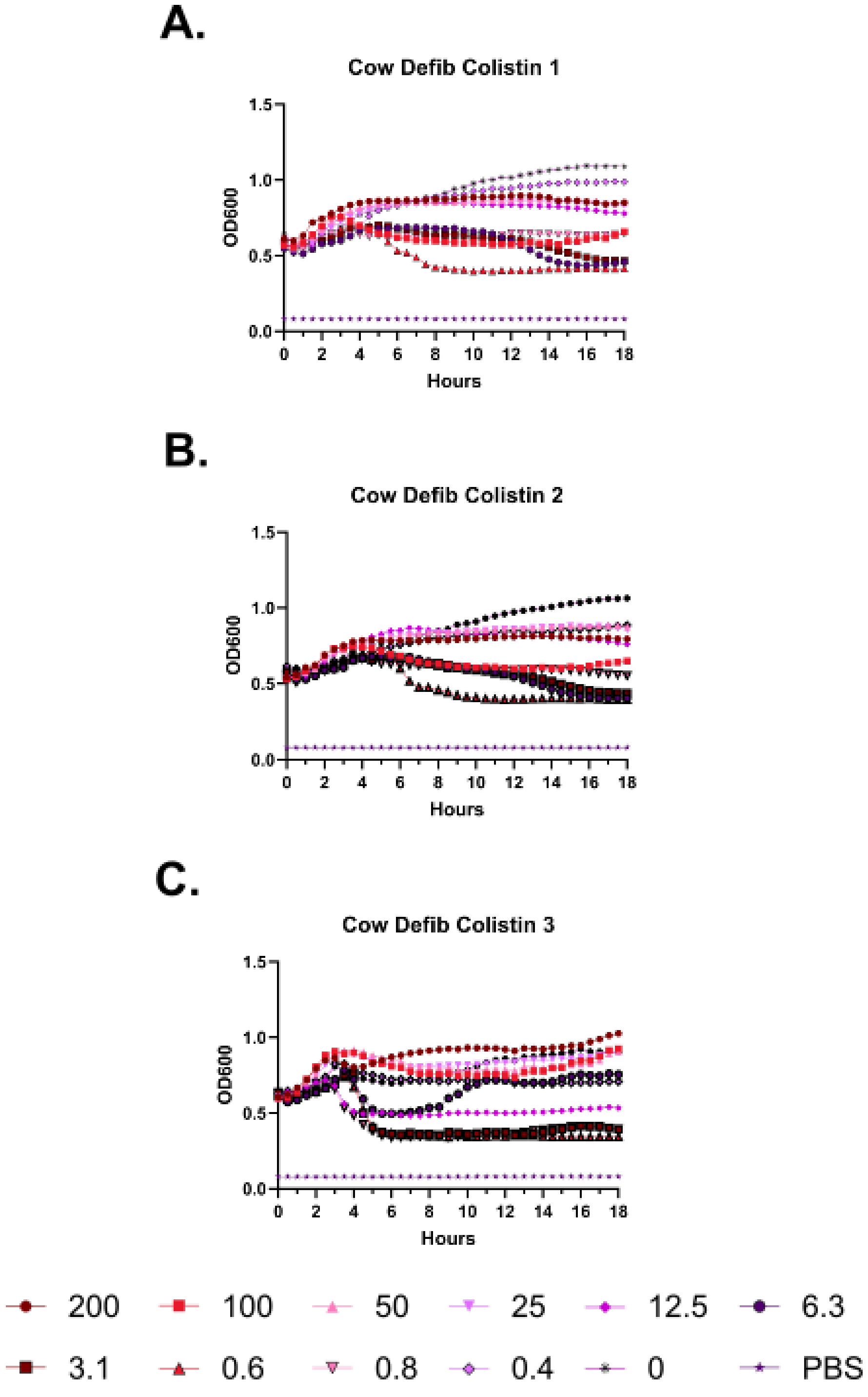
A.
Cow Defib Colistin 1
OD600
Hours
B.
Cow Defib Colistin 2
OD600
Hours
C.
Cow Defib Colistin 3
OD600
Hours
200
100
50
25
12.5
6.3
3.1
0.6
0.8
0.4
0
PBS

**Supplementary Figure 63 – Kinetic hemolysis assays for colistin in defibrinated cow blood. A-C.** Hemolysis was monitored by OD600 over 18 hours at 37 °C in the presence of the indicated peptide concentrations. Decreasing OD600 indicates increasing hemolysis. Peptide concentrations are given in μM. Each curve indicates an independent experiment.

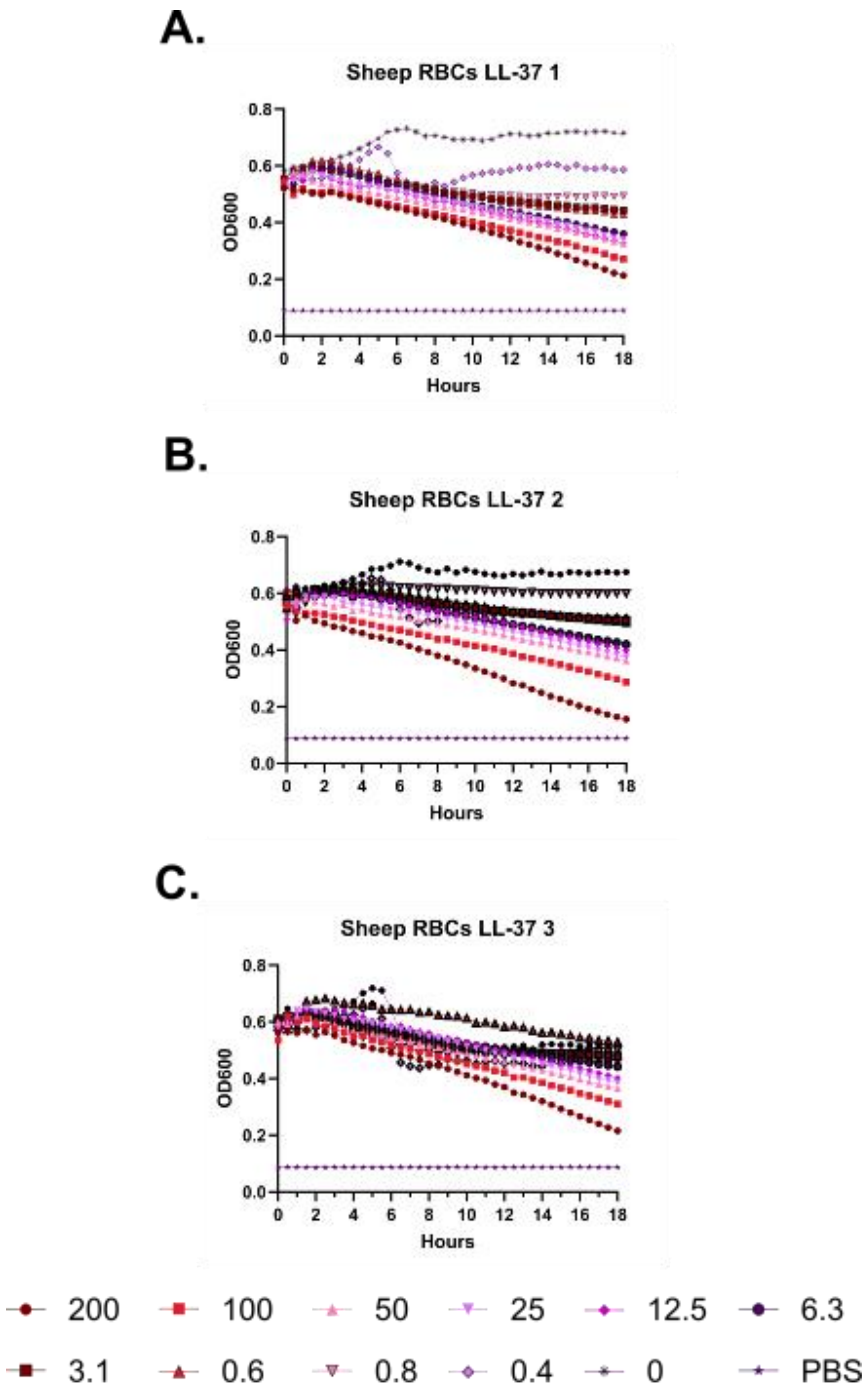
A.
Sheep RBCs LL-37 1
OD600
Hours
B.
Sheep RBCs LL-37 2
OD600
Hours
C.
Sheep RBCs LL-37 3
OD600
Hours
200
100
50
25
12.5
6.3
3.1
0.6
0.8
0.4
0
PBS

**Supplementary Figure 64 – Kinetic hemolysis assays for LL-37 in sheep red blood cells.** **A-C.** Hemolysis was monitored by OD600 over 18 hours at 37 °C in the presence of the indicated peptide concentrations. Decreasing OD600 indicates increasing hemolysis. Peptide concentrations are given in μM. Each curve indicates an independent experiment.

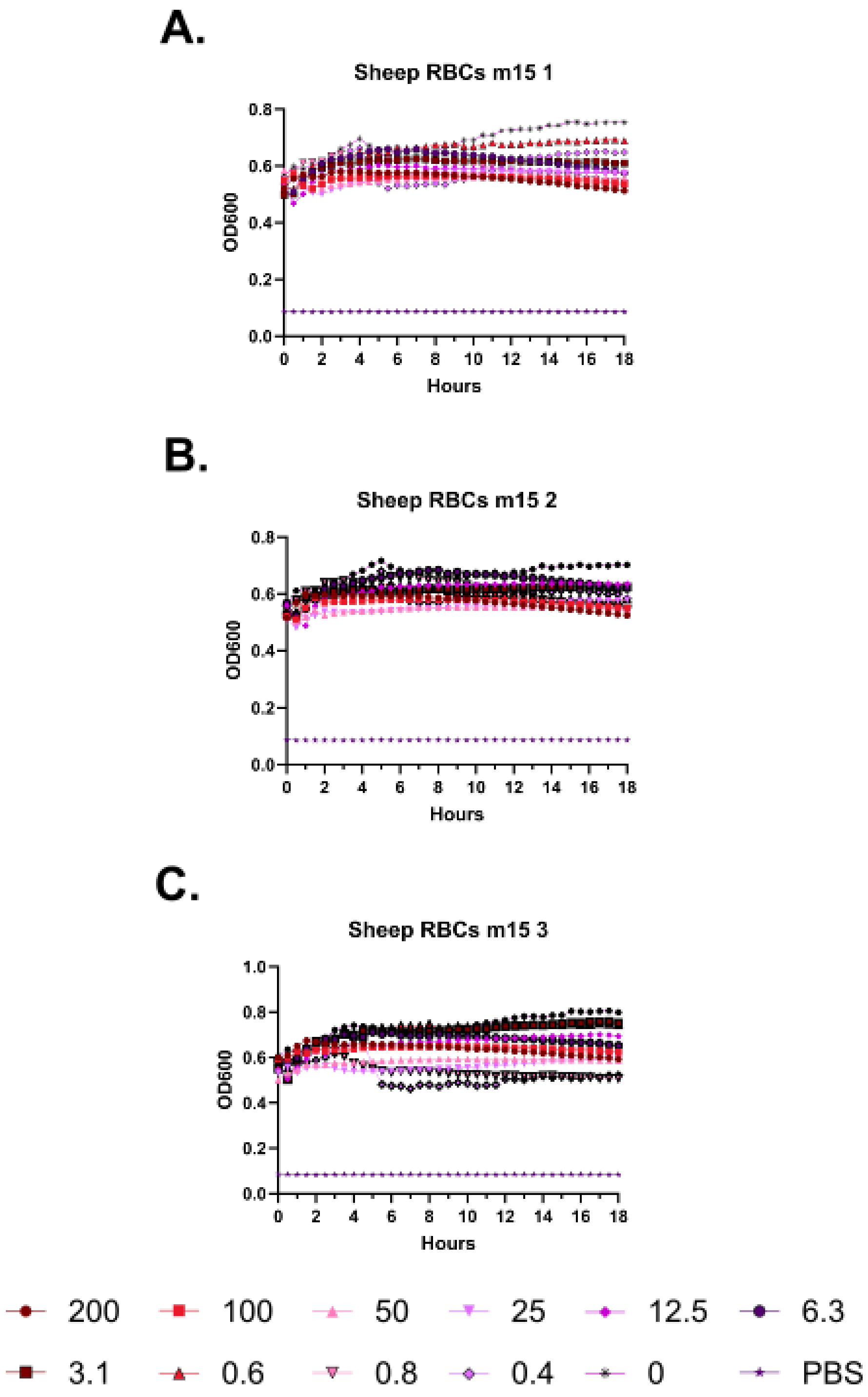
A.
Sheep RBCs m15 1
OD600
Hours
B.
Sheep RBCs m15 2
OD600
Hours
C.
Sheep RBCs m15 3
OD600
Hours
200
100
50
25
12.5
6.3
3.1
0.6
0.8
0.4
0
PBS

**Supplementary Figure 65 – Kinetic hemolysis assays for m15 in sheep red blood cells. A-C.** Hemolysis was monitored by OD600 over 18 hours at 37 °C in the presence of the indicated peptide concentrations. Decreasing OD600 indicates increasing hemolysis. Peptide concentrations are given in μM. Each curve indicates an independent experiment.

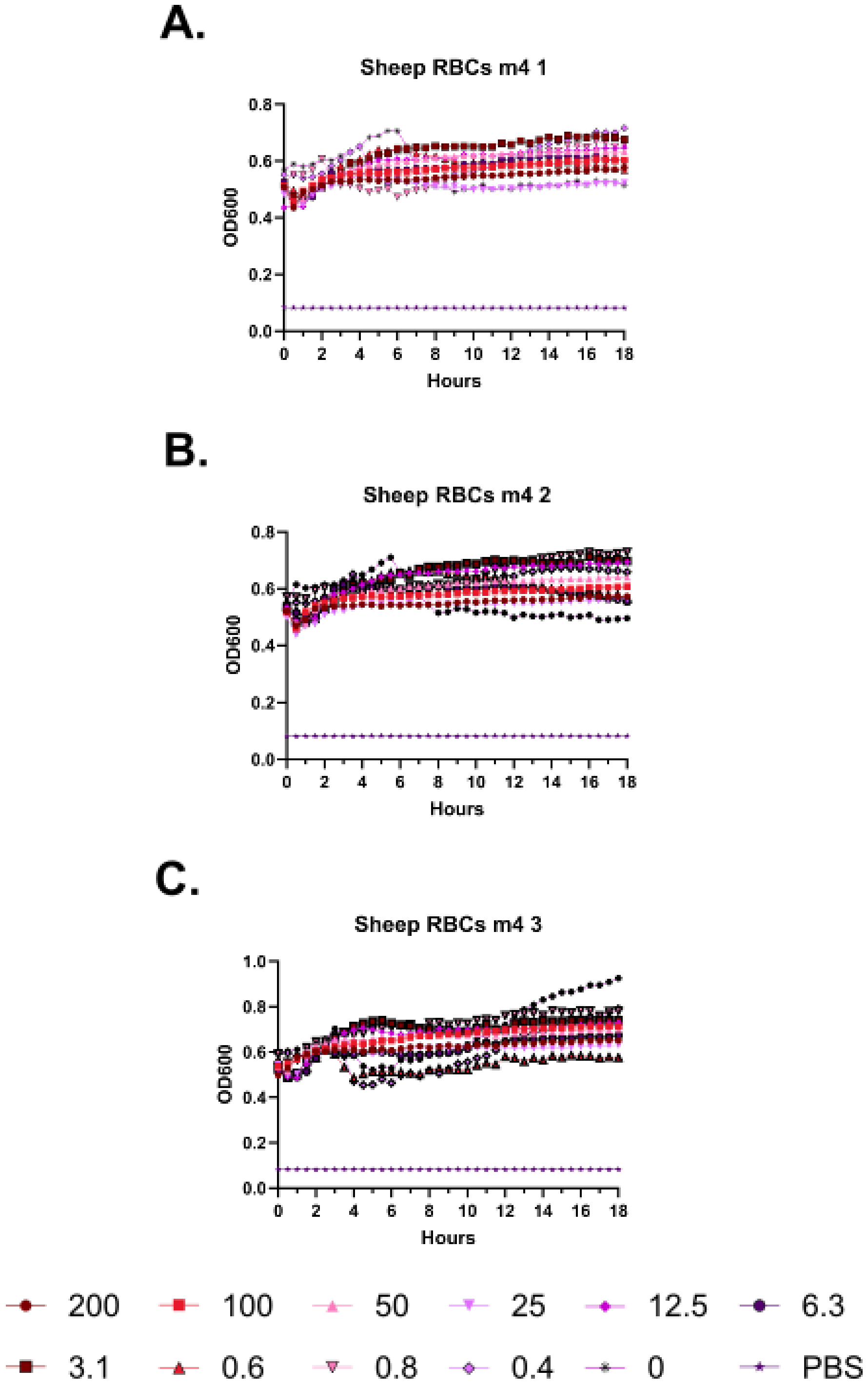
A.
Sheep RBCs m4 1
OD600
Hours
B.
Sheep RBCs m4 2
OD600
Hours
C.
Sheep RBCs m4 3
OD600
Hours
200
100
50
25
12.5
6.3
3.1
0.6
0.8
0.4
0
PBS

**Supplementary Figure 66 – Kinetic hemolysis assays for m4 in sheep red blood cells. A-C.** Hemolysis was monitored by OD600 over 18 hours at 37 °C in the presence of the indicated peptide concentrations. Decreasing OD600 indicates increasing hemolysis. Peptide concentrations are given in μM. Each curve indicates an independent experiment.

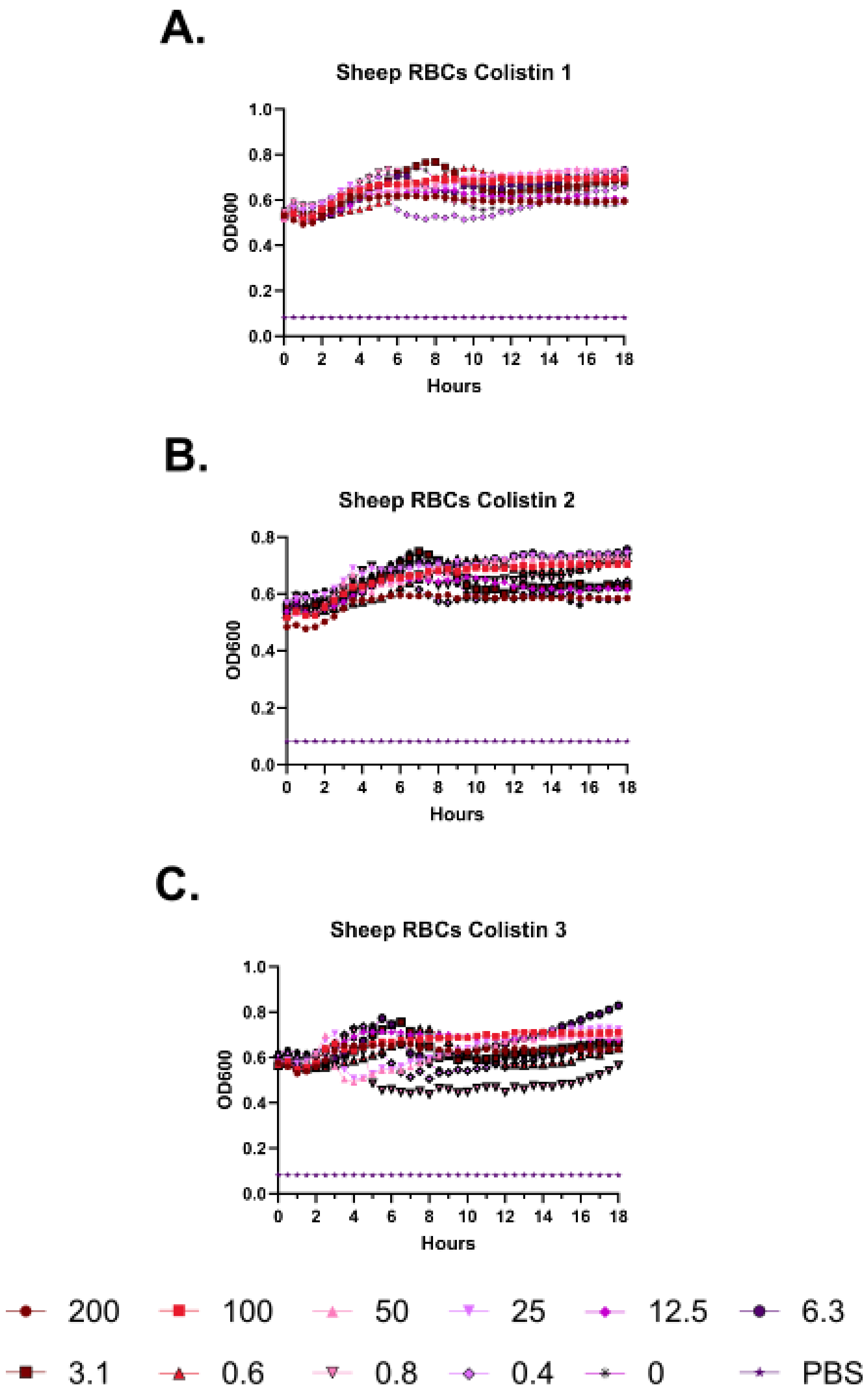
A.
Sheep RBCs Colistin 1
OD600
Hours
B.
Sheep RBCs Colistin 2
OD600
Hours
C.
Sheep RBCs Colistin 3
OD600
Hours
200
100
50
25
12.5
6.3
3.1
0.6
0.8
0.4
0
PBS

**Supplementary Figure 67 – Kinetic hemolysis assays for colistin in sheep red blood cells. A-C.** Hemolysis was monitored by OD600 over 18 hours at 37 °C in the presence of the indicated peptide concentrations. Decreasing OD600 indicates increasing hemolysis. Peptide concentrations are given in μM. Each curve indicates an independent experiment.

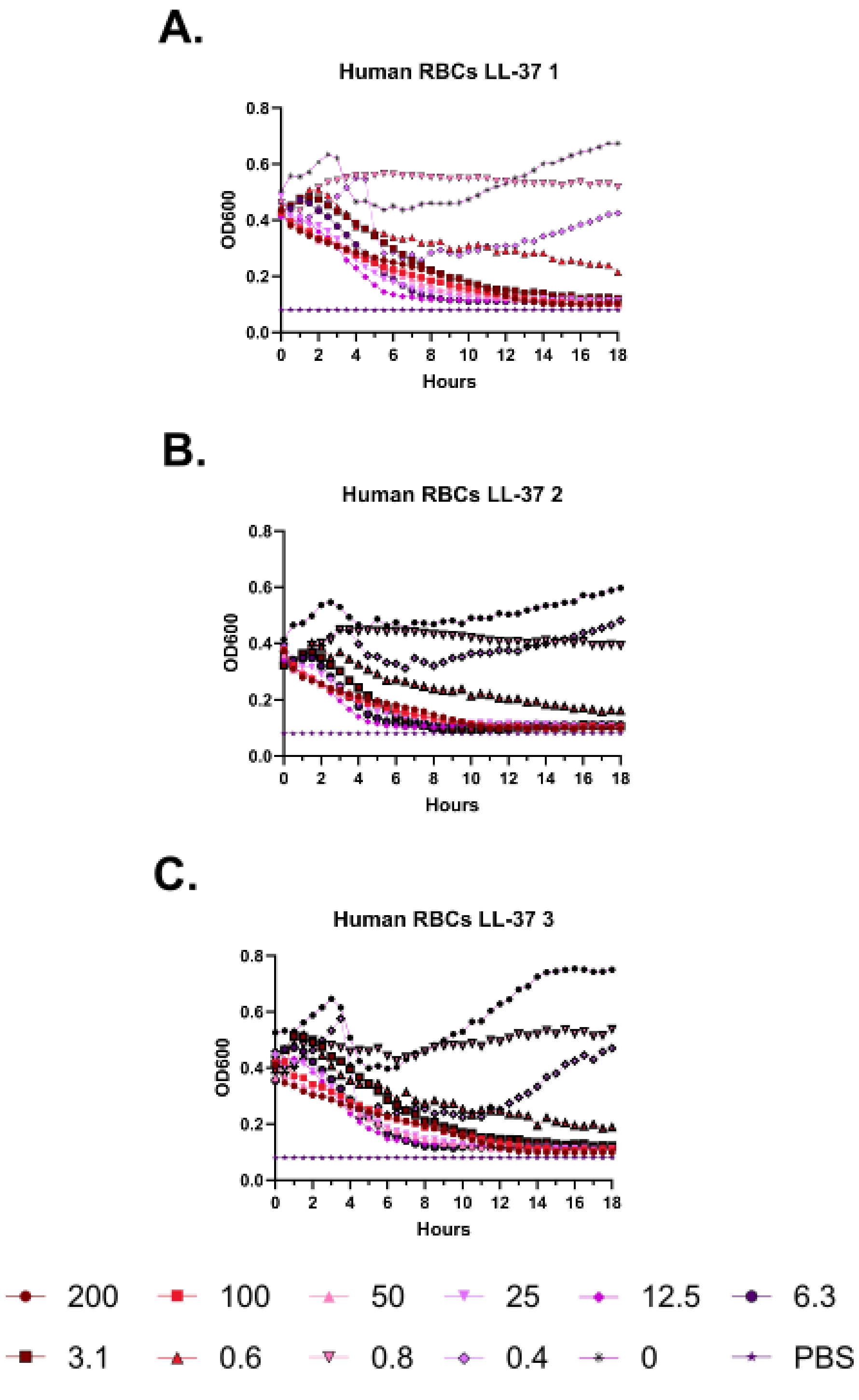
A.
Human RBCs LL-37 1
OD600
Hours
B.
Human RBCs LL-37 2
OD600
Hours
C.
Human RBCs LL-37 3
OD600
Hours
0.0
0.2
0.4
0.6
0.8
0
2
4
6
8
10
12
14
16
18
200
100
50
25
12.5
6.3
3.1
0.6
0.8
0.4
0
PBS

**Supplementary Figure 68 – Kinetic hemolysis assays for LL-37 in human red blood cells. A-C.** Hemolysis was monitored by OD600 over 18 hours at 37 °C in the presence of the indicated peptide concentrations. Decreasing OD600 indicates increasing hemolysis. Peptide concentrations are given in μM. Each curve indicates an independent experiment.

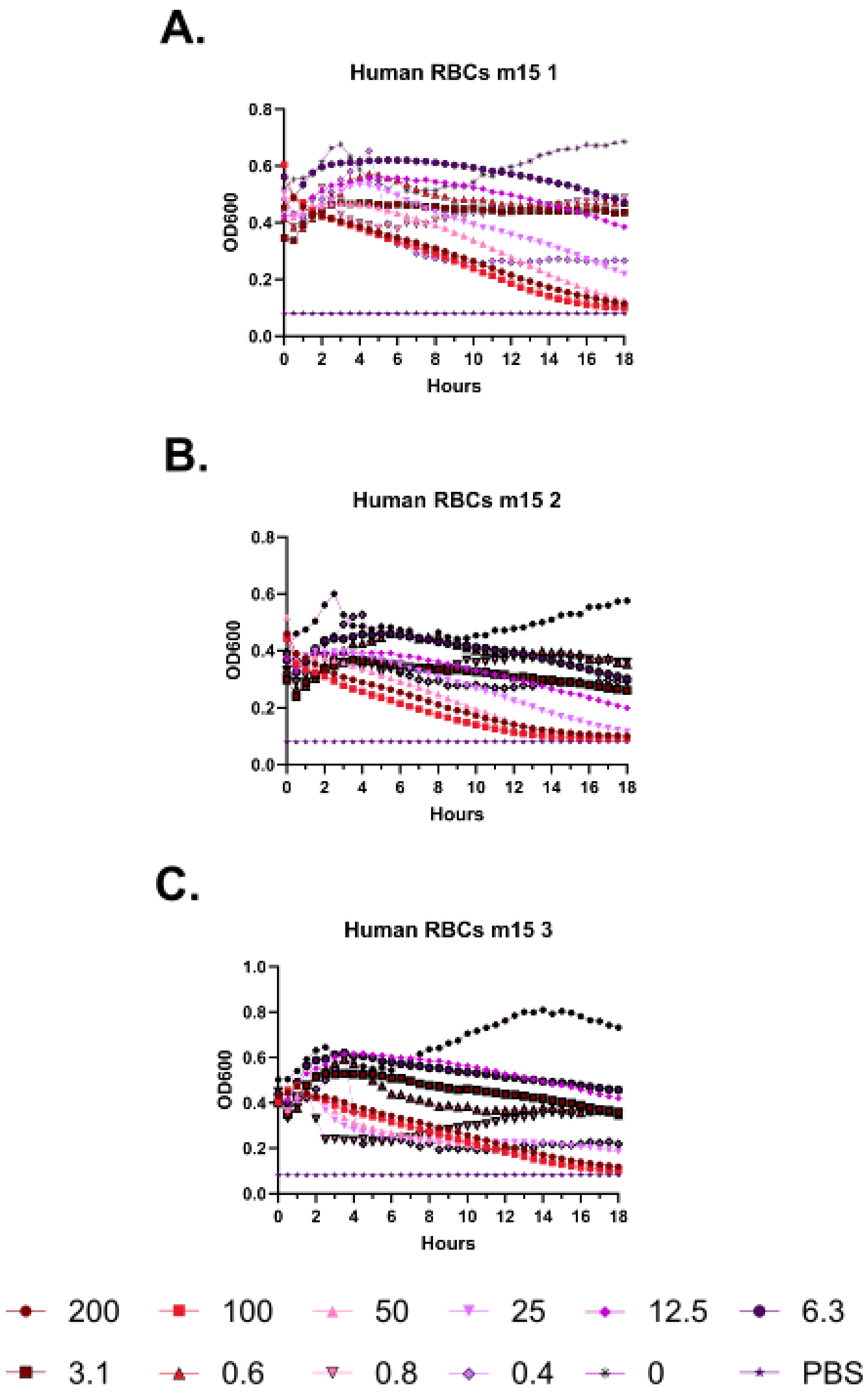

A.
Human RBCs m15 1
OD600
0.0
0.2
0.4
0.6
0.8
0 2 4 6 8 10 12 14 16 18
Hours
B.
Human RBCs m15 2
OD600
0.0
0.2
0.4
0.6
0.8
0 2 4 6 8 10 12 14 16 18
Hours
C.
Human RBCs m15 3
OD600
0.0
0.2
0.4
0.6
0.8
1.0
0 2 4 6 8 10 12 14 16 18
Hours
200
100
50
25
12.5
6.3
3.1
0.6
0.8
0.4
0
PBS

**Supplementary Figure 69 – Kinetic hemolysis assays for m15 in human red blood cells. A-C.** Hemolysis was monitored by OD600 over 18 hours at 37 °C in the presence of the indicated peptide concentrations. Decreasing OD600 indicates increasing hemolysis. Peptide concentrations are given in µM. Each curve indicates an independent experiment.

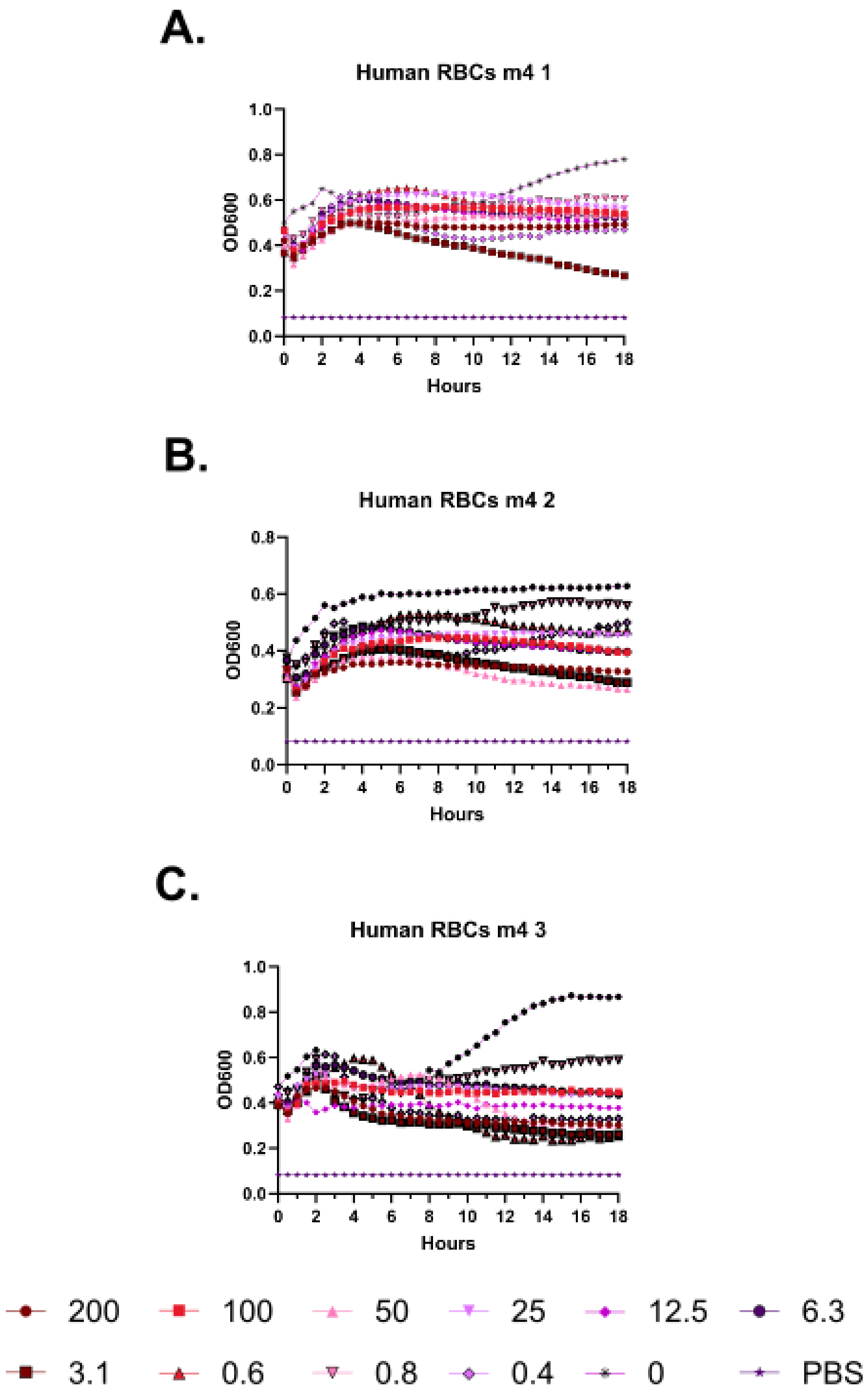
A.
Human RBCs m4 1
OD600
Hours
B.
Human RBCs m4 2
OD600
Hours
C.
Human RBCs m4 3
OD600
Hours
200
100
50
25
12.5
6.3
3.1
0.6
0.8
0.4
0
PBS

**Supplementary Figure 70 – Kinetic hemolysis assays for m4 in human red blood cells. A-C.** Hemolysis was monitored by OD600 over 18 hours at 37 °C in the presence of the indicated peptide concentrations. Decreasing OD600 indicates increasing hemolysis. Peptide concentrations are given in μM. Each curve indicates an independent experiment.

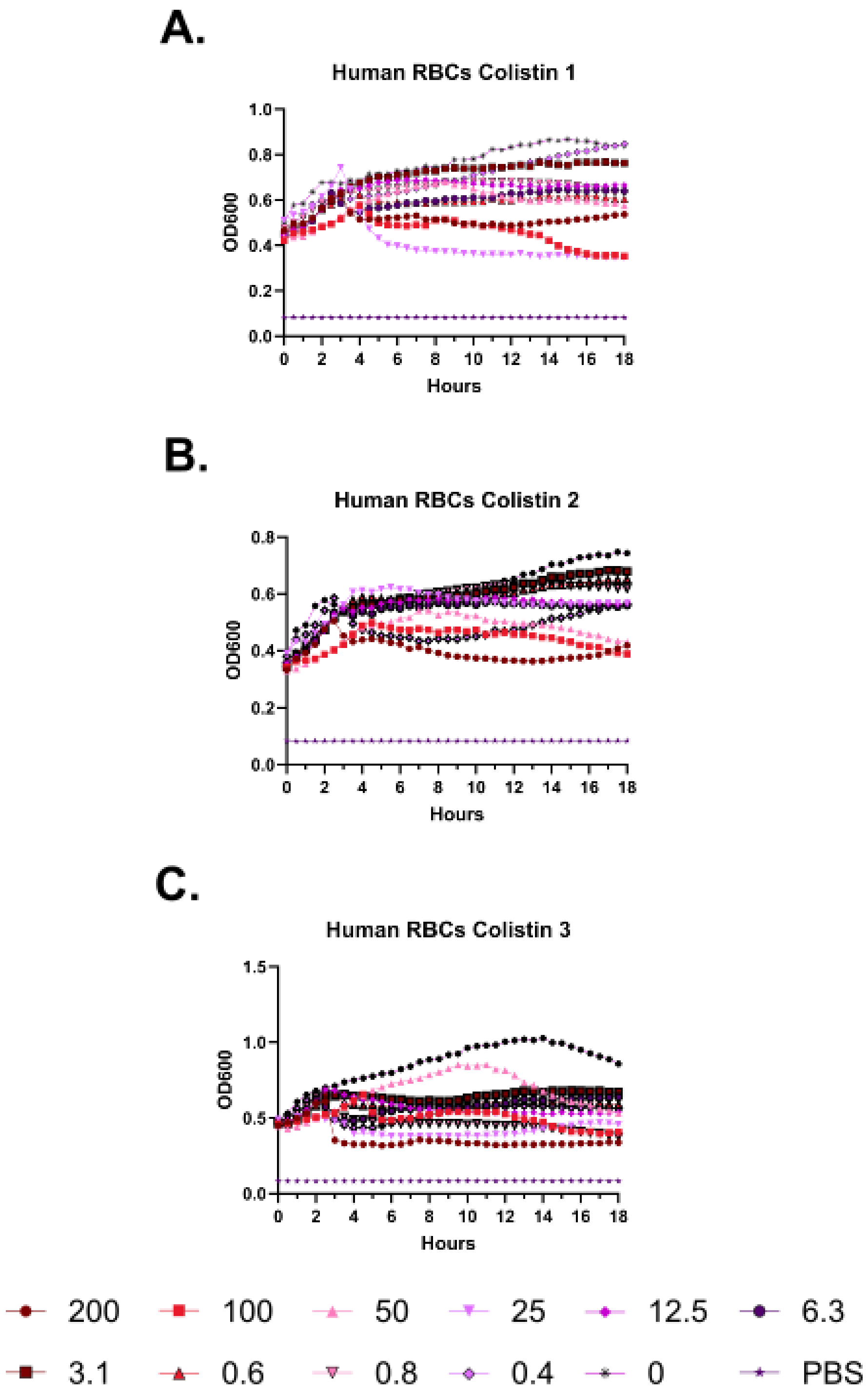
A.
Human RBCs Colistin 1
OD600
Hours
B.
Human RBCs Colistin 2
OD600
Hours
C.
Human RBCs Colistin 3
OD600
Hours
200
100
50
25
12.5
6.3
3.1
0.6
0.8
0.4
0
PBS

**Supplementary Figure 71 – Kinetic hemolysis assays for colistin in human red blood cells. A-C.** Hemolysis was monitored by OD600 over 18 hours at 37 °C in the presence of the indicated peptide concentrations. Decreasing OD600 indicates increasing hemolysis. Peptide concentrations are given in μM. Each curve indicates an independent experiment.

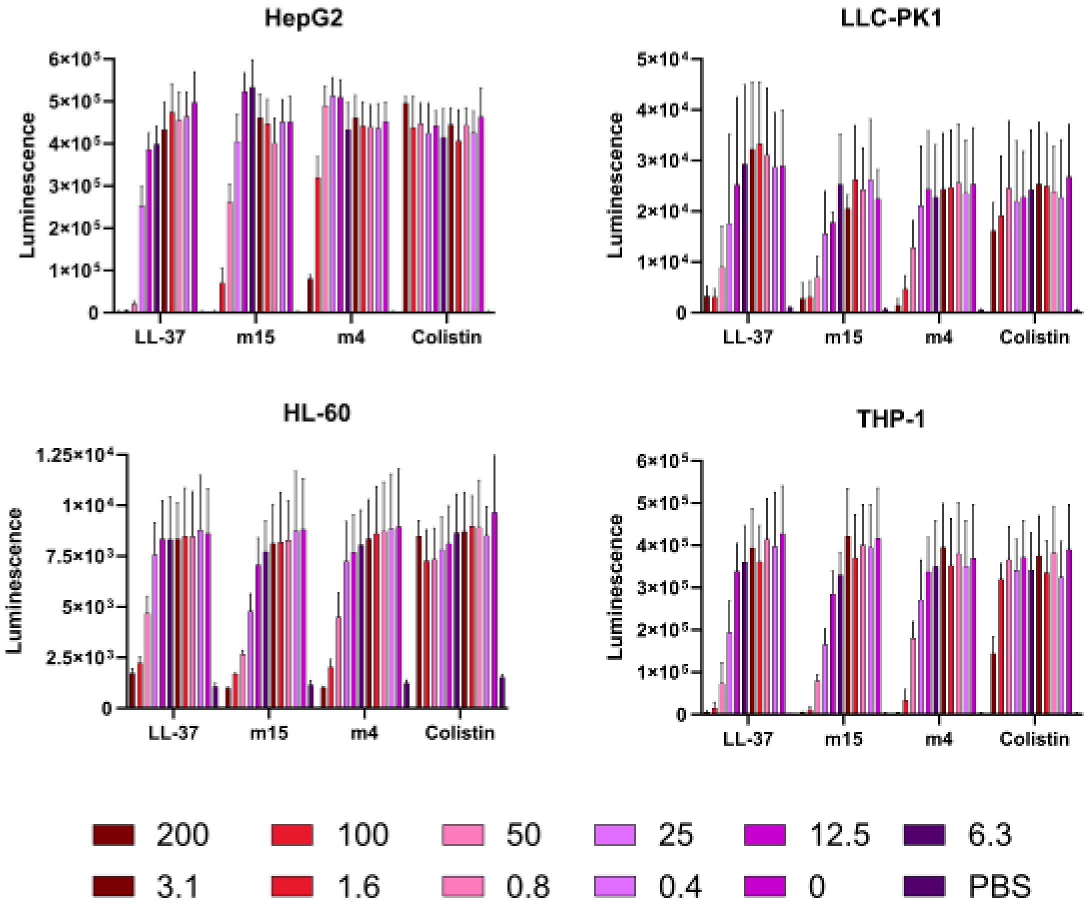
HepG2
Luminescence
6×10⁵
5×10⁵
4×10⁵
3×10⁵
2×10⁵
1×10⁵
0
LL-37
m15
m4
Colistin
LLC-PK1
Luminescence
5×10⁴
4×10⁴
3×10⁴
2×10⁴
1×10⁴
0
LL-37
m15
m4
Colistin
HL-60
Luminescence
1.25×10⁴
1×10⁴
7.5×10³
5×10³
2.5×10³
0
LL-37
m15
m4
Colistin
THP-1
Luminescence
6×10⁵
5×10⁵
4×10⁵
3×10⁵
2×10⁵
1×10⁵
0
LL-37
m15
m4
Colistin
200
100
50
25
12.5
6.3
3.1
1.6
0.8
0.4
0
PBS

**Supplementary Figure 72 – Histogram depiction of Figure 3E data. A.** Endpoint of RealTime Glo luciferase toxicity assay using the indicated cell types. Data are the mean and standard deviation from the raw luminescence measures of three independent experiments.

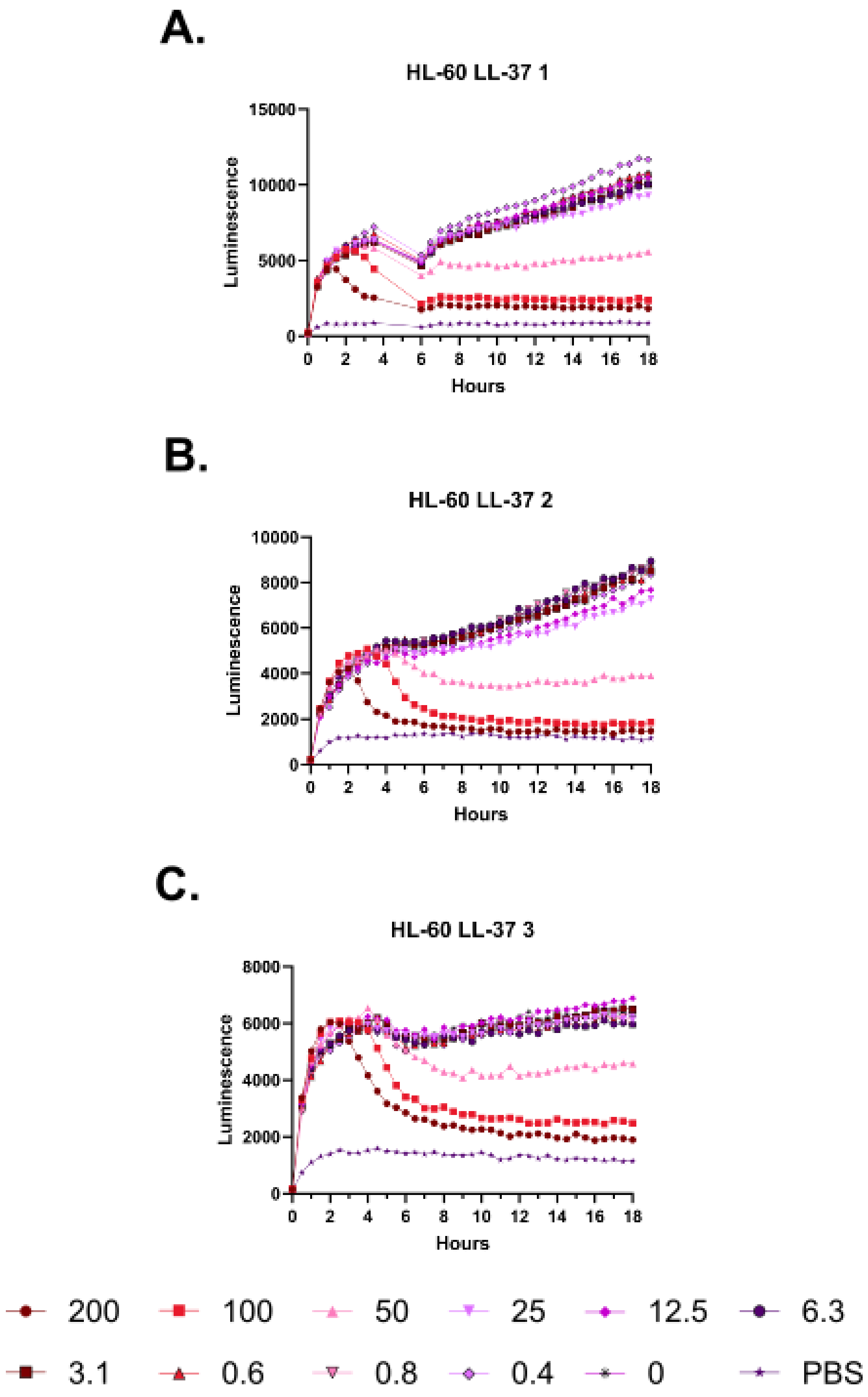
A.
HL-60 LL-37 1
Luminescence
Hours
B.
HL-60 LL-37 2
Luminescence
Hours
C.
HL-60 LL-37 3
Luminescence
Hours
200
100
50
25
12.5
6.3
3.1
0.6
0.8
0.4
0
PBS

**Supplementary Figure 73 – Kinetic luminescence toxicity assay for HL-60 treated with LL-37. A-C.** Luminescence was monitored over 18 hours at 37 °C with 5% $CO_2$ in a Small Humidity Cassette in the presence of the indicated peptide concentrations. Toxicity is indicated by curves that fall off of a steady upward trajectory. An instrument error in Experiment 1 resulted in no reads being taken during the gap in the curve. Peptide concentrations are given in μM. Each curve indicates an independent experiment.

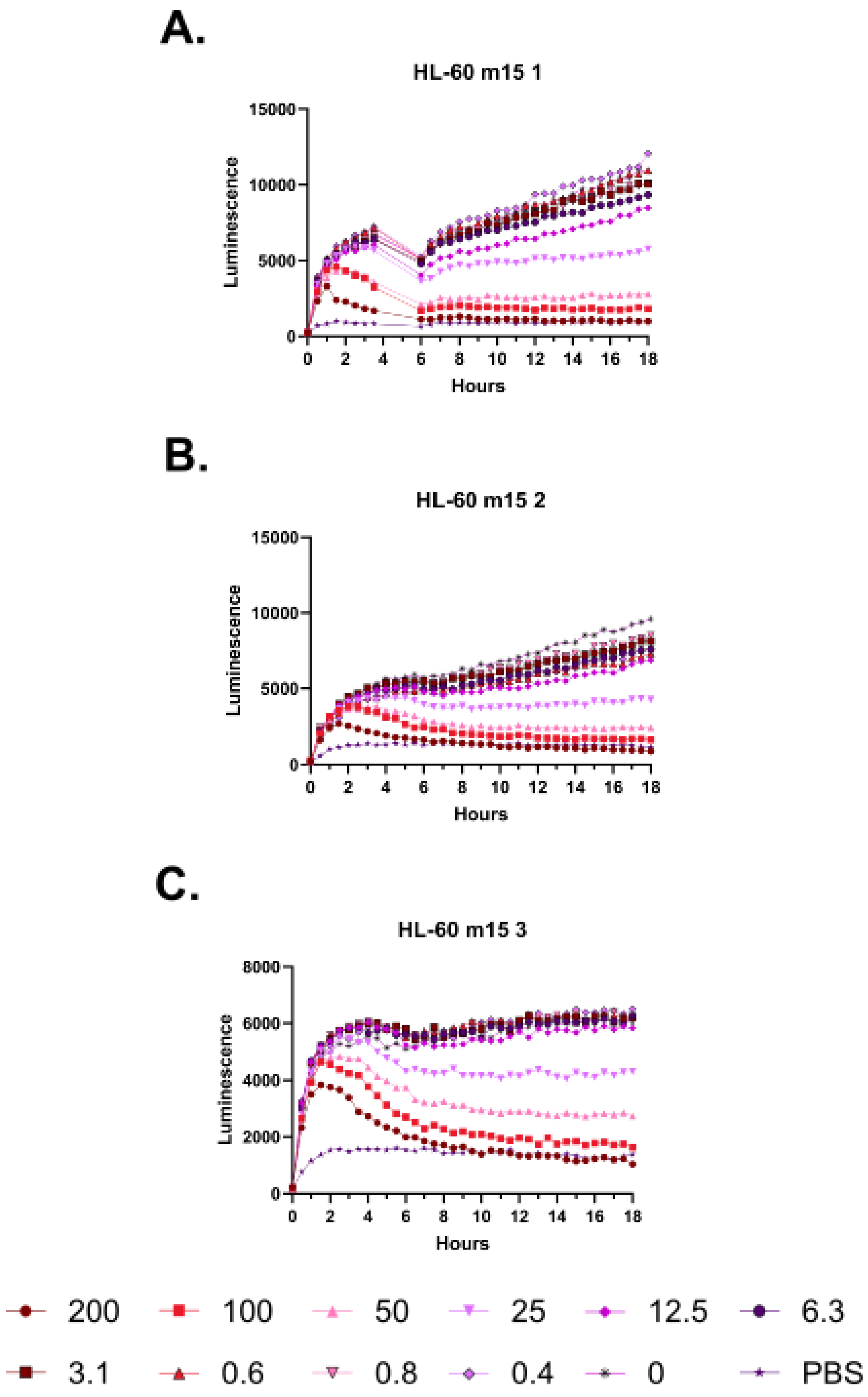
A.
HL-60 m15 1
B.
HL-60 m15 2
C.
HL-60 m15 3
Luminescence
Hours
200
100
50
25
12.5
6.3
3.1
0.6
0.8
0.4
0
PBS

**Supplementary Figure 74 – Kinetic luminescence toxicity assay for HL-60 treated with m15. A-C.** Luminescence was monitored over 18 hours at 37 °C with 5% $CO_2$ in a Small Humidity Cassette in the presence of the indicated peptide concentrations. Toxicity is indicated by curves that fall off of a steady upward trajectory. An instrument error in Experiment 1 resulted in no reads being taken during the gap in the curve. Peptide concentrations are given in μM. Each curve indicates an independent experiment.

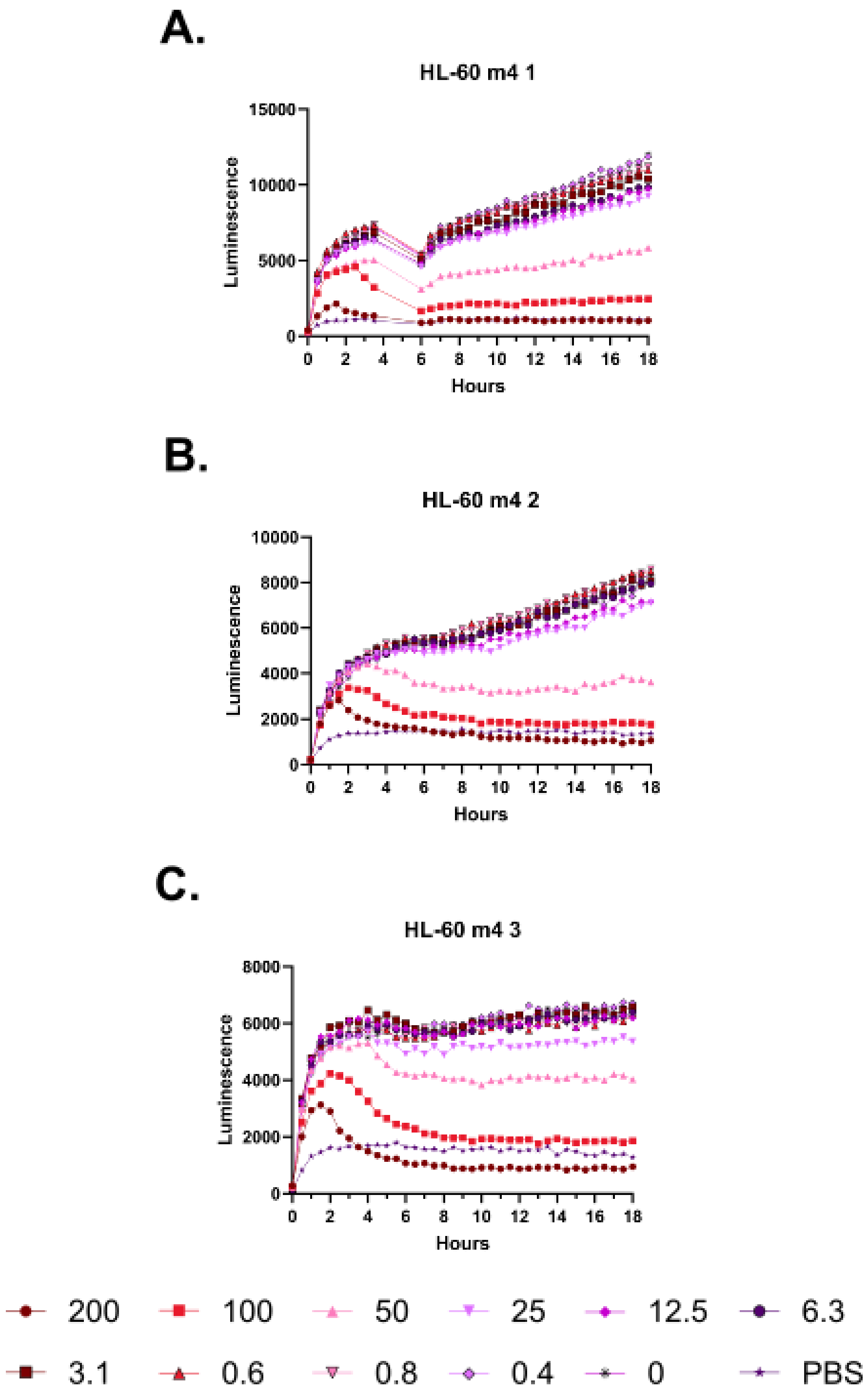
A.
HL-60 m4 1
Luminescence
Hours
B.
HL-60 m4 2
Luminescence
Hours
C.
HL-60 m4 3
Luminescence
Hours
200
100
50
25
12.5
6.3
3.1
0.6
0.8
0.4
0
PBS

**Supplementary Figure 75 – Kinetic luminescence toxicity assay for HL-60 treated with m4. A-C.** Luminescence was monitored over 18 hours at 37 °C with 5% $CO_2$ in a Small Humidity Cassette in the presence of the indicated peptide concentrations. Toxicity is indicated by curves that fall off of a steady upward trajectory. An instrument error in Experiment 1 resulted in no reads being taken during the gap in the curve. Peptide concentrations are given in μM. Each curve indicates an independent experiment.

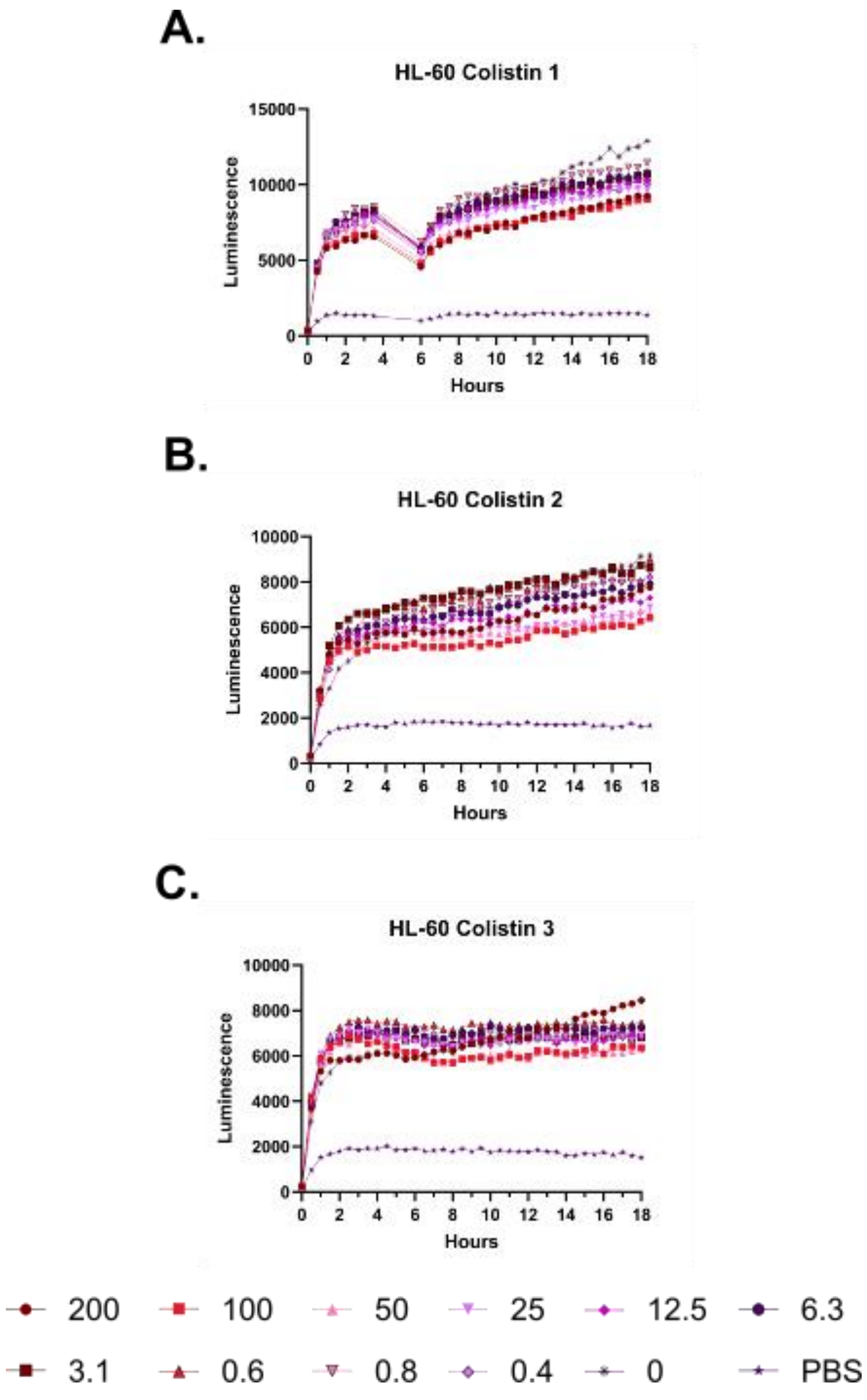

A.
HL-60 Colistin 1
Luminescence
Hours
B.
HL-60 Colistin 2
Luminescence
Hours
C.
HL-60 Colistin 3
Luminescence
Hours
200
100
50
25
12.5
6.3
3.1
0.6
0.8
0.4
0
PBS

**Supplementary Figure 76 – Kinetic luminescence toxicity assay for HL-60 treated with colistin. A-C.** Luminescence was monitored over 18 hours at 37 °C with 5% $CO_2$ in a Small Humidity Cassette in the presence of the indicated peptide concentrations. Toxicity is indicated by curves that fall off of a steady upward trajectory. An instrument error in Experiment 1 resulted in no reads being taken during the gap in the curve. Peptide concentrations are given in µM. Each curve indicates an independent experiment.

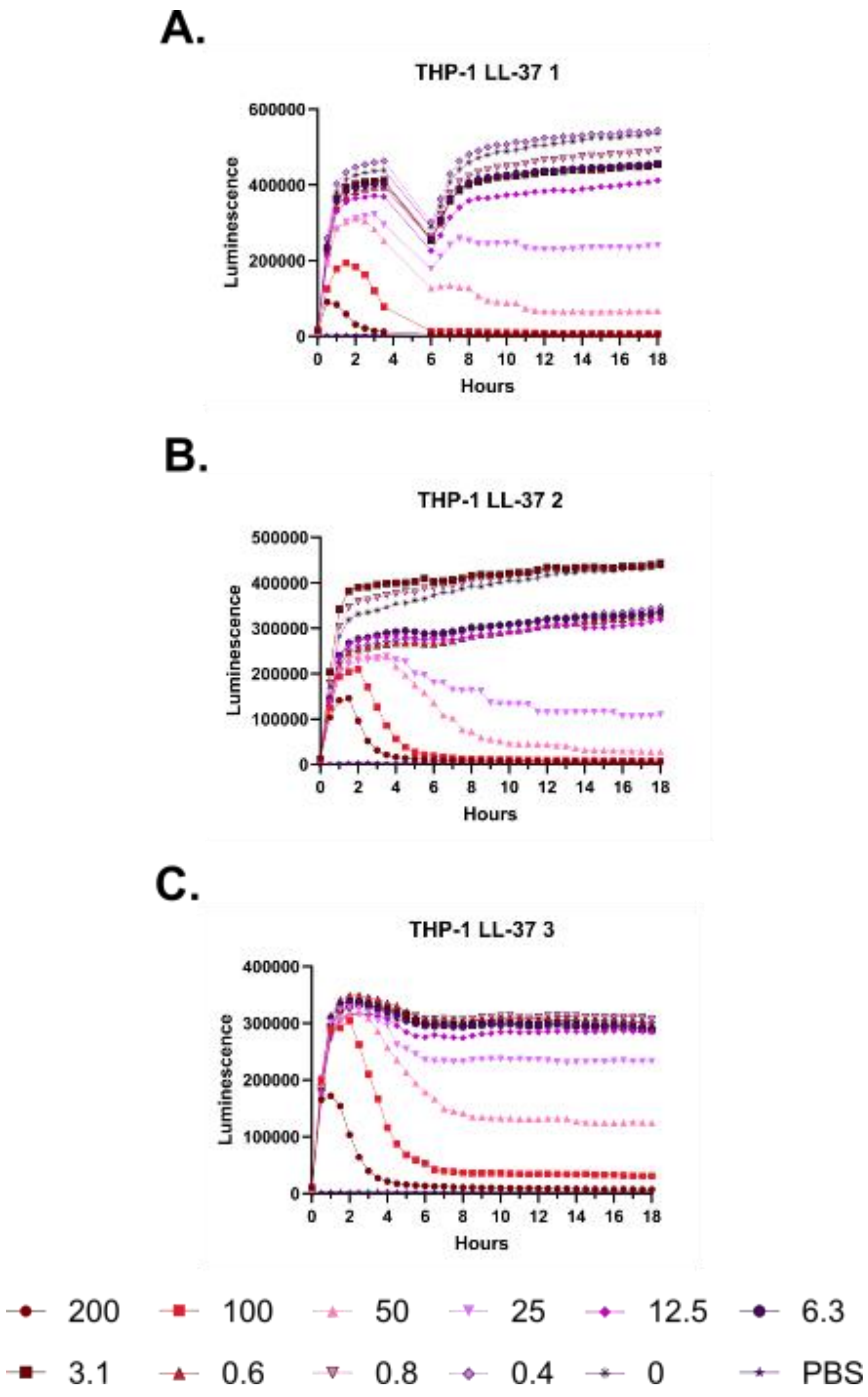
A.
THP-1 LL-37 1
Luminescence
Hours
B.
THP-1 LL-37 2
Luminescence
Hours
C.
THP-1 LL-37 3
Luminescence
Hours
200
100
50
25
12.5
6.3
3.1
0.6
0.8
0.4
0
PBS

**Supplementary Figure 77 – Kinetic luminescence toxicity assay for THP-1 treated with LL-37. A-C.** Luminescence was monitored over 18 hours at 37 °C with 5% $CO_2$ in a Small Humidity Cassette in the presence of the indicated peptide concentrations. Toxicity is indicated by curves that fall off of a steady upward trajectory. An instrument error in Experiment 1 resulted in no reads being taken during the gap in the curve. Peptide concentrations are given in μM. Each curve indicates an independent experiment.

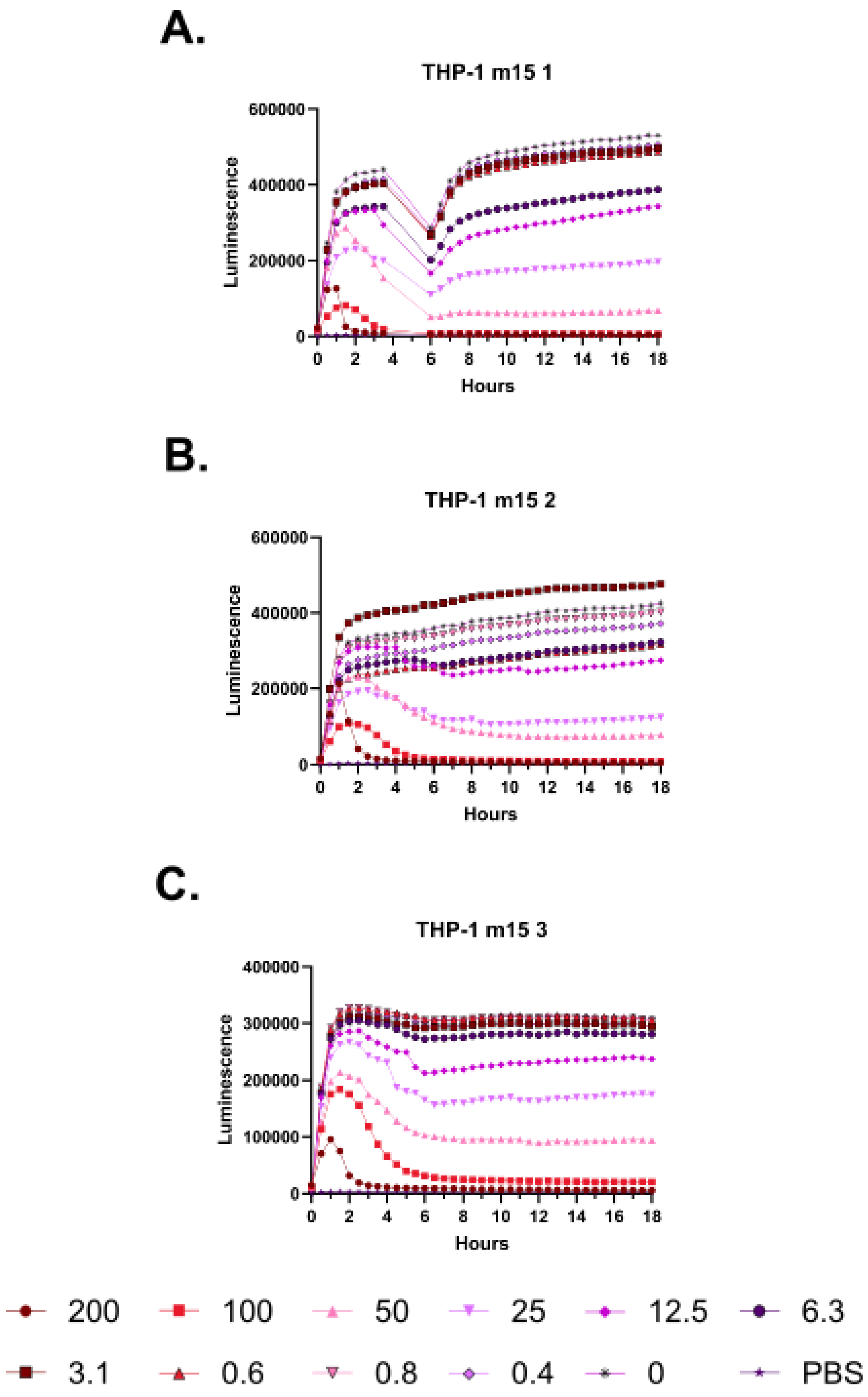

A.
THP-1 m15 1
600000
400000
200000
0
Luminescence
0 2 4 6 8 10 12 14 16 18
Hours
B.
THP-1 m15 2
600000
400000
200000
0
Luminescence
0 2 4 6 8 10 12 14 16 18
Hours
C.
THP-1 m15 3
400000
300000
200000
100000
0
Luminescence
0 2 4 6 8 10 12 14 16 18
Hours
200
100
50
25
12.5
6.3
3.1
0.6
0.8
0.4
0
PBS

**Supplementary Figure 78 – Kinetic luminescence toxicity assay for THP-1 treated with m15. A-C.** Luminescence was monitored over 18 hours at 37 °C with 5% $CO_2$ in a Small Humidity Cassette in the presence of the indicated peptide concentrations. Toxicity is indicated by curves that fall off of a steady upward trajectory. An instrument error in Experiment 1 resulted in no reads being taken during the gap in the curve. Peptide concentrations are given in μM. Each curve indicates an independent experiment.

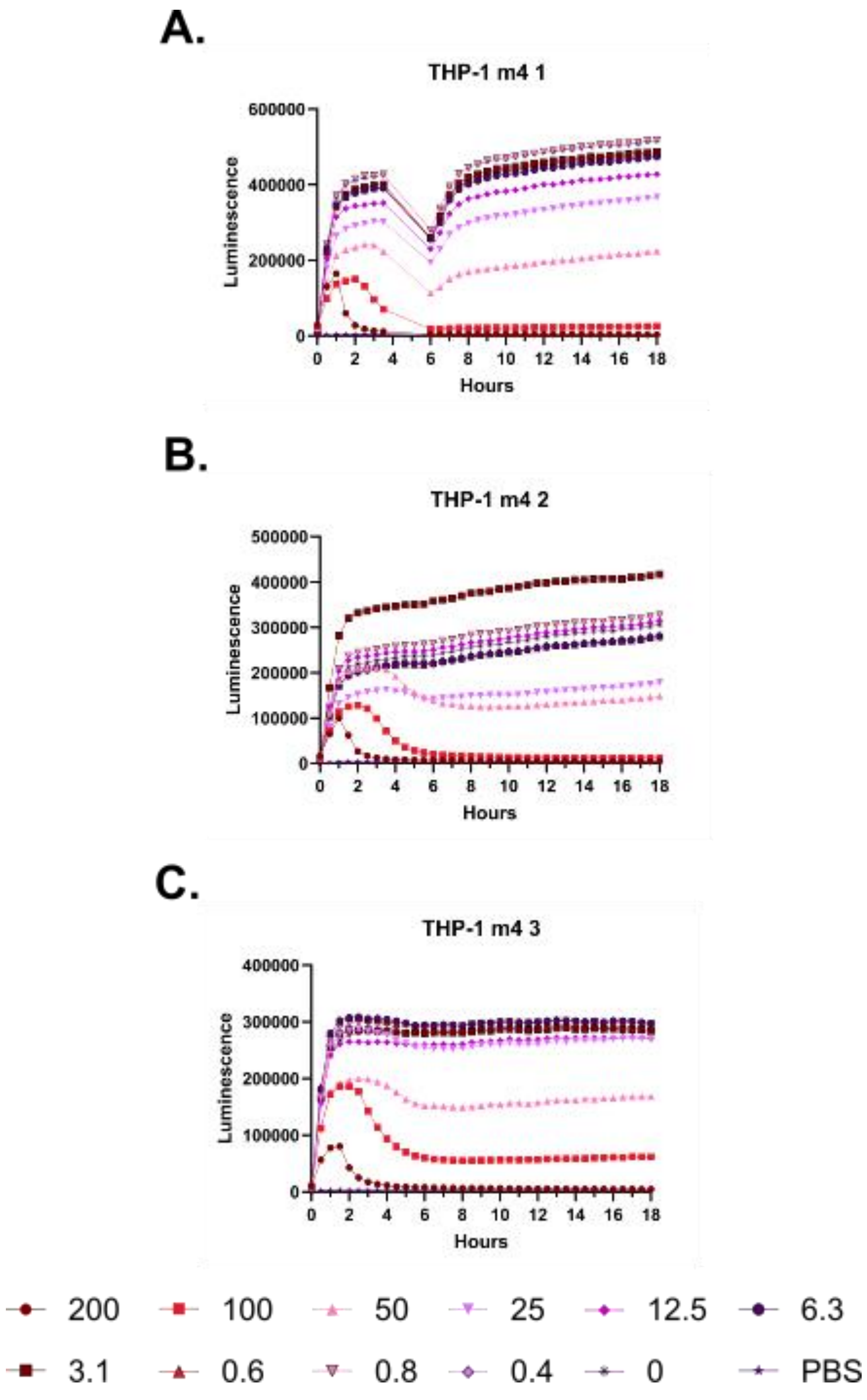
A.
THP-1 m4 1
Luminescence
Hours
B.
THP-1 m4 2
Luminescence
Hours
C.
THP-1 m4 3
Luminescence
Hours
200
100
50
25
12.5
6.3
3.1
0.6
0.8
0.4
0
PBS

**Supplementary Figure 79 – Kinetic luminescence toxicity assay for THP-1 treated with m4. A-C.** Luminescence was monitored over 18 hours at 37 °C with 5% $CO_2$ in a Small Humidity Cassette in the presence of the indicated peptide concentrations. Toxicity is indicated by curves that fall off of a steady upward trajectory. An instrument error in Experiment 1 resulted in no reads being taken during the gap in the curve. Peptide concentrations are given in μM. Each curve indicates an independent experiment.

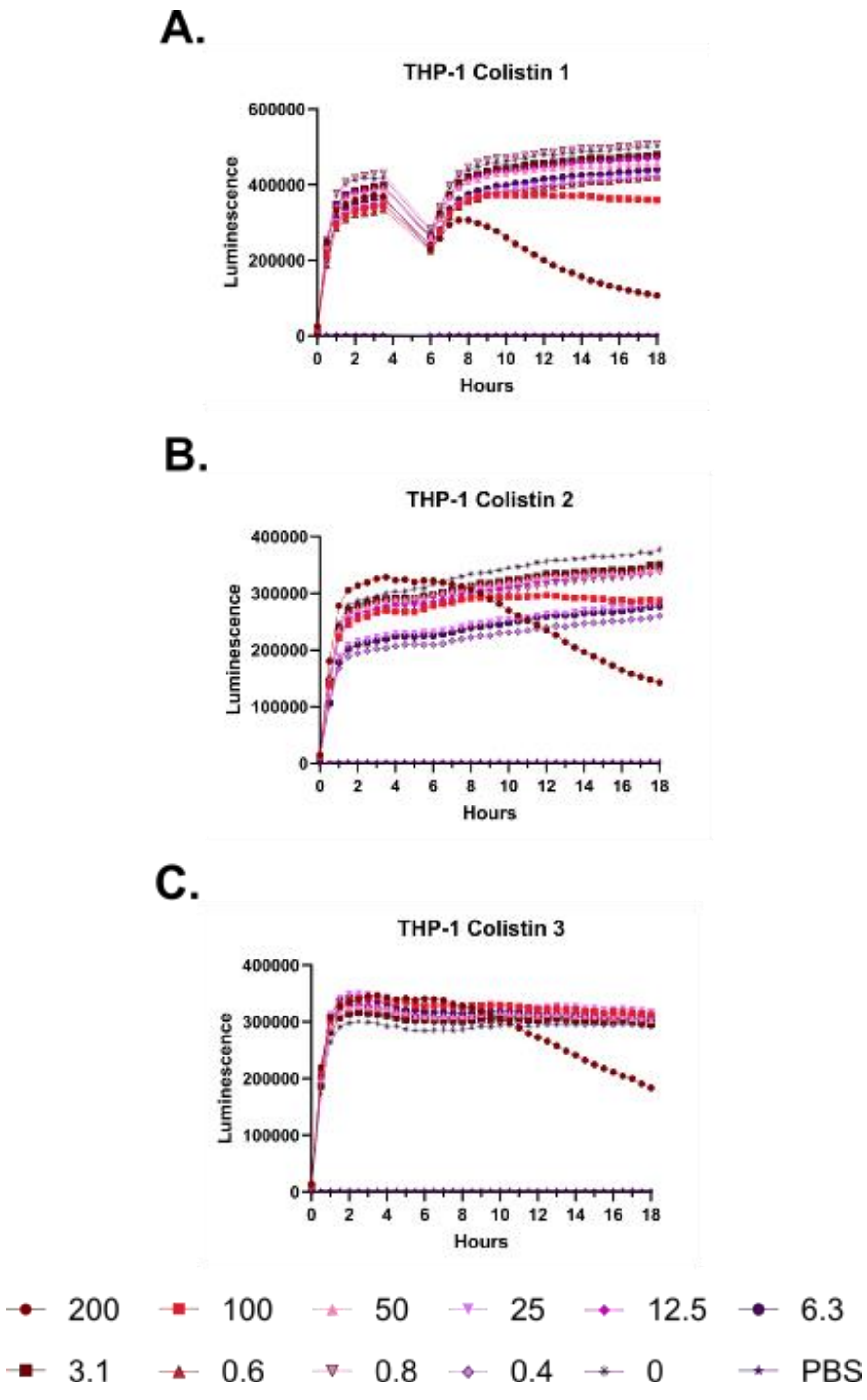

A.
THP-1 Colistin 1
Luminescence
600000
400000
200000
0
0 2 4 6 8 10 12 14 16 18
Hours
B.
THP-1 Colistin 2
Luminescence
400000
300000
200000
100000
0
0 2 4 6 8 10 12 14 16 18
Hours
C.
THP-1 Colistin 3
Luminescence
400000
300000
200000
100000
0
0 2 4 6 8 10 12 14 16 18
Hours
200
100
50
25
12.5
6.3
3.1
0.6
0.8
0.4
0
PBS

**Supplementary Figure 80 – Kinetic luminescence toxicity assay for THP-1 treated with colistin. A-C.** Luminescence was monitored over 18 hours at 37 °C with 5% $CO_2$ in a Small Humidity Cassette in the presence of the indicated peptide concentrations. Toxicity is indicated by curves that fall off of a steady upward trajectory. An instrument error in Experiment 1 resulted in no reads being taken during the gap in the curve. Peptide concentrations are given in μM. Each curve indicates an independent experiment.

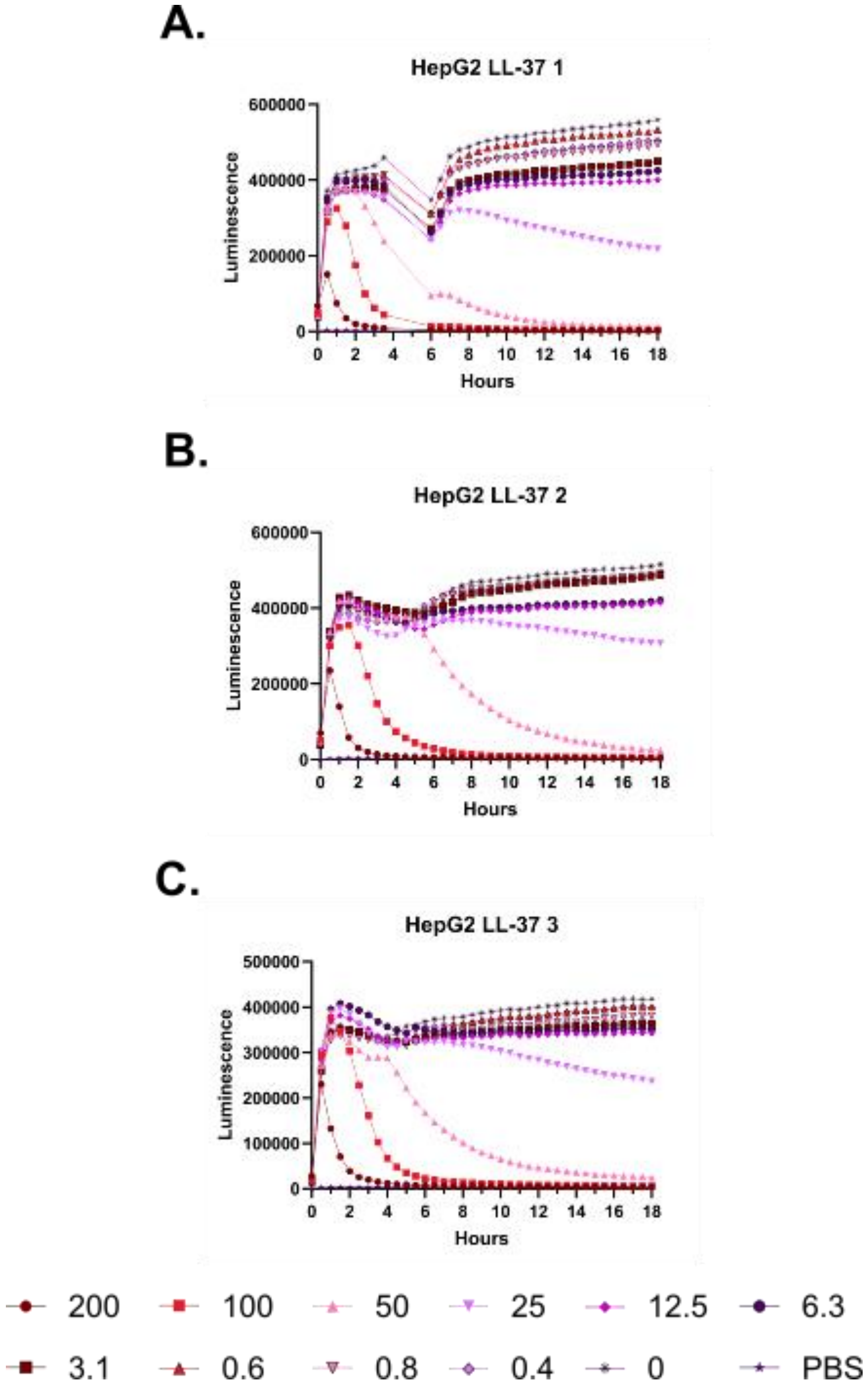
A.
HepG2 LL-37 1
Luminescence
Hours
B.
HepG2 LL-37 2
Luminescence
Hours
C.
HepG2 LL-37 3
Luminescence
Hours
200
100
50
25
12.5
6.3
3.1
0.6
0.8
0.4
0
PBS

**Supplementary Figure 81 – Kinetic luminescence toxicity assay for HepG2 treated with LL-37. A-C.** Luminescence was monitored over 18 hours at 37 °C with 5% $CO_2$ in a Small Humidity Cassette in the presence of the indicated peptide concentrations. Toxicity is indicated by curves that fall off of a steady upward trajectory. An instrument error in Experiment 1 resulted in no reads being taken during the gap in the curve. Peptide concentrations are given in μM. Each curve indicates an independent experiment.

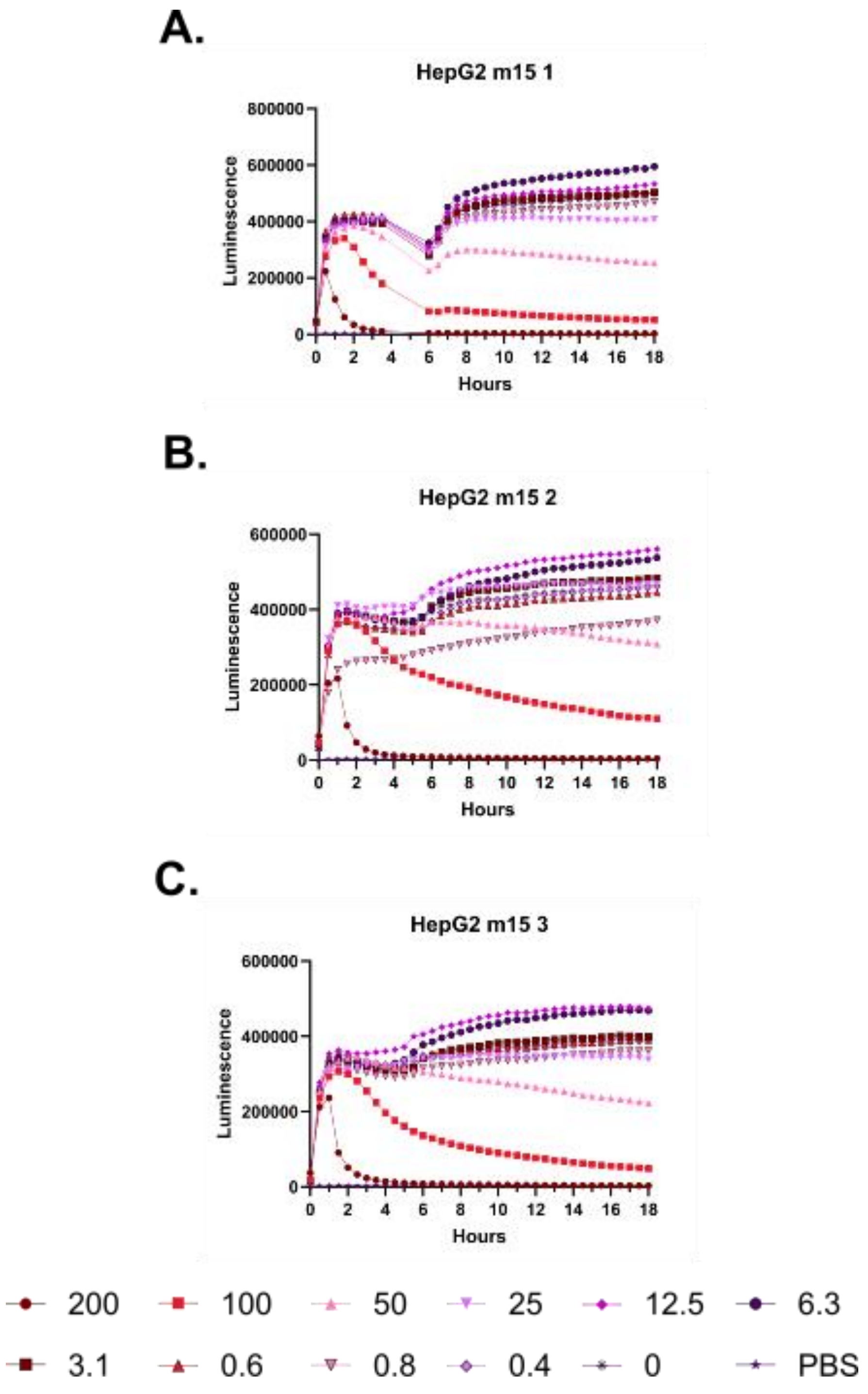
A.
HepG2 m15 1
Luminescence
800000
600000
400000
200000
0
0 2 4 6 8 10 12 14 16 18
Hours
B.
HepG2 m15 2
Luminescence
600000
400000
200000
0
0 2 4 6 8 10 12 14 16 18
Hours
C.
HepG2 m15 3
Luminescence
600000
400000
200000
0
0 2 4 6 8 10 12 14 16 18
Hours
200
100
50
25
12.5
6.3
3.1
0.6
0.8
0.4
0
PBS

**Supplementary Figure 82 – Kinetic luminescence toxicity assay for HepG2 treated with m15. A-C.** Luminescence was monitored over 18 hours at 37 °C with 5% $CO_2$ in a Small Humidity Cassette in the presence of the indicated peptide concentrations. Toxicity is indicated by curves that fall off of a steady upward trajectory. An instrument error in Experiment 1 resulted in no reads being taken during the gap in the curve. Peptide concentrations are given in μM. Each curve indicates an independent experiment.

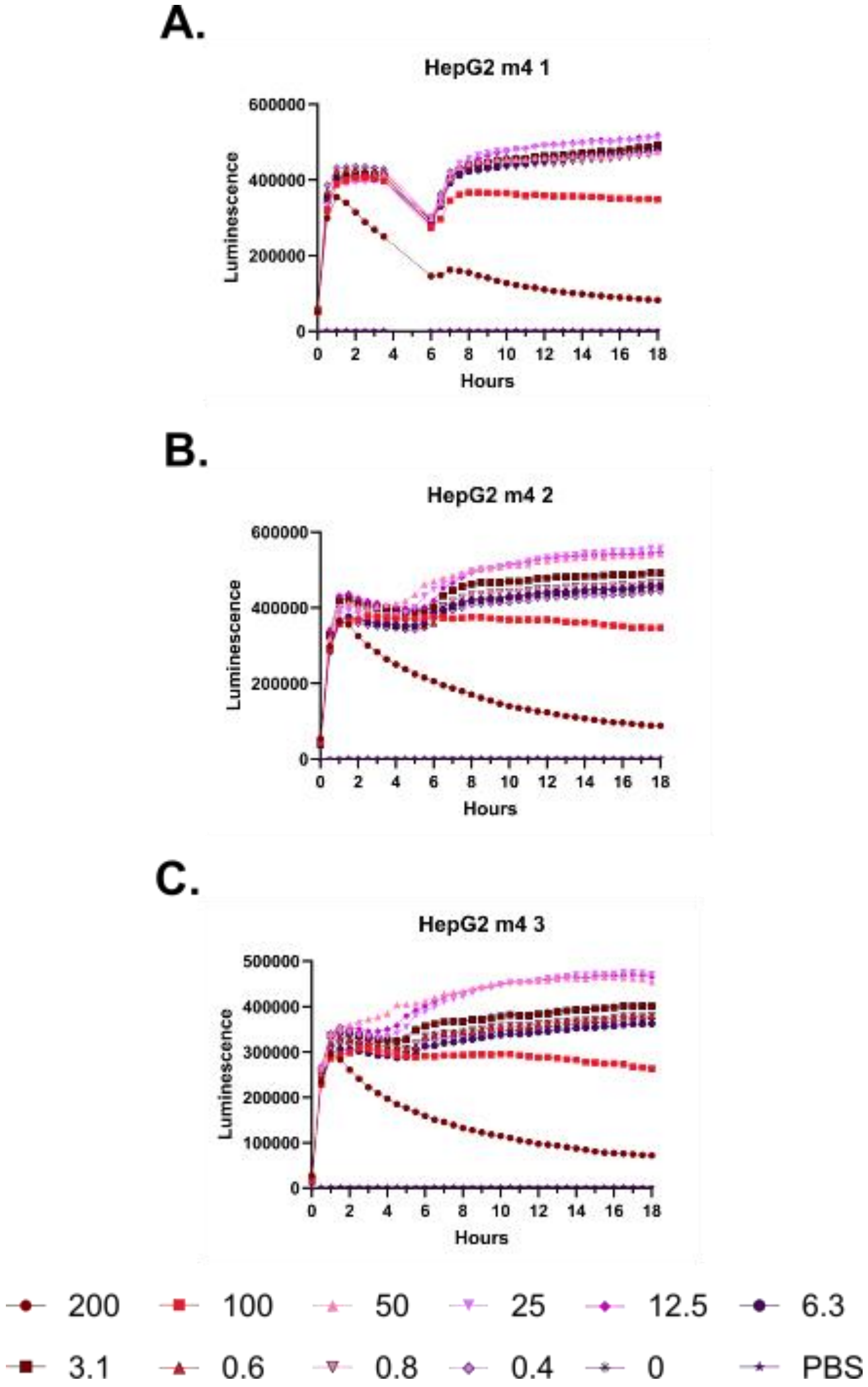
A.
HepG2 m4 1
Luminescence
Hours
B.
HepG2 m4 2
Luminescence
Hours
C.
HepG2 m4 3
Luminescence
Hours
200
100
50
25
12.5
6.3
3.1
0.6
0.8
0.4
0
PBS

**Supplementary Figure 82 – Kinetic luminescence toxicity assay for HepG2 treated with m4. A-C.** Luminescence was monitored over 18 hours at 37 °C with 5% $CO_2$ in a Small Humidity Cassette in the presence of the indicated peptide concentrations. Toxicity is indicated by curves that fall off of a steady upward trajectory. An instrument error in Experiment 1 resulted in no reads being taken during the gap in the curve. Peptide concentrations are given in μM. Each curve indicates an independent experiment.

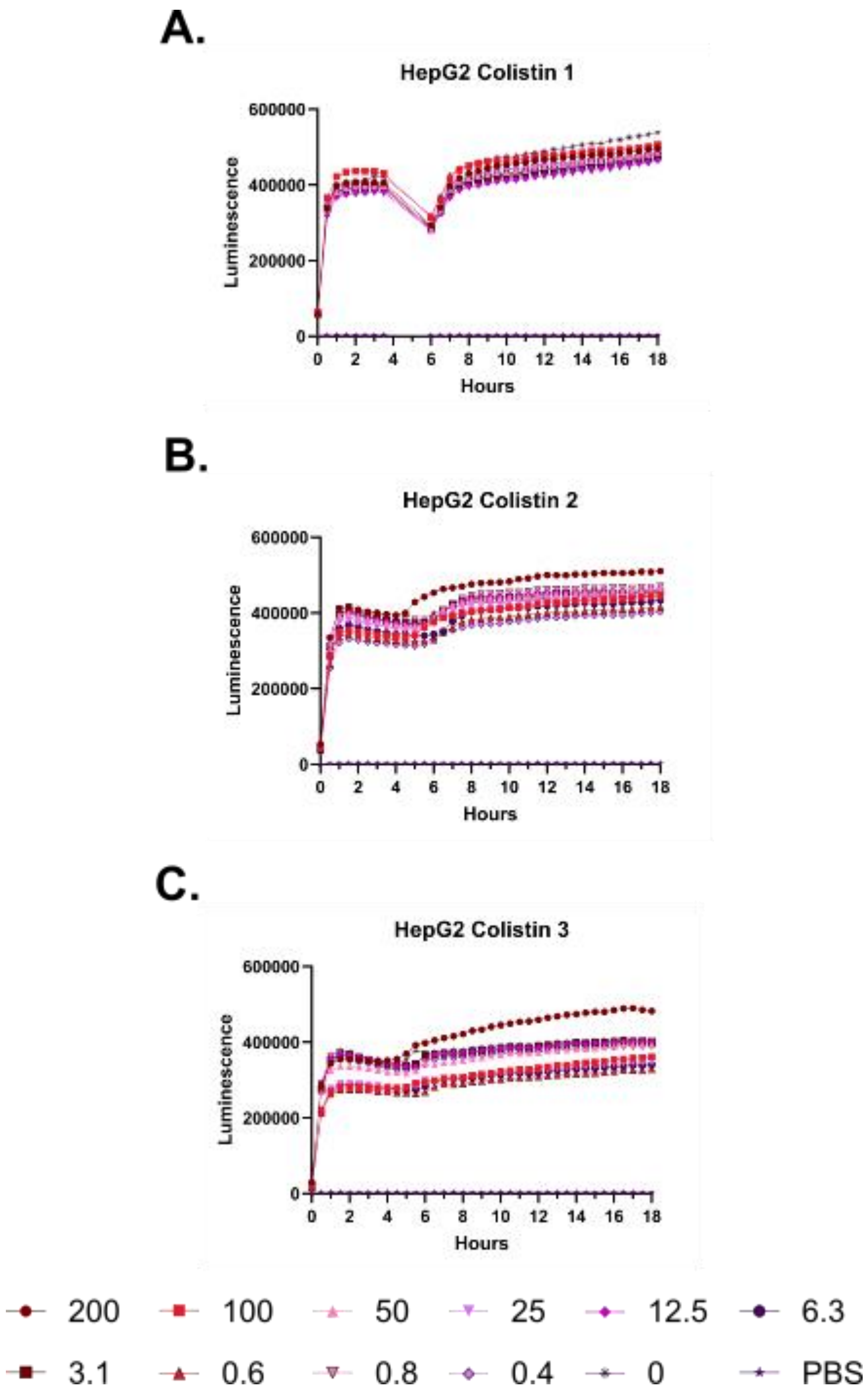

A.
HepG2 Colistin 1
Luminescence
Hours
B.
HepG2 Colistin 2
Luminescence
Hours
C.
HepG2 Colistin 3
Luminescence
Hours
200
100
50
25
12.5
6.3
3.1
0.6
0.8
0.4
0
PBS

**Supplementary Figure 84 – Kinetic luminescence toxicity assay for HepG2 treated with colistin. A-C.** Luminescence was monitored over 18 hours at 37 °C with 5% $CO_2$ in a Small Humidity Cassette in the presence of the indicated peptide concentrations. Toxicity is indicated by curves that fall off of a steady upward trajectory. An instrument error in Experiment 1 resulted in no reads being taken during the gap in the curve. Peptide concentrations are given in μM. Each curve indicates an independent experiment.

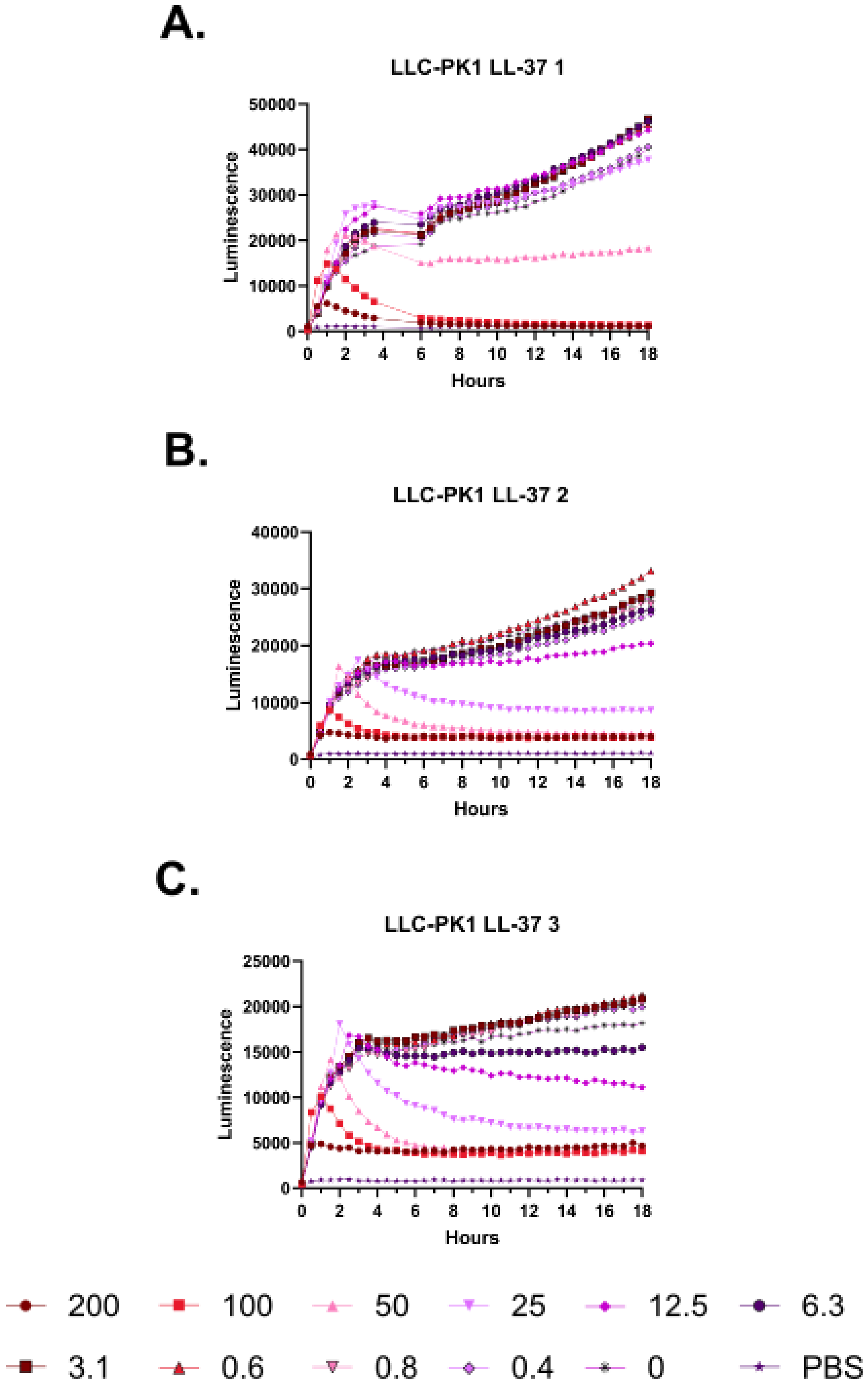
A.
LLC-PK1 LL-37 1
Luminescence
50000
40000
30000
20000
10000
0
0
2
4
6
8
10
12
14
16
18
Hours
B.
LLC-PK1 LL-37 2
Luminescence
40000
30000
20000
10000
0
0
2
4
6
8
10
12
14
16
18
Hours
C.
LLC-PK1 LL-37 3
Luminescence
25000
20000
15000
10000
5000
0
0
2
4
6
8
10
12
14
16
18
Hours
200
100
50
25
12.5
6.3
3.1
0.6
0.8
0.4
0
PBS

**Supplementary Figure 85 – Kinetic luminescence toxicity assay for LLC-PK1 treated with LL-37. A-C.** Luminescence was monitored over 18 hours at 37 °C with 5% $CO_2$ in a Small Humidity Cassette in the presence of the indicated peptide concentrations. Toxicity is indicated by curves that fall off of a steady upward trajectory. An instrument error in Experiment 1 resulted in no reads being taken during the gap in the curve. Peptide concentrations are given in μM. Each curve indicates an independent experiment.

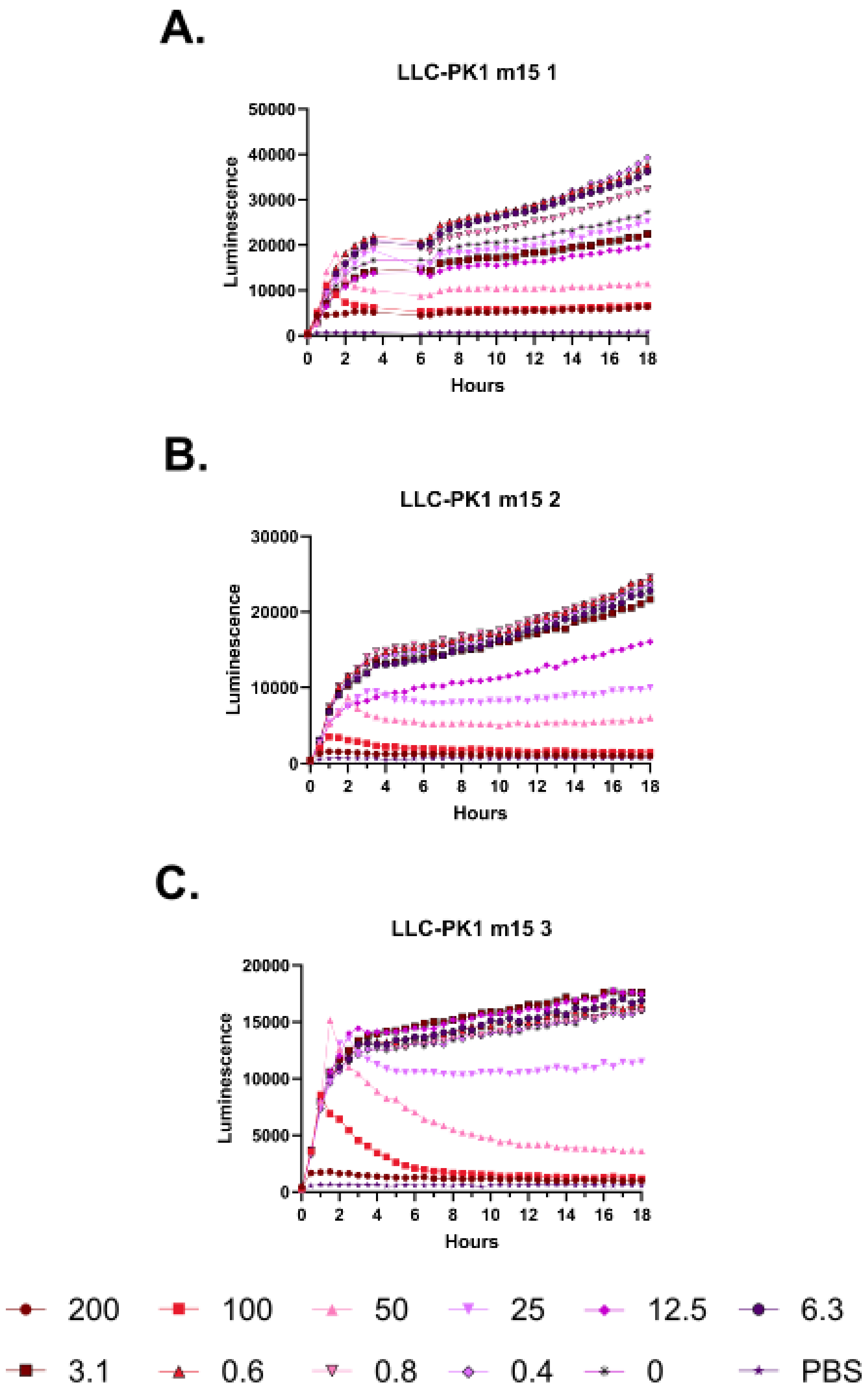
A.
LLC-PK1 m15 1
Luminescence
Hours
B.
LLC-PK1 m15 2
Luminescence
Hours
C.
LLC-PK1 m15 3
Luminescence
Hours
200
100
50
25
12.5
6.3
3.1
0.6
0.8
0.4
0
PBS

**Supplementary Figure 86 – Kinetic luminescence toxicity assay for LLC-PK1 treated with m15. A-C.** Luminescence was monitored over 18 hours at 37 °C with 5% $CO_2$ in a Small Humidity Cassette in the presence of the indicated peptide concentrations. Toxicity is indicated by curves that fall off of a steady upward trajectory. An instrument error in Experiment 1 resulted in no reads being taken during the gap in the curve. Peptide concentrations are given in µM. Each curve indicates an independent experiment.

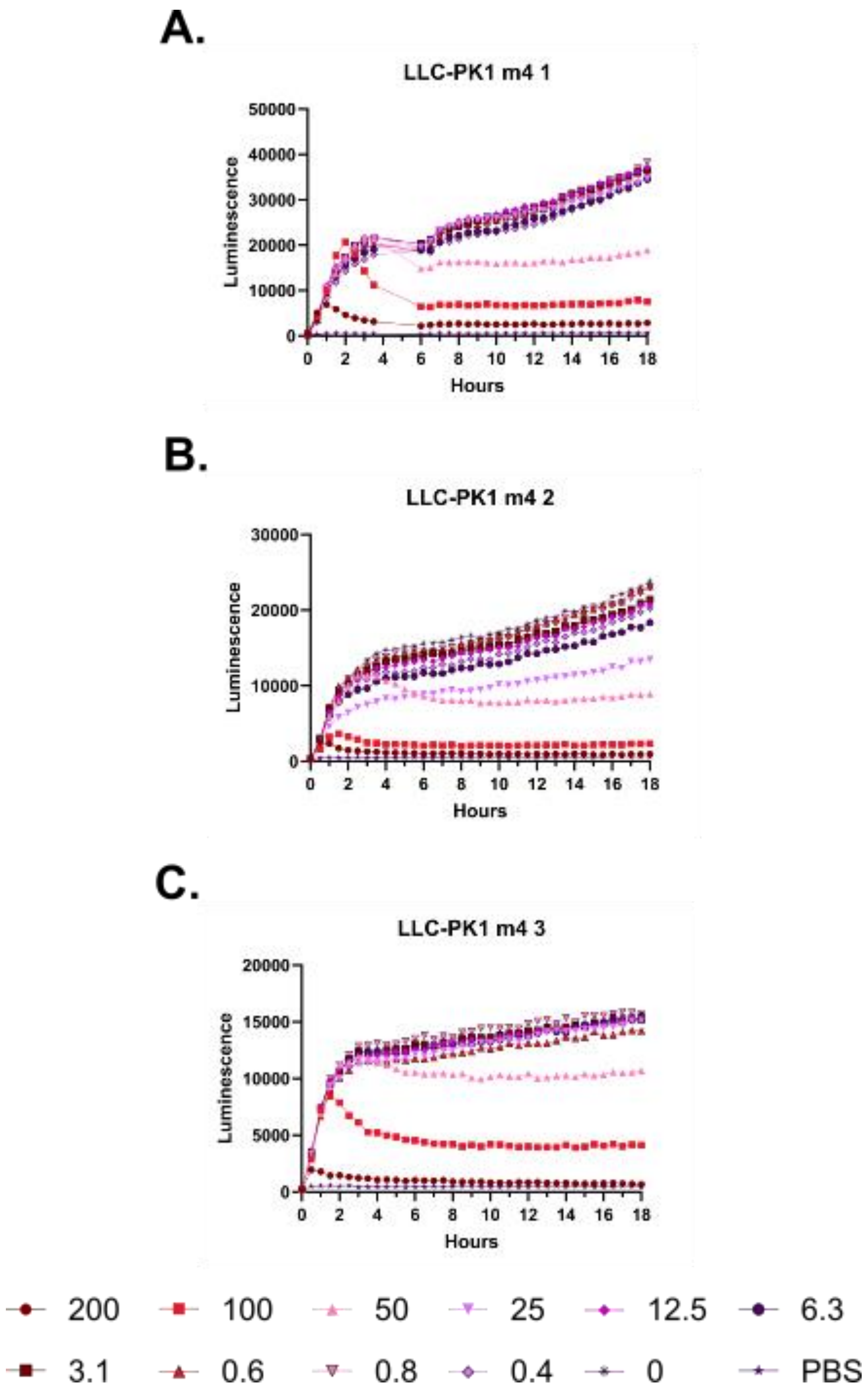
A.
LLC-PK1 m4 1
Luminescence
Hours
B.
LLC-PK1 m4 2
Luminescence
Hours
C.
LLC-PK1 m4 3
Luminescence
Hours
200
100
50
25
12.5
6.3
3.1
0.6
0.8
0.4
0
PBS

**Supplementary Figure 87 – Kinetic luminescence toxicity assay for LLC-PK1 treated with m4. A-C.** Luminescence was monitored over 18 hours at 37 °C with 5% $CO_2$ in a Small Humidity Cassette in the presence of the indicated peptide concentrations. Toxicity is indicated by curves that fall off of a steady upward trajectory. An instrument error in Experiment 1 resulted in no reads being taken during the gap in the curve. Peptide concentrations are given in μM. Each curve indicates an independent experiment.

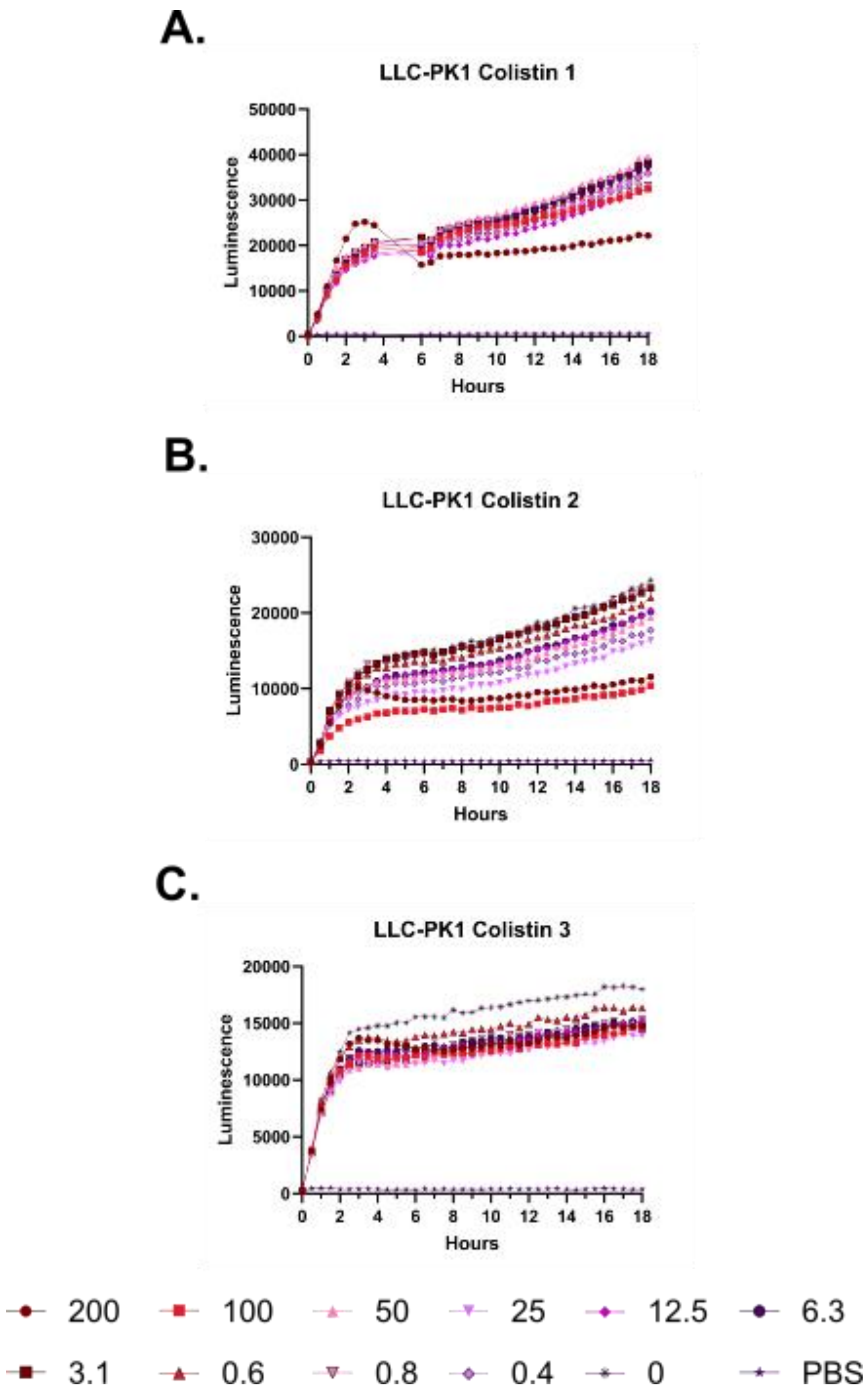
A.
LLC-PK1 Colistin 1
Luminescence
Hours
B.
LLC-PK1 Colistin 2
Luminescence
Hours
C.
LLC-PK1 Colistin 3
Luminescence
Hours
200
100
50
25
12.5
6.3
3.1
0.6
0.8
0.4
0
PBS

**Supplementary Figure 88 – Kinetic luminescence toxicity assay for LLC-PK1 treated with colistin. A-C.** Luminescence was monitored over 18 hours at 37 °C with 5% $CO_2$ in a Small Humidity Cassette in the presence of the indicated peptide concentrations. Toxicity is indicated by curves that fall off of a steady upward trajectory. An instrument error in Experiment 1 resulted in no reads being taken during the gap in the curve. Peptide concentrations are given in μM. Each curve indicates an independent experiment.

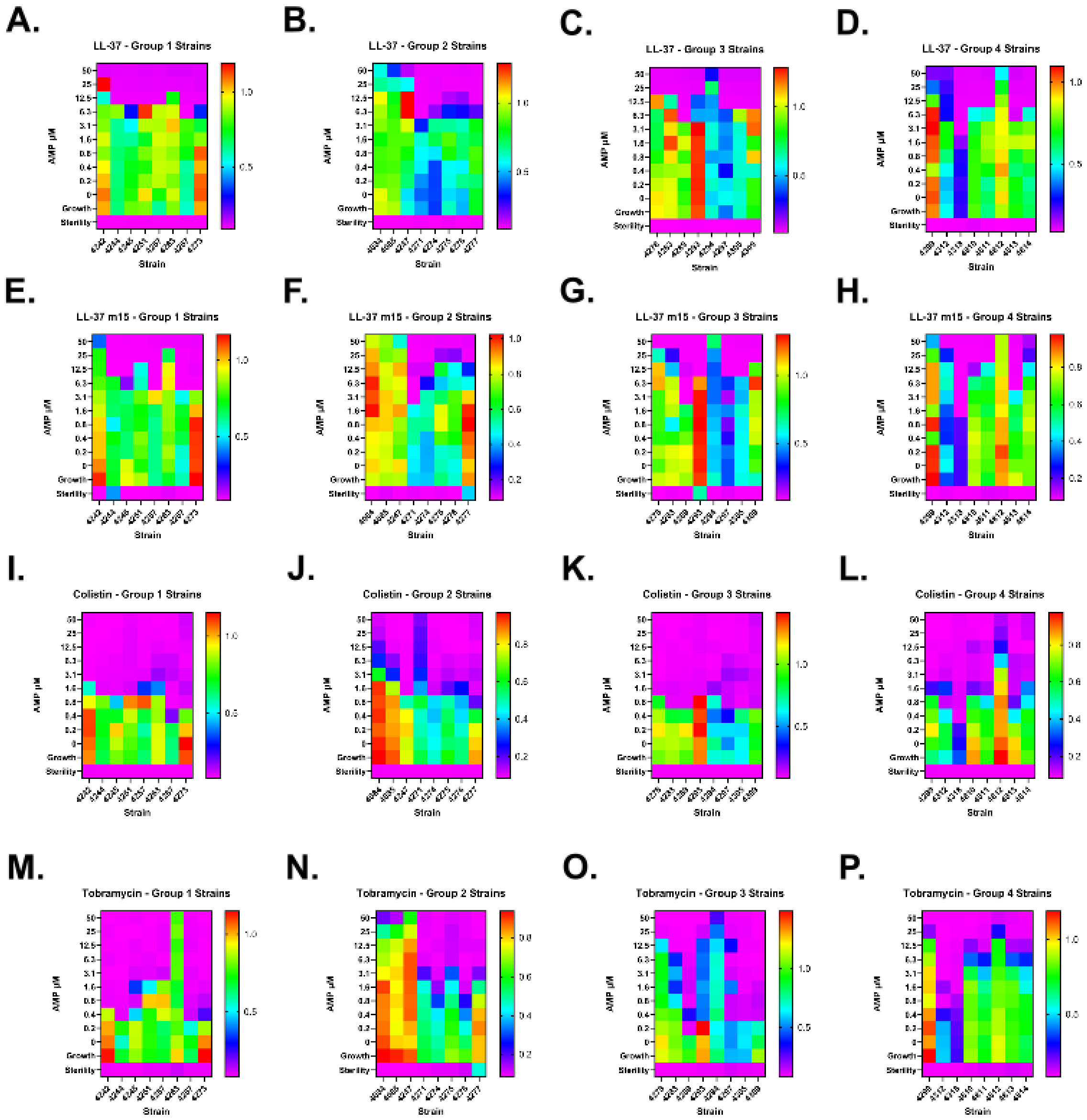
A.
LL-37 - Group 1 Strains
B.
LL-37 - Group 2 Strains
C.
LL-37 - Group 3 Strains
D.
LL-37 - Group 4 Strains
E.
LL-37 m15 - Group 1 Strains
F.
LL-37 m15 - Group 2 Strains
G.
LL-37 m15 - Group 3 Strains
H.
LL-37 m15 - Group 4 Strains
I.
Colistin - Group 1 Strains
J.
Colistin - Group 2 Strains
K.
Colistin - Group 3 Strains
L.
Colistin - Group 4 Strains
M.
Tobramycin - Group 1 Strains
N.
Tobramycin - Group 2 Strains
O.
Tobramycin - Group 3 Strains
P.
Tobramycin - Group 4 Strains
AMP µM
Strain
Growth
Sterility

**Supplementary Figure 89 – Endpoint antimicrobial susceptibility testing for LL-37, m15, colistin, and tobramycin against clinical strains of *Pseudomonas*.** Clinical strains of *Pseudomonas* from patients with cystic fibrosis as maintained in a Massachusetts General Hospital Cystic Fibrosis Center collection were subjected to antimicrobial susceptibility testing with each of the above antimicrobials. Strains were divided into groups of eight each as shown, with results showing the mean of two independent experiments. Cool colors indicate no / low bacterial growth, while warm colors indicate bacterial growth. **A-D** = LL-37 with each strain; **E-H** = m15 with each strain; **I-L** = colistin with each strain; **M-P** = tobramycin with each strain.

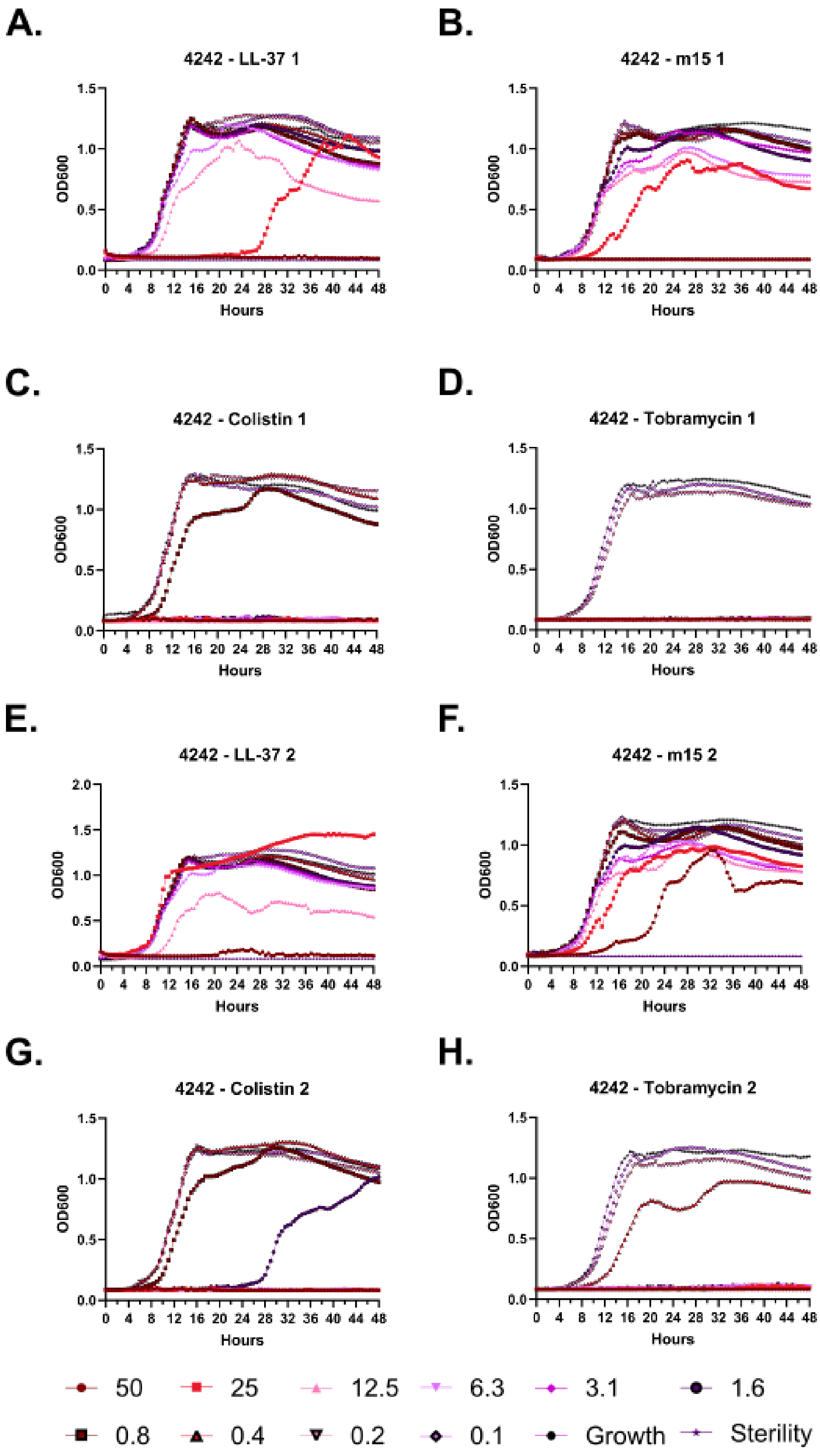
A.
4242 - LL-37 1
B.
4242 - m15 1
C.
4242 - Colistin 1
D.
4242 - Tobramycin 1
E.
4242 - LL-37 2
F.
4242 - m15 2
G.
4242 - Colistin 2
H.
4242 - Tobramycin 2
OD600
Hours
50
25
12.5
6.3
3.1
1.6
0.8
0.4
0.2
0.1
Growth
Sterility

**Supplementary Figure 90 – Kinetic antimicrobial susceptibility testing for *Pseudomonas* strain 4242. A-D.** Experiment 1. **E-H.** Experiment 2. Bacterial growth was monitored by OD600 over 48 hours at 37 °C in the presence of the indicated peptide concentrations. Peptide concentrations are given in μM.

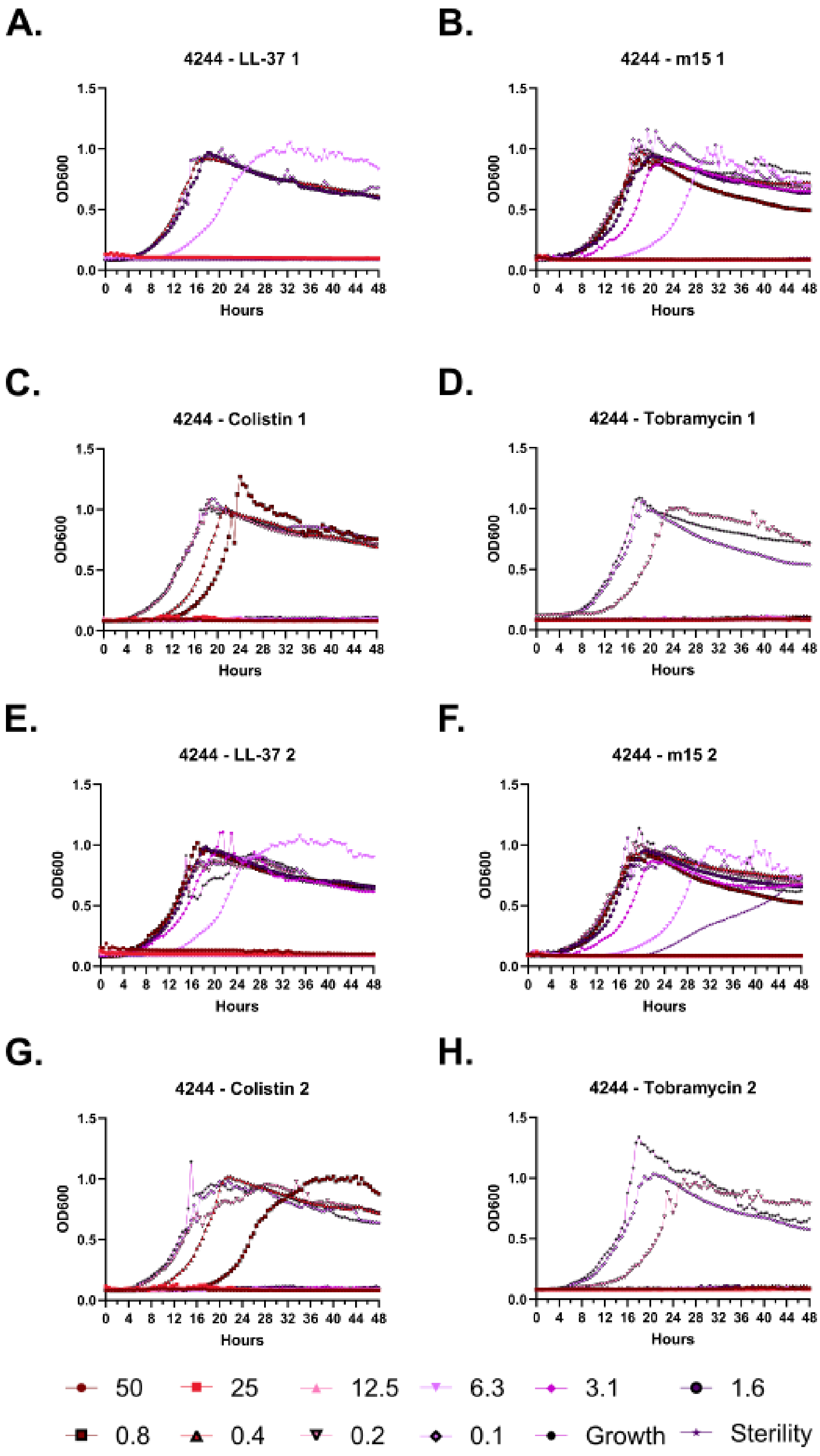
A.
4244 - LL-37 1
B.
4244 - m15 1
C.
4244 - Colistin 1
D.
4244 - Tobramycin 1
E.
4244 - LL-37 2
F.
4244 - m15 2
G.
4244 - Colistin 2
H.
4244 - Tobramycin 2
OD600
Hours
0.0
0.5
1.0
1.5
0 4 8 12 16 20 24 28 32 36 40 44 48
50
25
12.5
6.3
3.1
1.6
0.8
0.4
0.2
0.1
Growth
Sterility

**Supplementary Figure 91 – Kinetic antimicrobial susceptibility testing for *Pseudomonas* strain 4244. A-D.** Experiment 1. **E-H.** Experiment 2. Bacterial growth was monitored by OD600 over 48 hours at 37 °C in the presence of the indicated peptide concentrations. Peptide concentrations are given in μM.

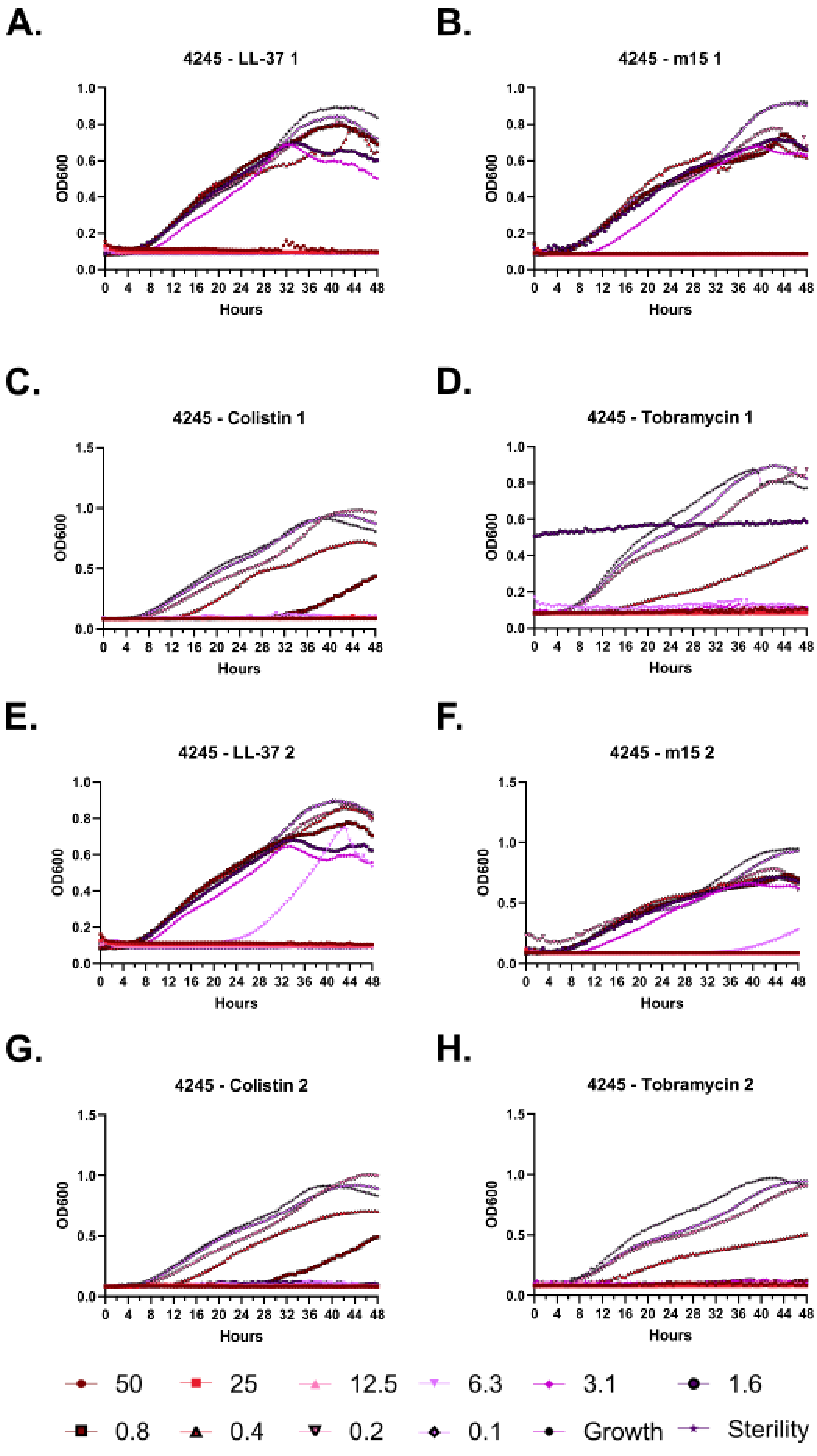

A.
4245 - LL-37 1
B.
4245 - m15 1
C.
4245 - Colistin 1
D.
4245 - Tobramycin 1
E.
4245 - LL-37 2
F.
4245 - m15 2
G.
4245 - Colistin 2
H.
4245 - Tobramycin 2
OD600
Hours
50
25
12.5
6.3
3.1
1.6
0.8
0.4
0.2
0.1
Growth
Sterility

**Supplementary Figure 92 – Kinetic antimicrobial susceptibility testing for *Pseudomonas* strain 4245. A-D.** Experiment 1. **E-H.** Experiment 2. Bacterial growth was monitored by OD600 over 48 hours at 37 °C in the presence of the indicated peptide concentrations. Peptide concentrations are given in μM.

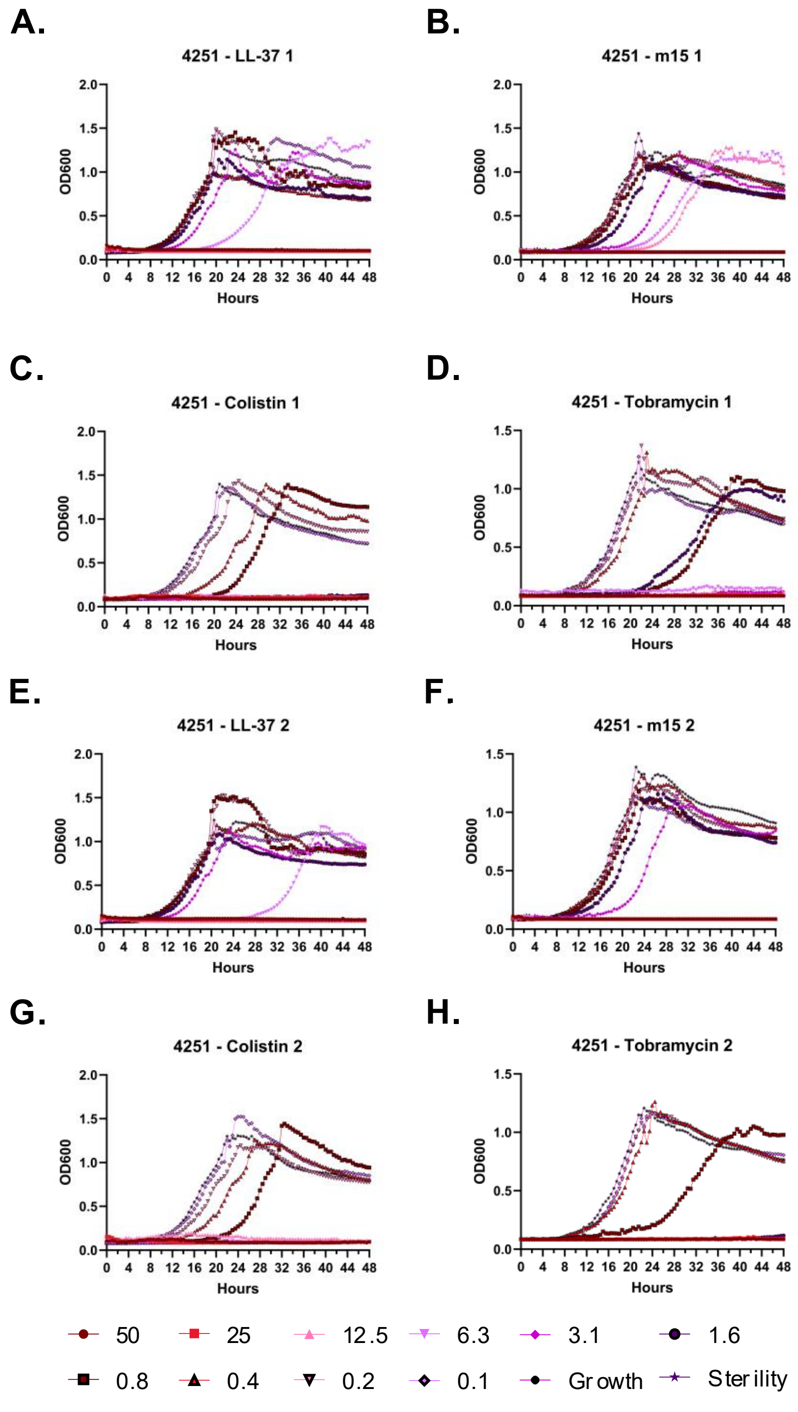
A.
4251 - LL-37 1
B.
4251 - m15 1
C.
4251 - Colistin 1
D.
4251 - Tobramycin 1
E.
4251 - LL-37 2
F.
4251 - m15 2
G.
4251 - Colistin 2
H.
4251 - Tobramycin 2
OD600
Hours
50
25
12.5
6.3
3.1
1.6
0.8
0.4
0.2
0.1
Growth
Sterility

**Supplementary Figure 93 – Kinetic antimicrobial susceptibility testing for *Pseudomonas* strain 4251. A-D.** Experiment 1. **E-H.** Experiment 2. Bacterial growth was monitored by OD600 over 48 hours at 37 °C in the presence of the indicated peptide concentrations. Peptide concentrations are given in μM.

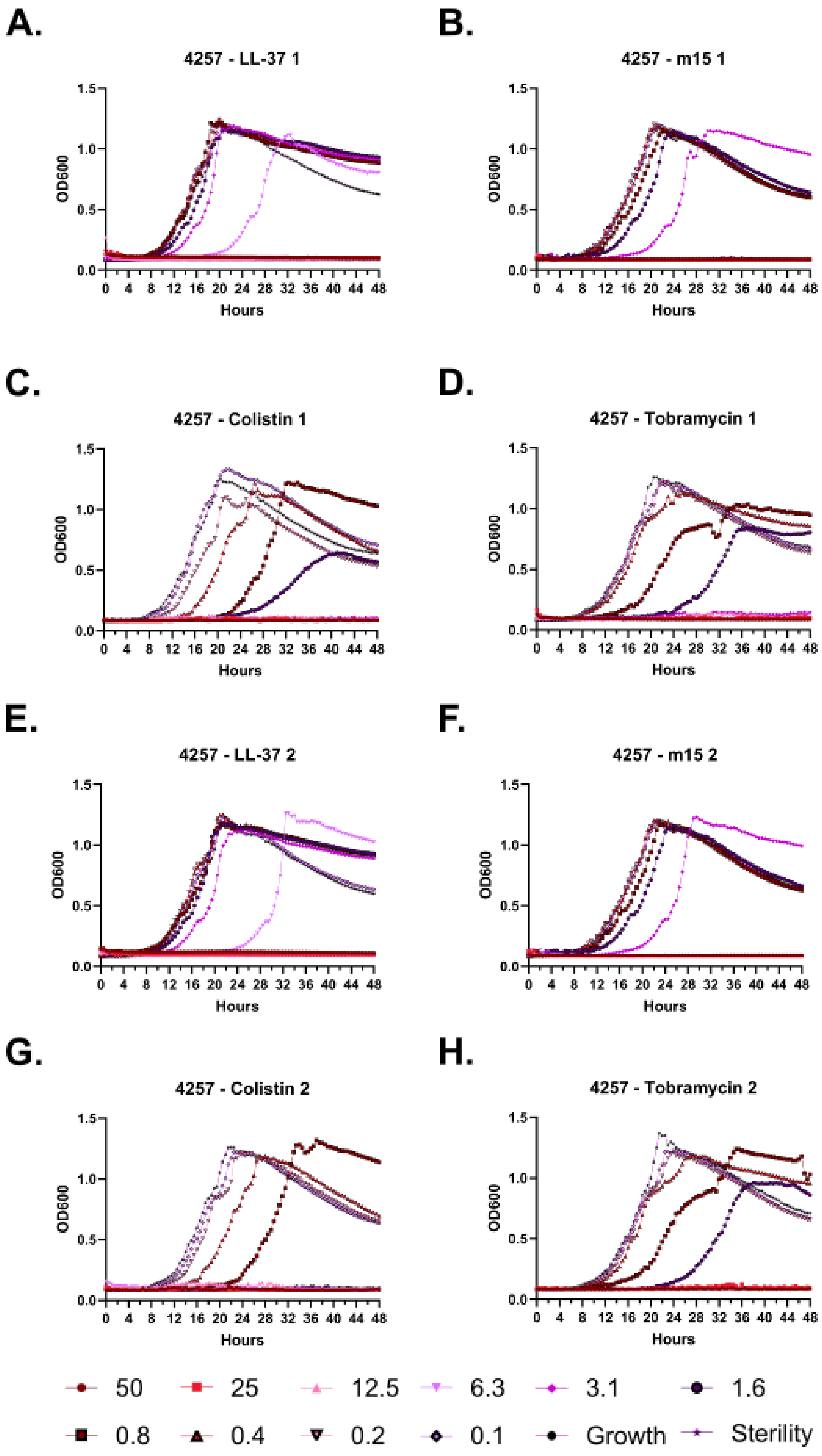
A.
4257 - LL-37 1
B.
4257 - m15 1
C.
4257 - Colistin 1
D.
4257 - Tobramycin 1
E.
4257 - LL-37 2
F.
4257 - m15 2
G.
4257 - Colistin 2
H.
4257 - Tobramycin 2
OD600
Hours
0.0
0.5
1.0
1.5
0 4 8 12 16 20 24 28 32 36 40 44 48
50
25
12.5
6.3
3.1
1.6
0.8
0.4
0.2
0.1
Growth
Sterility

**Supplementary Figure 94 – Kinetic antimicrobial susceptibility testing for *Pseudomonas* strain 4257. A-D.** Experiment 1. **E-H.** Experiment 2. Bacterial growth was monitored by OD600 over 48 hours at 37 °C in the presence of the indicated peptide concentrations. Peptide concentrations are given in μM.

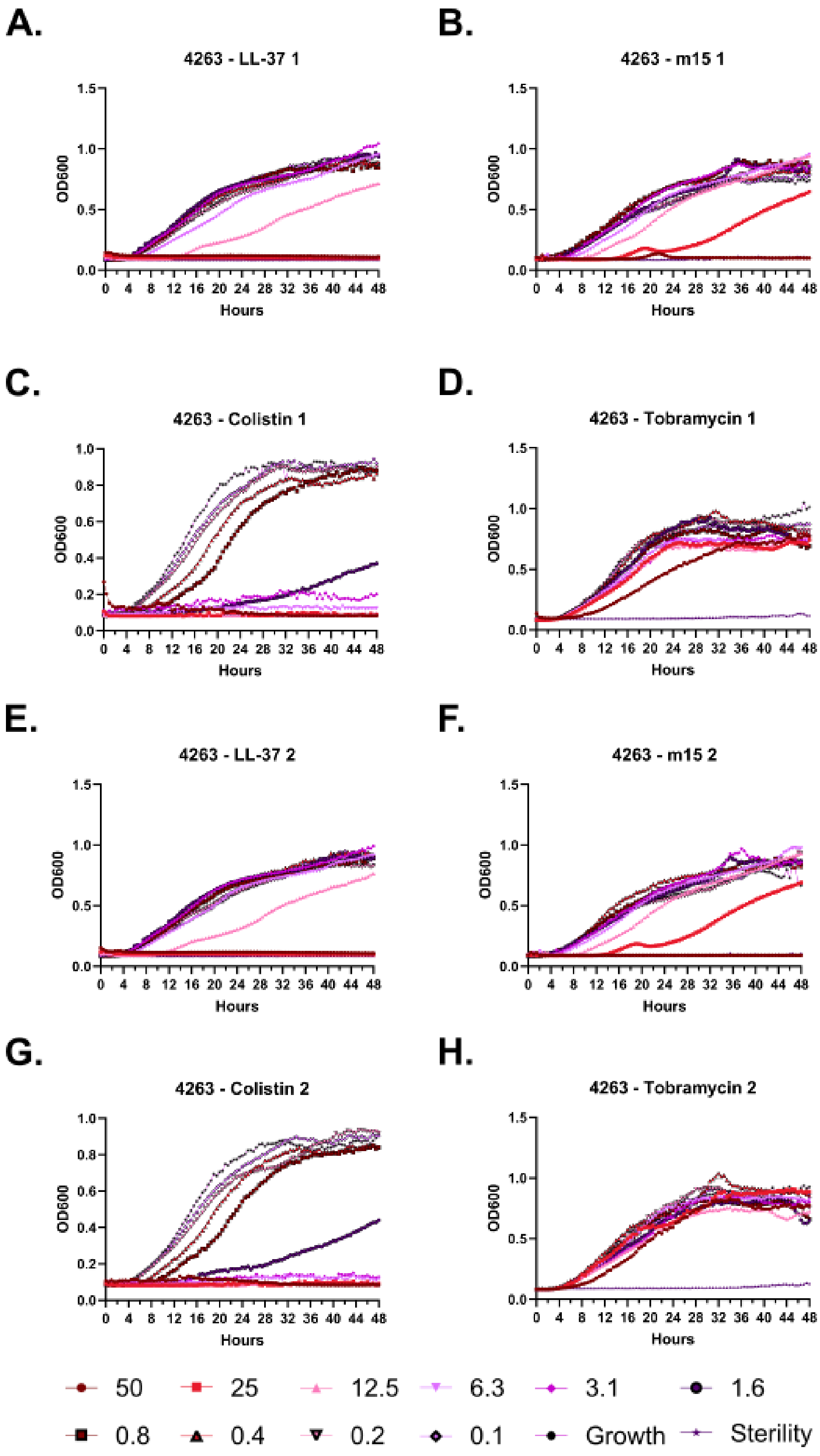

A.
4263 - LL-37 1
B.
4263 - m15 1
C.
4263 - Colistin 1
D.
4263 - Tobramycin 1
E.
4263 - LL-37 2
F.
4263 - m15 2
G.
4263 - Colistin 2
H.
4263 - Tobramycin 2
OD600
Hours
50
25
12.5
6.3
3.1
1.6
0.8
0.4
0.2
0.1
Growth
Sterility

**Supplementary Figure 95 – Kinetic antimicrobial susceptibility testing for *Pseudomonas* strain 4263. A-D.** Experiment 1. **E-H.** Experiment 2. Bacterial growth was monitored by OD600 over 48 hours at 37 °C in the presence of the indicated peptide concentrations. Peptide concentrations are given in μM.

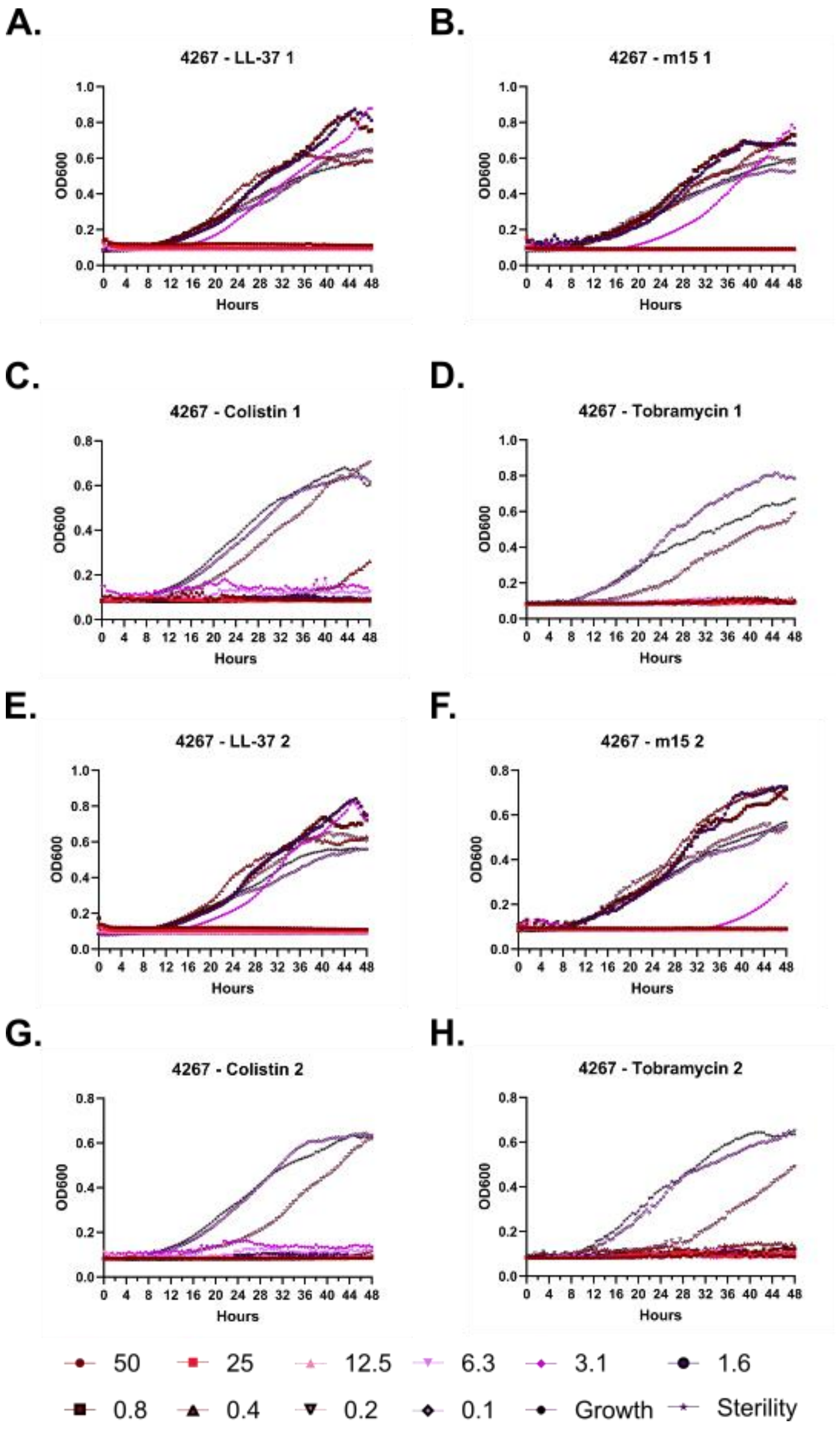
A.
4267 - LL-37 1
B.
4267 - m15 1
C.
4267 - Colistin 1
D.
4267 - Tobramycin 1
E.
4267 - LL-37 2
F.
4267 - m15 2
G.
4267 - Colistin 2
H.
4267 - Tobramycin 2
OD600
Hours
50
25
12.5
6.3
3.1
1.6
0.8
0.4
0.2
0.1
Growth
Sterility

**Supplementary Figure 96 – Kinetic antimicrobial susceptibility testing for *Pseudomonas* strain 4267. A-D.** Experiment 1. **E-H.** Experiment 2. Bacterial growth was monitored by OD600 over 48 hours at 37 °C in the presence of the indicated peptide concentrations. Peptide concentrations are given in μM.

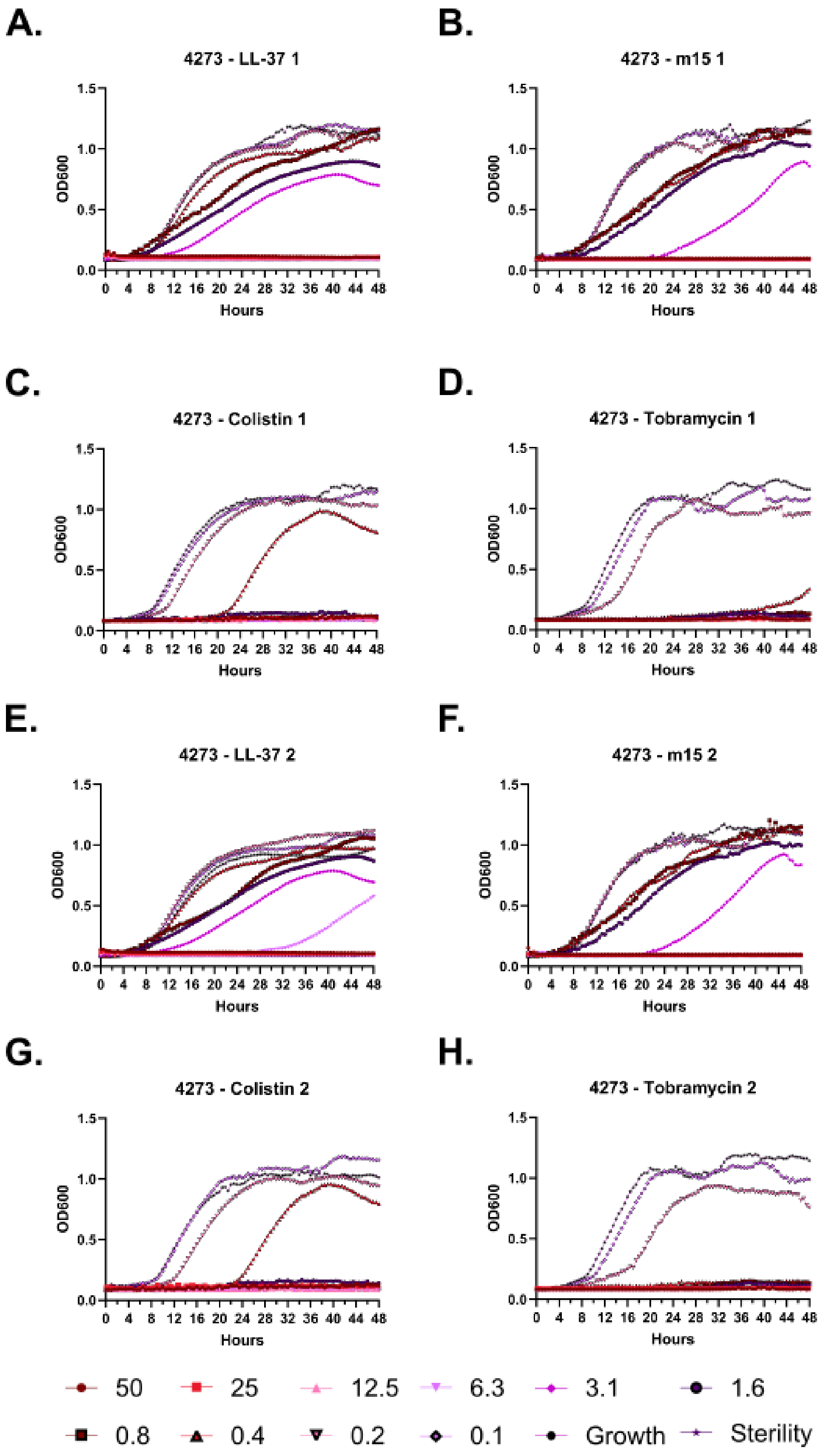

A.
4273 - LL-37 1
B.
4273 - m15 1
C.
4273 - Colistin 1
D.
4273 - Tobramycin 1
E.
4273 - LL-37 2
F.
4273 - m15 2
G.
4273 - Colistin 2
H.
4273 - Tobramycin 2
OD600
Hours
0.0
0.5
1.0
1.5
0 4 8 12 16 20 24 28 32 36 40 44 48
50
25
12.5
6.3
3.1
1.6
0.8
0.4
0.2
0.1
Growth
Sterility

**Supplementary Figure 97 – Kinetic antimicrobial susceptibility testing for *Pseudomonas* strain 4273. A-D.** Experiment 1. **E-H.** Experiment 2. Bacterial growth was monitored by OD600 over 48 hours at 37 °C in the presence of the indicated peptide concentrations. Peptide concentrations are given in μM.

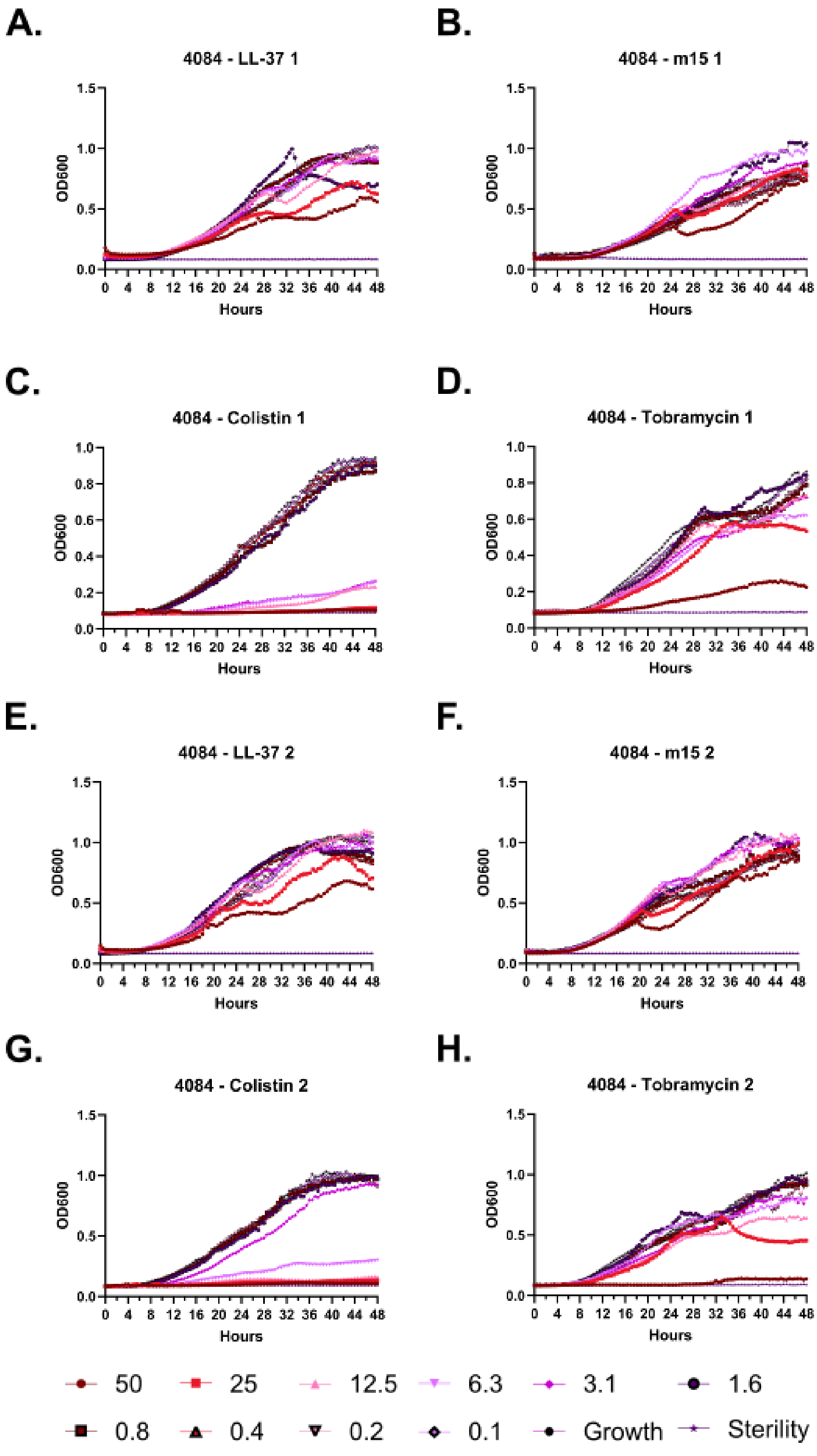
A.
4084 - LL-37 1
B.
4084 - m15 1
C.
4084 - Colistin 1
D.
4084 - Tobramycin 1
E.
4084 - LL-37 2
F.
4084 - m15 2
G.
4084 - Colistin 2
H.
4084 - Tobramycin 2
OD600
Hours
50
25
12.5
6.3
3.1
1.6
0.8
0.4
0.2
0.1
Growth
Sterility

**Supplementary Figure 98 – Kinetic antimicrobial susceptibility testing for *Pseudomonas* strain 4084. A-D.** Experiment 1. **E-H.** Experiment 2. Bacterial growth was monitored by OD600 over 48 hours at 37 °C in the presence of the indicated peptide concentrations. Peptide concentrations are given in μM.

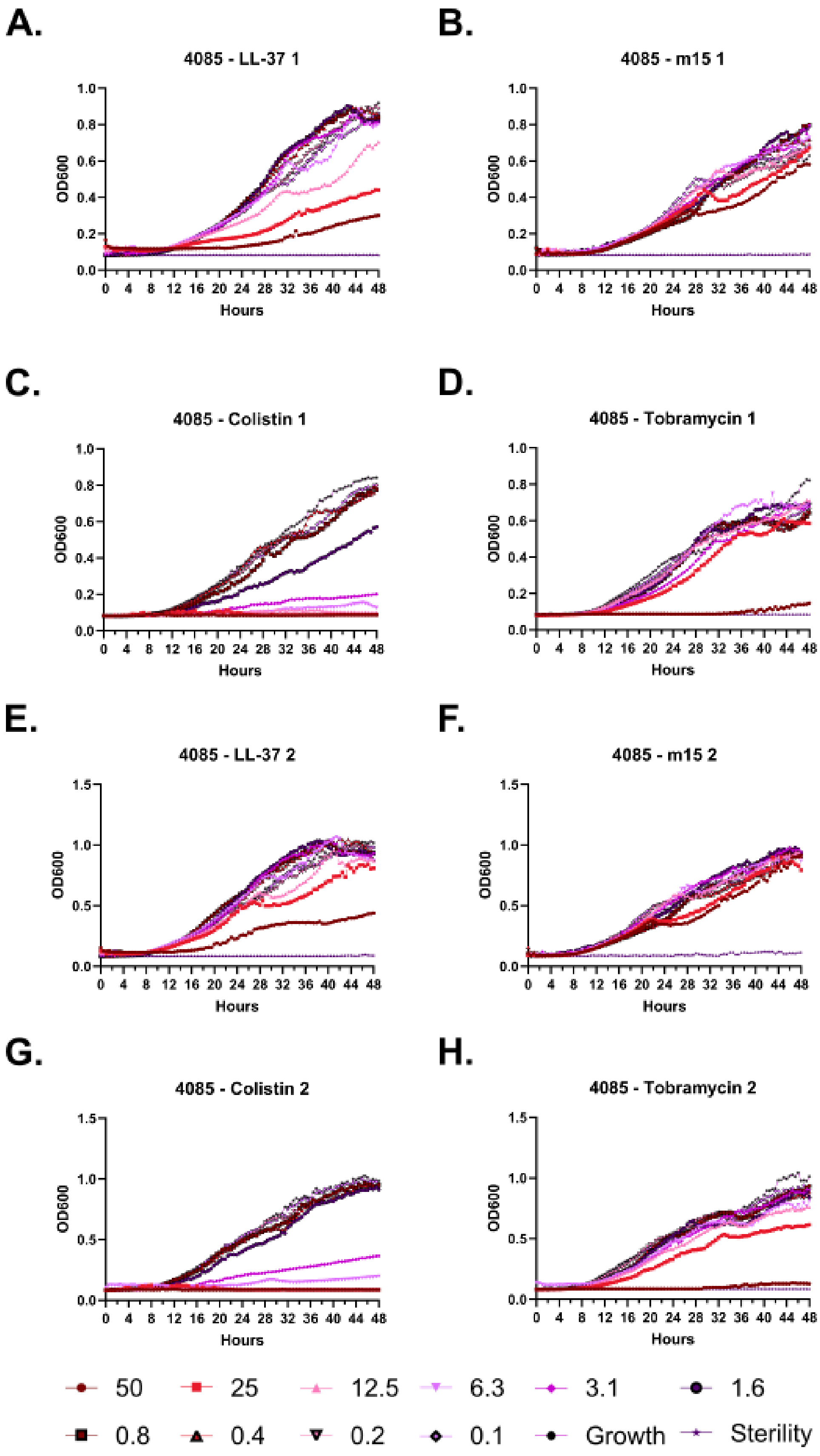

A.
4085 - LL-37 1
B.
4085 - m15 1
C.
4085 - Colistin 1
D.
4085 - Tobramycin 1
E.
4085 - LL-37 2
F.
4085 - m15 2
G.
4085 - Colistin 2
H.
4085 - Tobramycin 2
OD600
Hours
50
25
12.5
6.3
3.1
1.6
0.8
0.4
0.2
0.1
Growth
Sterility

**Supplementary Figure 99 – Kinetic antimicrobial susceptibility testing for *Pseudomonas* strain 4085. A-D.** Experiment 1. **E-H.** Experiment 2. Bacterial growth was monitored by OD600 over 48 hours at 37 °C in the presence of the indicated peptide concentrations. Peptide concentrations are given in μM.

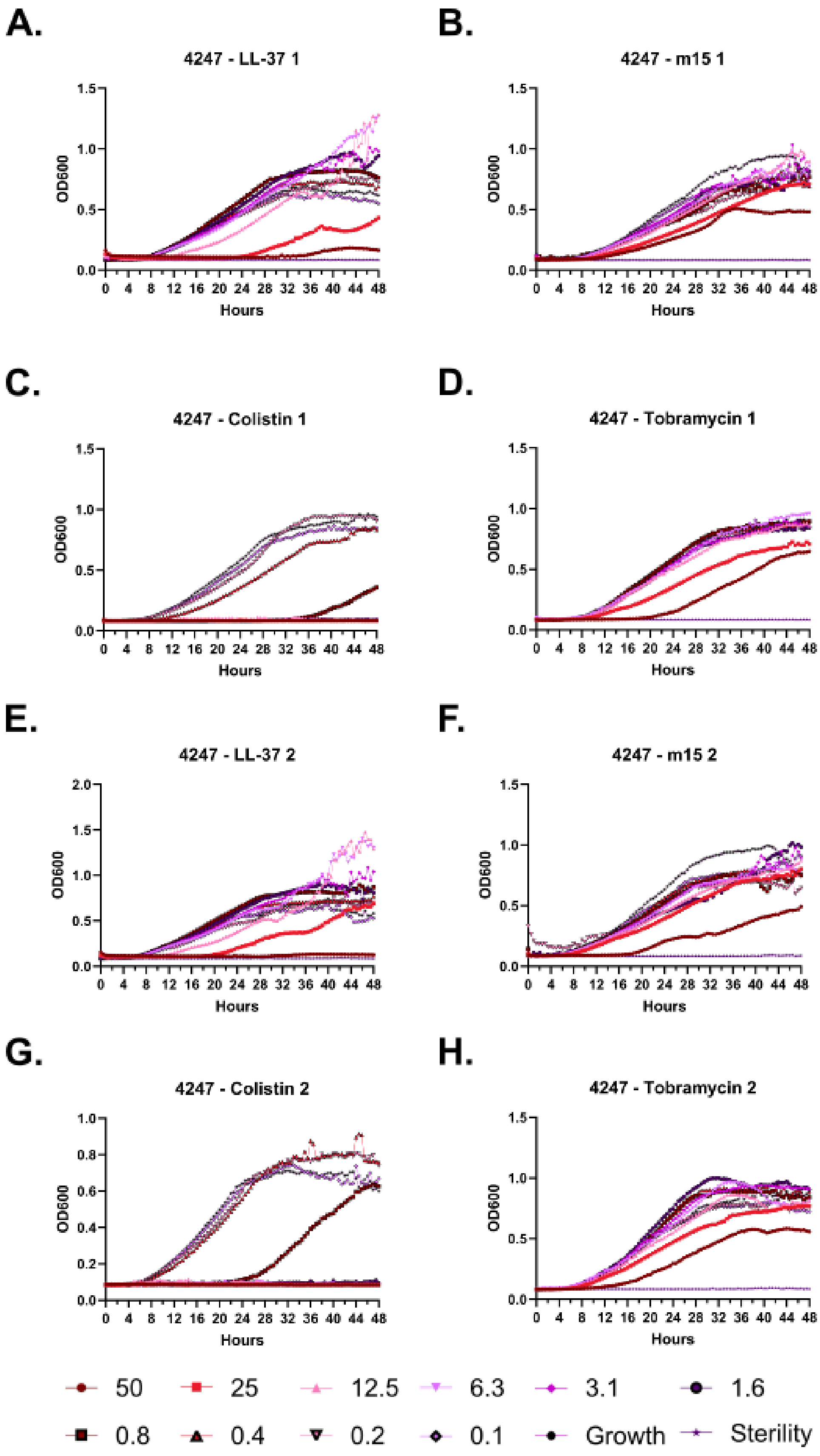
A.
4247 - LL-37 1
B.
4247 - m15 1
C.
4247 - Colistin 1
D.
4247 - Tobramycin 1
E.
4247 - LL-37 2
F.
4247 - m15 2
G.
4247 - Colistin 2
H.
4247 - Tobramycin 2
OD600
Hours
50
25
12.5
6.3
3.1
1.6
0.8
0.4
0.2
0.1
Growth
Sterility

**Supplementary Figure 100 – Kinetic antimicrobial susceptibility testing for *Pseudomonas* strain 4247. A-D.** Experiment 1. **E-H.** Experiment 2. Bacterial growth was monitored by OD600 over 48 hours at 37 °C in the presence of the indicated peptide concentrations. Peptide concentrations are given in μM.

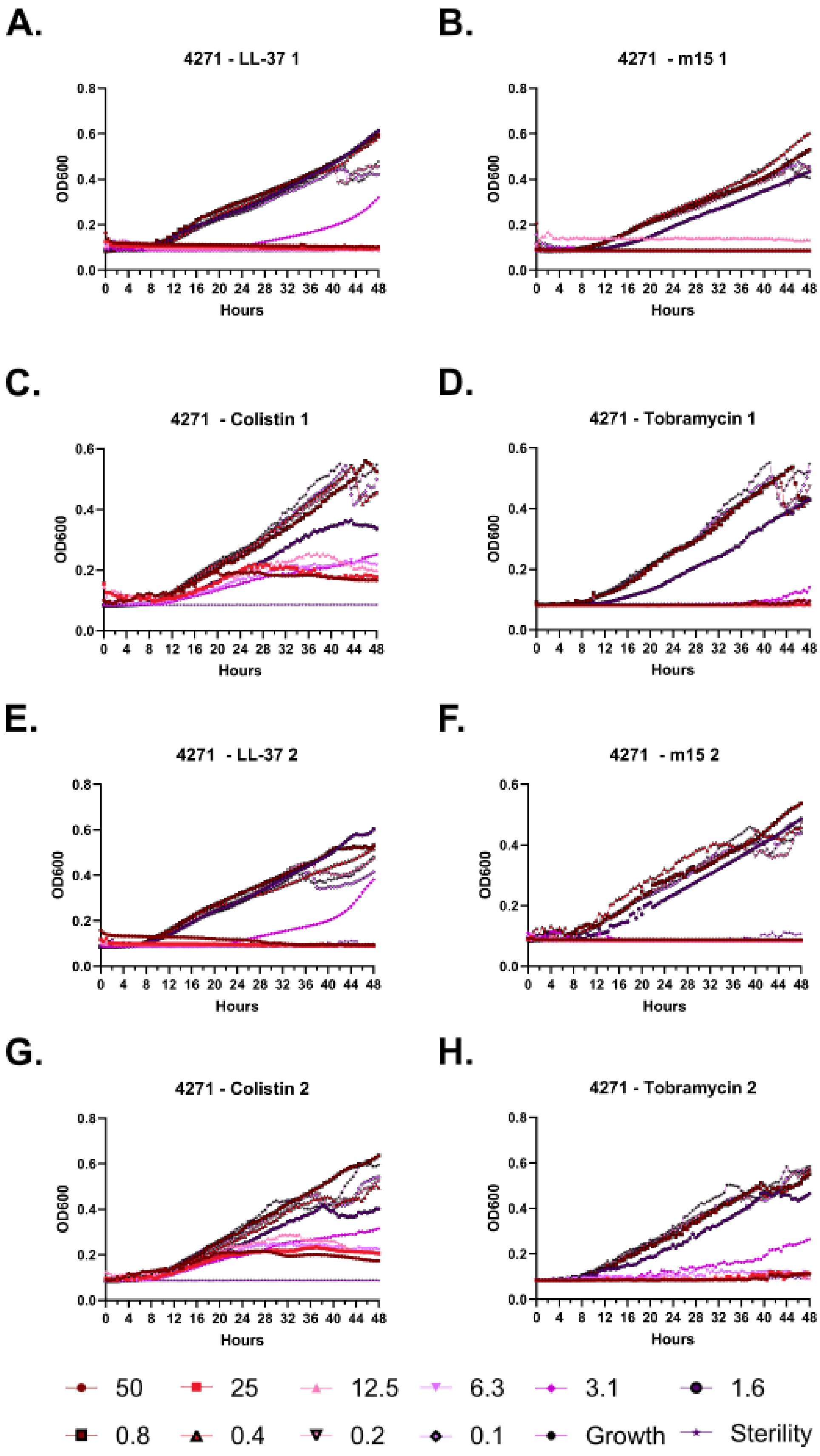

A.
4271 - LL-37 1
B.
4271 - m15 1
C.
4271 - Colistin 1
D.
4271 - Tobramycin 1
E.
4271 - LL-37 2
F.
4271 - m15 2
G.
4271 - Colistin 2
H.
4271 - Tobramycin 2
OD600
Hours
50
25
12.5
6.3
3.1
1.6
0.8
0.4
0.2
0.1
Growth
Sterility

**Supplementary Figure 101 – Kinetic antimicrobial susceptibility testing for *Pseudomonas* strain 4271. A-D.** Experiment 1. **E-H.** Experiment 2. Bacterial growth was monitored by OD600 over 48 hours at 37 °C in the presence of the indicated peptide concentrations. Peptide concentrations are given in μM.

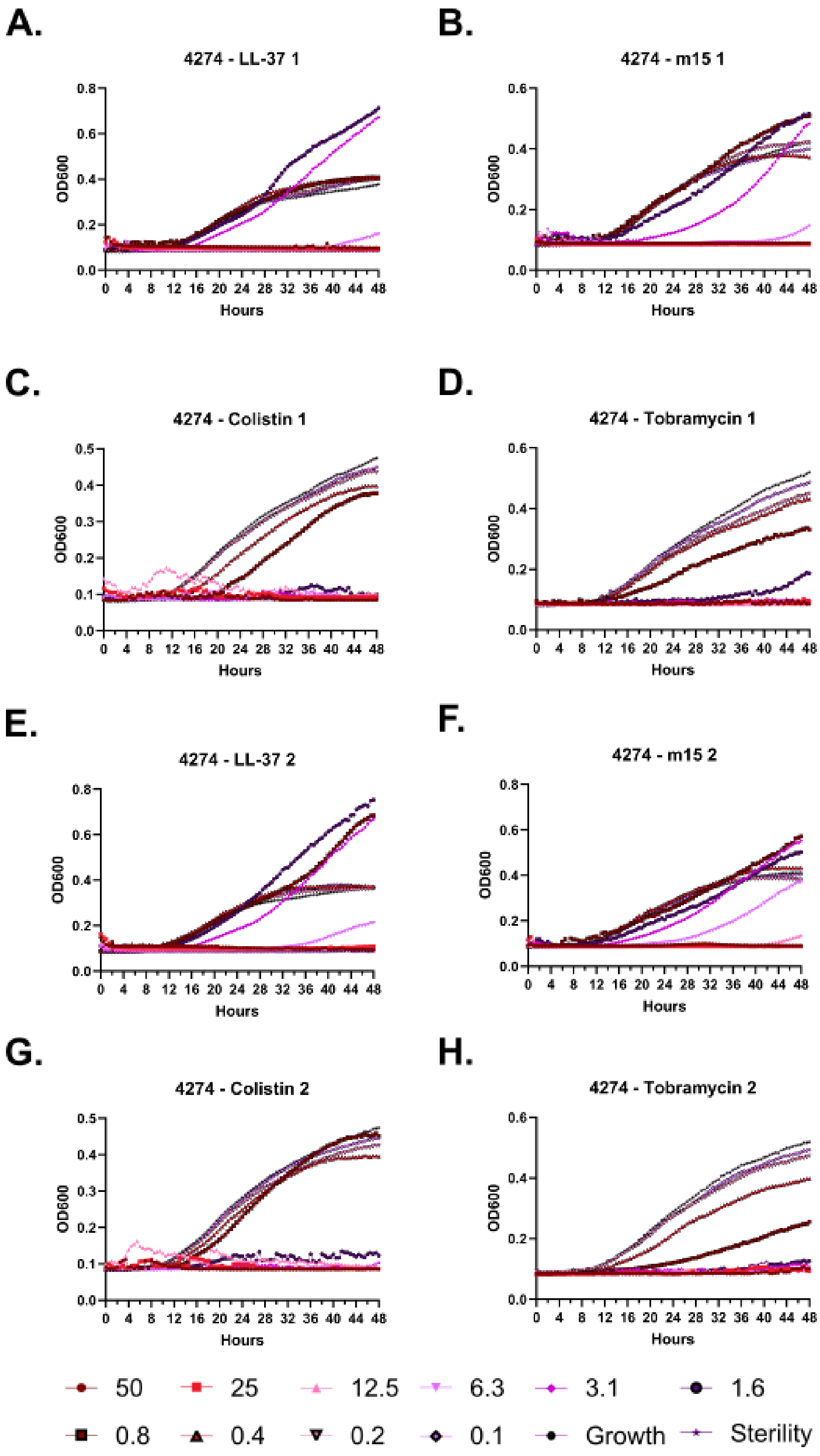

A.
4274 - LL-37 1
B.
4274 - m15 1
C.
4274 - Colistin 1
D.
4274 - Tobramycin 1
E.
4274 - LL-37 2
F.
4274 - m15 2
G.
4274 - Colistin 2
H.
4274 - Tobramycin 2
OD600
Hours
50
25
12.5
6.3
3.1
1.6
0.8
0.4
0.2
0.1
Growth
Sterility

**Supplementary Figure 102 – Kinetic antimicrobial susceptibility testing for *Pseudomonas* strain 4274. A-D.** Experiment 1. **E-H.** Experiment 2. Bacterial growth was monitored by OD600 over 48 hours at 37 °C in the presence of the indicated peptide concentrations. Peptide concentrations are given in μM.

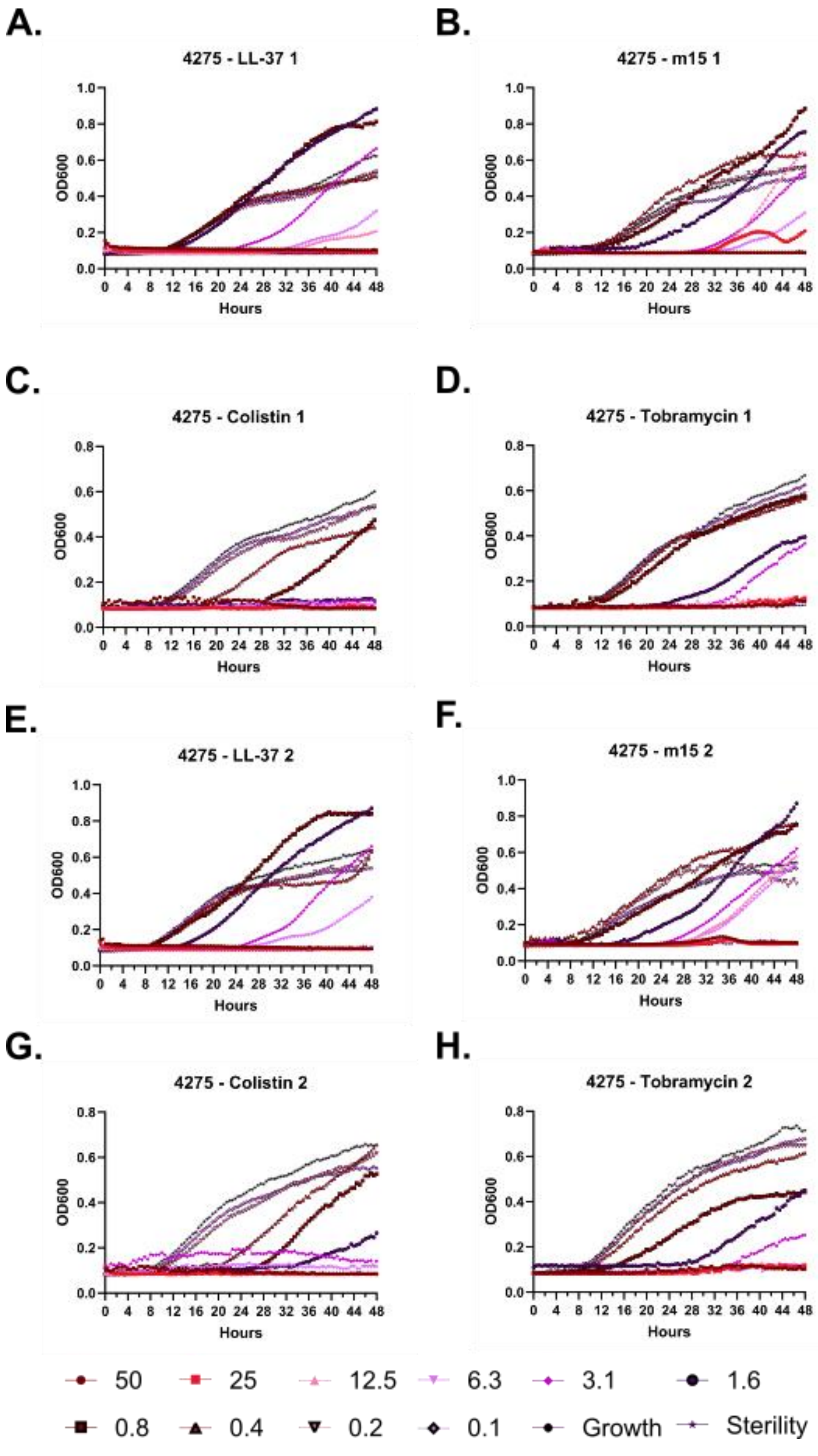
A.
4275 - LL-37 1
B.
4275 - m15 1
C.
4275 - Colistin 1
D.
4275 - Tobramycin 1
E.
4275 - LL-37 2
F.
4275 - m15 2
G.
4275 - Colistin 2
H.
4275 - Tobramycin 2
OD600
Hours
0 4 8 12 16 20 24 28 32 36 40 44 48
50
25
12.5
6.3
3.1
1.6
0.8
0.4
0.2
0.1
Growth
Sterility

**Supplementary Figure 103 – Kinetic antimicrobial susceptibility testing for *Pseudomonas* strain 4274. A-D.** Experiment 1. **E-H.** Experiment 2. Bacterial growth was monitored by OD600 over 48 hours at 37 °C in the presence of the indicated peptide concentrations. Peptide concentrations are given in μM.

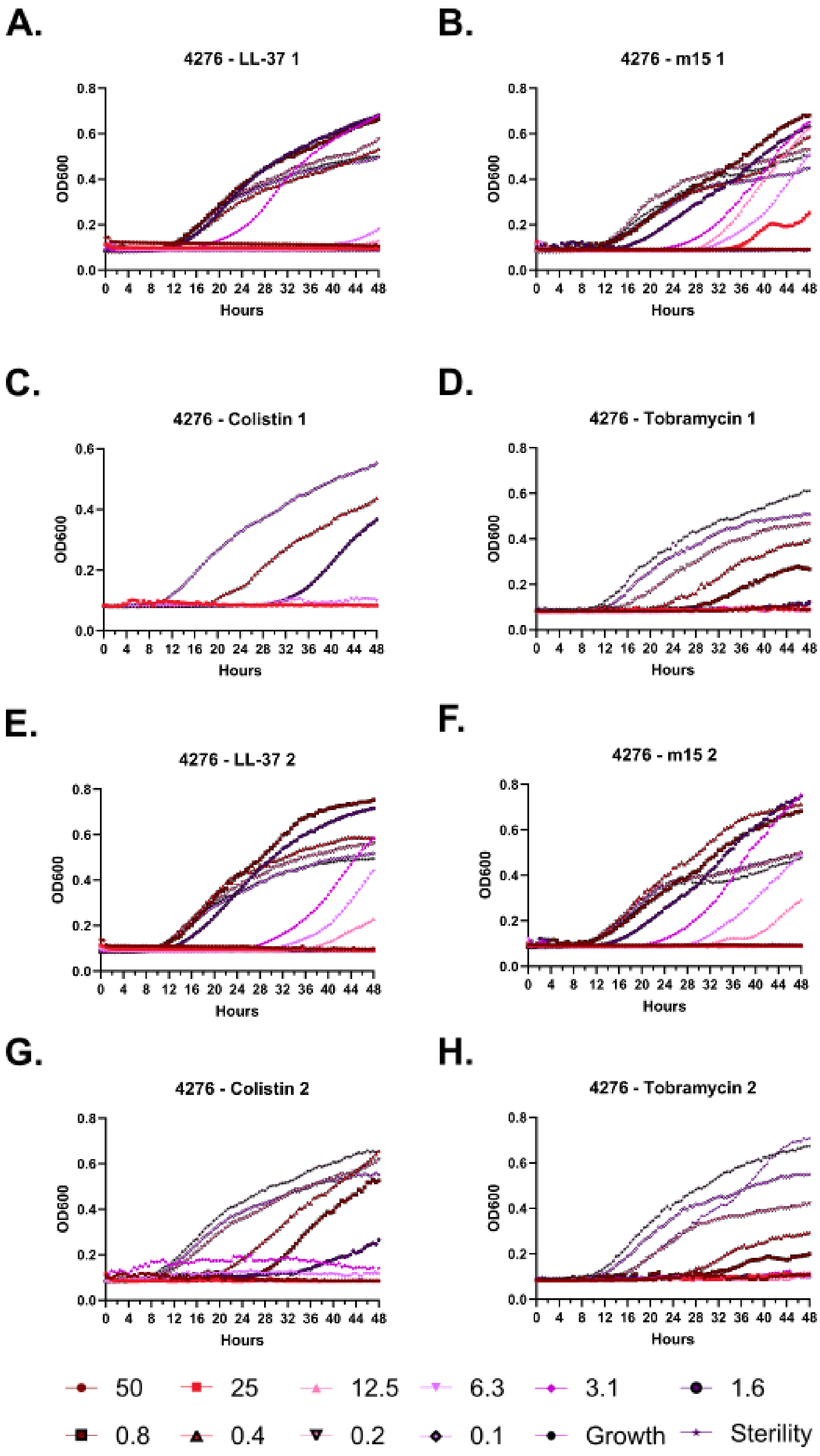

A.
4276 - LL-37 1
B.
4276 - m15 1
C.
4276 - Colistin 1
D.
4276 - Tobramycin 1
E.
4276 - LL-37 2
F.
4276 - m15 2
G.
4276 - Colistin 2
H.
4276 - Tobramycin 2
OD600
Hours
50
25
12.5
6.3
3.1
1.6
0.8
0.4
0.2
0.1
Growth
Sterility

**Supplementary Figure 104 – Kinetic antimicrobial susceptibility testing for *Pseudomonas* strain 4276. A-D.** Experiment 1. **E-H.** Experiment 2. Bacterial growth was monitored by OD600 over 48 hours at 37 °C in the presence of the indicated peptide concentrations. Peptide concentrations are given in μM.

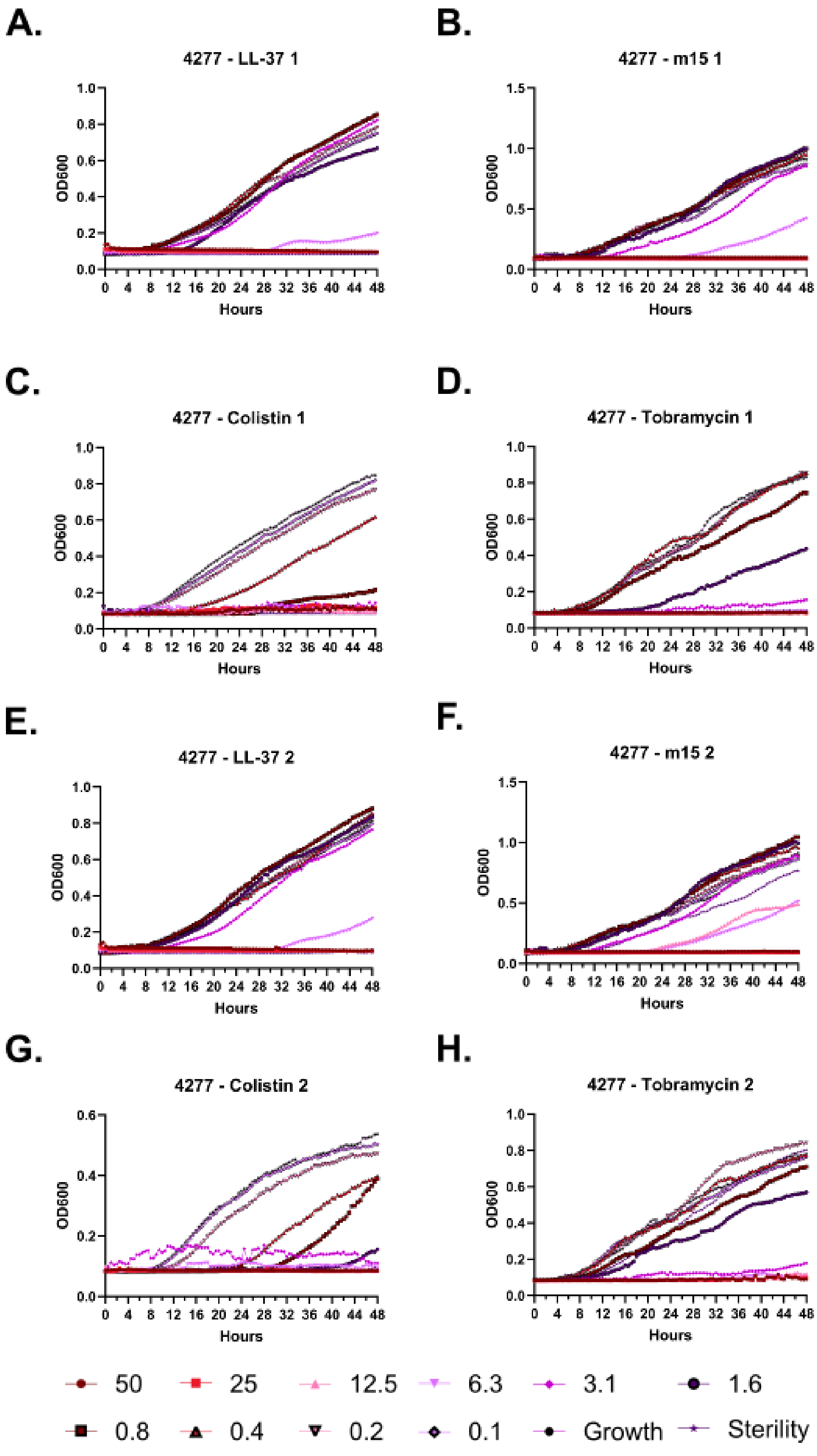
A.
4277 - LL-37 1
B.
4277 - m15 1
C.
4277 - Colistin 1
D.
4277 - Tobramycin 1
E.
4277 - LL-37 2
F.
4277 - m15 2
G.
4277 - Colistin 2
H.
4277 - Tobramycin 2
OD600
Hours
50
25
12.5
6.3
3.1
1.6
0.8
0.4
0.2
0.1
Growth
Sterility

**Supplementary Figure 105 – Kinetic antimicrobial susceptibility testing for *Pseudomonas* strain 4277. A-D.** Experiment 1. **E-H.** Experiment 2. Bacterial growth was monitored by OD600 over 48 hours at 37 °C in the presence of the indicated peptide concentrations. Peptide concentrations are given in μM.

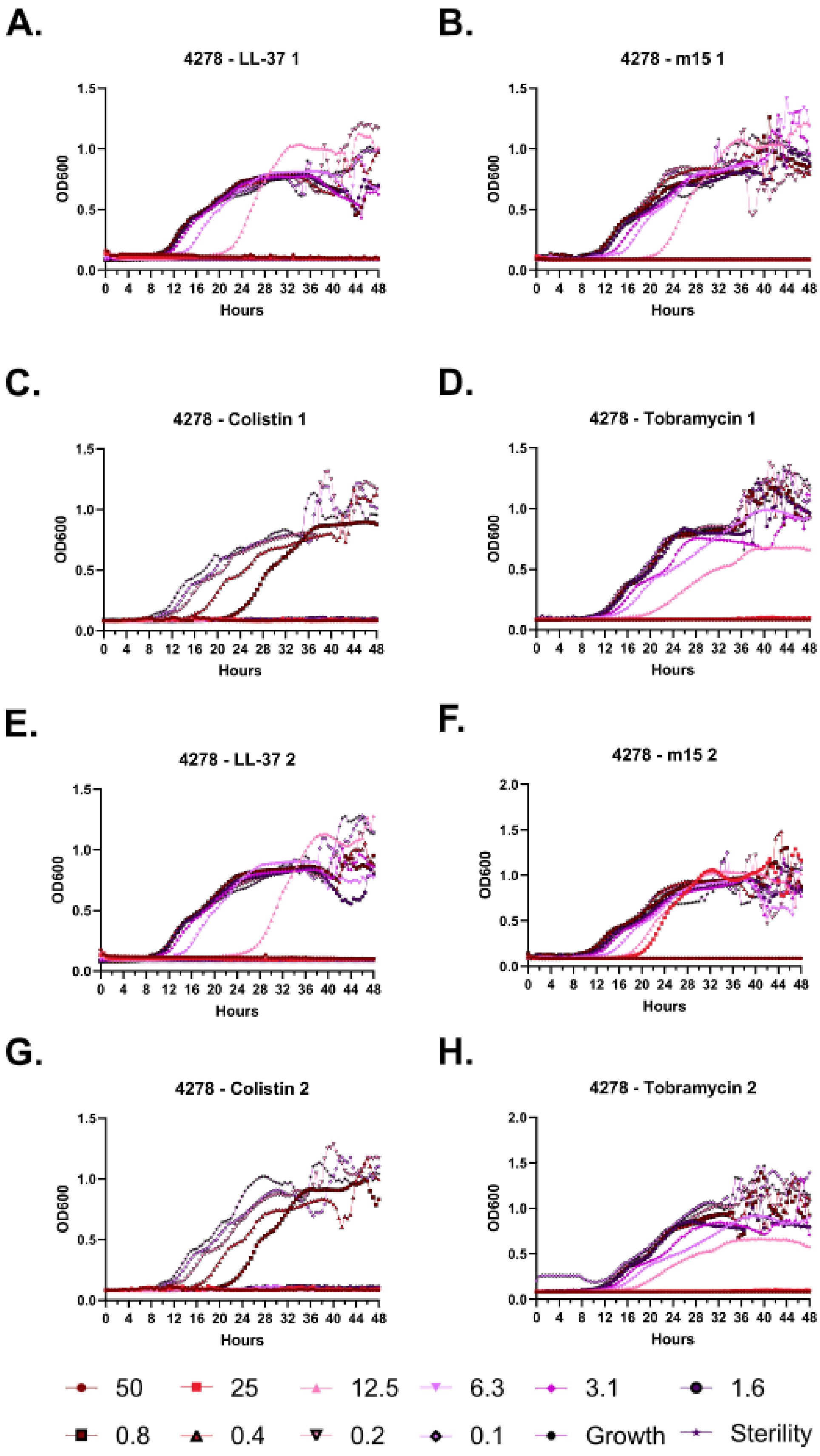
A.
4278 - LL-37 1
OD600
Hours
B.
4278 - m15 1
OD600
Hours
C.
4278 - Colistin 1
OD600
Hours
D.
4278 - Tobramycin 1
OD600
Hours
E.
4278 - LL-37 2
OD600
Hours
F.
4278 - m15 2
OD600
Hours
G.
4278 - Colistin 2
OD600
Hours
H.
4278 - Tobramycin 2
OD600
Hours
50
25
12.5
6.3
3.1
1.6
0.8
0.4
0.2
0.1
Growth
Sterility

**Supplementary Figure 106 – Kinetic antimicrobial susceptibility testing for *Pseudomonas* strain 4278. A-D.** Experiment 1. **E-H.** Experiment 2. Bacterial growth was monitored by OD600 over 48 hours at 37 °C in the presence of the indicated peptide concentrations. Peptide concentrations are given in μM.

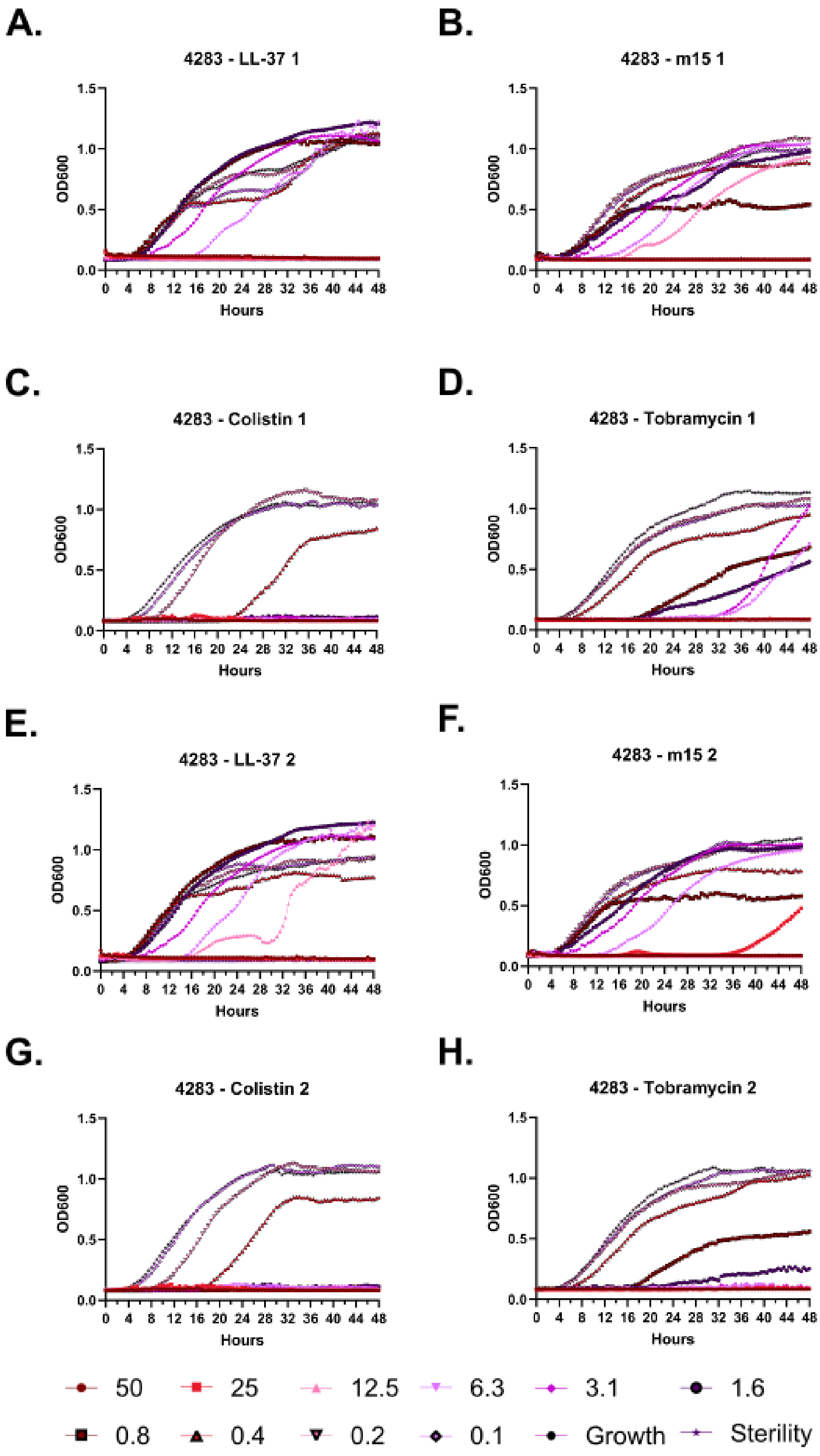
A.
4283 - LL-37 1
B.
4283 - m15 1
C.
4283 - Colistin 1
D.
4283 - Tobramycin 1
E.
4283 - LL-37 2
F.
4283 - m15 2
G.
4283 - Colistin 2
H.
4283 - Tobramycin 2
OD600
Hours
0.0 0.5 1.0 1.5
0 4 8 12 16 20 24 28 32 36 40 44 48
50
25
12.5
6.3
3.1
1.6
0.8
0.4
0.2
0.1
Growth
Sterility

**Supplementary Figure 107 – Kinetic antimicrobial susceptibility testing for *Pseudomonas* strain 4283. A-D.** Experiment 1. **E-H.** Experiment 2. Bacterial growth was monitored by OD600 over 48 hours at 37 °C in the presence of the indicated peptide concentrations. Peptide concentrations are given in μM.

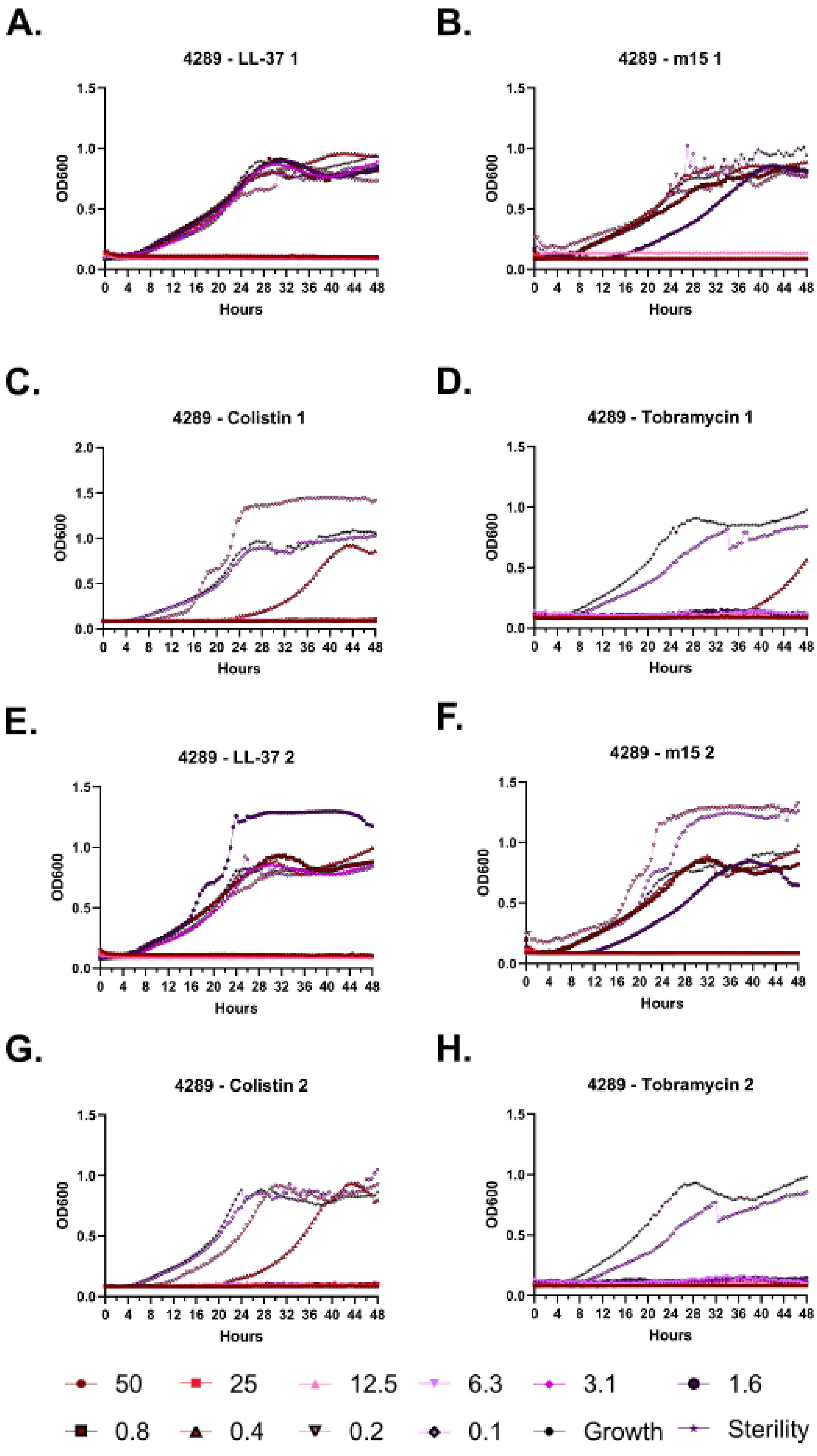

A.
4289 - LL-37 1
B.
4289 - m15 1
C.
4289 - Colistin 1
D.
4289 - Tobramycin 1
E.
4289 - LL-37 2
F.
4289 - m15 2
G.
4289 - Colistin 2
H.
4289 - Tobramycin 2
OD600
Hours
50
25
12.5
6.3
3.1
1.6
0.8
0.4
0.2
0.1
Growth
Sterility

**Supplementary Figure 108 – Kinetic antimicrobial susceptibility testing for *Pseudomonas* strain 4289. A-D.** Experiment 1. **E-H.** Experiment 2. Bacterial growth was monitored by OD600 over 48 hours at 37 °C in the presence of the indicated peptide concentrations. Peptide concentrations are given in μM.

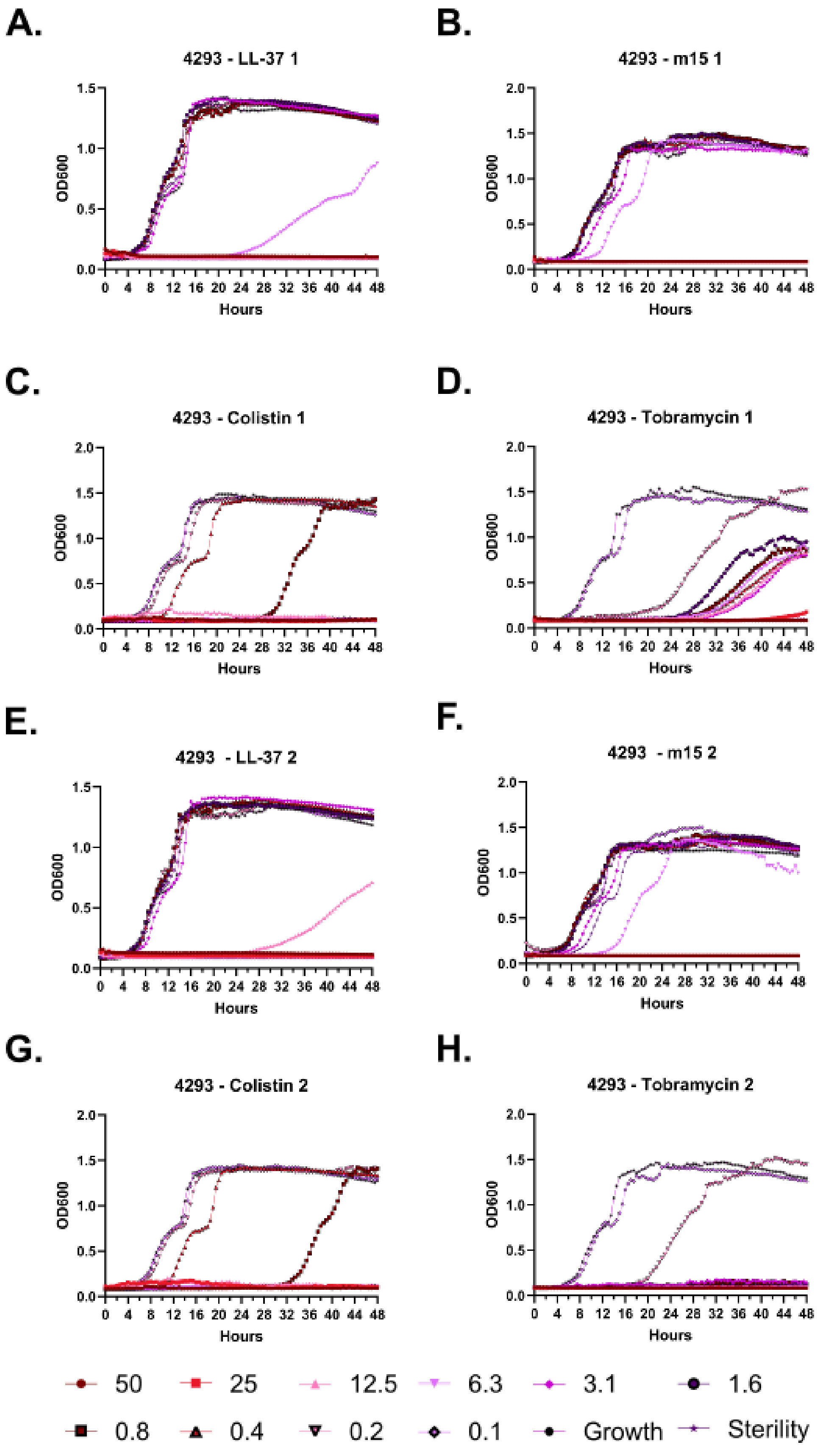
A.
4293 - LL-37 1
B.
4293 - m15 1
C.
4293 - Colistin 1
D.
4293 - Tobramycin 1
E.
4293 - LL-37 2
F.
4293 - m15 2
G.
4293 - Colistin 2
H.
4293 - Tobramycin 2
OD600
Hours
50
25
12.5
6.3
3.1
1.6
0.8
0.4
0.2
0.1
Growth
Sterility

**Supplementary Figure 109 – Kinetic antimicrobial susceptibility testing for *Pseudomonas* strain 4293. A-D.** Experiment 1. **E-H.** Experiment 2. Bacterial growth was monitored by OD600 over 48 hours at 37 °C in the presence of the indicated peptide concentrations. Peptide concentrations are given in μM.

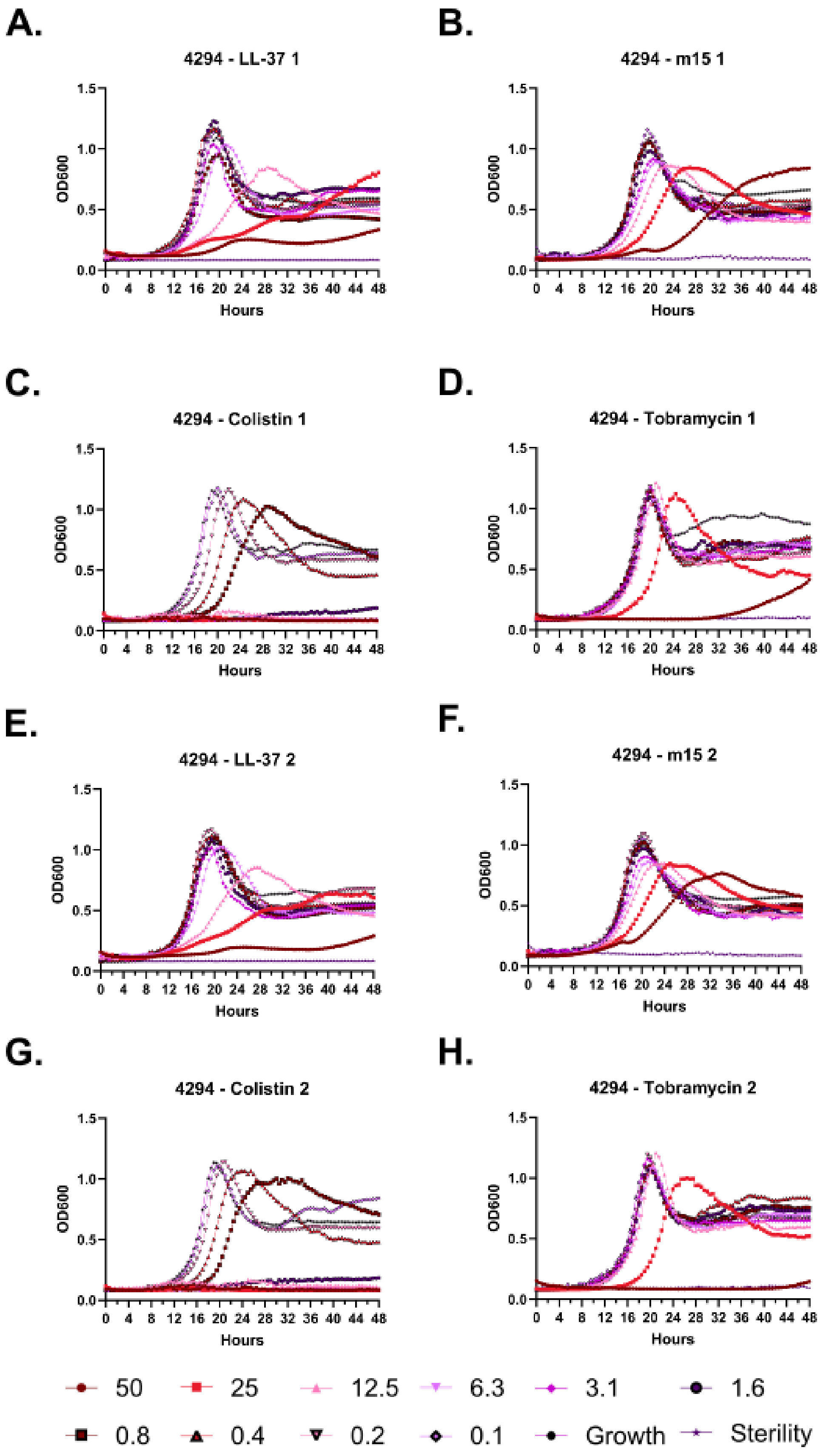
A.
4294 - LL-37 1
B.
4294 - m15 1
C.
4294 - Colistin 1
D.
4294 - Tobramycin 1
E.
4294 - LL-37 2
F.
4294 - m15 2
G.
4294 - Colistin 2
H.
4294 - Tobramycin 2
OD600
Hours
0 4 8 12 16 20 24 28 32 36 40 44 48
0.0 0.5 1.0 1.5
50
25
12.5
6.3
3.1
1.6
0.8
0.4
0.2
0.1
Growth
Sterility

**Supplementary Figure 110 – Kinetic antimicrobial susceptibility testing for *Pseudomonas* strain 4294. A-D.** Experiment 1. **E-H.** Experiment 2. Bacterial growth was monitored by OD600 over 48 hours at 37 °C in the presence of the indicated peptide concentrations. Peptide concentrations are given in μM.

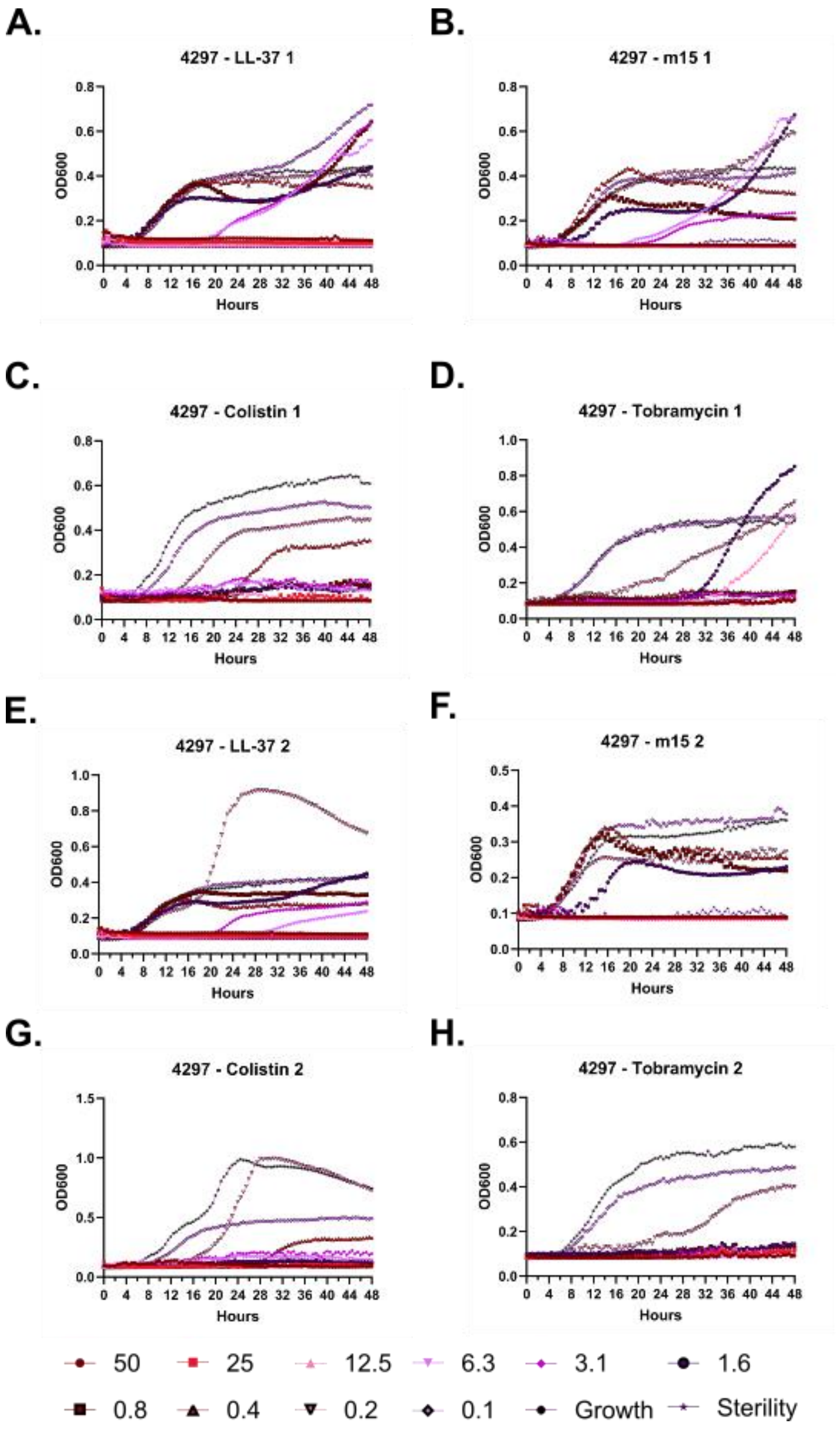
A.
4297 - LL-37 1
B.
4297 - m15 1
C.
4297 - Colistin 1
D.
4297 - Tobramycin 1
E.
4297 - LL-37 2
F.
4297 - m15 2
G.
4297 - Colistin 2
H.
4297 - Tobramycin 2
OD600
Hours
50
25
12.5
6.3
3.1
1.6
0.8
0.4
0.2
0.1
Growth
Sterility

**Supplementary Figure 111 – Kinetic antimicrobial susceptibility testing for *Pseudomonas* strain 4297. A-D.** Experiment 1. **E-H.** Experiment 2. Bacterial growth was monitored by OD600 over 48 hours at 37 °C in the presence of the indicated peptide concentrations. Peptide concentrations are given in μM.

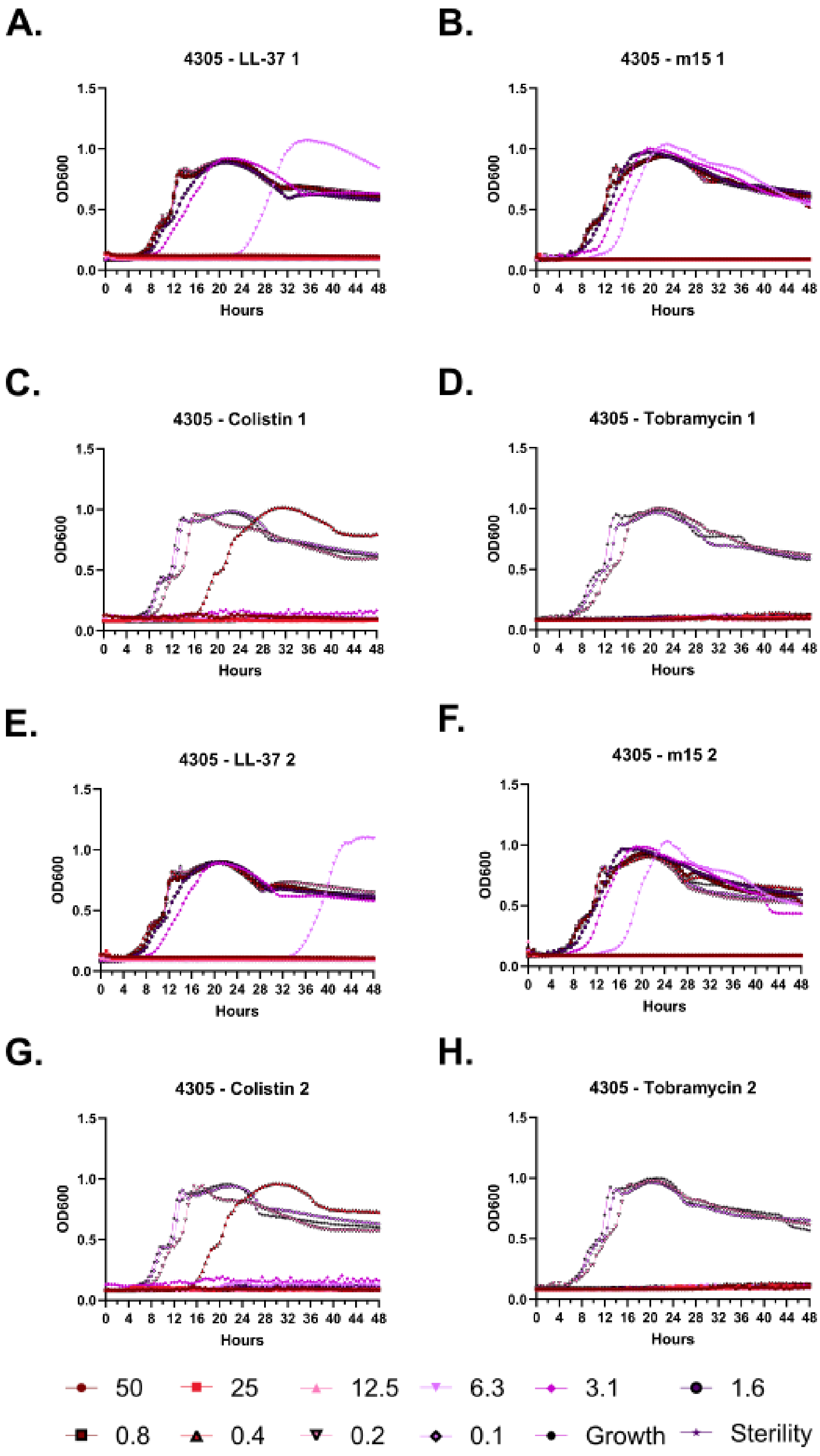
A.
4305 - LL-37 1
B.
4305 - m15 1
C.
4305 - Colistin 1
D.
4305 - Tobramycin 1
E.
4305 - LL-37 2
F.
4305 - m15 2
G.
4305 - Colistin 2
H.
4305 - Tobramycin 2
OD600
Hours
0.0
0.5
1.0
1.5
0 4 8 12 16 20 24 28 32 36 40 44 48
50
25
12.5
6.3
3.1
1.6
0.8
0.4
0.2
0.1
Growth
Sterility

**Supplementary Figure 112 – Kinetic antimicrobial susceptibility testing for *Pseudomonas* strain 4305. A-D.** Experiment 1. **E-H.** Experiment 2. Bacterial growth was monitored by OD600 over 48 hours at 37 °C in the presence of the indicated peptide concentrations. Peptide concentrations are given in μM.

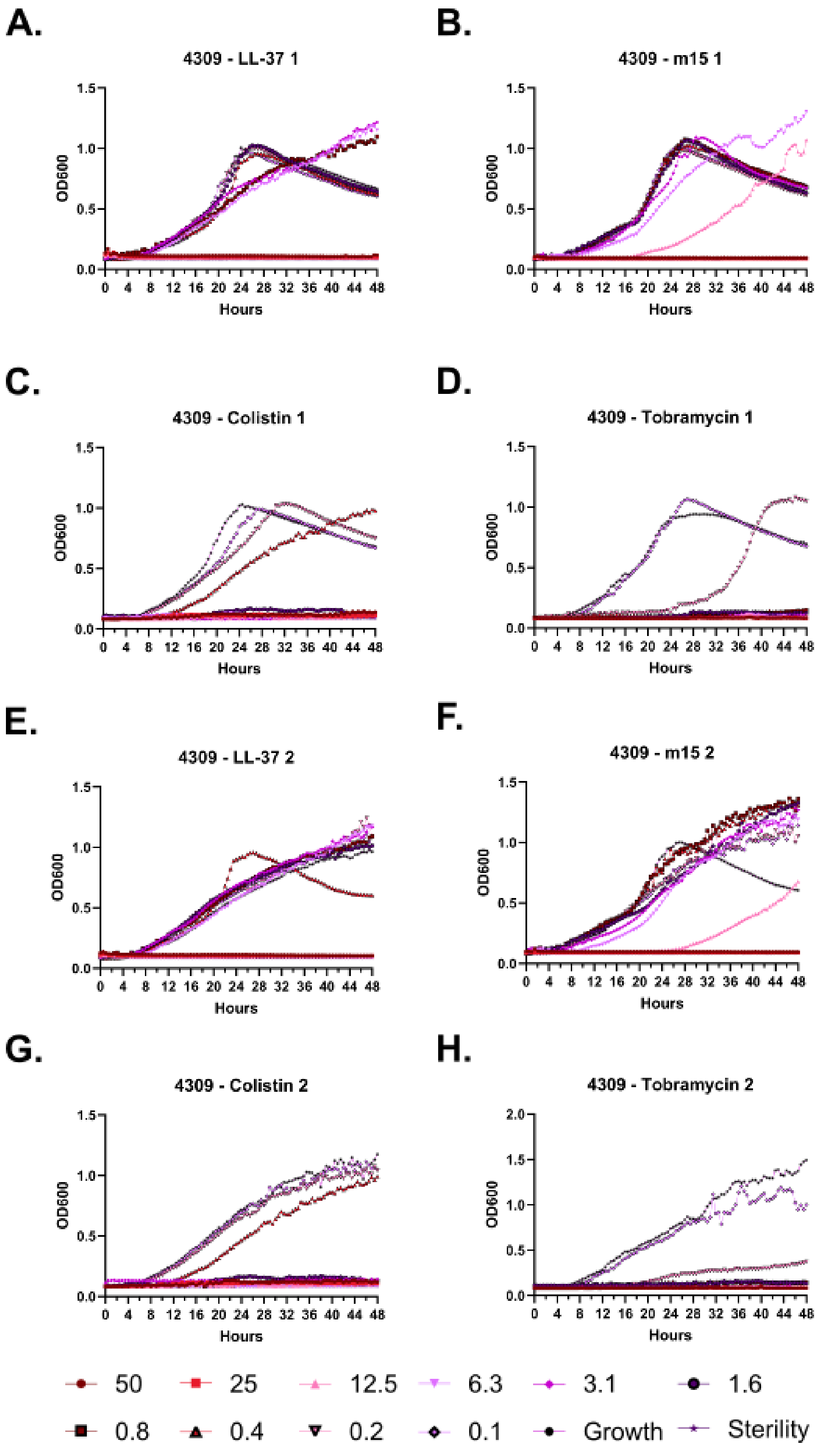
A.
4309 - LL-37 1
B.
4309 - m15 1
C.
4309 - Colistin 1
D.
4309 - Tobramycin 1
E.
4309 - LL-37 2
F.
4309 - m15 2
G.
4309 - Colistin 2
H.
4309 - Tobramycin 2
OD600
Hours
50
25
12.5
6.3
3.1
1.6
0.8
0.4
0.2
0.1
Growth
Sterility

**Supplementary Figure 113 – Kinetic antimicrobial susceptibility testing for *Pseudomonas* strain 4309. A-D.** Experiment 1. **E-H.** Experiment 2. Bacterial growth was monitored by OD600 over 48 hours at 37 °C in the presence of the indicated peptide concentrations. Peptide concentrations are given in μM.

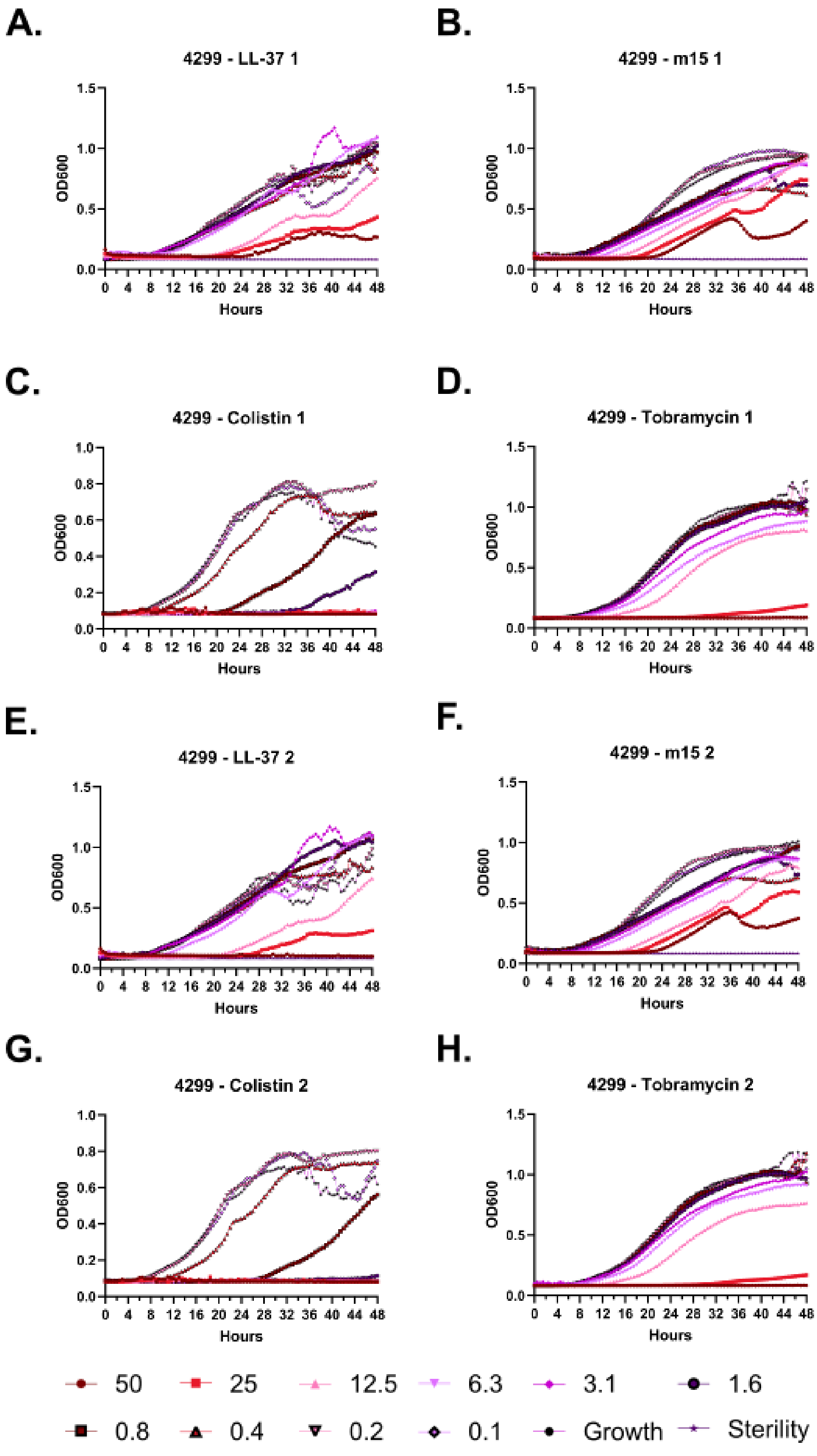
A.
4299 - LL-37 1
B.
4299 - m15 1
C.
4299 - Colistin 1
D.
4299 - Tobramycin 1
E.
4299 - LL-37 2
F.
4299 - m15 2
G.
4299 - Colistin 2
H.
4299 - Tobramycin 2
OD600
Hours
50
25
12.5
6.3
3.1
1.6
0.8
0.4
0.2
0.1
Growth
Sterility

**Supplementary Figure 114 – Kinetic antimicrobial susceptibility testing for *Pseudomonas* strain 4299. A-D.** Experiment 1. **E-H.** Experiment 2. Bacterial growth was monitored by OD600 over 48 hours at 37 °C in the presence of the indicated peptide concentrations. Peptide concentrations are given in μM.

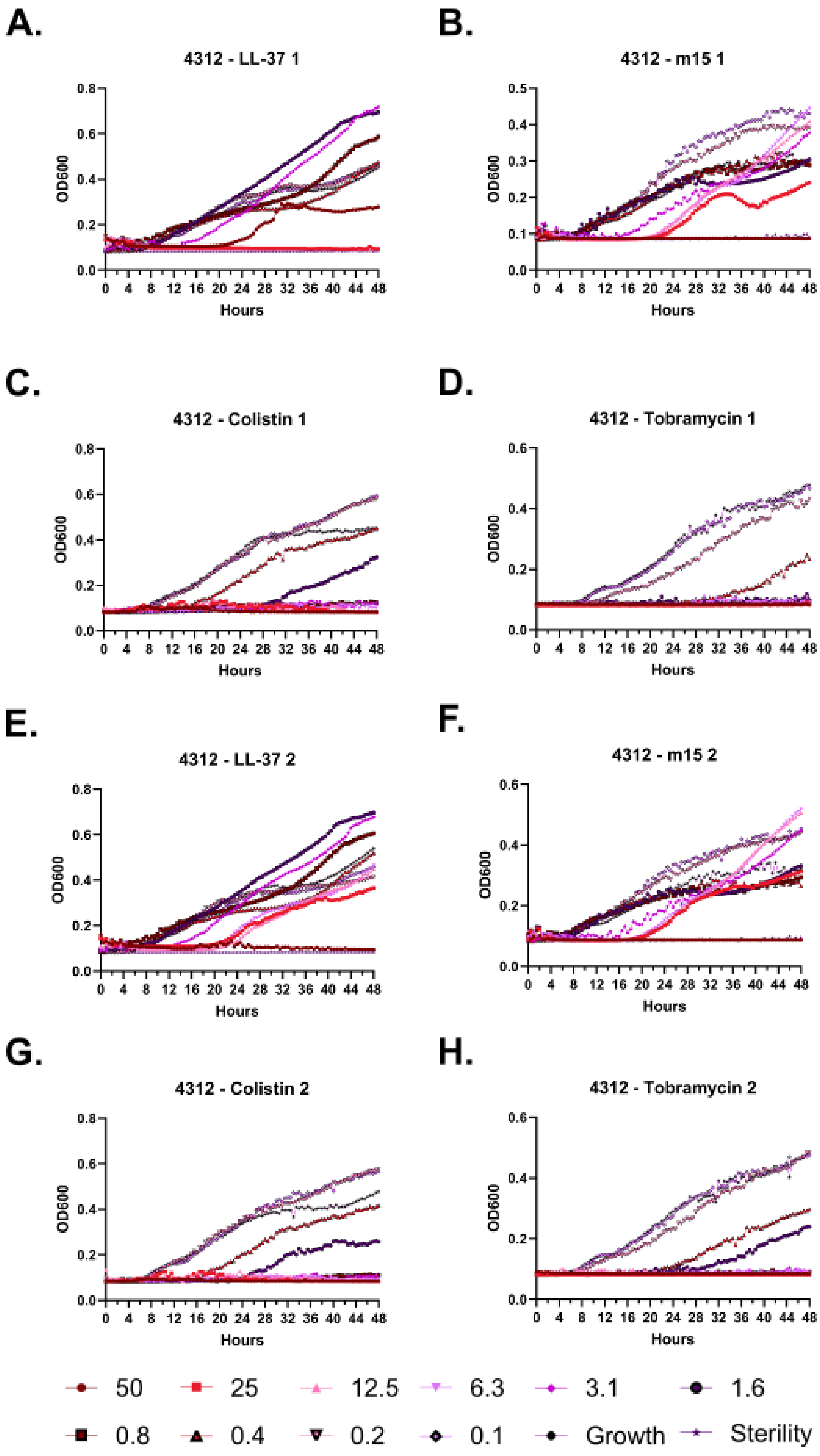
A.
4312 - LL-37 1
B.
4312 - m15 1
C.
4312 - Colistin 1
D.
4312 - Tobramycin 1
E.
4312 - LL-37 2
F.
4312 - m15 2
G.
4312 - Colistin 2
H.
4312 - Tobramycin 2
OD600
Hours
0 4 8 12 16 20 24 28 32 36 40 44 48
50
25
12.5
6.3
3.1
1.6
0.8
0.4
0.2
0.1
Growth
Sterility

**Supplementary Figure 115 – Kinetic antimicrobial susceptibility testing for *Pseudomonas* strain 4312. A-D.** Experiment 1. **E-H.** Experiment 2. Bacterial growth was monitored by OD600 over 48 hours at 37 °C in the presence of the indicated peptide concentrations. Peptide concentrations are given in μM.

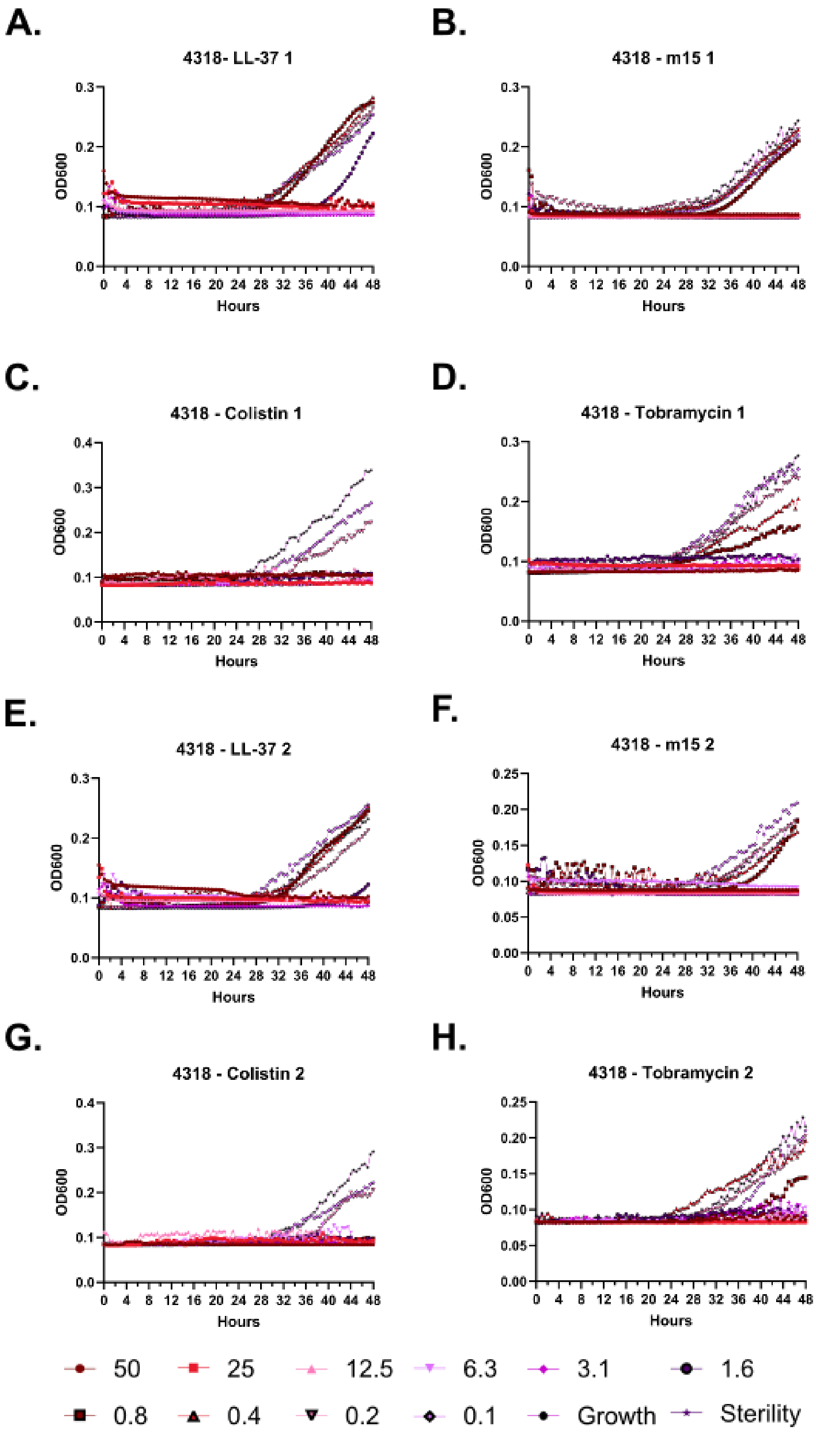
A.
4318- LL-37 1
B.
4318 - m15 1
C.
4318 - Colistin 1
D.
4318 - Tobramycin 1
E.
4318 - LL-37 2
F.
4318 - m15 2
G.
4318 - Colistin 2
H.
4318 - Tobramycin 2
OD600
Hours
0 4 8 12 16 20 24 28 32 36 40 44 48
50
25
12.5
6.3
3.1
1.6
0.8
0.4
0.2
0.1
Growth
Sterility

**Supplementary Figure 116 – Kinetic antimicrobial susceptibility testing for *Pseudomonas* strain 4318. A-D.** Experiment 1. **E-H.** Experiment 2. Bacterial growth was monitored by OD600 over 48 hours at 37 °C in the presence of the indicated peptide concentrations. Peptide concentrations are given in μM.

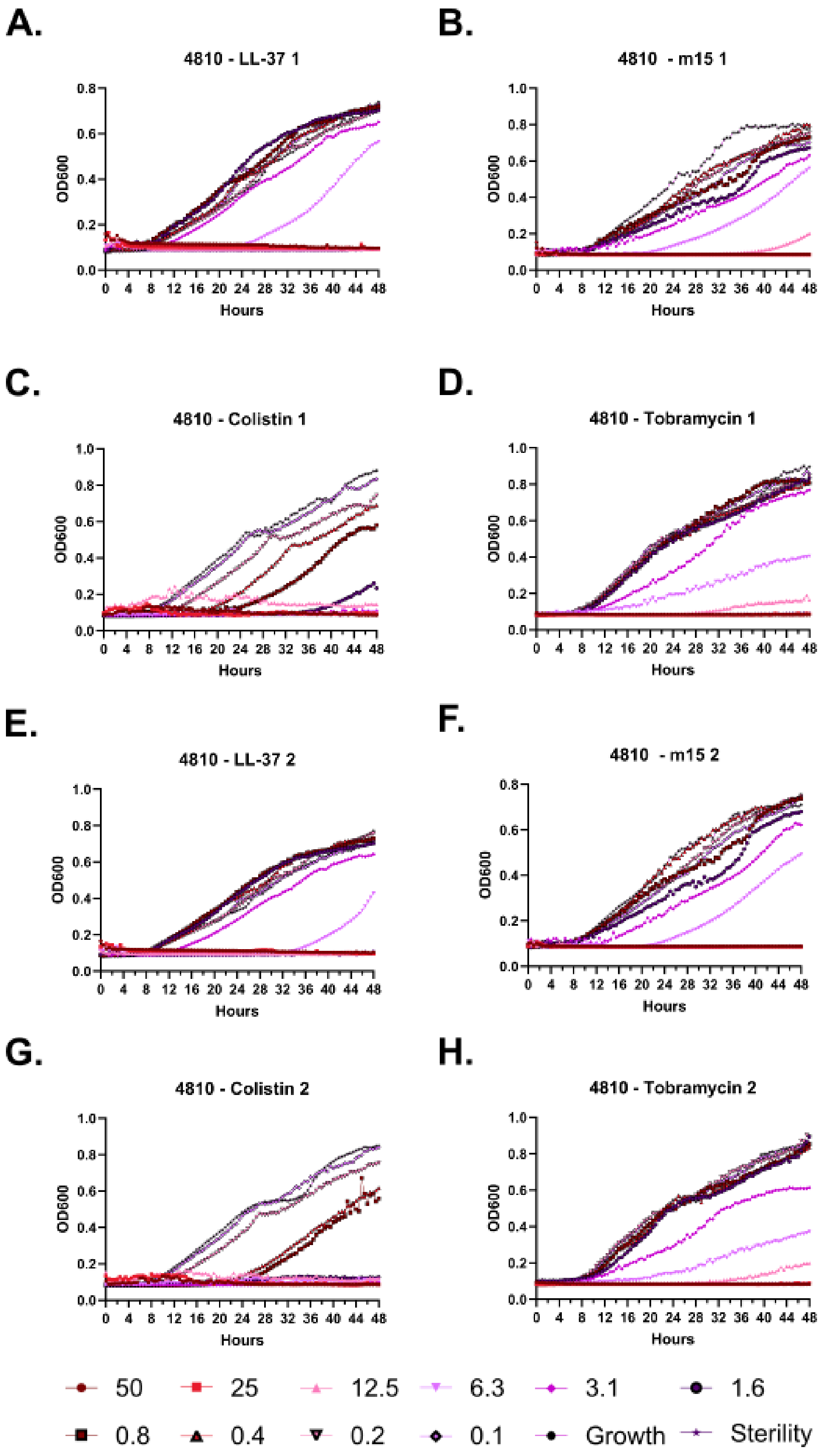
A.
4810 - LL-37 1
B.
4810 - m15 1
C.
4810 - Colistin 1
D.
4810 - Tobramycin 1
E.
4810 - LL-37 2
F.
4810 - m15 2
G.
4810 - Colistin 2
H.
4810 - Tobramycin 2
OD600
Hours
50
25
12.5
6.3
3.1
1.6
0.8
0.4
0.2
0.1
Growth
Sterility

**Supplementary Figure 117 – Kinetic antimicrobial susceptibility testing for *Pseudomonas* strain 4810. A-D.** Experiment 1. **E-H.** Experiment 2. Bacterial growth was monitored by OD600 over 48 hours at 37 °C in the presence of the indicated peptide concentrations. Peptide concentrations are given in μM.

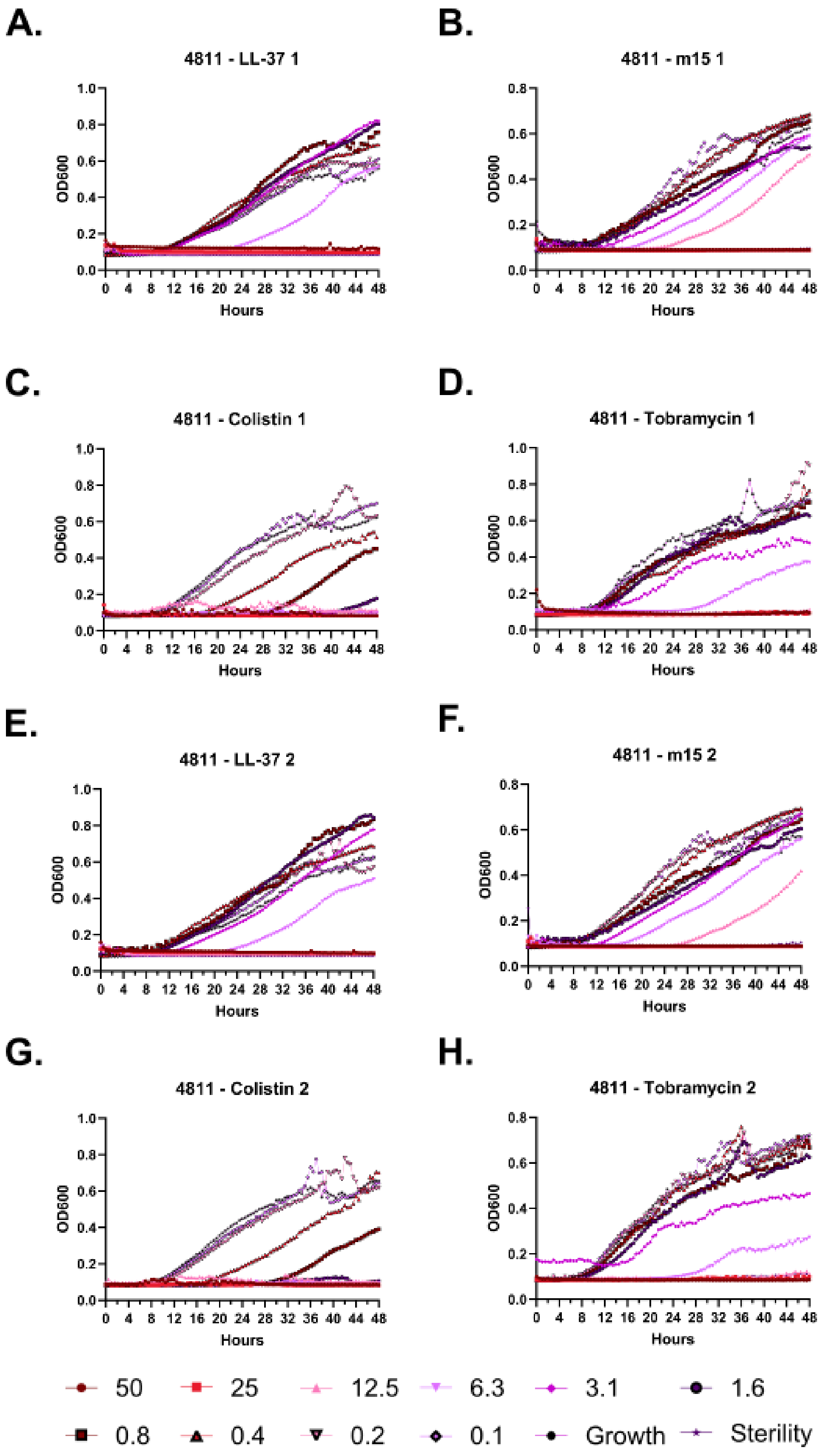
A.
4811 - LL-37 1
B.
4811 - m15 1
C.
4811 - Colistin 1
D.
4811 - Tobramycin 1
E.
4811 - LL-37 2
F.
4811 - m15 2
G.
4811 - Colistin 2
H.
4811 - Tobramycin 2
OD600
Hours
0 4 8 12 16 20 24 28 32 36 40 44 48
50
25
12.5
6.3
3.1
1.6
0.8
0.4
0.2
0.1
Growth
Sterility

**Supplementary Figure 118 – Kinetic antimicrobial susceptibility testing for *Pseudomonas* strain 4811. A-D.** Experiment 1. **E-H.** Experiment 2. Bacterial growth was monitored by OD600 over 48 hours at 37 °C in the presence of the indicated peptide concentrations. Peptide concentrations are given in µM.

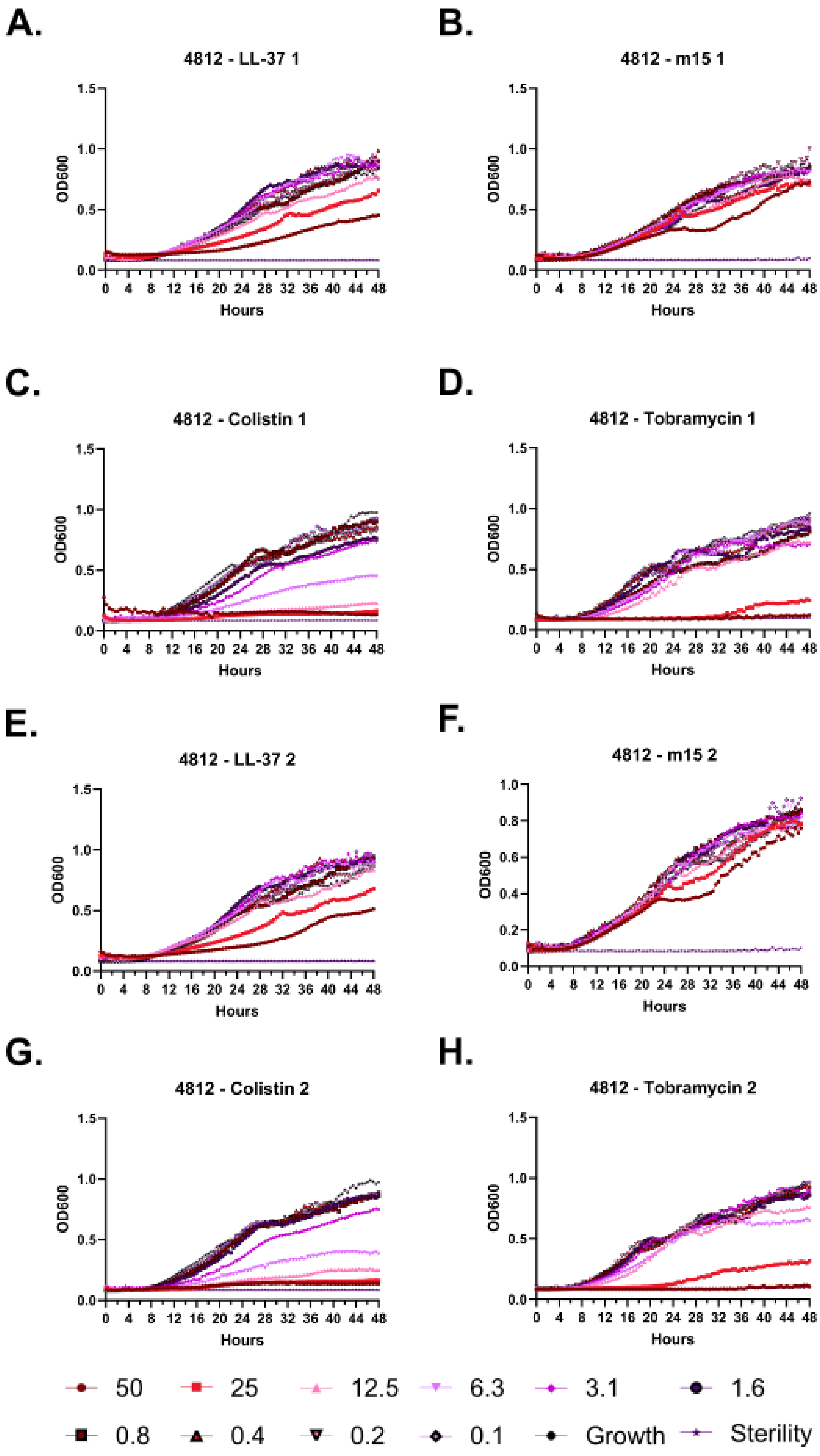
A.
4812 - LL-37 1
B.
4812 - m15 1
C.
4812 - Colistin 1
D.
4812 - Tobramycin 1
E.
4812 - LL-37 2
F.
4812 - m15 2
G.
4812 - Colistin 2
H.
4812 - Tobramycin 2
OD600
Hours
50
25
12.5
6.3
3.1
1.6
0.8
0.4
0.2
0.1
Growth
Sterility

**Supplementary Figure 119 – Kinetic antimicrobial susceptibility testing for *Pseudomonas* strain 4812. A-D.** Experiment 1. **E-H.** Experiment 2. Bacterial growth was monitored by OD600 over 48 hours at 37 °C in the presence of the indicated peptide concentrations. Peptide concentrations are given in μM.

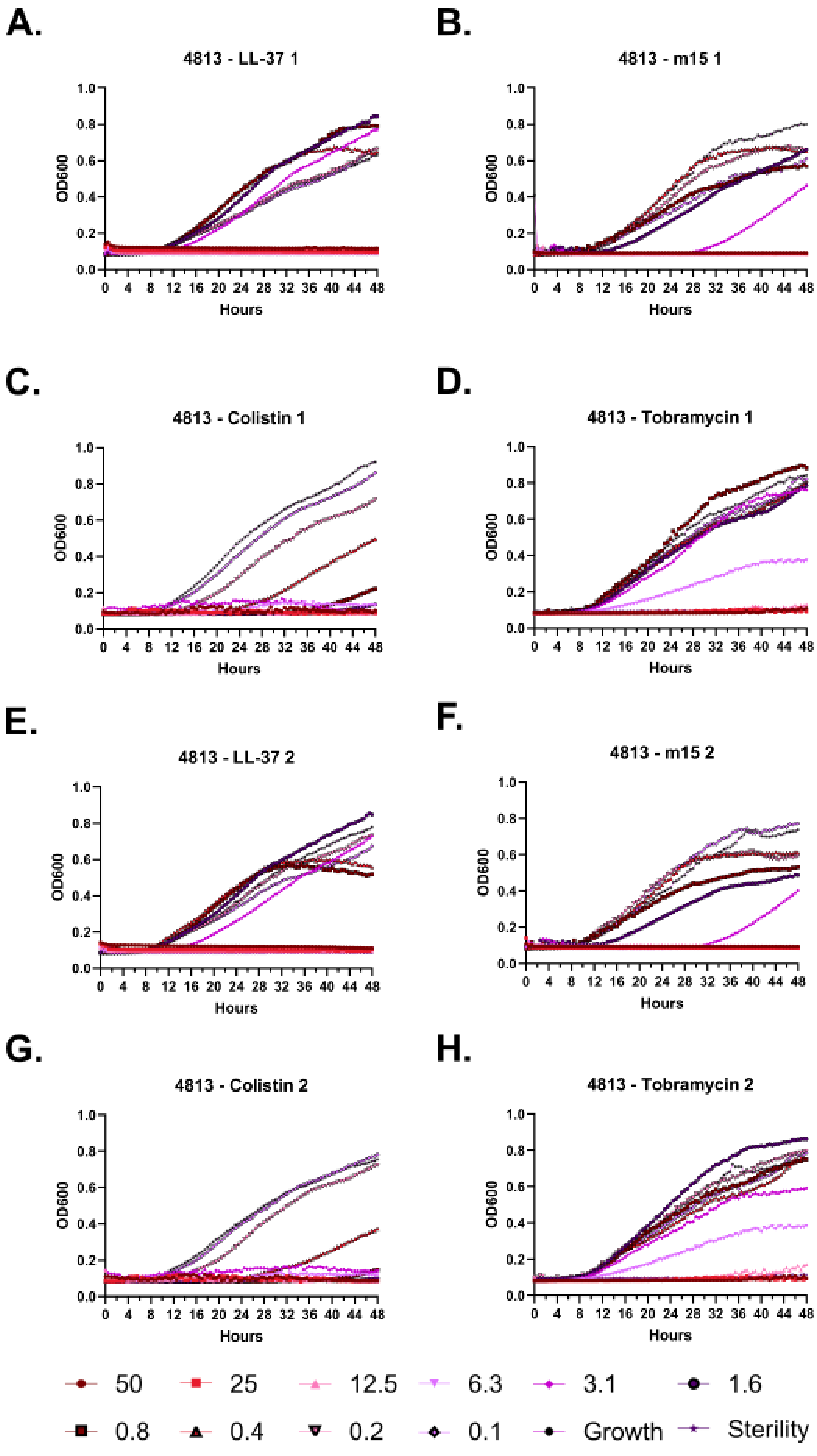
A.
4813 - LL-37 1
B.
4813 - m15 1
C.
4813 - Colistin 1
D.
4813 - Tobramycin 1
E.
4813 - LL-37 2
F.
4813 - m15 2
G.
4813 - Colistin 2
H.
4813 - Tobramycin 2
OD600
Hours
0.0 0.2 0.4 0.6 0.8 1.0
0 4 8 12 16 20 24 28 32 36 40 44 48
50
25
12.5
6.3
3.1
1.6
0.8
0.4
0.2
0.1
Growth
Sterility

**Supplementary Figure 120 – Kinetic antimicrobial susceptibility testing for *Pseudomonas* strain 4813. A-D.** Experiment 1. **E-H.** Experiment 2. Bacterial growth was monitored by OD600 over 48 hours at 37 °C in the presence of the indicated peptide concentrations. Peptide concentrations are given in μM.

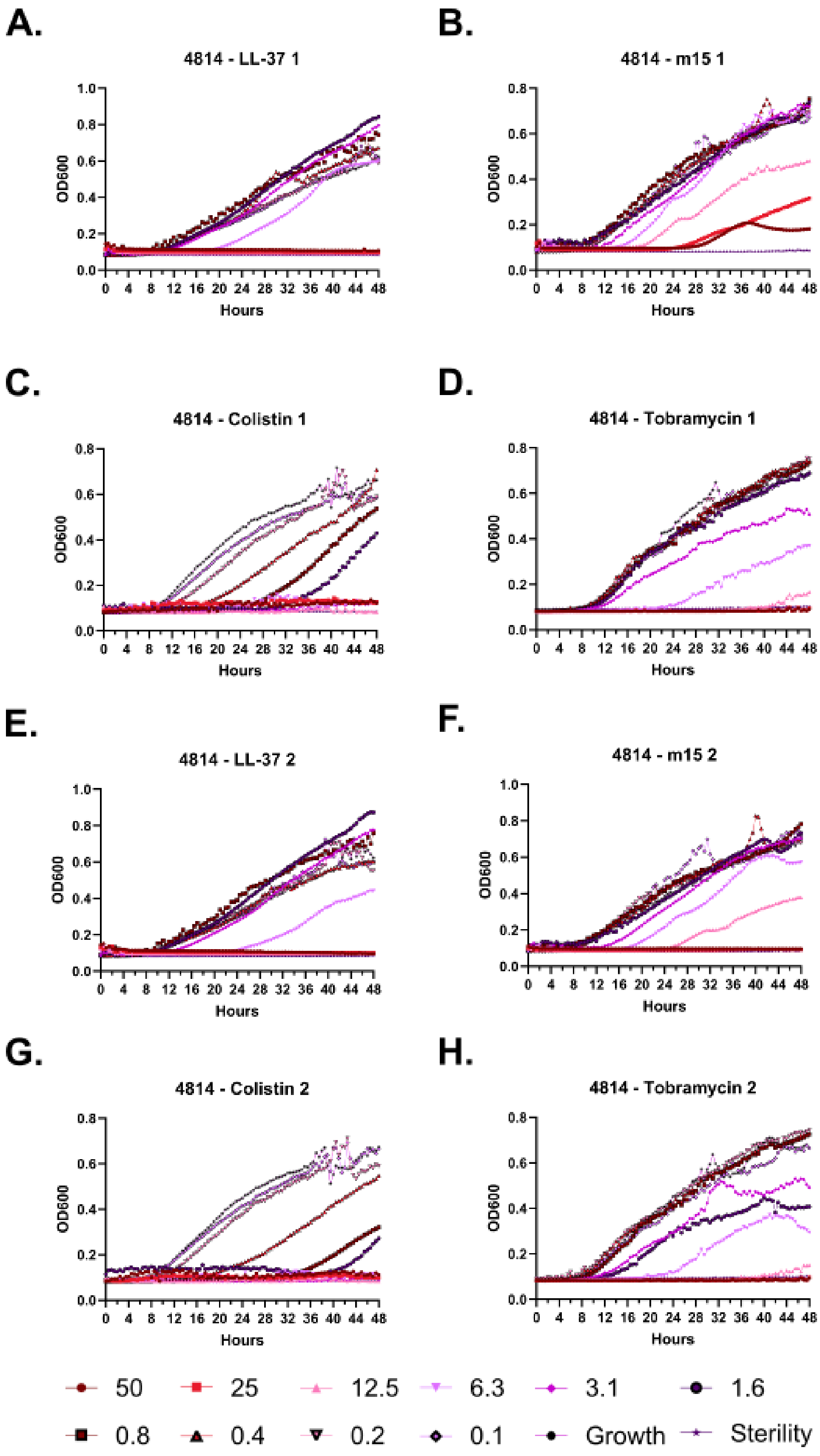
A.
4814 - LL-37 1
B.
4814 - m15 1
C.
4814 - Colistin 1
D.
4814 - Tobramycin 1
E.
4814 - LL-37 2
F.
4814 - m15 2
G.
4814 - Colistin 2
H.
4814 - Tobramycin 2
OD600
Hours
0 4 8 12 16 20 24 28 32 36 40 44 48
50
25
12.5
6.3
3.1
1.6
0.8
0.4
0.2
0.1
Growth
Sterility

**Supplementary Figure 121 – Kinetic antimicrobial susceptibility testing for *Pseudomonas* strain 4814. A-D.** Experiment 1. **E-H.** Experiment 2. Bacterial growth was monitored by OD600 over 48 hours at 37 °C in the presence of the indicated peptide concentrations. Peptide concentrations are given in μM.

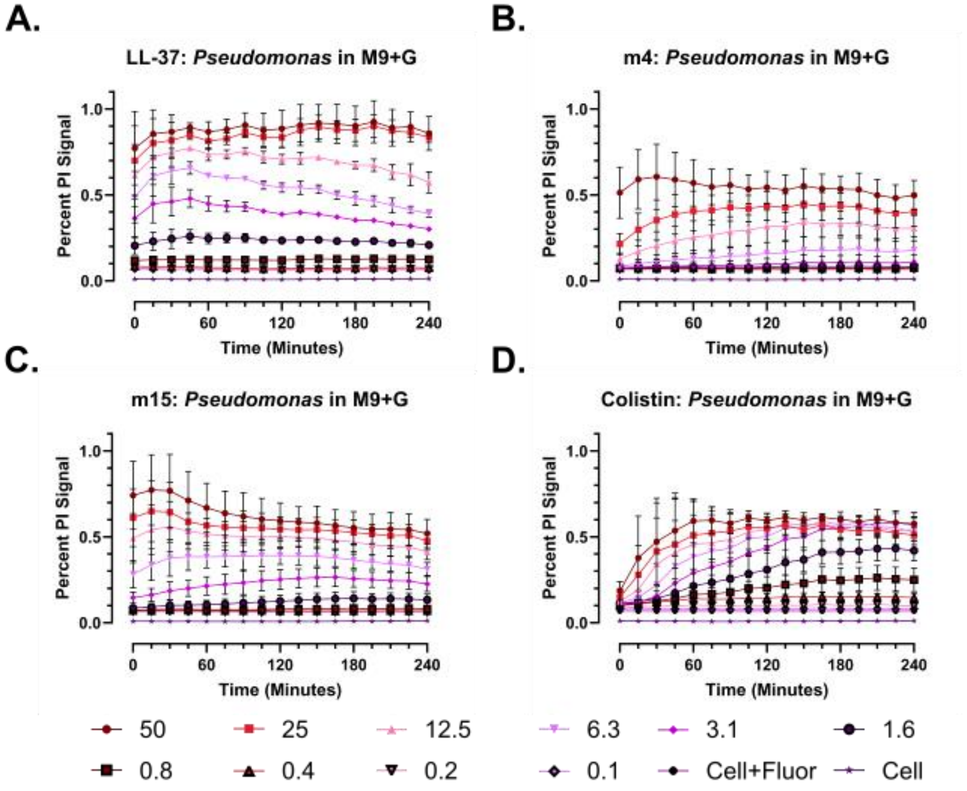
A.
LL-37: Pseudomonas in M9+G
B.
m4: Pseudomonas in M9+G
C.
m15: Pseudomonas in M9+G
D.
Colistin: Pseudomonas in M9+G
Percent PI Signal
Time (Minutes)
0 60 120 180 240
0.0 0.5 1.0
50
25
12.5
6.3
3.1
1.6
0.8
0.4
0.2
0.1
Cell+Fluor
Cell

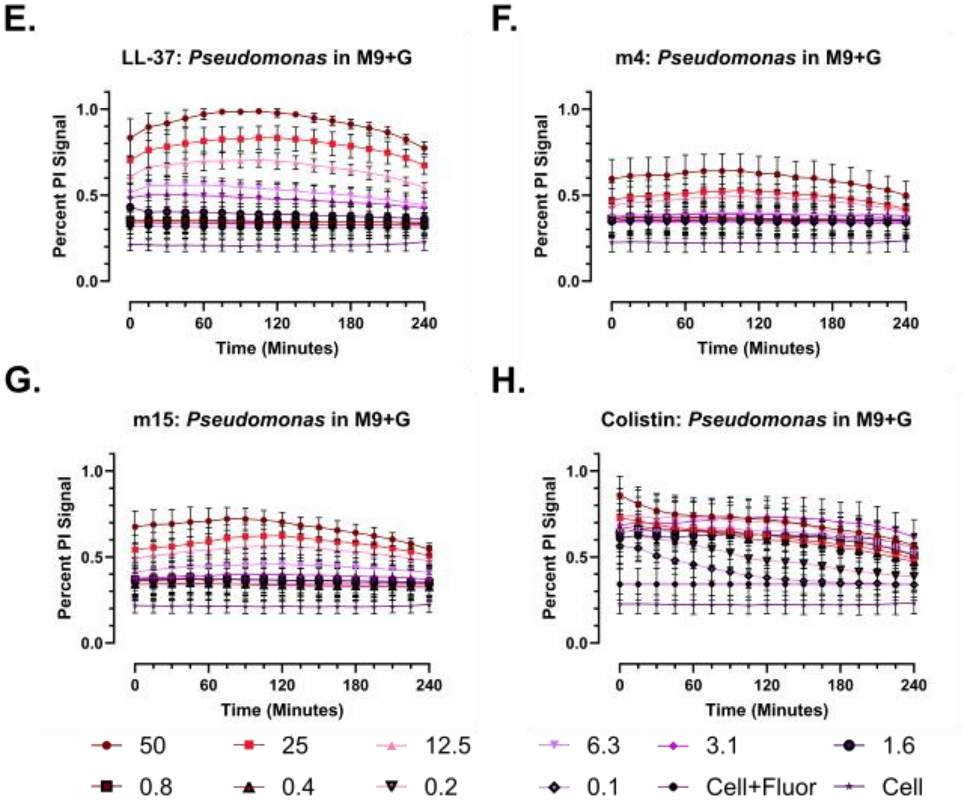
E.
LL-37: Pseudomonas in M9+G
F.
m4: Pseudomonas in M9+G
G.
m15: Pseudomonas in M9+G
H.
Colistin: Pseudomonas in M9+G
Percent PI Signal
Time (Minutes)
0 60 120 180 240
0.0 0.5 1.0
50
25
12.5
6.3
3.1
1.6
0.8
0.4
0.2
0.1
Cell+Fluor
Cell

**Figure 122 – Oligomerization-deficient mutants demonstrate attenuated membrane permeabilization activity in *Pseudomonas*. A-D.** Propidium iodide (PI) staining for inner membrane permeabilization over 4 hours in the presence of each peptide at the indicated concentrations given in μM. Percentage is based on calculation of a fold-change in PI fluorescent signal (535 / Em 617) over baseline within a condition. **E-H.** N-Phenyl-1-naphthylamine (NPN) staining for outer membrane permeabilization over 4 hours in the presence of each peptide at the indicated concentration given in in μM. Quantification proceeds as for PI, except that the highest NPN signal (350 / Em 420) timepoint is set to 100%. Data throughout show the mean and standard deviation of three independent experiments in 50 μL in black, transparent-bottom 384-well plates incubated at 37 °C.

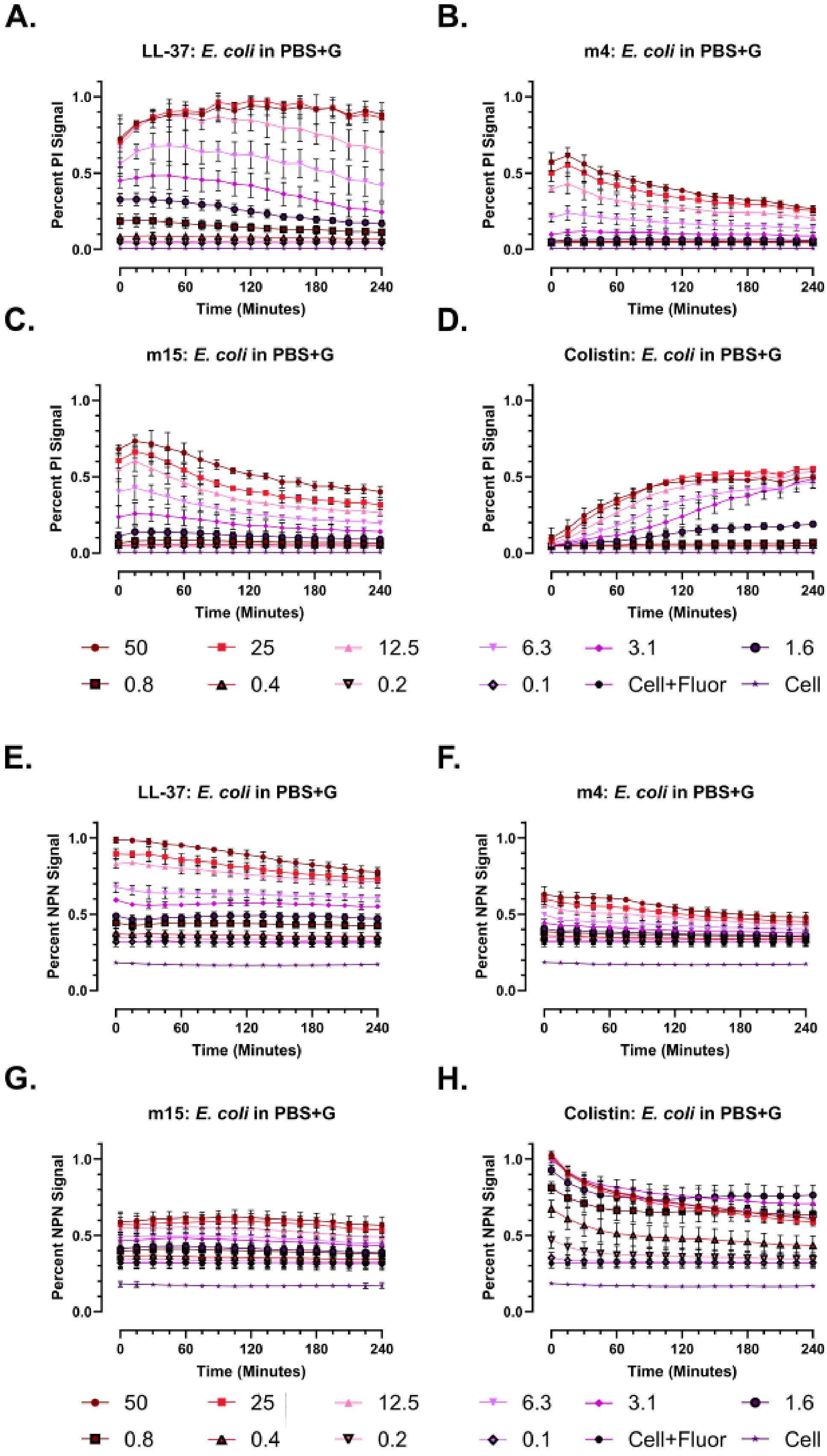
A.
LL-37: E. coli in PBS+G
B.
m4: E. coli in PBS+G
C.
m15: E. coli in PBS+G
D.
Colistin: E. coli in PBS+G
Percent PI Signal
Time (Minutes)
0.0
0.5
1.0
0
60
120
180
240
50
25
12.5
6.3
3.1
1.6
0.8
0.4
0.2
0.1
Cell+Fluor
Cell
E.
LL-37: E. coli in PBS+G
F.
m4: E. coli in PBS+G
G.
m15: E. coli in PBS+G
H.
Colistin: E. coli in PBS+G
Percent NPN Signal
Time (Minutes)

**Supplementary Figure 123 – Attenuation of membrane permeabilization among oligomerization-deficient mutants is less prominent in PBS+glucose. A-D.** Propidium iodide (PI, Ex 535 / Em 617) staining for inner membrane permeabilization over 4 hours in the presence of each peptide at the indicated concentrations given in µM. **E-H.** N-Phenyl-Naphthylamine (NPN, Ex 350 / Em 420) staining for outer membrane permeabilization over 4 hours in the presence of each peptide at the indicated concentration given in in µM. Data throughout show the mean and standard deviation of three independent experiments in 50 µL in black 384-well plates incubated at 37 °C.

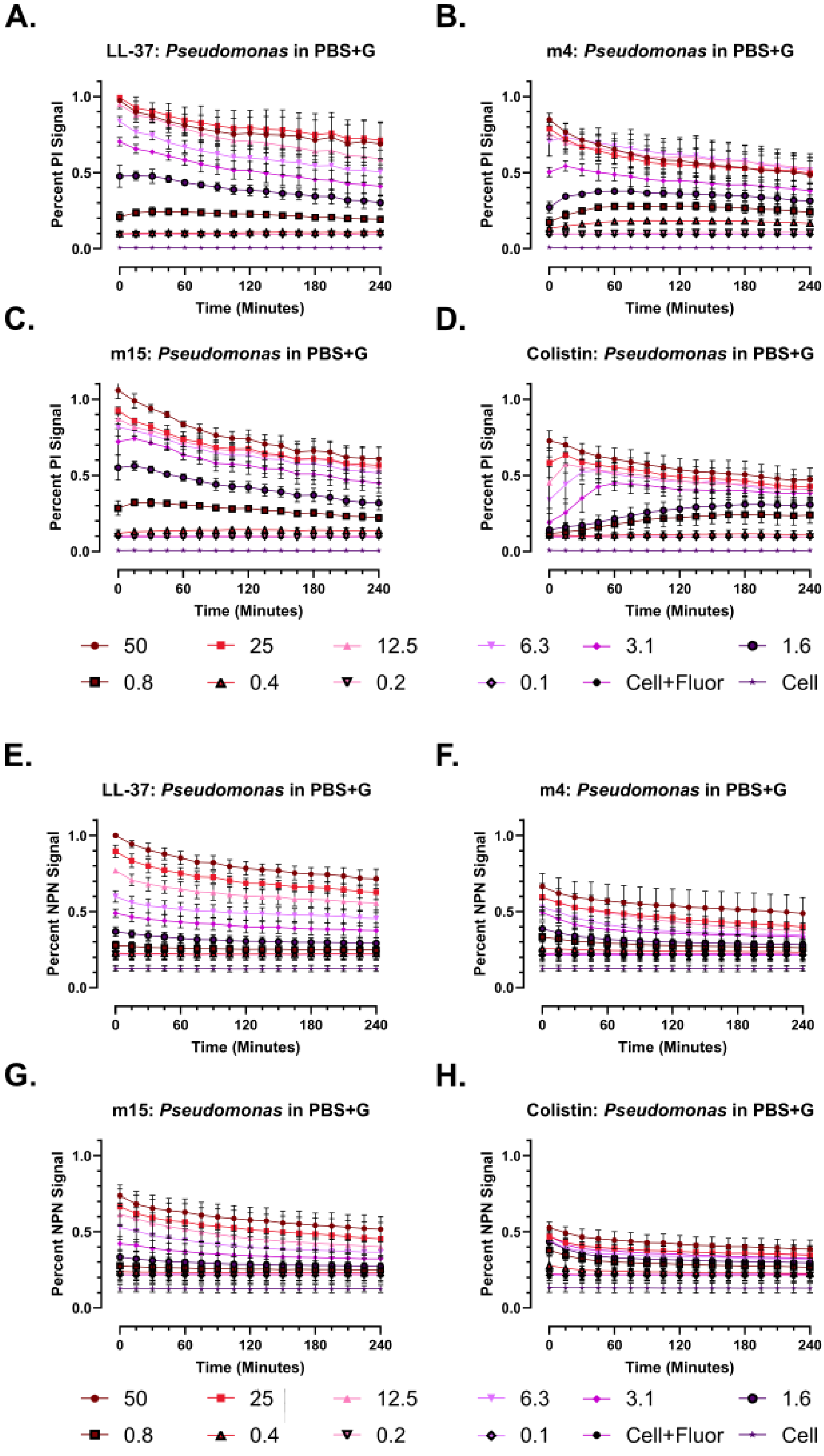
A.
LL-37: Pseudomonas in PBS+G
B.
m4: Pseudomonas in PBS+G
C.
m15: Pseudomonas in PBS+G
D.
Colistin: Pseudomonas in PBS+G
Percent PI Signal
Time (Minutes)
0
60
120
180
240
0.0
0.5
1.0
50
25
12.5
6.3
3.1
1.6
0.8
0.4
0.2
0.1
Cell+Fluor
Cell
E.
LL-37: Pseudomonas in PBS+G
F.
m4: Pseudomonas in PBS+G
G.
m15: Pseudomonas in PBS+G
H.
Colistin: Pseudomonas in PBS+G
Percent NPN Signal
Time (Minutes)

**Supplementary Figure 124 – Oligomerization-deficient mutants demonstrate membrane permeabilization comparable to wild-type LL-37 in *Pseudomonas* in PBS+glucose. A-D.** Propidium iodide (PI, Ex 535 / Em 617) staining for inner membrane permeabilization over 4 hours in the presence of each peptide at the indicated concentrations given in μM. **E-H.** N-Phenyl-1-Naphthylamine (NPN, Ex 350 / Em 420) staining for outer membrane permeabilization over 4 hours in the presence of each peptide at the indicated concentration given in in μM. Data throughout show the mean and standard deviation of three independent experiments in 50 μL in black, transparent-bottom 384-well plates incubated at 37 °C.

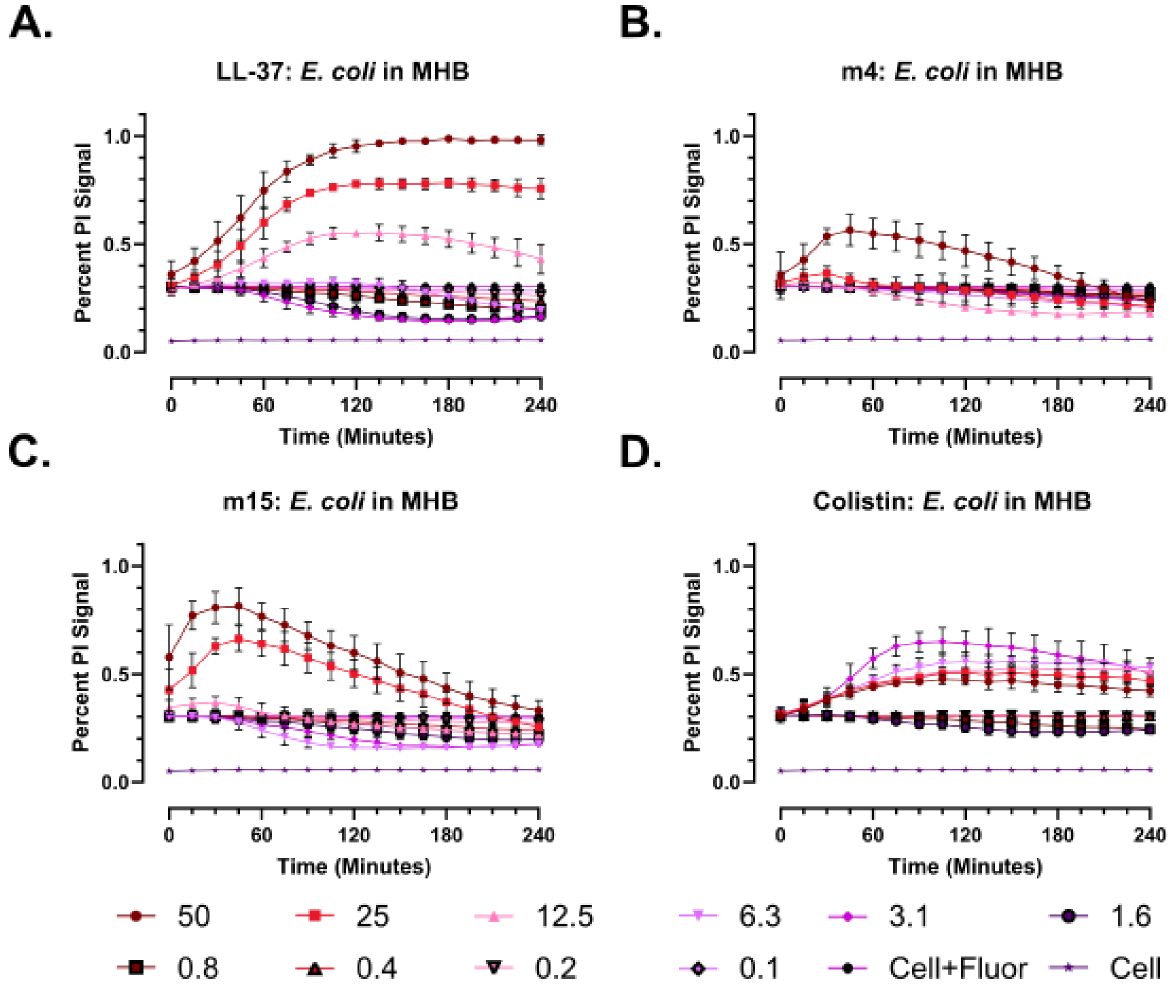
A.
LL-37: E. coli in MHB
B.
m4: E. coli in MHB
C.
m15: E. coli in MHB
D.
Colistin: E. coli in MHB
Percent PI Signal
Time (Minutes)
0.0
0.5
1.0
0
60
120
180
240
50
25
12.5
6.3
3.1
1.6
0.8
0.4
0.2
0.1
Cell+Fluor
Cell

**Supplementary Figure 125 – Oligomerization-deficient mutants demonstrate attenuated membrane permeabilization activity in *E. coli* in MHB. A-D.** Propidium iodide (PI, Ex 535 / Em 617) staining for inner membrane permeabilization over 4 hours in the presence of each peptide at the indicated concentrations given in μM. Outer membrane staining with N-Phenyl-1-Naphthylamine was unrevealing due to high background fluorescence in MHB (data not shown). Data throughout show the mean and standard deviation of three independent experiments in 50 μL in black, transparent-bottom 384-well plates incubated at 37 °C.

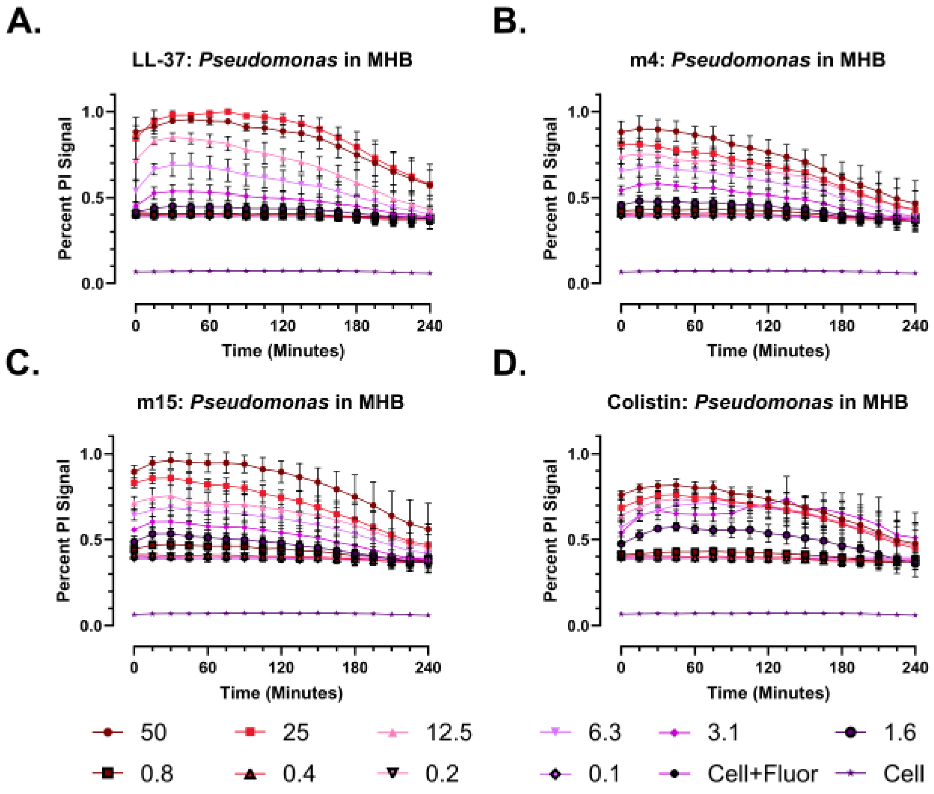
A.
LL-37: Pseudomonas in MHB
B.
m4: Pseudomonas in MHB
C.
m15: Pseudomonas in MHB
D.
Colistin: Pseudomonas in MHB
Percent PI Signal
0.0
0.5
1.0
Time (Minutes)
0
60
120
180
240
50
25
12.5
6.3
3.1
1.6
0.8
0.4
0.2
0.1
Cell+Fluor
Cell

**Supplementary Figure 126 – Oligomerization-deficient mutants demonstrate membrane permeabilization of *Pseudomonas* comparable to wild-type LL-37 in MHB. A-D.** Propidium iodide (PI, Ex 535 / Em 617) staining for inner membrane permeabilization over 4 hours in the presence of each peptide at the indicated concentrations given in μM. Outer membrane staining with N-Phenyl-1-Naphthylamine (NPN) was unrevealing due to high background fluorescence in MHB (data not shown). Data throughout show the mean and standard deviation of three independent experiments in 50 μL in black, transparent bottom 384-well plates incubated at 37 °C.

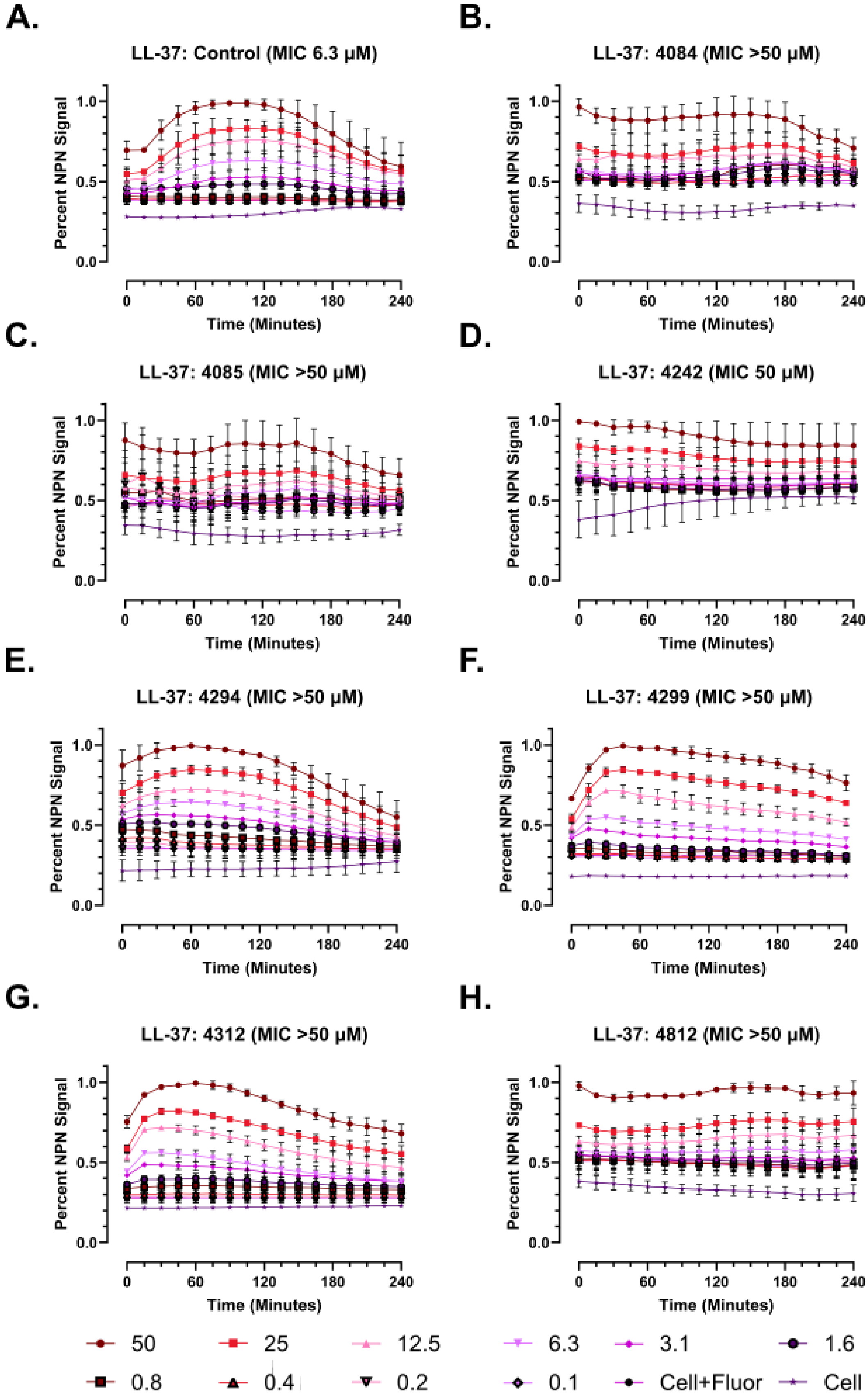
A.
LL-37: Control (MIC 6.3 µM)
B.
LL-37: 4084 (MIC >50 µM)
C.
LL-37: 4085 (MIC >50 µM)
D.
LL-37: 4242 (MIC 50 µM)
E.
LL-37: 4294 (MIC >50 µM)
F.
LL-37: 4299 (MIC >50 µM)
G.
LL-37: 4312 (MIC >50 µM)
H.
LL-37: 4812 (MIC >50 µM)
Percent NPN Signal
Time (Minutes)
0.0
0.5
1.0
0
60
120
180
240
50
25
12.5
6.3
3.1
1.6
0.8
0.4
0.2
0.1
Cell+Fluor
Cell

**Supplementary Figure 127 – The outer membranes of *Pseudomonas* with elevated LL-37 MICs are efficiently permeabilized by LL-37.** A control strain, ATCC 27853, and seven clinical strains of *Pseudomonas* from patients with cystic fibrosis were evaluated in dual membrane permeability assays. This figure shows the outer membrane staining (NPN signal) from the same experiments as **Figure 6**. **A-D.** Data throughout show the mean and standard deviation of three independent experiments in 50 μL in black, transparent-bottom 384-well plates incubated at 37 °C.

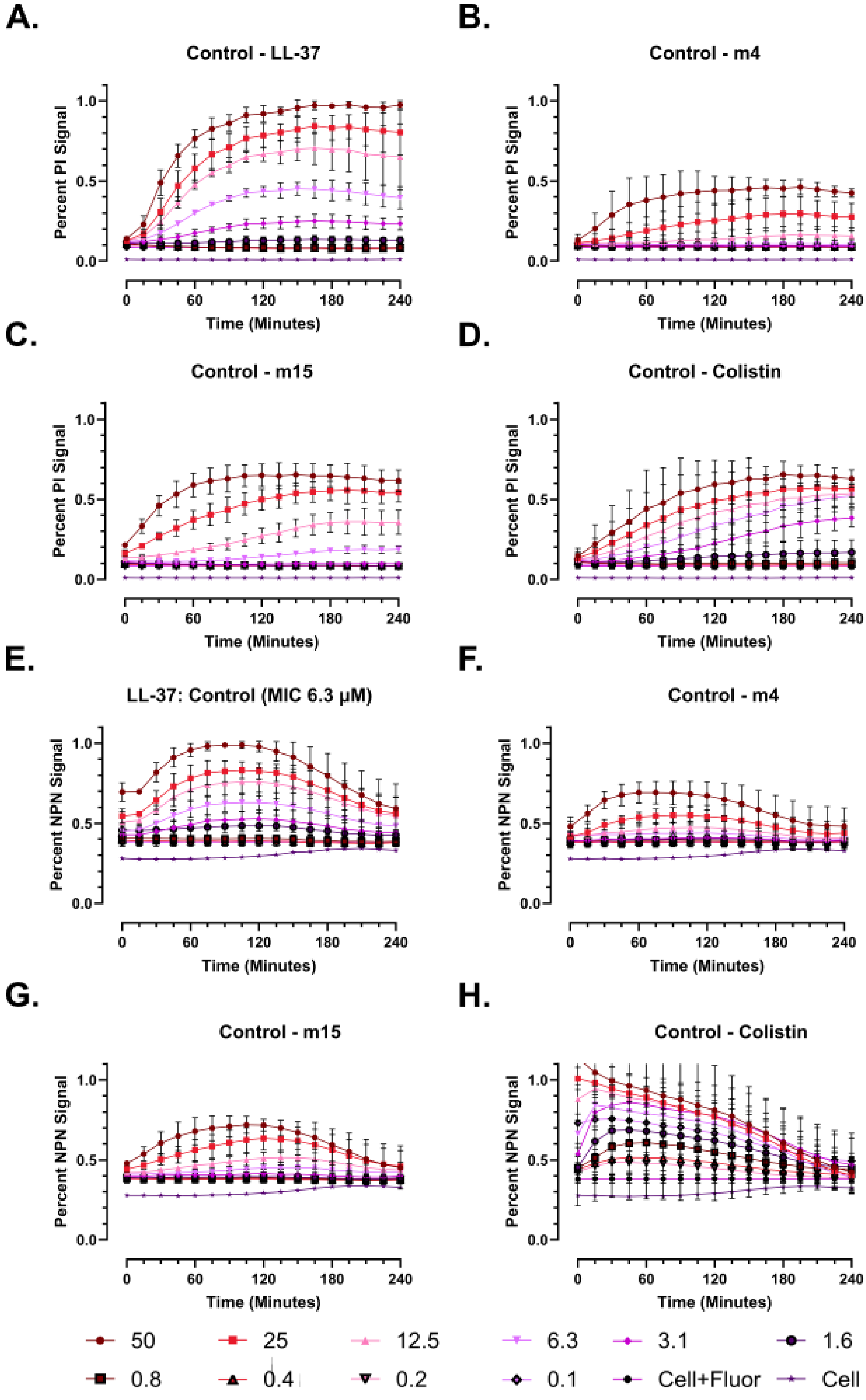
A.
Control - LL-37
Percent PI Signal
Time (Minutes)
B.
Control - m4
Percent PI Signal
Time (Minutes)
C.
Control - m15
Percent PI Signal
Time (Minutes)
D.
Control - Colistin
Percent PI Signal
Time (Minutes)
E.
LL-37: Control (MIC 6.3 µM)
Percent NPN Signal
Time (Minutes)
F.
Control - m4
Percent NPN Signal
Time (Minutes)
G.
Control - m15
Percent NPN Signal
Time (Minutes)
H.
Control - Colistin
Percent NPN Signal
Time (Minutes)
50
25
12.5
6.3
3.1
1.6
0.8
0.4
0.2
0.1
Cell+Fluor
Cell

**Supplementary Figure 128 – Membrane permeabilization patterns in control strain ATCC 27853.** Control strain ATCC 27853 was used in dual membrane permeabilization assays as described in the main text. **A-D.** Inner membrane permeabilization, PI signal (Ex 535 / Em 617). **E-H.** Inner membrane permeabilization, NPN signal (Ex 350 / Em 420).

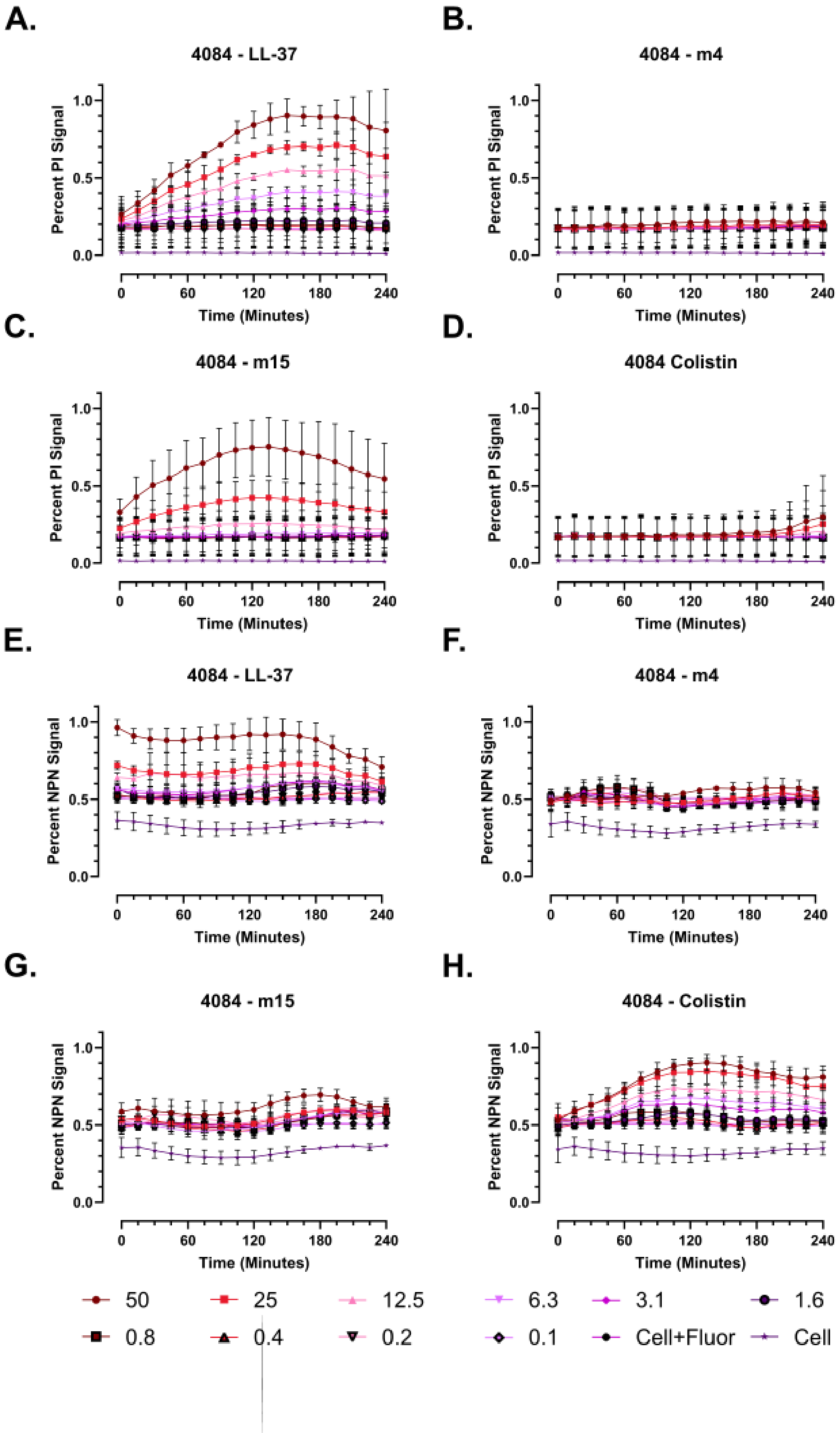

A.
4084 - LL-37
B.
4084 - m4
C.
4084 - m15
D.
4084 Colistin
E.
4084 - LL-37
F.
4084 - m4
G.
4084 - m15
H.
4084 - Colistin
Percent PI Signal
Percent NPN Signal
Time (Minutes)
0.0
0.5
1.0
0
60
120
180
240
50
25
12.5
6.3
3.1
1.6
0.8
0.4
0.2
0.1
Cell+Fluor
Cell

**Supplementary Figure 129 – Membrane permeabilization patterns in clinical strain 4084.** Clinical *Pseudomonas* strain 4084 was used in dual membrane permeabilization assays as described in the main text. **A-D.** Inner membrane permeabilization, PI signal (Ex 535 / Em 617). **E-H.** Inner membrane permeabilization, NPN signal (Ex 350 / Em 420).

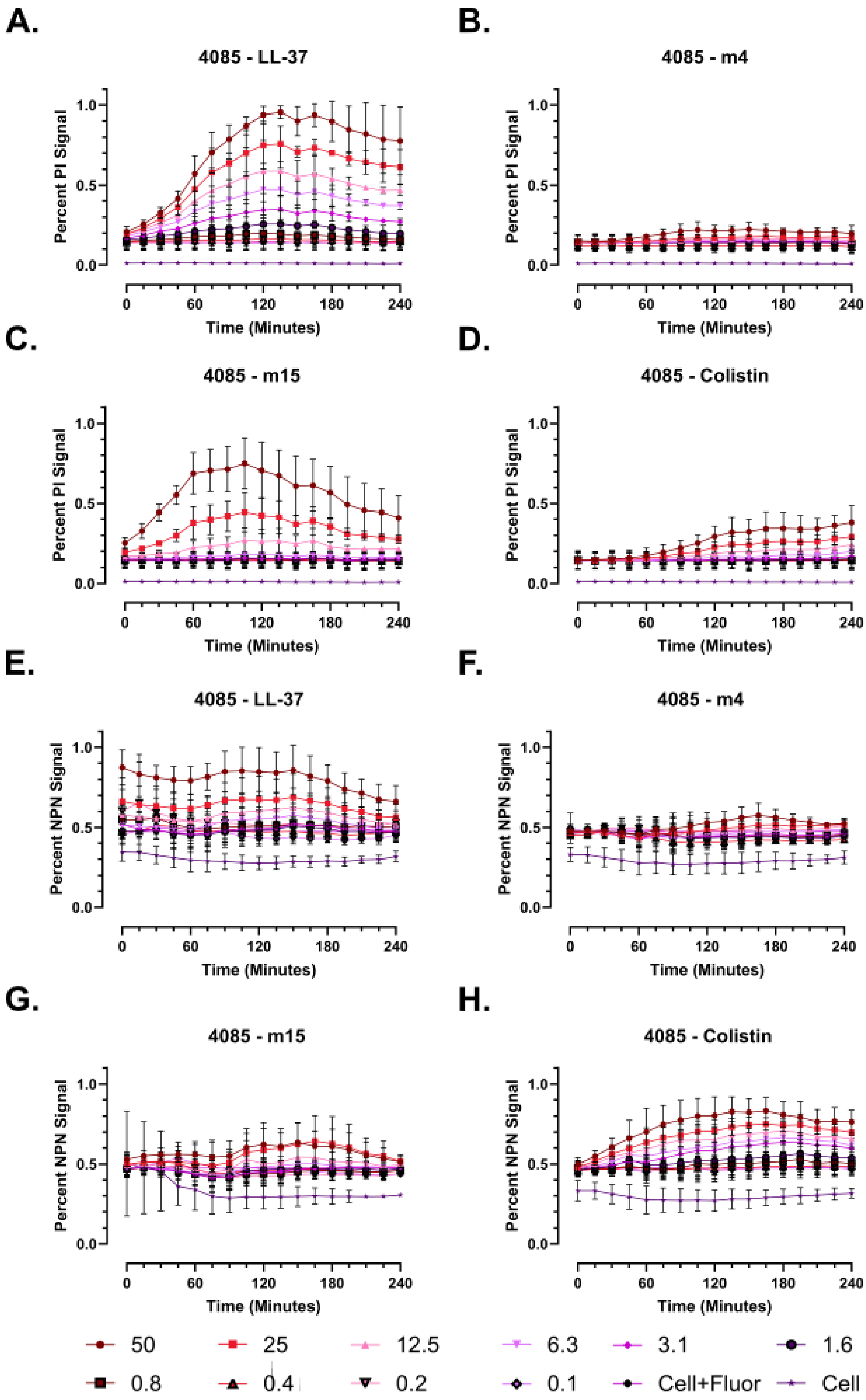

A.
4085 - LL-37
B.
4085 - m4
C.
4085 - m15
D.
4085 - Colistin
E.
4085 - LL-37
F.
4085 - m4
G.
4085 - m15
H.
4085 - Colistin
Percent PI Signal
Percent NPN Signal
Time (Minutes)
0.0
0.5
1.0
0
60
120
180
240
50
25
12.5
6.3
3.1
1.6
0.8
0.4
0.2
0.1
Cell+Fluor
Cell

**Supplementary Figure 130 – Membrane permeabilization patterns in clinical strain 4085.** Clinical *Pseudomonas* strain 4085 was used in dual membrane permeabilization assays as described in the main text. **A-D.** Inner membrane permeabilization, PI signal (Ex 535 / Em 617). **E-H.** Inner membrane permeabilization, NPN signal (Ex 350 / Em 420).

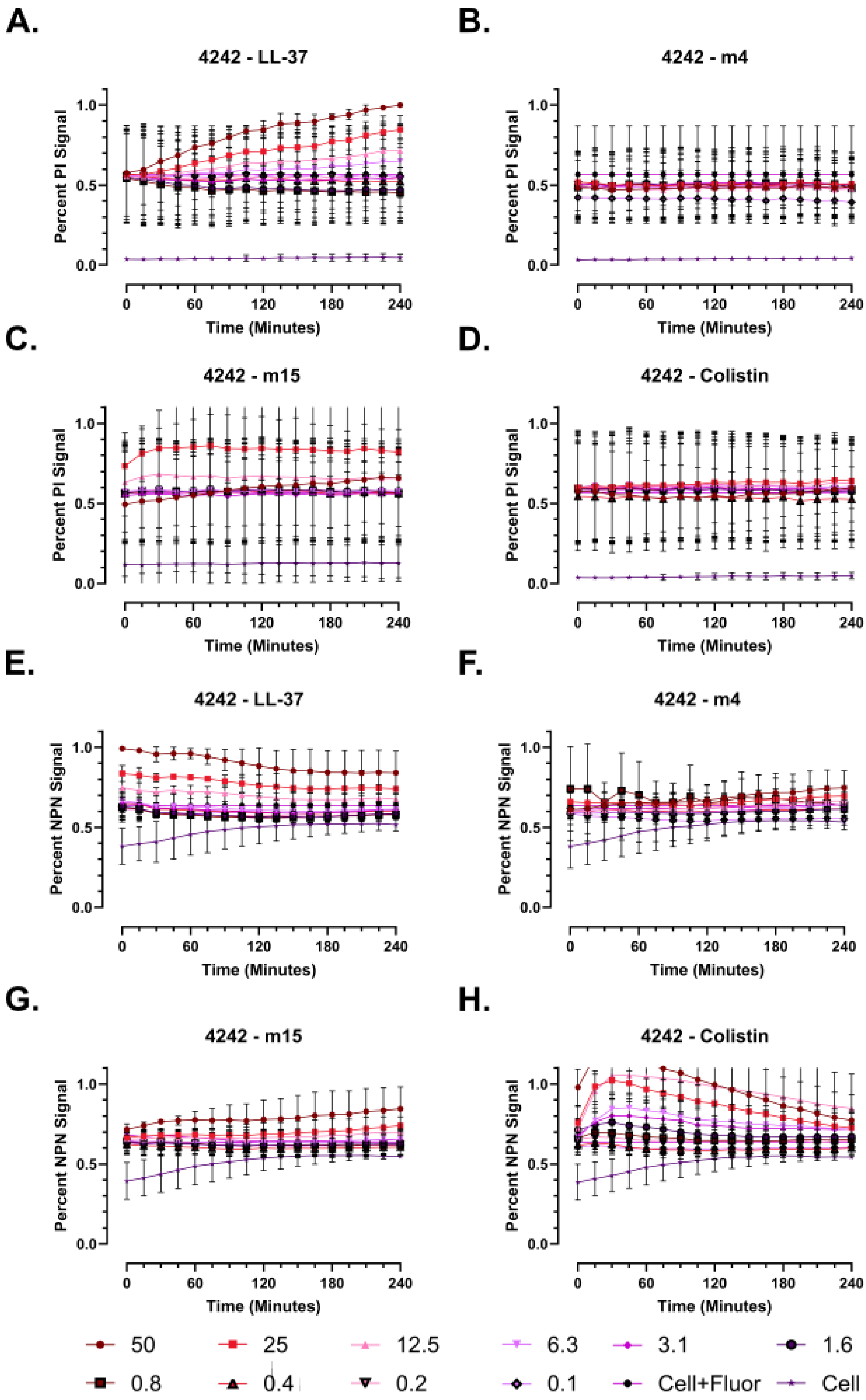
A.
4242 - LL-37
Percent PI Signal
Time (Minutes)
B.
4242 - m4
C.
4242 - m15
D.
4242 - Colistin
E.
4242 - LL-37
Percent NPN Signal
F.
4242 - m4
G.
4242 - m15
H.
4242 - Colistin
50
25
12.5
6.3
3.1
1.6
0.8
0.4
0.2
0.1
Cell+Fluor
Cell

**Supplementary Figure 131 – Membrane permeabilization patterns in clinical strain 4242.** Clinical *Pseudomonas* strain 4242 was used in dual membrane permeabilization assays as described in the main text. **A-D.** Inner membrane permeabilization, PI signal (Ex 535 / Em 617). **E-H.** Inner membrane permeabilization, NPN signal (Ex 350 / Em 420).

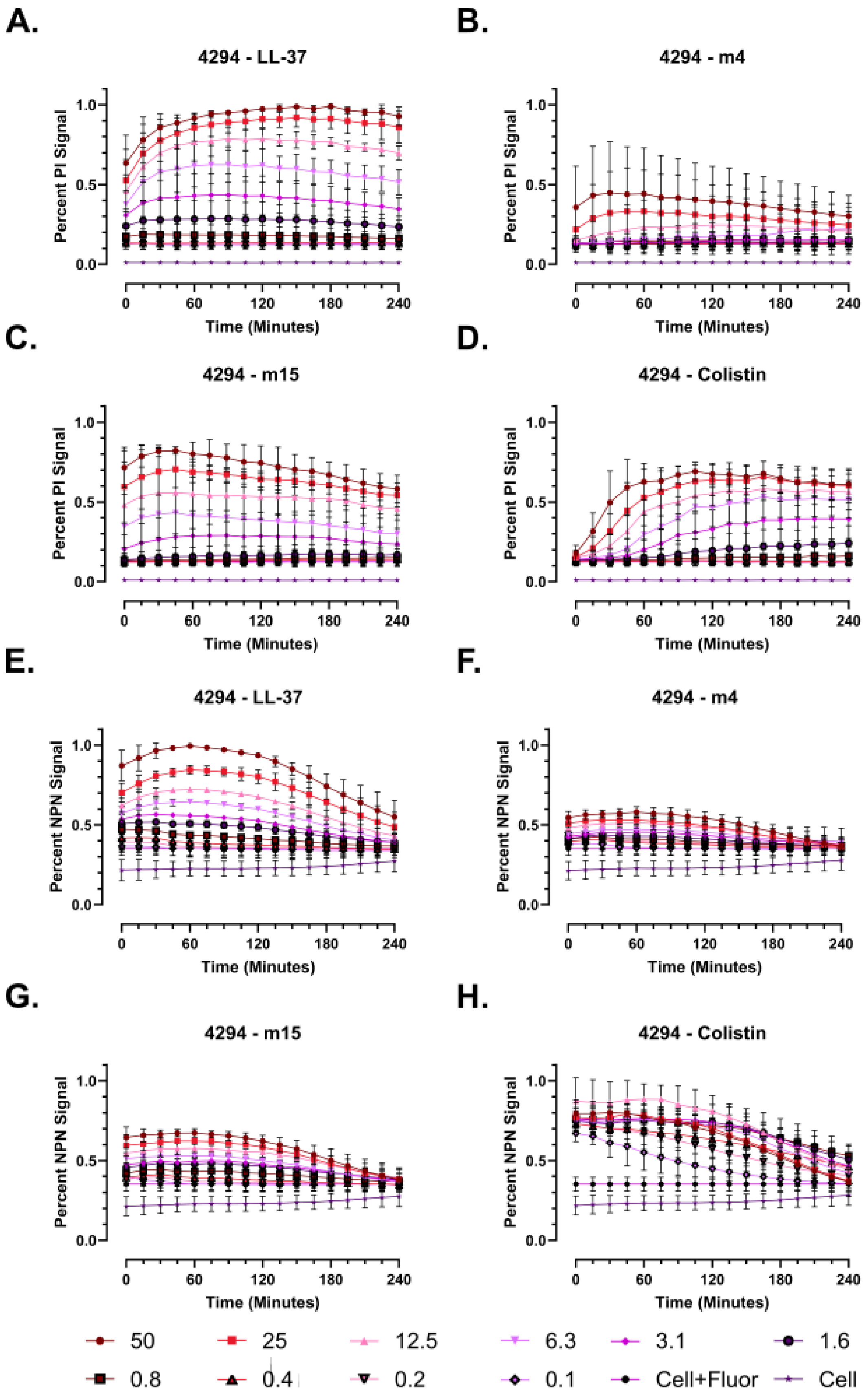
A.
4294 - LL-37
Percent PI Signal
Time (Minutes)
B.
4294 - m4
Percent PI Signal
Time (Minutes)
C.
4294 - m15
Percent PI Signal
Time (Minutes)
D.
4294 - Colistin
Percent PI Signal
Time (Minutes)
E.
4294 - LL-37
Percent NPN Signal
Time (Minutes)
F.
4294 - m4
Percent NPN Signal
Time (Minutes)
G.
4294 - m15
Percent NPN Signal
Time (Minutes)
H.
4294 - Colistin
Percent NPN Signal
Time (Minutes)
50
25
12.5
6.3
3.1
1.6
0.8
0.4
0.2
0.1
Cell+Fluor
Cell

**Supplementary Figure 132 – Membrane permeabilization patterns in clinical strain 4294.** Clinical *Pseudomonas* strain 4294 was used in dual membrane permeabilization assays as described in the main text. **A-D.** Inner membrane permeabilization, PI signal (Ex 535 / Em 617). **E-H.** Inner membrane permeabilization, NPN signal (Ex 350 / Em 420).

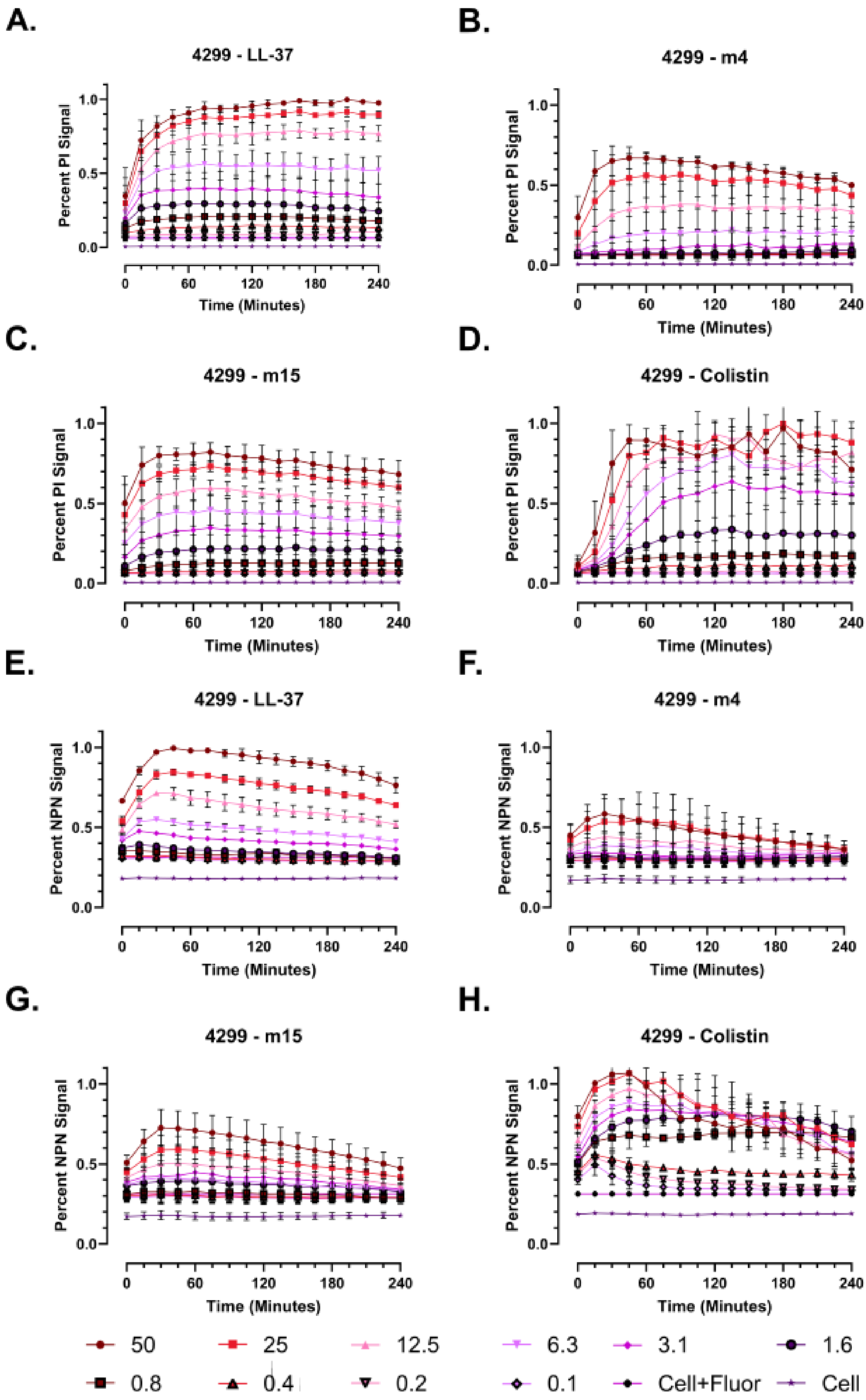

A.
4299 - LL-37
B.
4299 - m4
C.
4299 - m15
D.
4299 - Colistin
E.
4299 - LL-37
F.
4299 - m4
G.
4299 - m15
H.
4299 - Colistin
Percent PI Signal
Percent NPN Signal
Time (Minutes)
50
25
12.5
6.3
3.1
1.6
0.8
0.4
0.2
0.1
Cell+Fluor
Cell

**Supplementary Figure 133 – Membrane permeabilization patterns in clinical strain 4299.** Clinical *Pseudomonas* strain 4299 was used in dual membrane permeabilization assays as described in the main text. **A-D.** Inner membrane permeabilization, PI signal (Ex 535 / Em 617). **E-H.** Inner membrane permeabilization, NPN signal (Ex 350 / Em 420).

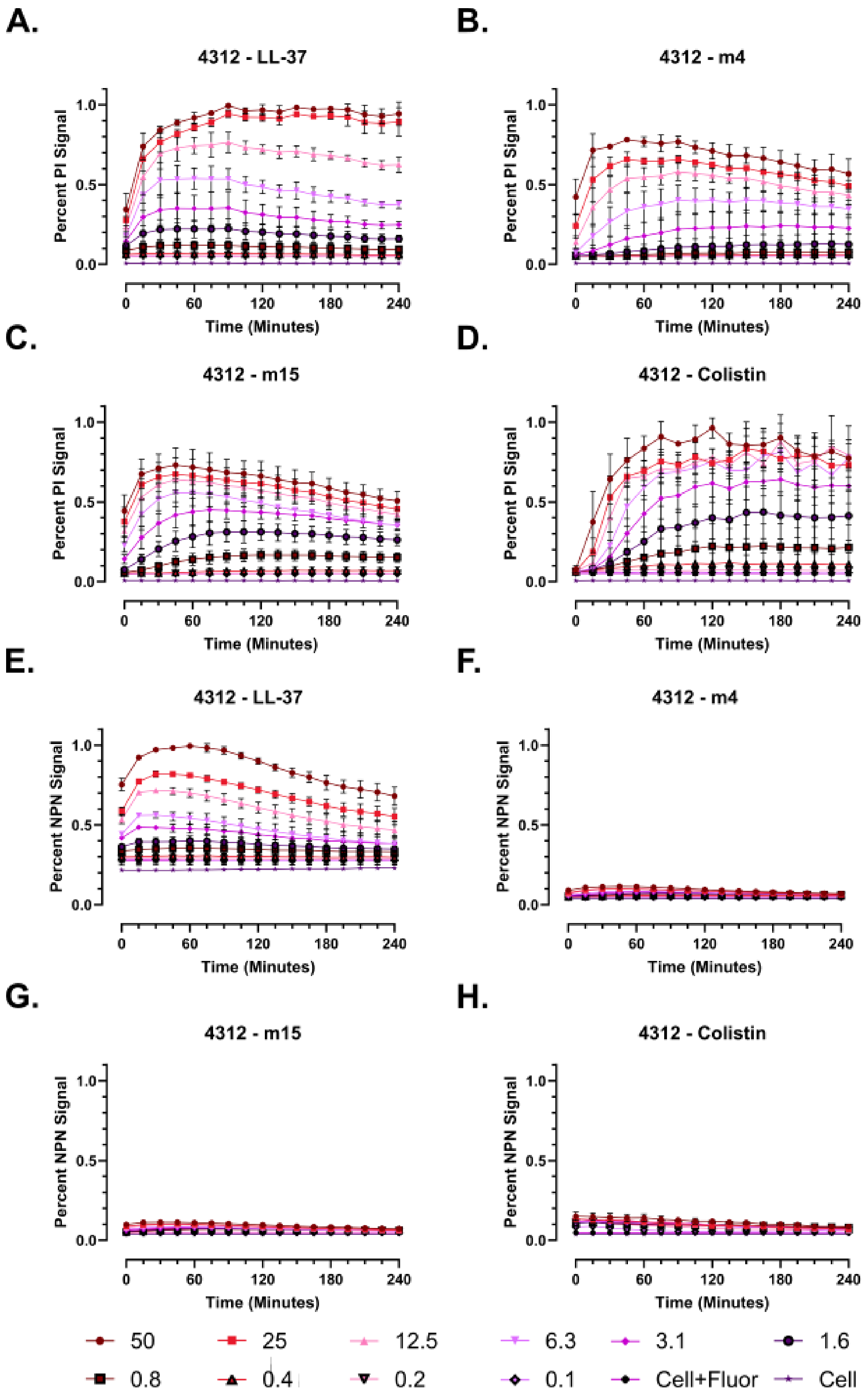
A.
4312 - LL-37
B.
4312 - m4
C.
4312 - m15
D.
4312 - Colistin
E.
4312 - LL-37
F.
4312 - m4
G.
4312 - m15
H.
4312 - Colistin
Percent PI Signal
Percent NPN Signal
Time (Minutes)
0.0
0.5
1.0
0
60
120
180
240
50
25
12.5
6.3
3.1
1.6
0.8
0.4
0.2
0.1
Cell+Fluor
Cell

**Supplementary Figure 134 – Membrane permeabilization patterns in clinical strain 4312.** Clinical *Pseudomonas* strain 4312 was used in dual membrane permeabilization assays as described in the main text. **A-D.** Inner membrane permeabilization, PI signal (Ex 535 / Em 617). **E-H.** Inner membrane permeabilization, NPN signal (Ex 350 / Em 420).

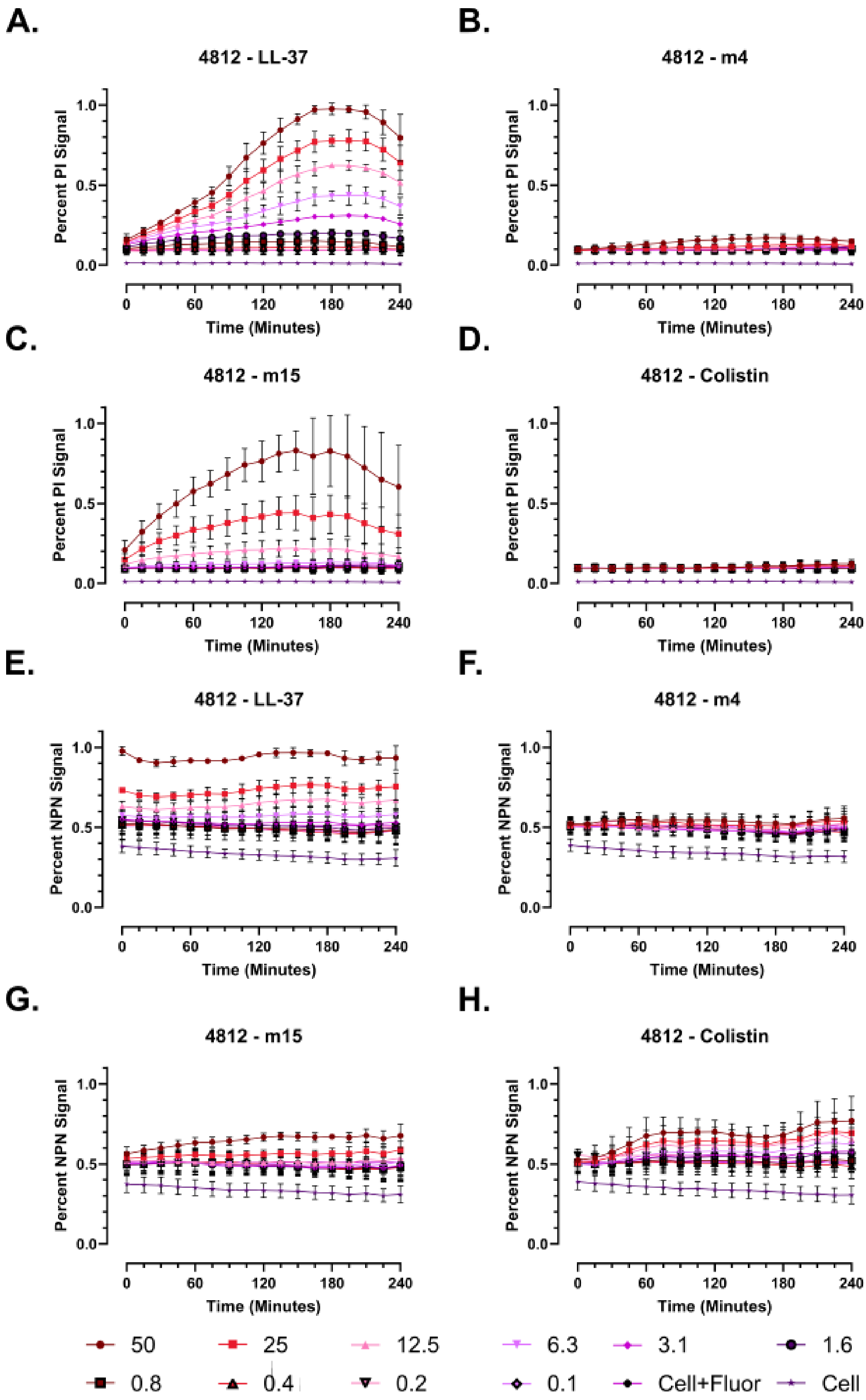
A.
4812 - LL-37
B.
4812 - m4
C.
4812 - m15
D.
4812 - Colistin
E.
4812 - LL-37
F.
4812 - m4
G.
4812 - m15
H.
4812 - Colistin
Percent PI Signal
Percent NPN Signal
Time (Minutes)
0.0
0.5
1.0
0
60
120
180
240
50
25
12.5
6.3
3.1
1.6
0.8
0.4
0.2
0.1
Cell+Fluor
Cell

**Supplementary Figure 135 – Membrane permeabilization patterns in clinical strain 4812.** Clinical *Pseudomonas* strain 4812 was used in dual membrane permeabilization assays as described in the main text. **A-D.** Inner membrane permeabilization, PI signal (Ex 535 / Em 617). **E-H.** Inner membrane permeabilization, NPN signal (Ex 350 / Em 420).
